# Biochemical propensity mapping for structural and functional anatomy of importin α IBB domain

**DOI:** 10.1101/2021.10.25.465662

**Authors:** Kazuya Jibiki, Mo-yan Liu, Chao-sen Lei, Takashi S. Kodama, Chojiro Kojima, Toshimichi Fujiwara, Noriko Yasuhara

## Abstract

Importin α has been described as a nuclear protein transport receptor that enables proteins synthesized in the cytoplasm to translocate into the nucleus. Besides its function in nuclear transport, an increasing number of studies have examined its non-nuclear transport functions. In both nuclear transport and non-nuclear transport, a functional domain called the IBB domain (importin β binding domain) plays a key role in regulating importin α behavior, and is a common interacting domain for multiple binding partners. However, it is not yet fully understood how the IBB domain interacts with multiple binding partners, which leads to the switching of importin α function. In this study, we have distinguished the location and propensities of amino acids important for each function of the importin α IBB domain by mapping the biochemical/physicochemical propensities of evolutionarily conserved amino acids of the IBB domain onto the structure associated with each function. We found important residues that are universally conserved for IBB functions across organisms and families, in addition to those previously known, as well as residues that are presumed to be responsible for the differences in complex-forming ability between families and for functional switching.

## Introduction

Importin α is a nuclear transport factor that mediates the translocation of nuclear proteins from the cytoplasm to the nucleus. There are three functional domains necessary for nuclear transport by importin α. The N-terminal domain known as the IBB domain (importin β binding domain) is necessary for the interaction with a partner transport factor, importin β (Weis et al., 1996). The main body is composed of 10 repeated structures called armadillo (ARM) repeats (Conti et al., 1998). This region also includes two recognition sites for the nuclear localization signal (NLS) of transport cargo proteins, called the major NLS binding site and the minor NLS binding site (Conti et al., 1998, Lange et al., 2007). The C-terminal part with ARM 9, 10 and unstructured region includes a binding site for Nup50 that stretches from ARM10 to ARM 4 (Matsuura & Stewart, 2005), which facilitates the release of the NLS from importin α. ARM 10 in the C-terminal also has a binding site for the specific export factor CAS/CSE1 (Cellular Apoptosis Susceptibility protein/Chromosome Segregation 1) (Kutay et al., 1997, Görlich et al., 1997, Hood & Silver, 1998, Solsbacher et al., 1998).

In the importin α dependent transport machinery, the importin α recognizes the NLS of the cargo proteins through major and/or minor NLS binding sites (Lange et al., 2007). Importin β is also recruited to the transport complex through binding to the IBB domain forming a ternary complex with importin α and the cargo (Stewart, 2007, Tran et al., 2014, Dickmanns et al., 2015). Like the cargo proteins, the IBB domain is rich in basic amino acids and can cover the NLS sites of importin α leading to autoinhibition. However, the association of importin β with the IBB prevents autoinhibition and exposes the NLS binding sites to facilitate the binding of NLS cargo to importin α (Harreman, Cohen, et al., 2003, Harreman, Hodel, et al., 2003). The ternary complex then translocates to the nucleus through the nuclear pore complex. The fates of importin α and the cargo proteins in the nucleus are determined by several interacting molecules. Cargo release is achieved by RanGTP binding to importin β that mediates the dissociation of importin β from the complex (Gilchrist et al., 2002, Floer et al., 1997, Vetter et al., 1999), and by Nup50 or CAS binding to importin α (Gilchrist et al., 2002, Sun et al., 2013). Moreover, very recently we reported that importin α-DNA binding can occur (with or without cargo), and part of the IBB acts as a subdomain named NAAT domain (Nuclear Acid Associating Trolley pole domain) (Jibiki et al., 2021). Finally, importin α is exported out of the nucleus through the export function of CAS and is recycled (Matsuura & Stewart, 2004).

Importin α forms a multi-gene family, and the number of genes varies depending on the organisms. There are organisms which has only one type of gene such as *Saccharomyces cerevisiae*, whereas humans have up to seven types of family genes. On the other hand, all family proteins have the typical domains described above and follow the typical cycle of nuclear transport. However, they have specific transport substrates and tissue expression (Pumroy & Cingolani, 2015, Köhler et al., 1999), which provide a selective system for the transport of nuclear proteins. The seven family proteins are further divided into three subtypes based on amino acid homology. Homology between subtypes in humans is around 50% (Oka & Yoneda, 2018).

In the above transport process, the IBB domain regulates functions of importin α, such as autoinhibition, importin β binding, DNA binding, CAS binding to form the nuclear export complex, and through the replacement of interacting molecules which causes a switch in the function of importin α (Lott & Cingolani, 2011).

Although the tertiary structures of the IBB complexed with importin β for nuclear translocation, complexed with CAS and Ran for nuclear export, and the autoinhibition form have been reported, the conformations of IBB in each of them are in different states, indicating structural polymorphism. Such stretches of the same amino acid sequence in different conformations are called chameleon sequences (ChSeqs) (Minor & Kim, 1996). Indeed, the IBB domain is shown as a region with low or very low confidence in the structural prediction by AlfaFold for any family member of any organisms, even though the presence of helices was predicted (e.g., P52292, human KPNA2). As intrinsically disordered proteins/regions (IDPs/IDRs) have been shown as regions with a low or very low confidence level in the structure prediction by AlfaFold (Ruff & Pappu, 2021), this implies that IBB has IDR-like propensities and is highly polymorphic with a chameleon sequence.

Thus, the IBB domain of the importin α family is multifaceted in terms of both structure and function. However, it is not yet fully understood how a single IBB domain interacts with multiple binding partners, leading to the distinction and switching of importin α function in certain situations. It has been reported that the function of import α via IBB varies among families and organisms (Lott & Cingolani, 2011, Pumroy & Cingolani, 2015, Miyamoto et al., 2016, Yasuhara & Yoneda, 2017, Oka & Yoneda, 2018), and some reports suggest that the amino acid sequence of IBB can be directly linked to differences in propensities among families (Zienkiewicz et al., 2013). However, a comprehensive understanding of the underlying mechanisms has not yet been achieved. Therefore, in this study, we focused our analysis on the IBB domain.

In this study, we aimed to distinguish the location and propensities of amino acids important for each function in the importin α IBB domain, a multifunctional ChSeq, by mapping the biochemical/physicochemical propensities of evolutionarily conserved amino acids to different conformations corresponding to each function. The result enabled discrimination and scrutiny of the contribution of each residue to the multiple functions. As a result, we have revealed several previously unknown propensities of the residues that are important for complex formation related to each function.

## Result and Discussion

### Generation of importin α family protein consensus sequences

The IBB domain is localized in the N-terminal region before the ARM repeats in the typical importin α protein (Figure 1A). Interactions with binding partners are mediated by the IBB domain and ARM repeats (Figure 1B), thus the existence of these two domains is one of the basic characteristics of importin α family proteins. In this study, the importin α protein is indicated by gene name KPNA1 (importin α1, NPI1, importin α5 in humans), KPNA2 (importin α2, Rch1, and importin α1 in humans), KPNA3 (importin α3, Qip2, and importin α4 in humans), KPNA4 (importin α4, Qip1, and importin α3 in humans), KPNA6 (importin α6, NPI2, and importin α7 in humans), and KPNA7 (importin α8), and subtypes including KPNA1, KPNA5, and KPNA6 are referred to as the subtype 1, KPNA3 and KPNA4 as the subtype 3, and KPNA2 and KPNA7 as the subtype 2.

**FIGURE 1.**
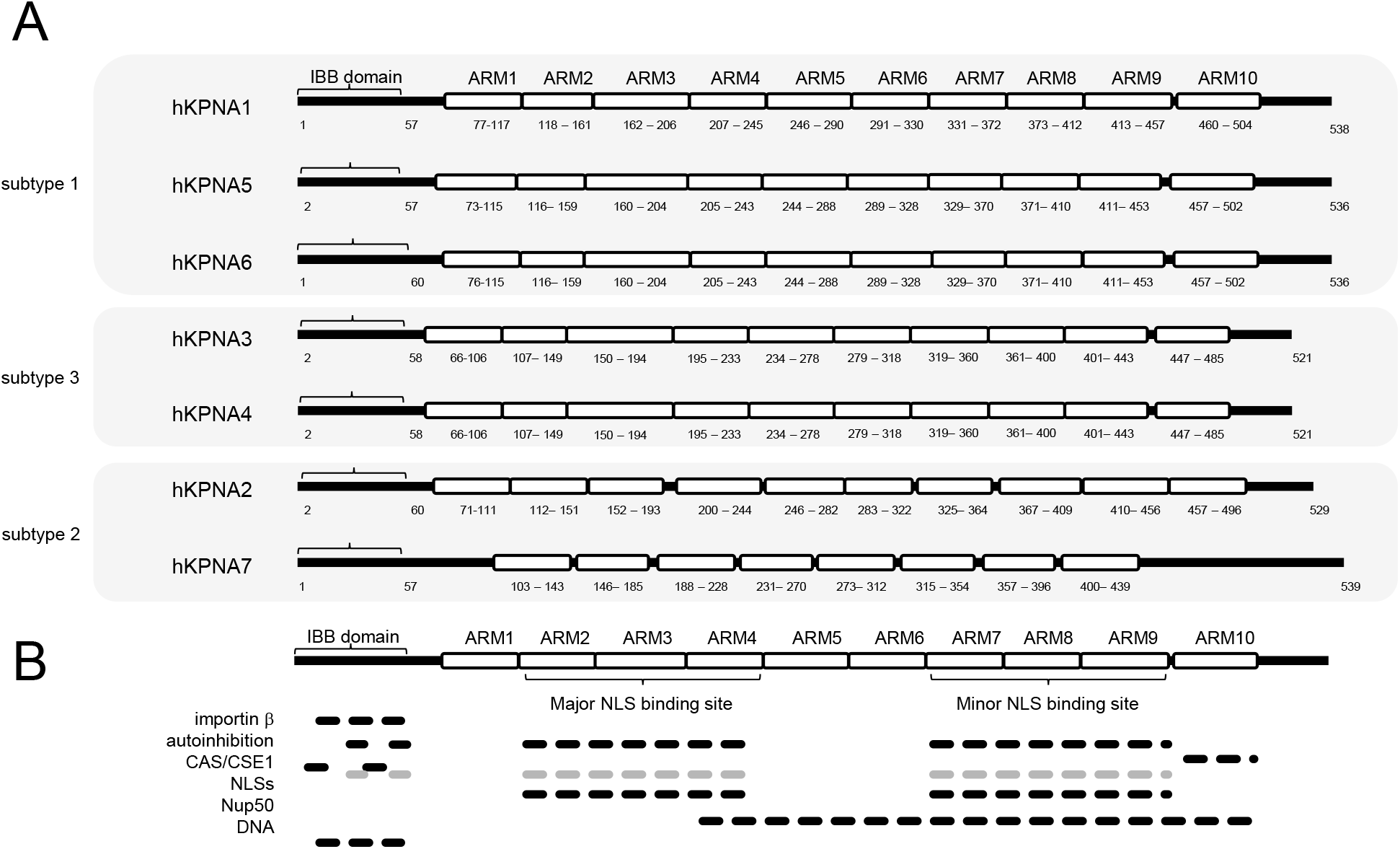
Schematic diagram of the domain architecture and interaction sites of the importin α family of human. (A) The families are grouped and arranged by subtype. The region of each domain is indicated by the position of the amino acids at the N- and C-termini of the domain. (B) Sites involved in intermolecular/interdomain interactions are shown by black bars for importin β binding, autoinhibition form formation, CAS/CSE1 complex formation, NLSs binding, Nup50 binding, and DNA association. Interaction sites between IBB and ARM repeats in IBB/ CAS/CSE1 complexes are represented by gray bars. The region of the major and the minor NLS binding sites are also depicted.

We collected multiple IBB domain sequences of diverse organisms to compare the amino acid conservation and create consensus sequences that can be used as unified sequences for each family (Figure S1, see also Supporting file 1). The collected sequence sets were subjected to multiple alignment, and the IBB domain and the full-length consensus sequence composed of the most conserved amino acids at each position were generated for each family member for the subsequent analysis, see Supporting file 2 for IBB domain. We named the consensus sequences of IBB domain or full-length for each importin α family member as cIBB1–7 and cKPNA1–7, respectively (Figure 2A, see also Supporting file 3). Overall, mammalian was the dominant organisms in the sequence set, but actinopterygii, which is comprise over 50% of vertebrate organisms, was the most abundant organisms in the KPNA2 sequence set. For KPNA2 IBB, when the consensus sequence for each class was used, the identity between the actinopterygii consensus sequence and the mammalian consensus sequence was 73% (data not shown). However, the sequence set also included many sequences of reptiles which showed a significant homology to those of mammalian compared to actinopterygii and eventually the identity between the consensus sequence cIBB2 determined for all organisms and that of mammalian sequences was 100% (data not shown). Thus, it confirmed that differences in the distribution of organisms in the sequence set did not create a cIBB2-specific bias.

**FIGURE 2.**
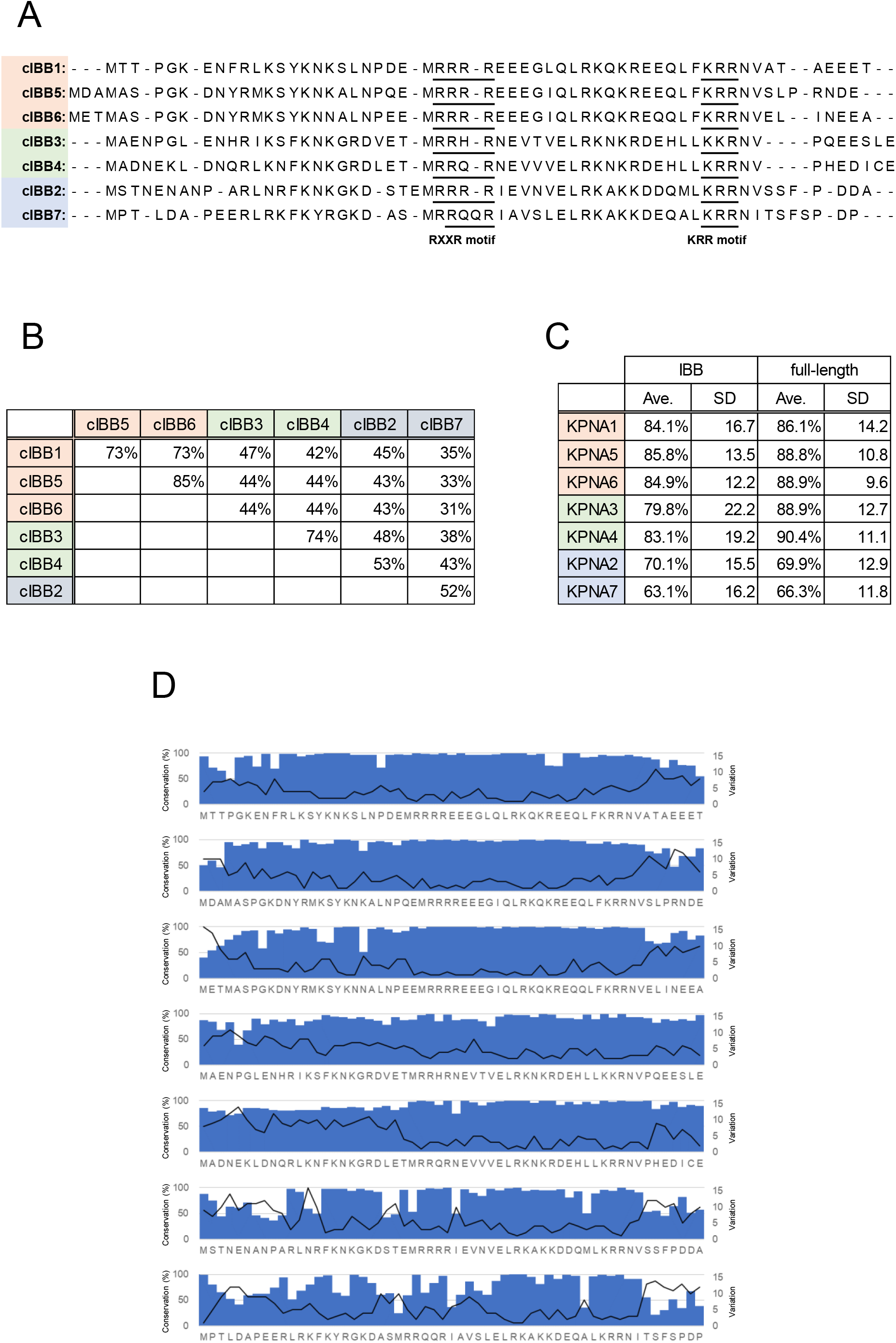
Conservation of the IBB domain residues among organisms. (A) The multiple alignment of IBB domain consensus sequences of importin α family. RXXR motif and KRR or KKR are indicated by underlines. (B) The percent identity scores in the pairwise alignment between each IBB domain consensus sequence. (C) The average percent identity scores of the pairwise alignments and standard deviation (SD) among IBB domain or full-length sequences included in the sequence set for each importin α family member. (D) The degree of the conservation of each amino acid of the consensus sequence (bar graphs) and the number of amino acid types that appeared at each position (line graphs), among the organisms in the sequence sets for each importin α family member.

### Amino acid variations in the IBB domain among organisms

To classify the amino acid sequences, the IBB domains and the full-length consensus sequences were then subjected to multiple alignments to produce phylogenetic trees (Figure 2A, S2, S3, see also Supporting file 3). In the phylogenetic trees of the full-length consensus sequences obtained by either the NJ method or the maximum likelihood method, all families were classified in the same way with the previously described canonical subtype classification that have been reported for human importin α (Figure S3) (Pumroy & Cingolani, 2015, Kelley et al., 2010, Miyamoto et al., 2016). This supports the fact that this full-length consensus sequence retains the same propensities as the conventional general classification. On the other hand, the phylogenetic trees of the IBB domain consensus sequences obtained by the NJ method and the maximum likelihood method had different branches (Figure S2). Taken together, this suggests that features that can be obtained explicitly from the sequence are not sufficient to characterize the IBB domain in relation to its function.

Other evidence suggests that the consensus sequences obtained in this study are a good representative of the nature of each family of importin α. Among the consensus sequence of the IBB domain generated in this study, cIBB7 showed low homology to other family members and was highly diverse in the amino acid substitution among the organisms (Figure 2B, 2C). The propensity was consistent with a previous study on the IBB domain of human KPNA7 and this also supported the hypothesis that the KPNA7 sequence may have evolved under various types of selection pressures.

We focused on the conservation and variation of individual residues between organisms. The degree of preservation of each amino acid in the consensus sequence and the diversity of the amino acid appearing at each position were calculated (Figure 2D), see also Supporting file 4. It has previously been established that the best-preserved features of the IBB domain are basic patches RXXR and KRR or KKR, corresponding to residues indicated by the underline in cIBB indicated in Figure 2A, which occupy their own minor NLS binding site and major NLS binding site, respectively, in the nuclear export complex with CAS and RanGTP, and in the autoinhibition from of importin α (Figure 1B) (Matsuura & Stewart, 2004, Kobe, 1999, Chang et al., 2013). All IBB domain consensus sequences generated in this study contained both the basic patches and their constituent arginine or lysine was found to be conserved at least 95.7% in all families (Supporting file 4).

### The interface of the IBB domain for each binding partner

We built model complex structures of three different functional complexes, the importin β binding form, a nuclear export complex with CAS/CSE1, and Ran-GTP, using the IBB consensus sequences cIBB1–7. Since the redundant consensus sequences of the non-IBB portion for cKPNA2 and cKPNA7 prevented us from using the consensus sequences for the modeling of the nuclear export complex and the autoinhibition form, we decided to use chimeric sequences prepared by replacing the IBB domain portion of the full-length human importin α family sequences with the corresponding IBB domain consensus sequence (Supporting file 5). The residues in the non-IBB portion of this chimeric sequence that showed the top 30% of absolute values of the total energy of nonBonded and electrostatic interactions were more than 84.7% identical to the consensus sequence of any family. Therefore, we concluded that there is no significant impact of using this chimeric sequence for the analysis of molecular interactions (data not shown). Note, however, that despite the large proportion of mammal in the sequence set, a small number of residues differ between the human and consensus sequences in places. For all modeled structures it was confirmed that there was no distortion or steric hindrance in the interface of the generated complex model by computing the energy with GROMOS96 43B1 force field using Swiss-Pdb Viewer 4.1.0. The GMQEs (Global Model Quality Estimates) for the model and the RMSD of all atoms between the model and the template are shown in Figure S4-S9.

The contact area of each residue of the IBB domain to the binding partner protein in the complexes was calculated to reveal the contribution of each residue to the binding (Figure S10). The total interface areas for the three structures are shown in Figure 3A. The estimated total interface area for each structure did not differ significantly among the family members, suggesting that the overall degree of interaction with the binding partners was conserved among the IBB domains of all family members (Figure 3A). In general, the standard area of the protein-protein interaction surface of a stable complex is 1600 (±400) Å2 (Conte et al., 1999). By this criterion, the interaction surface between IBB and importin β reflects a relatively stable complex (Figure 3A-I). Although the interaction between the IBB and CAS in the nuclear export complex (Figure 3A-IIa), and the interaction with ARM (Figure 3A-IIb), have both standard area size of interaction surfaces, considering that these interactions occur cooperatively, the net interface area is significantly large (Figure 3A-II), and the entire export complex would have sufficient stability. For the autoinhibition form, the area is rather small when only the visible part of the crystal structure (PDB ID:1IAL) is considered (Figure 3A-III), suggesting that this form is transient and unstable, or that the missing part of the crystal structure contributes to the interaction. For each model structure, we calculated the free energy changes of nonBonded interaction, electrostatic interaction, and solvation for each residue due to the formation of complex interface (Figure S11–13). The results showed a high correlation between the contact area and the sum of the energies of the nonBonded and electrostatic interactions (Figure S14). This is consistent with the fact that there is a correlation between the interface area and the strength of the binding (Henrick & Thornton, 1998). An exceptionally low correlation coefficient was obtained for the nuclear export complex of cIBB4, but this was thought to be due to the strong repulsion of charge caused by truncation of the N-terminal residue during modeling. Therefore, we excluded this residue and recalculated the correlation coefficients for all interactions. The GRSOMOS energy calculation does not include the hydration term, but it is generally difficult to estimate the free energy change due to solvation/desolvation, especially the absolute value of the entropy term. However, as mentioned above, there is a strong correlation between the contact area and nonBonded and electrostatic interactions, and there is a natural correlation between the change in hydration and the change in molecular surface area. Therefore, we decided to analyze the contribution of each residue to the interaction based on the interface area.

**FIGURE 3.**
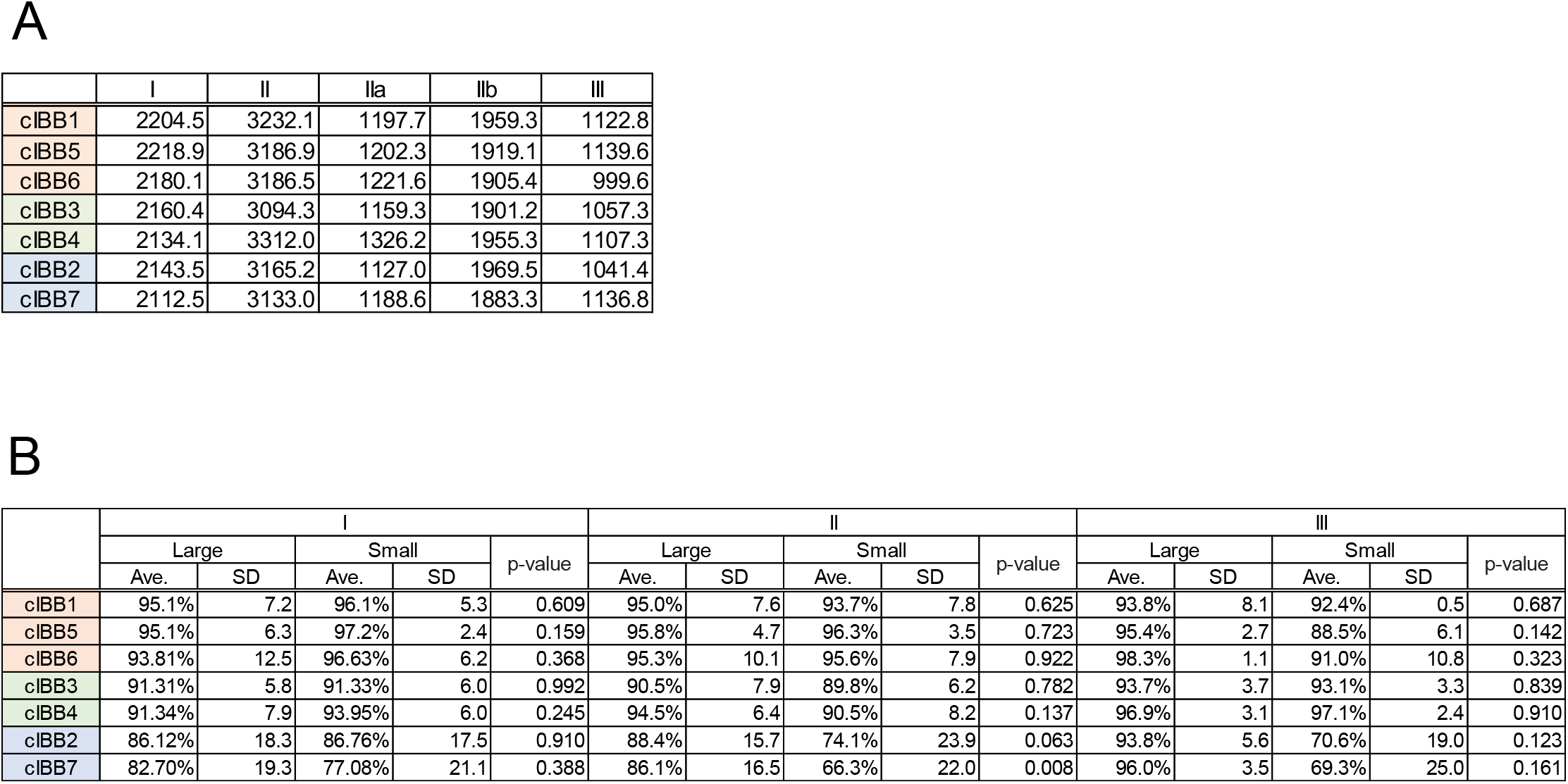
Interface in the functional complex between IBB domain and interacting molecules. (A) The total contact area between (I) IBB and importin β, (II) IBB and CAS/ARM repeat, (IIa) IBB and CAS, (IIb) IBB and ARM repeat in the nuclear export complex, and (III) IBB and ARM repeat in the autoinhibition formed by importin α alone. The contact area of residues 13–54 for I, 8–16 and 26–53 for II, and 44–54 for III were summed. (B) The average degrees of conservation over all residues at the position where at least one family had a relatively large contact area at the interface (Large) or all residues at the position where none of the family had a relatively large contact area at the interface (Small). See method for criteria to determine whether the contact area at the interface is relatively large or not. Standard deviation (SD) and p-values were also shown.

The relationship between the conservation of amino acids at each position and the contact area of the residue in each structure was evaluated (Figure 3B). For all three structures for each family, residues that showed a relatively large contact area at the interface also had highly averaged conservation level, suggesting the existence of strong selection pressures due to the maintenance of the interaction (Figure 3B-Large). On the other hand, the residues that showed a relatively small contact area at the interface in each complex also showed a relatively high average degree of conservation (Figure 3B-Small). This can be interpreted in the context of the multifunctionality of the IBB domain. Residues that are not important for interaction with one molecule may be important for interaction with another molecule and vice versa. Many residues will be important for at least one of the many different functions that IBB performs, and selection pressure will ensure that they are evolutionarily conserved. Among all family members, subtype 2 composed of KPNA2 and KPNA7 showed a relatively low overall amino acid conservation compared to other families (Figure 3B-Small, Supporting file 4). This likely reflects the fact that they have evolved to be specialized with a limited number of functions. Interestingly, we found residues that showed a very high degree of conservation, even though they contributed very little to the interface in any of the complexes. This implies that they are required for a function other than that of the three complexes discussed here.

### The mode of conservation of biochemical/physicochemical propensities differs from that of the amino acid sequence itself

To investigate the biochemical/physicochemical propensities of the residues of the IBB domain consensus sequence that are important for the interaction for each structure, we mapped the biochemical/biophysical propensities to each residue of the IBB domain consensus sequences. The residues of each consensus sequence were scored based on the score index for α-helix preference, beta-sheet preference, beta-turn preference, coil preference, bulkiness, polarity, hydrophobicity, and average flexibility (Figure S15). The propensity heat maps of IBB consensus sequences, average scores and standard deviations among organisms are shown in Figure S16–18. To examine the similarity of the IBB domains of the seven families by biochemical/physicochemical propensities, the cIBBs were clustered using the score of each propensity (Figure S19). The results showed that all families were classified by canonical subtypes, except for α-helix, β-sheet, and helix breaker. This indicates the fact that several biochemical/physicochemical propensities are conserved for the subtype IBB domains.

### Characteristics of the IBB interface in the importin β binding structure

The IBB domain in complex with importin β forms a structure consisting of a 310-helix and an approximate 30-residue α-helix connected by a short loop (Cingolani et al., 1999). The modeled structure of the consensus sequence shows a 310-helix at positions 14–16 and an α-helix at positions 24-51 (Figure S10). The tendency index to form α-helices was found to be relatively high at positions 24-49 in all the IBB domain consensus sequences (Figure S16). This tendency was also seen in the average score among all organisms (Figure S17). The relatively low coil formation propensity index between positions 26-49 in both the consensus sequence score and the interorganisms average score, and the relatively high index at positions outside these positions suggests the presence of evolutionary selection pressure for the maintenance of such secondary structures (Figure S16, S17).

Although the crystal structure used as a template for homology modeling (1QGK) has a clear helix structure, this part has an extended coil-like conformation in the crystal structure of the nuclear export complex (PDB:1WA5) and is frequently reported as a missing part in the crystal structure of the autoinhibition form on its monomer (PDB:1IAL and others). As is evident from the Ramachandran plots of the IBB region in these structures shown in Figure S20, the dihedral angles of many of the peptide main chains in the IBB region switch dramatically between their respective functional structures. Considering these facts together with the results of the present analysis, it seems that this region of IBB has an evolutionarily conserved chameleon-like nature as a multifunctional ChSeq that can change into multiple conformations, including the α-helix structure, when necessary, through the induced fitting with specific binding partners.

For the consensus sequences of the IBB the residues at the interface of the interaction with importin β were examined for common and differential propensities among the family members (Figure S21). Briefly, the residues R13, K18, R28, R31, R39, K40, and R51 which contact with importin β were common in all the consensus sequences and are highly conserved (Figure 4A, 4D-I, S21). Near those residues and residues at positions 14, 17, 43, 50, and 53, all the corresponding residues on the binding partner importin β side were also placed in the model structures as in the template structure 1QGK (Figure S21). Even in the case in which the type of amino acid varied at several positions among the family, the contact target residues on the surface of importin β were common and the estimated total interaction surface area was almost conserved (Figure 3A, 4A, 4D-I, S21). Even when the amino acid identity among the consensus sequences is low, the biochemical/physicochemical propensities of the amino acids are commonly conserved at the contact sites with importin β, e.g., hydrophobicity at positions 14, 17, and 53 and basicity at positions 43 and 50. However, in cIBB7, the hydrophobicity at position 14 fluctuated among organisms as methionine was quite abundant (Figure S16–18, see also Supporting file 4). These indicates that the interface regions among the family proteins and among the various organisms maintain certain common propensities even if the amino acids themselves are not conserved. Also, selective pressure on the type of amino acid at each position is diversified in a way that is specific to various biochemical/physicochemical propensities. This perspective seemed to be indispensable when considering the complex-forming capacity of IBB in relation to its function.

**FIGURE 4.**
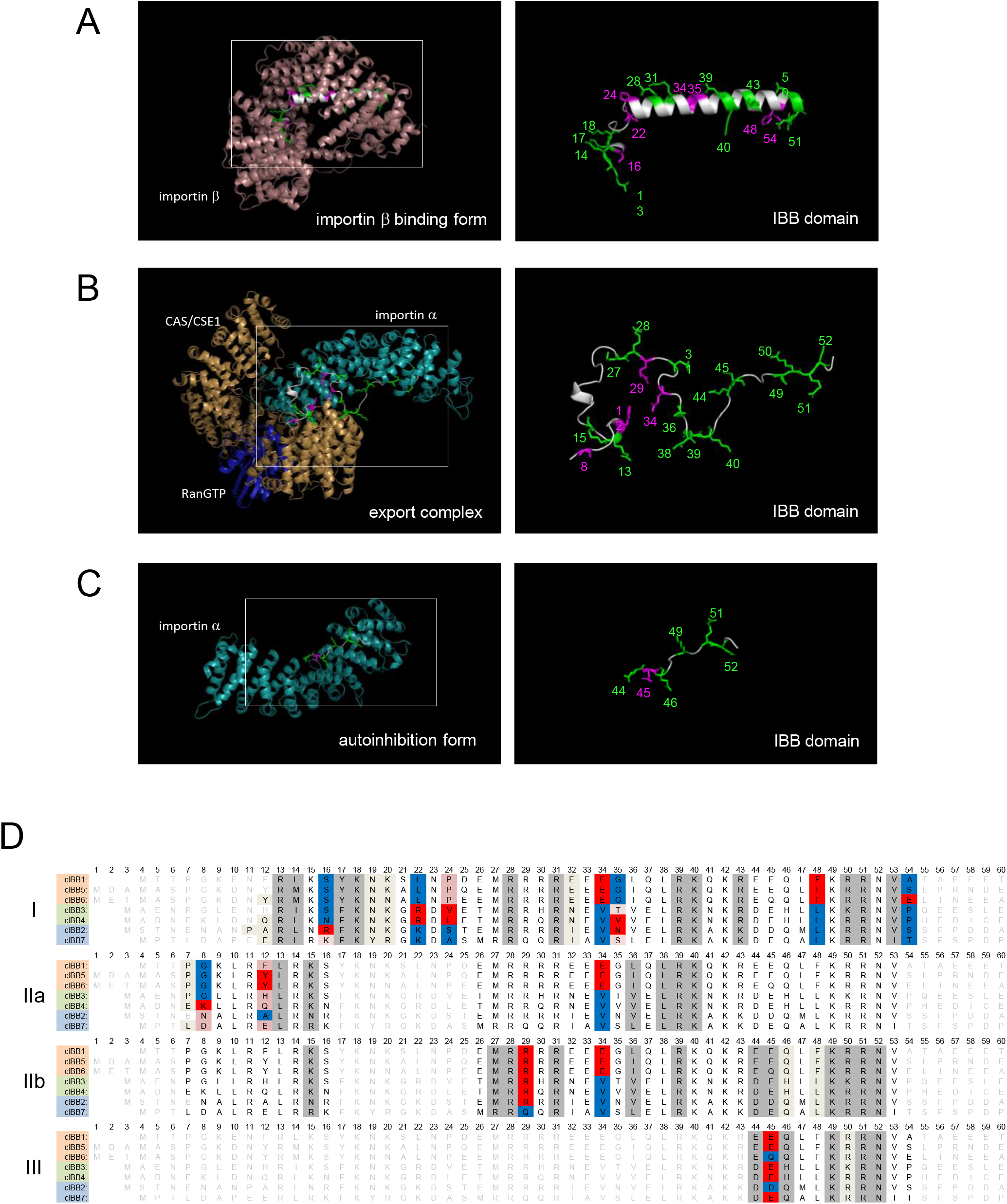
Characterization of the interface for each functional complex structure and the origin of family specificity. Residues at important common positions among the family (green) and those at a position defined as a specifically important positions for some family members (magenta) were mapped onto the cIBB of KPNA1 in model structures of the importin β binding form (A), the nuclear export complex (B), and the autoinhibition form (C). See method for criteria to determine whether the position is commonly important or specifically important. The expanded views of the IBB domain are shown on the right-side panels. Sidechains of the important residues are shown in a stick model and the residue numbers are shown. (D) Important residues for multifunctionality and functional switching of the IBB domain. Residues are shown for (I) the interface between IBB and importin β, (IIa) IBB and CAS, (IIb) IBB and ARM repeat in the nuclear export complex, and (III) IBB and ARM repeat in the autoinhibition formed by importin α alone according to the following rules: Gray: Positions where the residues commonly had the same large contact area among family. Red, light brown and blue: Positions where at least one family had large contact area and difference among family was large. The residue that had a relatively large contact area, intermediate area, and small area are colored with red, light brown, and blue, respectively. Ivory: Positions where some residues have a relatively large interface, but the difference among family is small. White: Positions where none of the family had relatively large contact area. If the residue was in a missing section of the template structure the letters are colored in pale gray. See also the materials and methods for detailed information on the selected residues.

As mentioned above, the fundamental propensities of the IBB for interaction with importin β are preserved not by conservation of the amino acid sequence itself, but by the conservation of biochemical/physicochemical propensities at the position of each residue. On the other hand, the presence of residues that may be responsible for the subtle differences between the families with respect to their ability to bind importin β was also revealed (Figure 4A, 4D-I, S21). At position 43, only cIBB2 and cIBB7, which belong to subtype 2, have lysine instead of arginine, suggesting that the contact area with importin β was relatively small compared to other family members. The residues on the importin β side with which they interact were different from those of other subtype members (Figure S10-I, S21). In addition, the presence of basicity or arginine rather than lysine at positions 16 and 22 could also give subtype 2 and subtype 3 a unique interaction with importin β. These predictions are consistent with the calculated energies of electrostatic interaction. However, the conservation of the basicity in KPNA2 at position 16 was low at 57.2% (Figure S11, S17). At positions 24 and 35, it appeared that the bulkiness of the amino acids simply characterized the contact area in each family, but in terms of diversity, the bulkiness score was more variable among the organisms for residue 24 than that for residue 35 (Figure S10-I, S16–18). Thus, this feature appeared to be more important for residue 35 than for residue 24.

In the model structures for all consensus sequences, no matter what amino acid is at position 34, there is a basic amino acid in its vicinity on the side of the importin β (Figure S21). In the IBBs in KPNA1, KPNA5, and KPNA6, which belong to subtype 1, an acidic amino acid was placed here with strong interorganisms conservation (Figure S17). At position 34, the basic amino acid of importin β was present in the vicinity for all consensus sequences. In the consensus sequence where glutamic acid was located at that position, R593 of importin β was present in the vicinity (Figure S21). The placement of acidic amino acids at this position probably favors the formation of an interface area due to charge compatibility. Similarly, at position 54, the acidic glutamic acid appears to characterize the interaction capacity with ARM. For example, interorganisms conservation of E54 in cIBB5 was only 78.0%, but the acidic nature is conserved at a much higher rate for KPNA5 (Figure S17). Residue 48 is phenylalanine in subtype 1, cIBB1, cIBB5, and cIBB6, and leucine in the others, and there is a considerable difference in contact area in the model structures (Figure S10-I). As the degree of interorganisms conservation in each family is high (>88.8%), it seems likely that this residue also evolved in relation to the regulation of the interactions (Supporting file 4). Thus, it is likely that residues 22, 34, 35, 48, 54, and in some organisms, 16 and 24, cooperatively give the families their individuality in the importin β binding capacity of IBB.

As for importin β binding, it has been reported that the substitution of 34RRRR (corresponding to 28RXXR in Figure S10) and 45RKAKR (corresponding to 39RKXKR(K) in Figure S10) of budding yeast SRP1 with alanine decreases the affinity for importin β (Harreman, Hodel, et al., 2003). The results of the present analysis show that R39 and R43 (K43) for this RXXR motif (28–31) and RKXKR/K (39-43) of IBB have a large contact area for importin β binding in all families. For the central section of the RKXKR/K motif, K40 to K42, the contact area was not very large for all families (Figure S10-I). Therefore, alanine substitutions of the RXXR and RKXKR(K) motifs, especially when the leading and trailing Rs are substituted, are likely to have a significant effect on importin β binding.

### Characteristics of the IBB interface in the nuclear export complex structure

In the importin α export complex, the IBB domain is known to bind to ARM repeats of importin α itself at two binding sites and simultaneously interact with CAS/CSE1 at additional two binding sites (Figure 1B) (Matsuura & Stewart, 2004). When the IBB is attached to the surface of the ARM repeat of importin α itself, the basic amino acids RXXR and KR(K)R in the IBB contact in a similar manner as the NLS of cargo proteins. For the contact area in the nuclear export complex the amino acid identity of each residue and the degree of conservation between the various organisms are shown in Figure S22 together with a biochemical/physicochemical trend heat map (Figure S16). The residues which had a large contact area were found at positions 13, 15, 27, 28, 31, 36, 38, 39, 40, 44, 45, 49, 50, 51, and 52 (Figure 4B, 4D-IIa, -IIb, S22). At positions 13, 28, 31, 38, 39, 40, 49, 51, and 52, residues R, R, R, L, R, K, K, R, and N were found, respectively, and were common among all IBB domain consensus sequences.

For all families, positions 15, 27, 36, 44, 45, and 50 existed in the interface, but the amino acids differed among the consensus sequences. However, in terms of biochemical/physicochemical propensities, there was considerable commonality. At position 15 all consensus sequences appeared to be more than moderately hydrophilic (Figure S16). At position 36 all consensus sequences have high scores for hydrophobicity and bulkiness and the interorganisms fluctuation was small, although the interspecific conservation of consensus sequences is only 60.4–97.3% (Figure S16–18, see also Supporting file 4). At position 44 all consensus sequences were acidic amino acids. Although the interorganisms conservation of the residue in each consensus sequence is only 74.5–99.3%, acidity was highly conserved (Figure S17, see also Supporting file 4). Residue 45 was moderately or highly hydrophilic with a similar contact area over all family members. In the family with glutamic acid and aspartic acid at this position, the formation of interfacial regions may be enhanced by electrostatic interactions, since in the model structures of all consensus sequences the arginine of its own ARM repeat is present near the residue at this position, while the difference did not seem to have a significant effect on the size of the contact area. The residue at this position showed more than 81.3% conservation among the organisms for each family (Figure S16, S22, see also Supporting file 4). At position 50, all the consensus sequences had basic amino acids, as described in the section of the importin β binding form, and basicity appeared to be conserved among many of the organisms in the sequence set (Figure S17).

The positions of 8, 12, 29, and 34 had a different contact area in the interface of the IBB-importin β complex among the families (Figure 4B, 4D-IIa, -IIb, S22). In position 8, asparagine and aspartic acid in cIBB2 and cIBB7, which belong to subtype 2, had twice the contact area of glycine in cIBB1, cIBB5, IBB6, and cIBB3. Furthermore, lysine in cIBB4 had twice the contact area (Figure S10-II). For this residue in cIBB4, the contact area is large, but it is unfavorable in terms of the free energy changes of nonBonded and electrostatic interaction, which suggests that the energy for hydration has a distinct characteristic (Figure S12). The presence of glutamic acid in CAS in the vicinity of this residue in the model complex structures of all the consensus sequences suggests that the positive charge may act through an electrostatic interaction (Figure S22). Position 29 was involved in the interaction with the minor NLS-binding site. The presence of glutamic acid in the ARM repeat in the vicinity of R29 in cIBB1 -6 suggests that basic amino acids were favorable for the interface. The interorganisms conservation of arginine in KPNA1-6 at this position is high (98.7-100%), while the conservation of glutamine in KPNA7 is low (64.6%) (Figure S22, see also Supporting file 4). At position 34, the contact area of glutamic acid in cIBB1, cIBB5, and cIBB6 is higher than that of other consensus sequences (Figure S10-II). At this position, acidic amino acids appear to be favored because of the presence of lysine in both ARM repeat and CAS in the vicinity of the residue (Figure S22). Regarding CAS binding, it has been reported that the substitution of arginine corresponding to R39 of Figure S10 with acidic amino acids in human and yeast importin α reduced the affinity for CAS (Sun et al., 2013, Lange et al., 2020). This is consistent with the fact that R39 has a large contact area with CAS and is commonly conserved among all family members, as revealed by the present analysis (Figure S10-II, see also Supporting file 4).

### Characteristics of the IBB interface in the autoinhibition structure taken by importin α alone

In the autoinhibition form, the basic amino acids in the latter part of IBB bind to the major-NLS binding site of ARM (Kobe, 1999). Our analysis revealed that the residues at positions 44, 46, 49, 51, and 52 of IBB were in the interface between IBB and ARM in all consensus sequences (Figure 4C, 4D-III, S23). Furthermore, the residues were common among the consensus sequences at the position 49, 51, and 52. The residues 49 and 52 were also important in the interface with ARM repeat in the nuclear export complex, and residue 51 was important in both the interface with ARM, in the nuclear export complex, and with importin β. These residues were conserved 96.3-100%, 93.3-100%, and 90.6-98.7% among organisms for each importin α family member (Supporting file 4). Although the amino acid was not identical at positions 44 and 46 among the consensus sequences, acidic amino acids were arranged in all consensus sequences at position 44 as previously mentioned. Since basic amino acids were placed in the ARM repeat in the vicinity of the acidic residue of the IBB, this conservation appears to be for electrostatic interaction (Figure S23). For residue 46, the degrees of hydrophilicity and bulkiness was conserved 89.5-99.6% among organisms in each family (Figure S16, Supporting file 4). Residue 45 was in a slightly different situation for each family in the monomeric autoinhibition form compared to that in the protein export complexes (Figure S10-III). In this position, glutamic acid showed 81.3–100% conservation among organisms, suggesting that this amino acid was particularly favored in terms of interactions in interface formation (Supporting file 4).

The substitution of K54 and R55 (corresponding to K49 and R50 in the consensus sequence) with alanine in budding yeast SRP1 reduced autoinhibition, and this effect was particularly pronounced for the substitution of K54 (Harreman, Cohen, et al., 2003). This effect is significant even with arginine substitution, which has synonymous with basicity, suggesting that the interaction at this position is lysine specific. These findings agree with the expected propensities of the IBB consensus sequence in the autoinhibition form, which has a large contact area with the ARMs at K49 and R50 (Figure S10-III). Furthermore, the presence of a sequential large contact area from K49 to R52, which is common among the families, suggests that the interaction in this region is cooperative and highly selective. This also explains why the substitution of K54 in SRP1, corresponding to K49 in the consensus, to a similarly charged amino acid, R, significantly destabilizes the autoinhibition form.

It has been reported that KPNA4, which has the RXXR motif in humans as RRQR, has a weaker autoinhibition than KPNA2, which has the RRRR motif (Pumroy et al., 2015). In addition to human KPNA4, Plasmodium falciparum, Toxoplasma gondii, A. thaliana and human KPNA7, have been reported to lack RXXR motif and/or KRR motifs and might have weak autoinhibition (Dey & Patankar, 2018, Bhatti & Sullivan, 2005, Hübner et al., 1999, Oostdyk et al., 2019). These are consistent with the finding that these motifs in the consensus sequences are conserved among families and have a large contact area in the autoinhibition form (Figure S10-III, see also Supporting file 4).

In the nuclear export complex, the IBB domain is expected to occupy the same position as in the autoinhibition form on the surface of the ARM repeat, but the H and Q in the middle of the RRRR motifs of KPNA3, KPNA4, and KPNA7 are estimated to have a lower contact area than the R of the other family members from our study (Figure S10-IIb). When 29R replaces Q and 30R replaces H, the nonBonded interaction, electrostatic interaction, and hydration energy seem to be greatly affected, respectively and in KPNA7, the second Q seems to contribute more to the decrease in autoinhibition than the third Q (Figure S12). In fact, the report regarding the weak autoinhibition in human KPNA7, suggested that IBB adopts an open state free from the ARM repeat (Oostdyk et al., 2019). It has also been shown that the open state of KPNA7 reduces the affinity for CAS and the efficiency of nuclear export (Oostdyk et al., 2019). The human KPNA7 autoinhibition form is relatively unstable, had reduced affinity for CAS and reduced efficiency of nuclear export, suggesting that the IBB adopts an open state, detached from ARM repeats, while the affinity for importin β is not particularly attenuated (Oostdyk et al., 2019). Conversely, it has also been reported that KPNA7 has a higher affinity for importin β than KPNA2, but that the affinity of the IBB domain of KPNA7 alone for importin β was not different from that of the IBB domain of KPNA2 alone (Oostdyk et al., 2019). Residues of the RXXR motif do not appear to be involved in importin β binding, which is consistent with the present analysis. These results are consistent with our prediction that KPNA7 has a low capacity to form the autoinhibition form, as described above, while the middle two residues of the RXXR motif are unlikely to be involved in importin β binding (Figure S10-I, -IIb).

In the autoinhibition form, the KRR motif of IBB fills the major NLS binding pocket with a large interface in all families in the model structure. The residue 50 in the middle of this motif is K in KPNA3 and has a contact area of 50 square angstroms less than the R in the other families, making the hydration energy less favorable than the families with R in this position (Figure S10-III, S13). This may be related to a previous study that showed that the middle R affects autoinhibition form formation (Harreman, Cohen, et al., 2003).

### Comparison of binding propensities of the IBB domain for partner proteins

As mentioned above, the structure of IBB in the three complexes studied here is quite different from each other (Figure S20). When the IBB binds to importin β, IBB forms a helix, and many of the conserved basic amino acids on IBB are concentrated on one side of the long helix, which is surrounded by acidic residues of importin β, resulting in a large interaction surface. In addition, when the nuclear export complex is formed, the peptide main chain of the IBB domain dropped in a completely different dihedral angle region. Furthermore, in this complex, the residues on the IBB alternate between patches of residues that have a large contact area with CAS and patches of residues that have a large contact area with ARM, with the IBB acting like a double-sided tape (Figure S10-II). In this case, at the point where the contact to CAS switches to the contact to ARM, there are adjacent parts with the same charge, such as 34EEE in subtype 1 and 42KR/K in all consensus sequences (Figure S16, S17). The charge repulsion at these sites may cause the side chains to move away from each other, creating a situation in which IBB can intrinsically adopt the above conformation. Thus, IBB retains the ability to form a helix over a long region, but at the same time retains the ability to extend and adopt conformations that reverse the orientation of the side chains at regular intervals, as well as the overall hydrophilicity that would allow the process of switching between them. All of these propensities seem to be essential for IBB to function as a chameleon sequence, and our results show that all of these propensities are evolutionary conserved in an exquisite manner.

Next, to examine the similarity of each interaction between families, clustering was performed among IBB domain consensus sequences using the interface area for each structure (Figure 5A). When the interface of each amino acid of the IBB domain in the three structures is correspondingly lined up, the amino acids that form the interfaces of each structure are arranged intricately in the IBB domain sequence. The positions where the interface areas were similar between the consensus sequences in each structure and the positions where the interface areas were different between the families were compared between the structures (Figure 5B).

**FIGURE 5.**
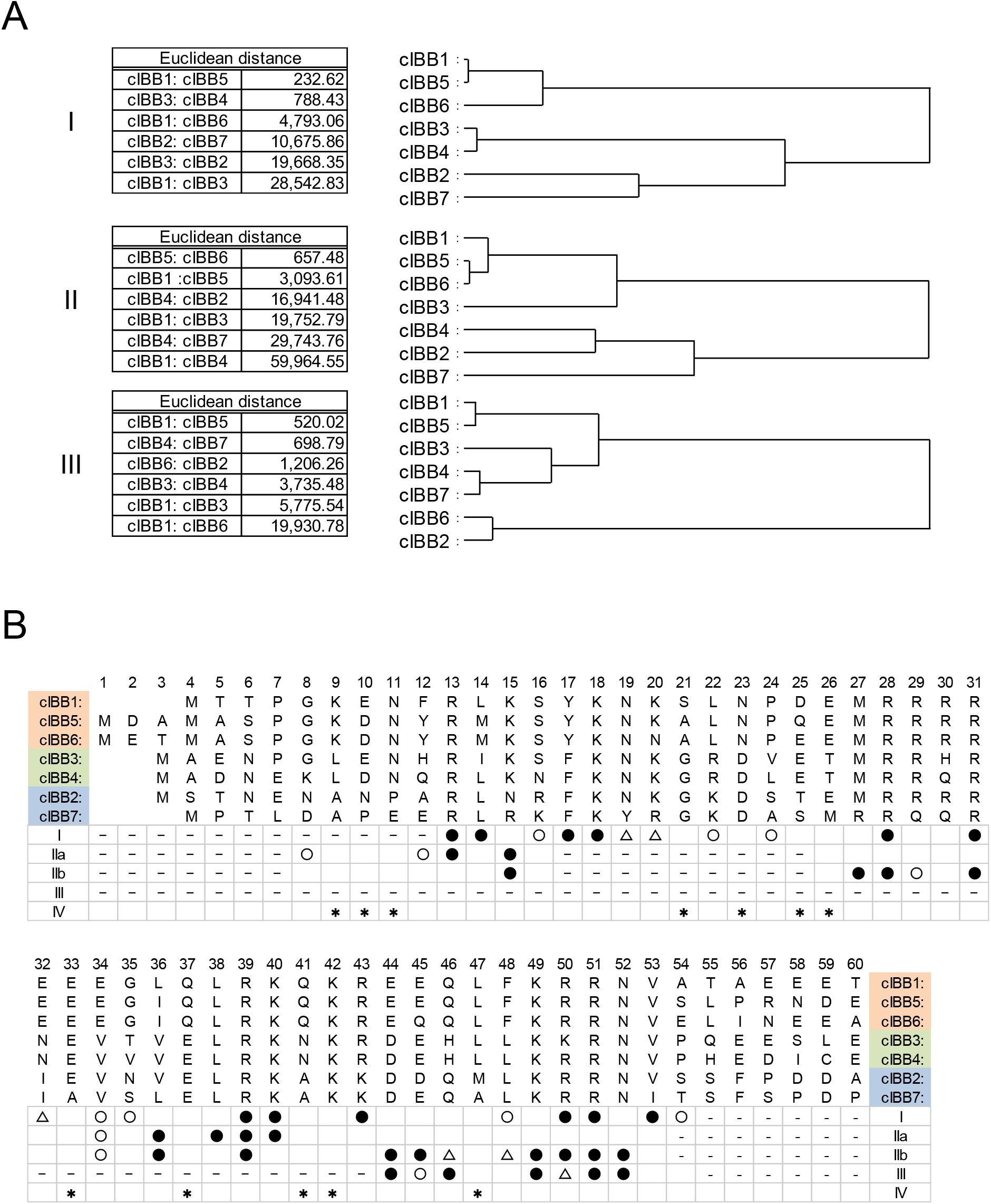
Consensus sequences of each family and important residues for the functional complex formation, functional switching, and functional difference among families. (A) Euclidean distance among the IBB consensus sequences obtained from the contact area at each position in each family. A dendrogram was drawn based on this Euclidean distance. (B) Interface between (I) IBB and importin β, (II) IBB and CAS and ARM repeat in the nuclear export complex, and (III) IBB and ARM repeat in the autoinhibition formed by importin α alone. ●: The positions where all family had commonly large contact area. ○: Positions where at least one family had large contact area and difference among family was large. Δ: Positions where some residues have a large interface, but the difference among family is small. Blank: Positions where none of family had large contact area. -: Positions where the residue was in a missing part in the template structure. *: Positions where none of family have a large interface in all structures.

The contact area distribution pattern in the interface with importin β was divided into three groups, KPNA1, 5, 6, KPNA 3, 4, and KPNA2, 7, which was the same as the canonical subtype classification (Figure 5A). The interface pattern in the nuclear export complex was divided in the same way as the phylogenetic tree of the primary sequence of the IBB domain consensus sequence inferred using the Neighbor-Joining method (Figure 5A, S2A). The interface pattern in autoinhibition neither correlated with the canonical subtype classification of whole importin α nor the phylogenetic tree of the primary sequence of the consensus sequence. The fact that clustering by the distribution pattern of contact surface area, as well as by the primary sequence of the IBB alone, results in a different grouping from the conventional classification, suggested that a more rigorous mode of interaction is implicitly conserved.

The fact that there are elements to deliver differences between the members of a family in interaction with each binding partners suggests that the balancing mode between the interactions may also be capable of producing complexed personalities. For example, the autoinhibition form includes interactions of the basic amino acids 49, 50, and 51 in the IBB domain with the amino acids in the major NLS site as seen in the 1IAL interface (Figure S10-III). This interaction is also observed when the export complex is formed with CAS/CSE1 as seen in 1WA5 (Figure S10-IIb). It is likely that the IBB must switch its conformation from the autoinhibition form to the export complex in the nuclear transport sequence, but it is not yet clear whether interaction of IBB with the major NLS binding site is once released or not before the export complex is formed. However, our analysis suggests that differences in the binding mode of each family member in the complex may lead to differences in efficiency on either side, which in turn may affect the recycling of importin α (Figure S4D-IIb, III). For example, although the autoinhibition form taken by importin α and the autoinhibition form used in complex with CAS are often considered to be the same, our analysis indicated that there may in fact be differences in the status of IBB in these forms, and these may also differ between families.

Differences in binding propensities such as those revealed here are also be found in other IBB domain binding molecules, such as DNA (Jibiki et al., 2021) and Rbbp4 (Tsujii et al., 2015). Moreover, as importin α plays an important role in multiple cellular events (Oka & Yoneda, 2018), there may exist additional unknown binding partners. It is possible that differences between the families affect the competition and switching involving these binding partners.

In addition to the three important basic clusters in the IBB, this study has identified amino acids or biochemical/physicochemical propensities that are significant for the interaction of each functional structure or that characterize the family (Figure 5B). Interestingly, we found organisms in the sequence set that did not have basic residues which are important for this interaction. For example, the arginine at position 39 was 100% conserved in the organisms in the sequence set of KPNA1, KPNA2, KPNA5, KPNA6, and KPNA7, but KPNA3 in *Takifugu rubripes* (Japanese pufferfish) and KPNA4 in *Callorhinchus milii* (Ghost shark) were D and Q, respectively (Supporting file 1, Supporting file 2, Supporting file 4). These substitutions may have an inhibitory effect on the binding to CAS, especially in *Takifugu rubripes* (Japanese pufferfish) as the substitution to D may cause electrostatic repulsion with the ARM repeat.

In this analysis, it was revealed that there are several residue positions that were predicted to have reduced involvement in the formation of the interface between the three functional structures (Figure 5B). The interorganisms conservation of amino acid residues in all these positions is high for all family members except KPNA2 and KPNA7, which belong to subtype 2. For example, K42 is retained by all consensus sequences and is highly conserved among organisms (Supporting file 4). This residue is thought to be involved in binding to DNA (Jibiki et al., 2021) and Rbbp4 (Tsujii et al., 2015) and may be important for such functions. In addition, the biochemical/physicochemical propensity of small bulkiness was conserved at position 21, although the residue differed among families (Figure S17, S18). In addition, positions 23 and 37 retained acidic amino acids only in subtype 3 and subtype 2. These residues were all highly conserved and may be important for specific interactions giving the families their individual functions (Supporting file 4). On the other hand, for KPNA2 and KPNA7, some residues have a high degree of interorganisms conservation, yet others have a very low degree (Figure 2D, 3B). These results suggest that there is evolutionary selection pressure on the IBB domain to maintain functions other than the three functions analyzed here and that for some of these functions, selection pressure is significantly weaker in KPNA2 and KPNA7. This suggests that KPNA2 and KPNA7, especially KPNA7 with its significantly less conserved residues, may have lost some of these extra functions and specialized in only a limited number of functions.

In this study, we have determined the consensus sequences of the IBB domain from sequence information of a wide range of organisms and by using information on the structure of functionally relevant complexes. We were able to distinguish and scrutinize the contribution of each residue to multiple functions, and the origin of IBB’s propensities as a multifunctional ChSeq was demonstrated in detail. We have also identified residues that are presumed to be responsible for differences in complex-forming ability among families. Furthermore, we not only gained a detailed understanding of the residues involved in these functions, but we found that there are residues that are universally important for IBB functions across organisms and families, in addition to those previously known. However, there are likely many other noncanonical IBB domains of proteins that have not yet been identified. The information on the importance of position-specific biochemical/physicochemical propensities provided by this study will be useful for predicting the function of such IBB domains.

## Experimental procedures

### Database

The amino acid sequence of each importin α family member included in the sequence set was obtained from UniProtKB (Bateman et al., 2021). The amino acid sequence of human KPNA1 (UniProtKB: P52294), human KPNA2 (UniProtKB: P52292), human KPNA3 (UniProtKB: O00505), human KPNA4 (UniProtKB: O00629), human KPNA5 (UniProtKB: O15131), human KPNA6 (UniProtKB: O60684), human KPNA7 (UniProtKB: A9QM74), human KPNB1 (UniProtKB: Q14974), human RAN (UniProtKB: P62826), human CSE1L (UniProtKB: P55060), Baker’s yeast SRP1 (UniProtKB: Q02821), mouse KPNA2 (UniProtKB: P52293), dog RAN (UniProtKB: P62825) and Baker’s yeast CSE1 (UniProtKB: P33307) were also obtained from UniProtKB as target sequences or template sequences in homology modeling. The structure of the IBB domain and importin β (PDB ID: 1QGK) complex, the tripartite complex of CSE1, importin α and RanGTP (PDB ID: 1WA5), and importin α monomer (PDB ID: 1IAL) were obtained from PDBj (Protein Data Bank Japan) (Kinjo et al., 2017, Kinjo et al., 2018) and used as a template structure for homology modeling.

### Creating a sequence set

We first extracted 8268 entries with a PROSITE ID (Sigrist et al., 2013) of IBB domain PS51214 from UniProtKB (release date 2021_02). From them, 1644 entries that have “KPNA” as their gene name were extracted. Entries that have an IBB domain sequences shorter than 50 residues were left out as truncated fragments and identical sequences of the same organisms were clustered.

### Multiple sequence alignment

The multiple sequence alignment was performed using CULUSTALW 2.1 at GenomeNet (Larkin et al., 2007). The pairwise alignment was always conducted with slow-accurate mode, and the weight matrix was fixed to BLOSUM for PROTEIN both in the pairwise and the multiple alignment. The gap open penalty and the gap extension penalty were set to 0.5 and 0.1, respectively, for the pairwise and the multiple alignment. Only multiple alignments for homology modeling were performed under the following conditions. The gap open penalty and the gap extension penalty was set to 10.0 and 0.1, respectively, for pairwise alignment. The gap open penalty and the gap extension penalty was set to 10.0 and 0.05, respectively, for multiple alignment.

### Phylogenetic analysis

The evolutionary history of the IBB or full-length consensus sequences were inferred using the Neighbor-Joining method (Saitou & Nei, 1987) or Maximum Likelihood method and Poisson correction model (ZUCKERKANDL & PAULING, 1965). In the Neighbor-Joining method, the optimal tree is shown. The percentage of replicate trees in which the associated taxa clustered together in the bootstrap test (500 replicates) are shown next to the branches (Felsenstein, 1985). The tree was drawn with branch lengths in the same units as those of the evolutionary distances used to infer the phylogenetic tree. The evolutionary distances were computed using the Poisson correction method (ZUCKERKANDL & PAULING, 1965) and are in the units of the number of amino acid substitutions per site. All ambiguous positions were removed for each sequence pair (pairwise deletion option). There were a total of 68 and 560 positions in the final dataset for IBB or full-length consensus sequences, respectively. Evolutionary analyses were conducted in MEGA X 10.2.5 (Kumar et al., 2018). In the Maximum Likelihood method and Poisson correction model, the trees for cIBBs and cKPNAs with the highest log likelihood (−703.46) or (−6081.23) were shown, respectively. The percentage of trees in which the associated taxa clustered together is shown next to the branches. Initial tree(s) for the heuristic search were obtained automatically by applying Neighbor-Join and BioNJ algorithms to a matrix of pairwise distances estimated using the Poisson model, and then selecting the topology with superior log likelihood value. The tree is drawn to scale, with branch lengths measured in the number of substitutions per site. The phylogenetic tree of cIBBs was constructed using 7 different sequences of 68 residues in length (including gaps inserted by alignment). For full-length cKPNAs, phylogenetic trees were obtained using 31 different sequences of 560 residues in length (including gaps inserted by alignment) for seven family members because there were multiple equivalent consensus sequences, two patterns of consensus sequences for cKPNA2 and 24 patterns for cKPNA7, for the non-IBB portion. Evolutionary analyses were conducted in MEGA X 10.2.5 (Kumar et al., 2018).

### Homology modeling

Modeling of the three-dimensional structure of the IBB consensus sequence was conducted by the ProMod3 3.2.0 on Swiss-Model modeling server (Waterhouse et al., 2018, Studer et al., 2021). Importin β binding form (PDB ID: 1QGK), nuclear export complexed with CAS/CSE1 and Ran-GTP (PDB ID: 1WA5), and the autoinhibition form (PDB ID: 1IAL) were used for template structures. For the modeling of the nuclear export complex and the autoinhibition form, chimeric sequences were prepared by replacing the IBB domain portion of the full-length human importin α family sequences (UniProtKB: P52294, P52292, O00505, O00629, O15131, O60684, A9QM74) with the corresponding IBB domain consensus sequence. The target sequences of interacting molecules have been unified to human sequences. The importin β binding form was constructed using the IBB domain consensus sequence and the human KPNB1 sequence (UniProtKB: Q14974) as hetero targets. For the construction of the nuclear export complex and the autoinhibition form, the chimeric sequences, human RAN sequence (UniProtKB: P62826), and human CSE1L sequence (UniProtKB: P55060) were subjected to multiple sequence alignment with Baker’s yeast SRP1 sequence (UniProtKB: Q02821) or mouse KPNA2 sequence (UniProtKB: P52293), dog RAN sequence (UniProtKB: P62825), and Baker’s yeast CSE1 sequence (UniProtKB: P33307), respectively, using ClustalW. Their structures in the nuclear export complex or the autoinhibition form were constructed individually with alignment mode. The alignment between the target sequence and the template sequence which was adopted in the modeling on Swiss-Model is shown in Supporting file 4. For the nuclear export complex, the structures of chimeric sequences, human RAN sequence, and human CSE1L were merged with Swiss-Pdb Viewer4.1.0. (Guex & Peitsch, 1997). Furthermore, to confirm the consistency of the interface of each molecule, hetero-target modeling of the chimeric sequences and human RAN sequence (UniProtKB: P62826), and human CSE1L sequence (UniProtKB: P55060) was performed using the PDB file generated with Swiss-Pdb Viewer as a template. The RMSD (Root Mean Square Deviation) of all atoms between the model and the template was calculated by the PyMOL Molecular Graphics System, Version 2.5.2 Schrödinger, LLC.

### Calculation of contact area for each residue of IBB domain in the interface

The solvent accessible surface area of each residue of the IBB domain in the three-dimensional structure with or without interacting molecule was calculated by STRIDE (Heinig & Frishman, 2004). The difference in the solvent accessible surface area of each residue between those structures was defined as the contact area to the binding partner molecule for each residue.

### Calculation of energy for each residue of IBB domain in the interface

The nonBonded and electrostatic constraint terms in the interaction energy of each amino acid residue in complex formation were calculated as the difference in energy between the presence and absence of each binding partner using the GROMOS96 43B1 force field in Swiss-Pdb Viewer 4.1.0. The solvation free energy between cIBB and each binding partner was calculated by ‘Protein interfaces, surfaces and assemblies’ service PISA v1.52 at the European Bioinformatics Institute (http://www.ebi.ac.uk/pdbe/prot_int/pistart.html) (Krissinel & Henrick, 2007).

### Scoring of amino acids in the IBB domain by biochemical propensities

The score indices for polarity (Zimmerman et al., 1968), hydrophobicity (Kyte & Doolittle, 1982), bulkiness (Zimmerman et al., 1968), average flexibility (BHASKARAN & PONNUSWAMY, 2009), α-helix (Chou & Fasman, 2006), β-sheet (Chou & Fasman, 2006), β-turn (Chou & Fasman, 2006), and coil (Deléage & Roux, 1987) shown in Figure S15A were standardized to give a mean of 5 and a standard deviation of 1 for each biochemical/biophysical propensity.

For the scoring of the consensus sequences, each residue in the IBB domain consensus sequence was scored using the normalized score index in Figure S15B. To calculate the average score for each residue position, each residue of the whole IBB domain sequence in the sequence set was also scored using the normalized score index in Figure S15B and averaged over all organisms. The relative positions were the same as those in the multiple sequence alignment generated in the process of the consensus sequence determination, and gaps were ignored.

The acidic, basic, and helix breaker amino acids were represented by a dummy variable 1 for aspartic acid and glutamic acid, lysine and arginine, and glycine and proline, respectively. Other amino acids and gaps were given as a dummy variable 0 and averaged over all organisms. A final indicator of the tendency to form each secondary structure, α-helix, β-sheet, β-turn, and coil, was calculated by taking the geometric mean of the above values over five residues along the primary sequence.

### Cluster analysis

Biochemical/biophysical propensities of the consensus sequences of seven importin α family members were analyzed by cluster analysis using the average score over all organisms of each biochemical propensity of the residue at each position of the IBB domain. The contact area for each residue of the IBB domain consensus sequence at the interface in the model structures were also clustered. The clustering was performed by Ward’s method based on the Euclidean distances of these metrics. The range of residues used in the clustering of the propensities was 4-60 in Figure S16, and 6-58 were used for the analysis of tendency to form each secondary structure, α-helix, β-sheet, β-turn, and coil. For the clustering of contact area distribution, the residues 13–54, 8–16, and 26–53, 44–54 of Figure S10 were used for importin β binding form, nuclear export complex, and autoinhibition form, respectively.

### Ramachandran plot

The ramachandran plots of IBB domain residues in 1QGK, 1WA5, and 1IAL used as templates for homology modeling were displayed by Swiss-Pdb Viewer 4.1.0.

### Screening for similarities and differences in consensus sequences for each residue at the interface

Using all residues of all family members including non-contacting residues, the average value of the contact area per residue was calculated for the model structures of the importin β binding form, nuclear export complex, and autoinhibition form. In order to extract the differences between families at each residue position for each complex structure, the ratio of the maximum and minimum contact area of the whole family at each position was calculated. After that, the position of the residue which is important for the interaction between the IBB domain and the binding partner molecule were screened as follows. First, the positions where at least one contact area of the residue was equal or higher than the average value were extracted. Second, if the ratio of maximum value to minimum value among the family was below the median, the position is defined as a commonly important position among the family. Third, if the ratio of maximum value to minimum value among the family was greater than or equal to 2, the position is defined as specifically important position for some family members. In the later criteria, residues were further screened by the difference from the minimum or the maximum values of contact area at each position. First, the value obtained by dividing the difference between the minimum value and the maximum value by 3 was used as the reference value. If the difference between the value of the residue and the difference between the maximum values is smaller than the reference value, the residue was defined as having a relatively large contact at interface. If the difference from the maximum value is smaller than the reference value, the residue was defined as having a relatively small contact at interface. The residue that takes a value of contact area between the two was defined as having medium contact at interface.

## Abbreviations

KPNA: karyopherin alpha
IBB domain: importin β binding domain
ARM repeats: armadillo repeats
NAAT domain: Nuclear Acid Associating Trolley pole domain
ChSeqs: chameleon sequences
IDPs: intrinsically disordered proteins
IDRs: intrinsically disordered regions
NLS: nuclear localization signal
CAS: Cellular Apoptosis Susceptibility protein
CSE1: Chromosome Segregation 1
Rbbp4: RB Binding Protein 4
Nup50: Nucleoporin 50

