## Supporting figure for "Biochemical propensity mapping for structural and functional anatomy of importin α IBB domain"

### FIGURE S1

| Class | KPNA1 | KPNA5 | KPNA6 | KPNA3 | KPNA4 | KPNA2 | KPNA7 |
|---|---|---|---|---|---|---|---|
| Anthozoa | -- | -- | 1 (0.4%) | -- | 1 (0.7%) | 1 (0.5%) | -- |
| Hydrozoa | -- | -- | 1 (0.4%) | 1 (0.8%) | -- | 1 (0.5%) | -- |
| Trematoda | -- | -- | -- | 1 (0.8%) | -- | -- | -- |
| Enoplea | -- | -- | -- | 7 (5.6%) | 1 (0.7%) | -- | -- |
| Chromadorea | -- | 1 (0.7%) | -- | -- | -- | -- | -- |
| Arachnida | -- | 1 (0.7%) | 1 (0.4%) | -- | 2 (1.4%) | 1 (0.5%) | -- |
| Malacostraca | -- | -- | -- | -- | 1 (0.7%) | 2 (1.1%) | -- |
| Insecta | 1 (0.6%) | 1 (0.7%) | 7 (2.8%) | 3 (2.4%) | 6 (4.1%) | 5 (2.7%) | -- |
| Hexanauplia | -- | -- | 1 (0.4%) | -- | -- | 1 (0.5%) | -- |
| Ascidiacea | 1 (0.6%) | -- | 1 (0.4%) | -- | 1 (0.7%) | -- | -- |
| Chondrichthyes | -- | 3 (2.0%) | 2 (0.8%) | -- | 3 (2.0%) | 1 (0.5%) | 1 (1.0%) |
| Actinopterygii | 37 (23.0%) | 28 (18.7%) | 60 (24.1%) | 35 (28.2%) | 33 (22.3%) | 83 (44.4%) | 3 (3.1%) |
| Sarcopterygii | 1 (0.6%) | 1 (0.7%) | -- | 1 (0.8%) | 1 (0.7%) | 1 (0.5%) | -- |
| Amphibia | 8 (5.0%) | -- | 7 (2.8%) | 5 (4.0%) | 4 (2.7%) | 7 (3.7%) | 5 (5.2%) |
| Reptilia | 31 (19.3%) | 28 (18.7%) | 33 (13.3%) | 19 (15.3%) | 23 (15.5%) | 25 (13.4%) | 24 (25.0%) |
| Mammalia | 82 (50.9%) | 85 (56.7%) | 135 (54.2%) | 52 (41.9%) | 72 (48.6%) | 59 (31.6%) | 63 (65.6%) |
| Magnoliopsida | -- | 2 (1.3%) | -- | -- | -- | -- | -- |
| Total | 161 | 150 | 249 | 124 | 148 | 187 | 96 |

**FIGURE S1 A table of the class distributions of each KPNA family protein in the sequence sets used in this study.** The number of organisms belonging to each class and their percentage of the total is shown in parentheses. The double hyphens indicate that the organisms in the corresponding class were not included in the corresponding KPNA family protein sequence set.

### FIGURE S2

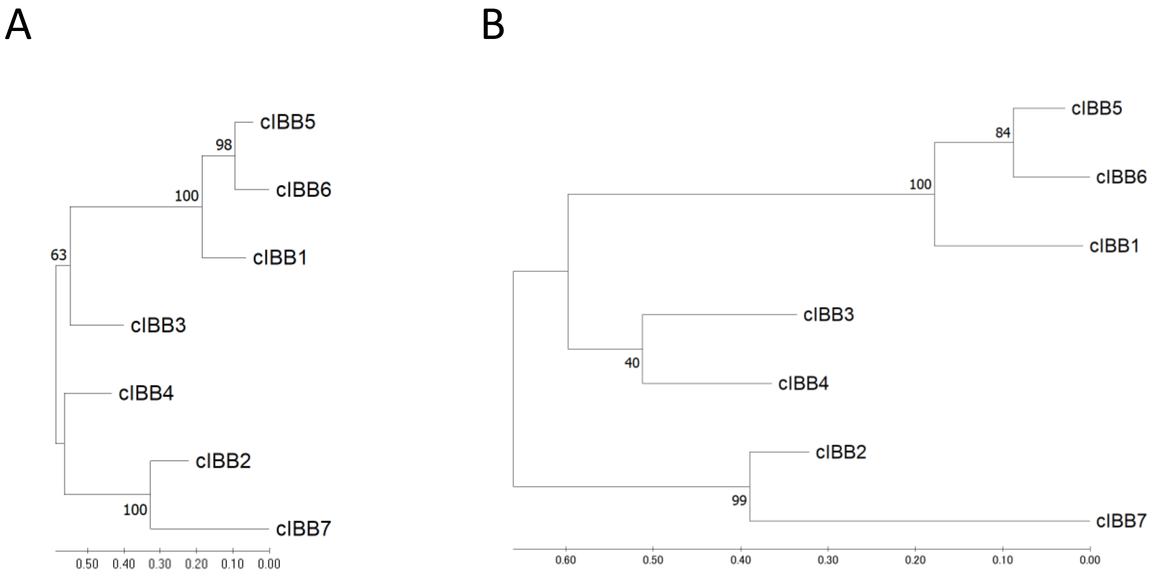


**FIGURE S2 The Phylogenetic analysis of IBB domain consensus sequences.** The phylogenetic tree of IBB consensus sequences of the importin $\alpha$ family inferred using the Neighbor-Joining method (A), or Maximum Likelihood method and Poisson correction model (B).

### FIGURE S3

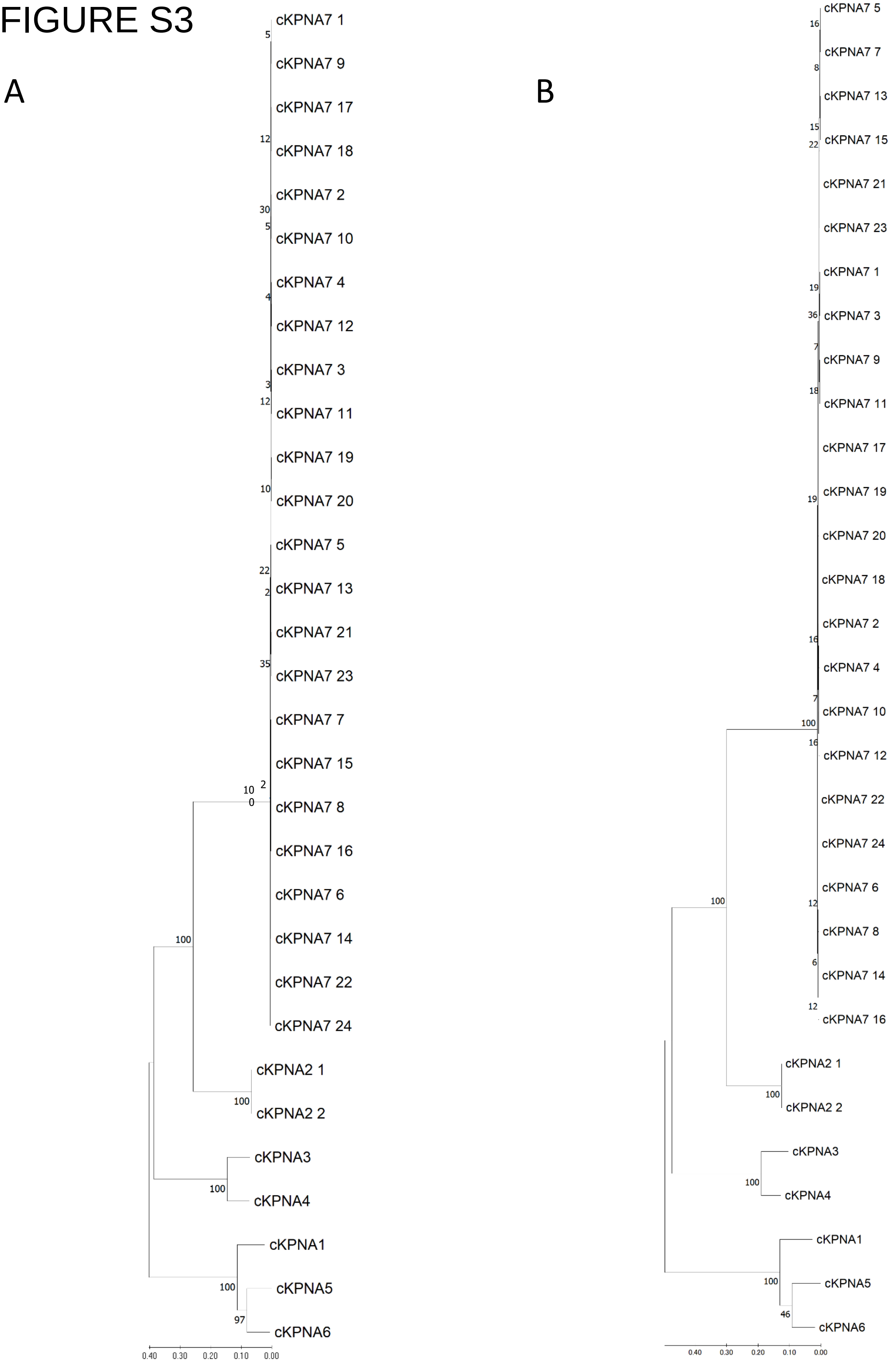


**FIGURE S3 The Phylogenetic analysis of full-length consensus sequences.** The phylogenetic tree of full-length consensus sequences of the importin $\alpha$ family inferred using the Neighbor-Joining method (A), or Maximum Likelihood method and Poisson correction model (B). This analysis involved 31 amino acid sequences including two and 24 variants for KPNA2 and KPNA7, respectively.

### FIGURE S4

## A

| Target | template | GMQE |
|---|---|---|
| cIBB1 | chain B | 0.89 |
| human KPNB1 | chain A | |
| cIBB5 | chain B | 0.89 |
| human KPNB1 | chain A | |
| cIBB6 | chain B | 0.89 |
| human KPNB1 | chain A | |
| cIBB3 | chain B | 0.89 |
| human KPNB1 | chain A | |
| cIBB4 | chain B | 0.89 |
| human KPNB1 | chain A | |
| cIBB2 | chain B | 0.89 |
| human KPNB1 | chain A | |
| cIBB7 | chain B | 0.89 |
| human KPNB1 | chain A | |

## B

| Target | template | GMQE |
|---|---|---|
| human RAN | chain A | 0.74 |
| chimeric KPNA1 | chain B | 0.76 |
| chimeric KPNA5 | chain B | 0.76 |
| chimeric KPNA6 | chain B | 0.77 |
| chimeric KPNA3 | chain B | 0.74 |
| chimeric KPNA4 | chain B | 0.74 |
| chimeric KPNA2 | chain B | 0.76 |
| chimeric KPNA7 | chain B | 0.74 |
| human CSE1L | chain C | 0.73 |

## C

| Target | template | GMQE |
|---|---|---|
| chimeric KPNA1 | chain B | 0.84 |
| human RAN | chain A | |
| human CSE1L | chain C | |
| chimeric KPNA5 | chain B | 0.83 |
| human RAN | chain A | |
| human CSE1L | chain C | |
| chimeric KPNA6 | chain B | 0.84 |
| human RAN | chain A | |
| human CSE1L | chain C | |
| chimeric KPNA3 | chain B | 0.83 |
| human RAN | chain A | |
| human CSE1L | chain C | |
| chimeric KPNA4 | chain B | 0.83 |
| human RAN | chain A | |
| human CSE1L | chain C | |
| chimeric KPNA2 | chain B | 0.83 |
| human RAN | chain A | |
| human CSE1L | chain C | |
| chimeric KPNA7 | chain B | 0.84 |
| human RAN | chain A | |
| human CSE1L | chain C | |

## D

| Target | template | GMQE |
|---|---|---|
| chimeric KPNA1 | chain A | 0.69 |
| chimeric KPNA5 | chain A | 0.69 |
| chimeric KPNA6 | chain A | 0.70 |
| chimeric KPNA3 | chain A | 0.74 |
| chimeric KPNA4 | chain A | 0.73 |
| chimeric KPNA2 | chain A | 0.82 |
| chimeric KPNA7 | chain A | 0.76 |

**FIGURE S4 GQME scores for modeled structures.** The GMQE of the modeled structures of cIBBs and human KPNA1 using 1QGK as the template (A), human RAN, chimeric KPNAs, and human CSE 1L using each chain of 1WA5 as the template (B), chimeric KPNAs, human RAN, and human CSE1L using the chimeric KPNAs / human RAN / human CSE1L complex merged by Swiss-Pdb Viewer as the template (C), and chimeric KPNAs using 1IAL as the template (D) are shown.

### FIGURE S5

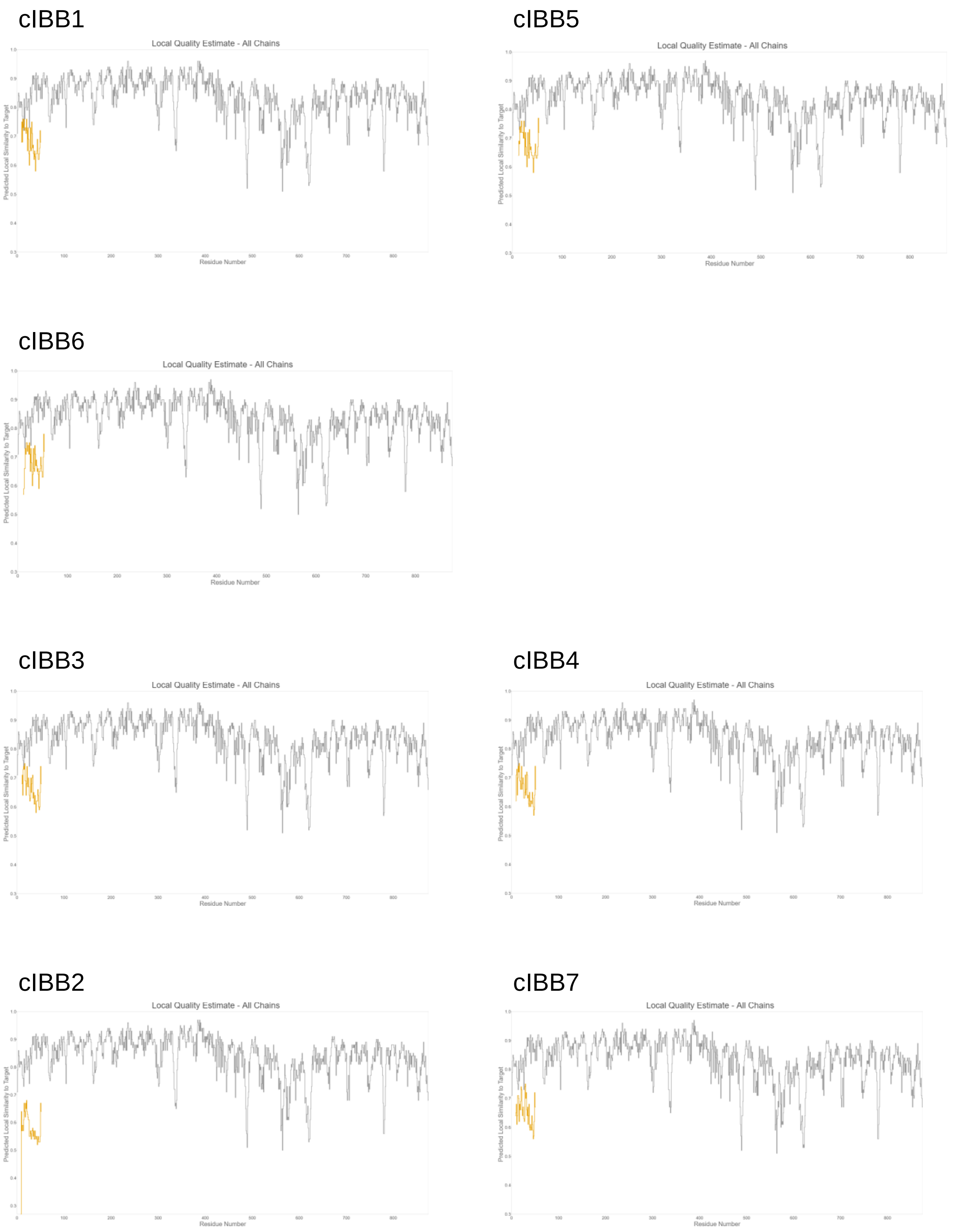


**FIGURE S5 GMQE scores for each residue in the moleded import β binding complex.** GMQE scores of each residue in the modeled cIBBs and human KPNA1 using 1QGK as a template. The yellow and gray line graphs represents cIBBs and human KPNB1, respectively.

### FIGURE S6

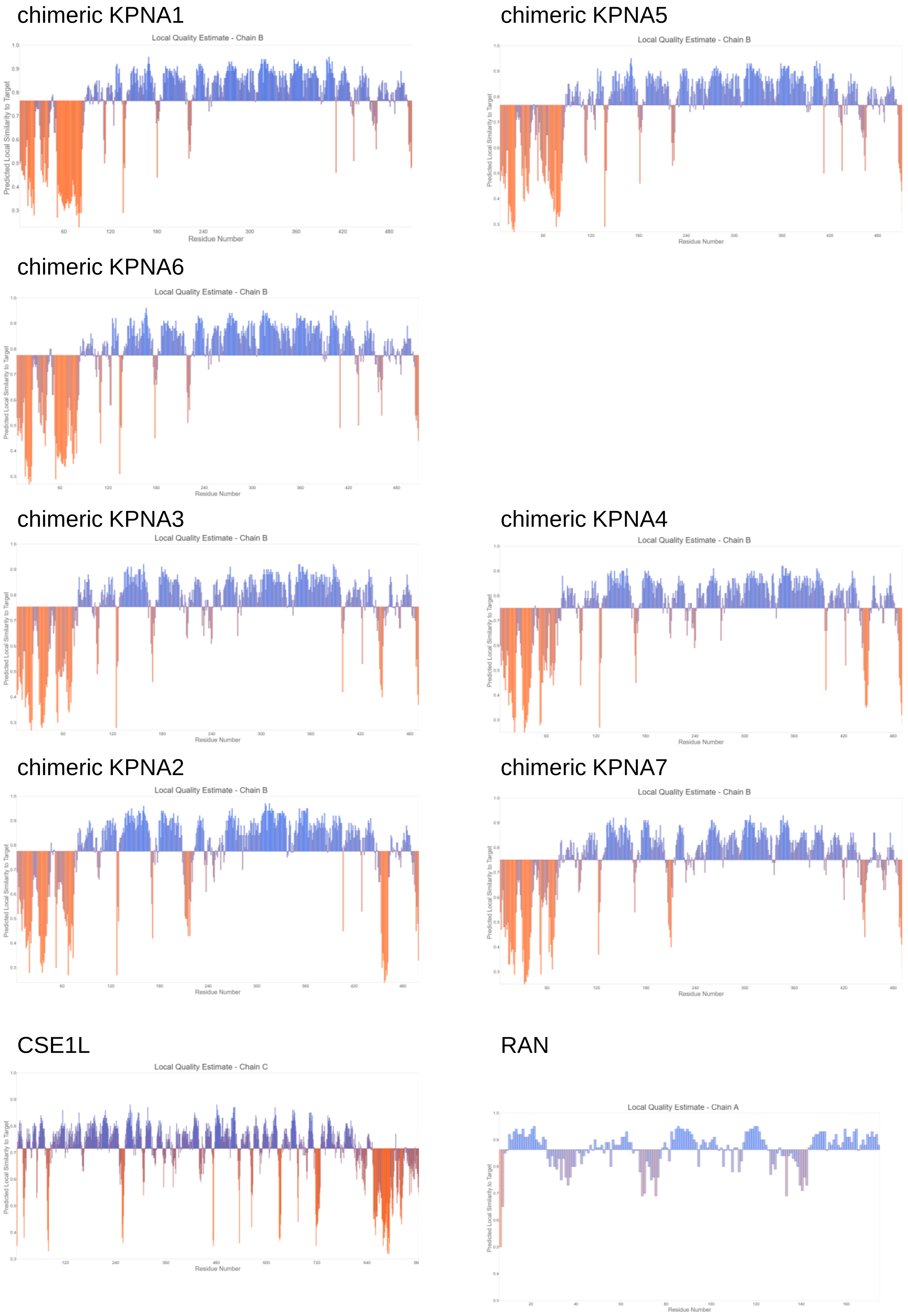


**FIGURE S6 GMQE scores for each residue in the modeled chimeric KPNAs, human RAN and human CSE1L in the nuclear export complex.** GMQE scores of each residue in the modeled human RAN, chimeric KPNAs and human CSE 1L using each chain of 1WA5 as the template.

### FIGURE S7

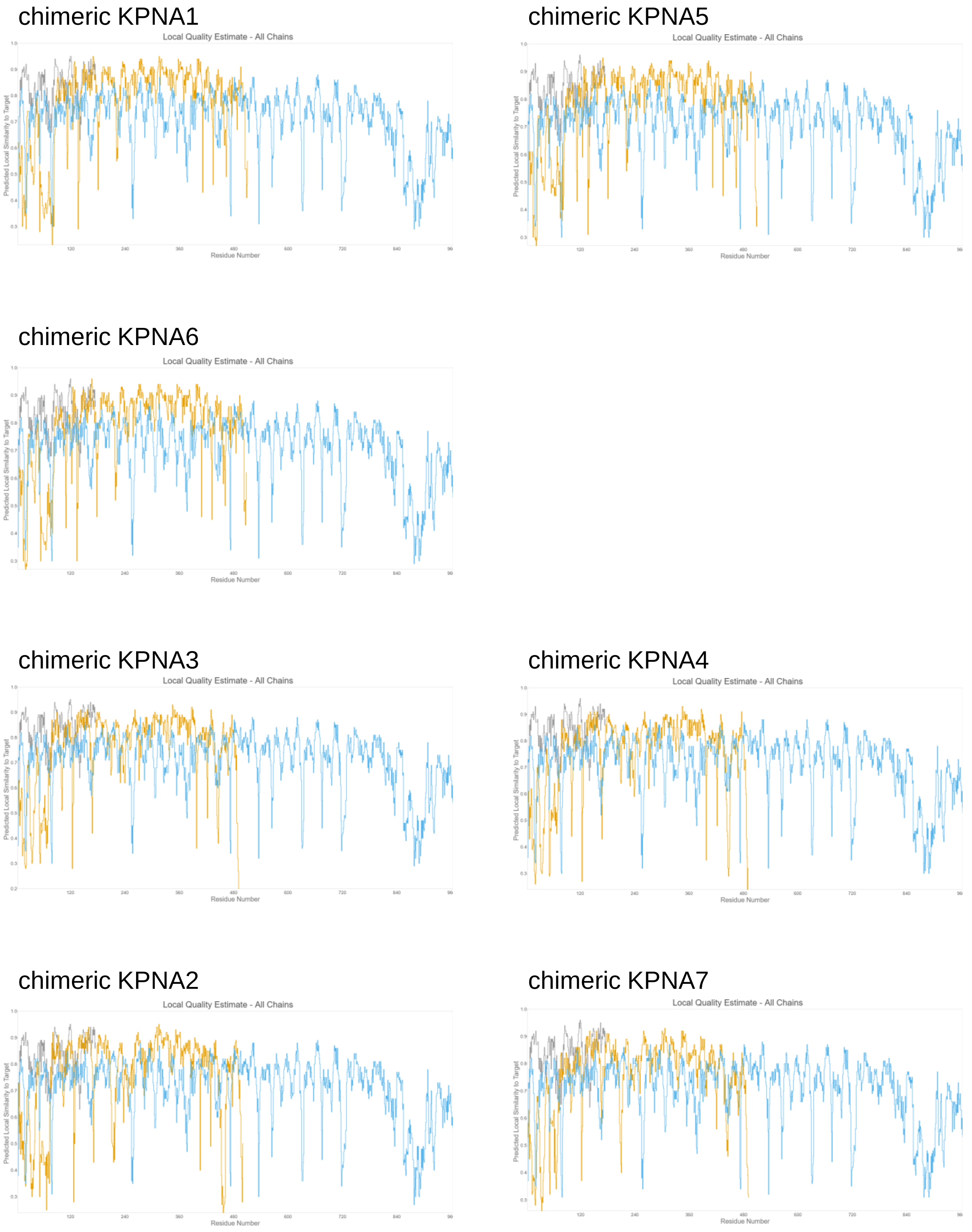


**FIGURE S7 GMQE scores for each residue in the modeled nuclear export complex.** GMQE scores of each residue in the modeled chimeric KPNAs, human RAN, and human CSE1L using the chimeric KPNAs / human RAN / human CSE1L complex merged by Swiss-Pdb Viewer as the template. The yellow, gray and light blue lines represents chimeric KPNAs, human RAN and human CSE1L, respectively.

### FIGURE S8

chimeric KPNA1

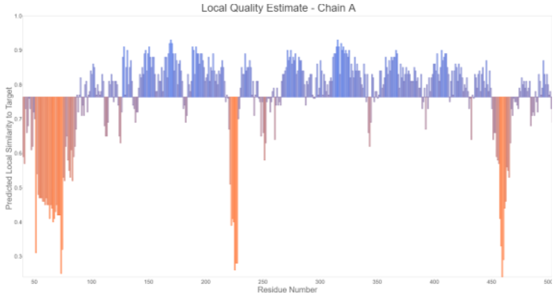


chimeric KPNA5

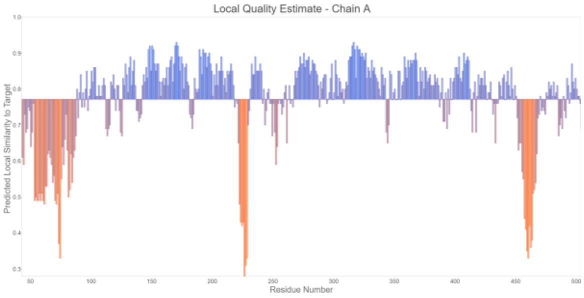


chimeric KPNA6

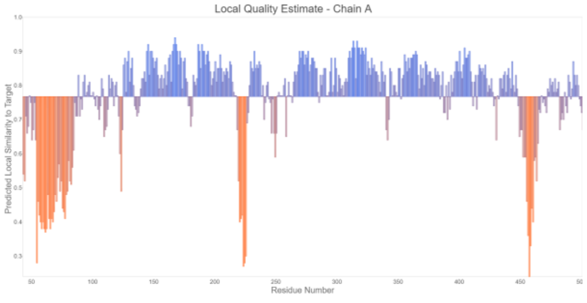


chimeric KPNA3

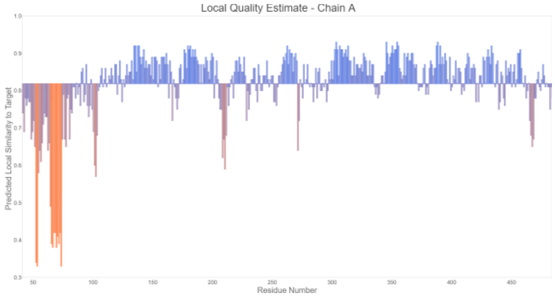


chimeric KPNA4

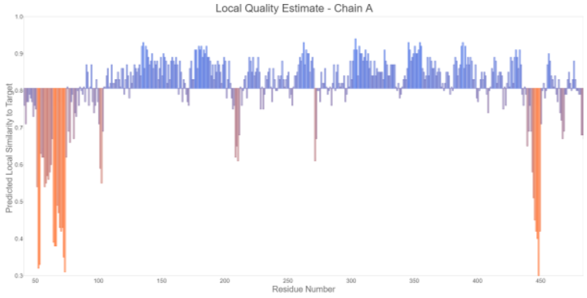


chimeric KPNA2

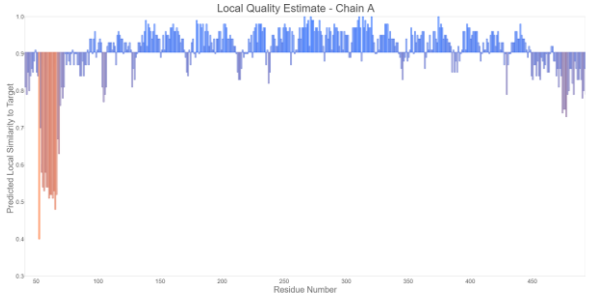


chimeric KPNA7

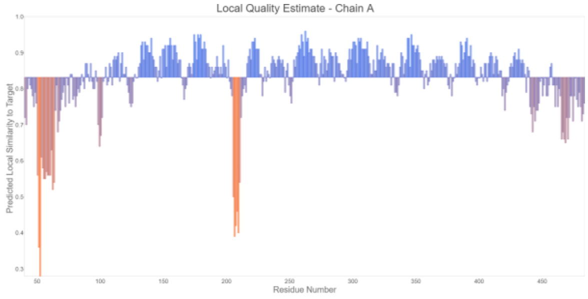


**FIGURE S8 GMQE scores for each residue in the modeled autoinhibition form.** GMQE scores of each residue in the modeled chimeric KPNAs using 1IAL as the template.

### FIGURE S9

| Template | Target | | RMSD (Å) |
|---|---|---|---|
| 1QGK | cIBB1 | human KPNB1 | 0.099 |
| | cIBB5 | human KPNB1 | 0.109 |
| | cIBB6 | human KPNB1 | 0.109 |
| | cIBB3 | human KPNB1 | 0.090 |
| | cIBB4 | human KPNB1 | 0.093 |
| | cIBB2 | human KPNB1 | 0.134 |
| | cIBB7 | human KPNB1 | 0.115 |

| Template | Target | | | RMSD (Å) |
|---|---|---|---|---|
| 1WA5 | chimeric KPNA1 | human RAN | human CSE1L | 0.296 |
| | chimeric KPNA5 | human RAN | human CSE1L | 0.298 |
| | chimeric KPNA6 | human RAN | human CSE1L | 0.220 |
| | human RAN | human RAN | human CSE1L | 0.272 |
| | chimeric KPNA4 | human RAN | human CSE1L | 0.227 |
| | chimeric KPNA2 | human RAN | human CSE1L | 0.284 |
| | chimeric KPNA7 | human RAN | human CSE1L | 0.272 |

| Template | Target | RMSD (Å) |
|---|---|---|
| 1IAL | chimeric KPNA1 | 0.151 |
| | chimeric KPNA5 | 0.150 |
| | chimeric KPNA6 | 0.156 |
| | human RAN | 0.151 |
| | chimeric KPNA4 | 0.143 |
| | chimeric KPNA2 | 0.093 |
| | chimeric KPNA7 | 0.137 |

**FIGURE S9 The deviations between the coordinate values of modeled and corresponding template structures.** The RMSD of all atoms between the modeled structures and the corresponding template structures are shown.

### FIGURE S10

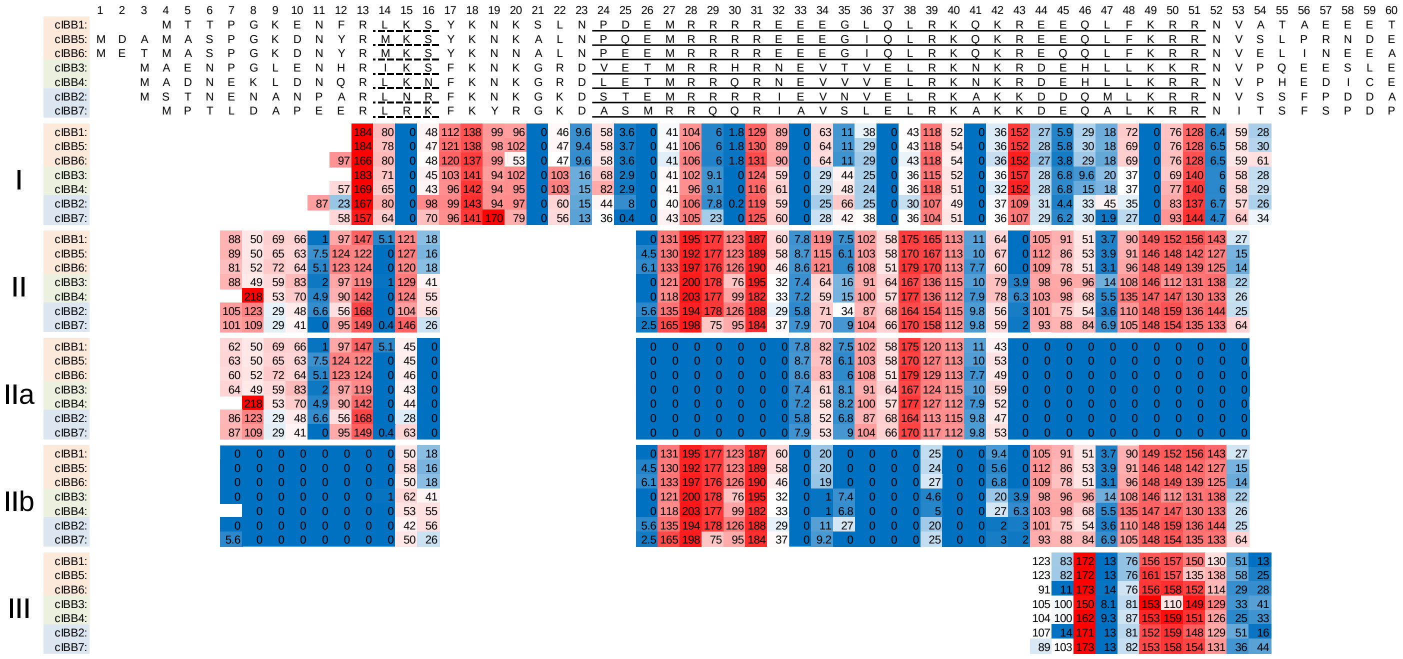


**FIGURE S10 Interface contributions of residues of cIBB domains in binding complexes with partner proteins.** The contributions for each residue of cIBBs in the interface in the complex with the binding partners, importin β binding form (I), export complex with CAS/CSE1 (IIa), ARM repeats (IIb) separated and total of CAS/CSE1, ARM repeats and RAN (II), and autoinhibition form (III) is mapped as a per-residue interface area (Å$^2$) in the IBB domain. The Heat map is presented with red (high) and blue (low) with the median in white. The residues which all the cIBBs formed 310 helix and α-helix are underlined with dotted and solid lines, respectively.

### FIGURE S11

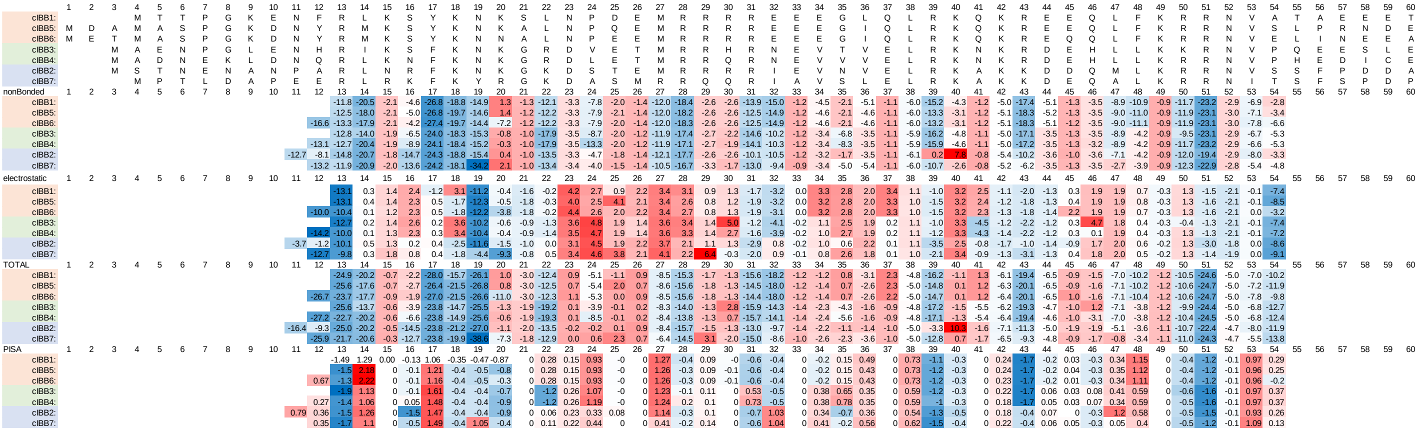


**FIGURE S11 The free energy change for each residue of cIBBs in the importin β binding complex.** The free energy changes of nonBonded interaction, electrostatic interaction, and the total of them and solvation for each residue of cIBBs due to interaction with importin β in the modeled importin β binding complexes were calculated. The energy unit of nonBonded interaction and electrostatic interaction are shown in (KJ / mol), and the energy unit of solvation is shown in (kcal / mol). The Heat map is presented with red (high) and blue (low) with the median in white.

### FIGURE S12

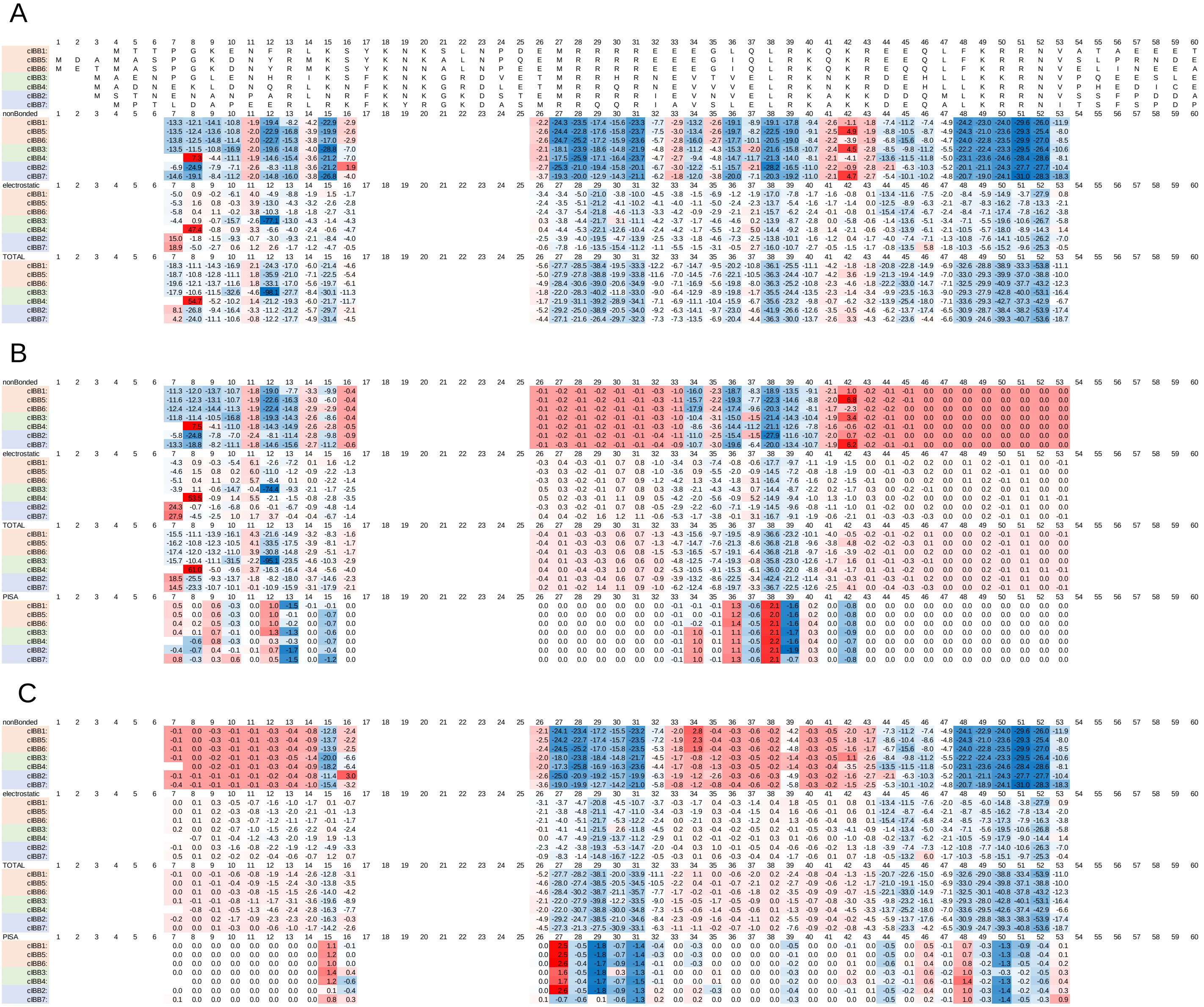


**FIGURE S12 The free energy change for each residue of cIBBs in the nuclear export complex.** The free energy changes of nonBonded interaction, electrostatic interaction, total of them and solvation for each residue of cIBBs due to interaction with RAN/CAS/ARM (A), CAS (B), and ARM (C) in the modeled nuclear export complex were calculated. The energy unit of nonBonded interaction and electrostatic interaction are shown in (KJ / mol), and the energy unit of solvation is shown in (kcal / mol). The Heat map is presented with red (high) and blue (low) with the median in white.

### FIGURE S13

| | 1 | 2 | 3 | 4 | 5 | 6 | 7 | 8 | 9 | 10 | 11 | 12 | 13 | 14 | 15 | 16 | 17 | 18 | 19 | 20 | 21 | 22 | 23 | 24 | 25 | 26 | 27 | 28 | 29 | 30 |
|---|---|---|---|---|---|---|---|---|---|---|---|---|---|---|---|---|---|---|---|---|---|---|---|---|---|---|---|---|---|---|
| cIBB1: | | | | M | T | T | P | G | K | E | N | F | R | L | K | S | Y | K | N | K | S | L | N | P | D | E | M | R | R | R |
| cIBB5: | M | D | A | M | A | S | P | G | K | D | N | Y | R | M | K | S | Y | K | N | K | A | L | N | P | Q | E | M | R | R | R |
| cIBB6: | M | E | T | M | A | S | P | G | K | D | N | Y | R | M | K | S | Y | K | N | N | A | L | N | P | E | E | M | R | R | R |
| cIBB3: | | | M | A | E | N | P | G | L | E | N | H | R | I | K | S | F | K | N | K | G | R | D | V | E | T | M | R | R | H |
| cIBB4: | | | M | A | D | N | E | K | L | D | N | Q | R | L | K | N | F | K | N | K | G | R | D | L | E | T | M | R | R | Q |
| cIBB2: | | | M | S | T | N | E | N | A | N | P | A | R | L | N | R | F | K | N | K | G | K | D | S | T | E | M | R | R | R |
| cIBB7: | | | | M | P | T | L | D | A | P | E | E | R | L | R | K | F | K | Y | R | G | K | D | A | S | M | R | R | Q | Q |
| nonBonded | 1 | 2 | 3 | 4 | 5 | 6 | 7 | 8 | 9 | 10 | 11 | 12 | 13 | 14 | 15 | 16 | 17 | 18 | 19 | 20 | 21 | 22 | 23 | 24 | 25 | 26 | 27 | 28 | 29 | 30 |
| cIBB1: | | | | | | | | | | | | | | | | | | | | | | | | | | | | | | |
| cIBB5: | | | | | | | | | | | | | | | | | | | | | | | | | | | | | | |
| cIBB6: | | | | | | | | | | | | | | | | | | | | | | | | | | | | | | |
| cIBB3: | | | | | | | | | | | | | | | | | | | | | | | | | | | | | | |
| cIBB4: | | | | | | | | | | | | | | | | | | | | | | | | | | | | | | |
| cIBB2: | | | | | | | | | | | | | | | | | | | | | | | | | | | | | | |
| cIBB7: | | | | | | | | | | | | | | | | | | | | | | | | | | | | | | |
| electrostatic | 1 | 2 | 3 | 4 | 5 | 6 | 7 | 8 | 9 | 10 | 11 | 12 | 13 | 14 | 15 | 16 | 17 | 18 | 19 | 20 | 21 | 22 | 23 | 24 | 25 | 26 | 27 | 28 | 29 | 30 |
| cIBB1: | | | | | | | | | | | | | | | | | | | | | | | | | | | | | | |
| cIBB5: | | | | | | | | | | | | | | | | | | | | | | | | | | | | | | |
| cIBB6: | | | | | | | | | | | | | | | | | | | | | | | | | | | | | | |
| cIBB3: | | | | | | | | | | | | | | | | | | | | | | | | | | | | | | |
| cIBB4: | | | | | | | | | | | | | | | | | | | | | | | | | | | | | | |
| cIBB2: | | | | | | | | | | | | | | | | | | | | | | | | | | | | | | |
| cIBB7: | | | | | | | | | | | | | | | | | | | | | | | | | | | | | | |
| TOTAL | 1 | 2 | 3 | 4 | 5 | 6 | 7 | 8 | 9 | 10 | 11 | 12 | 13 | 14 | 15 | 16 | 17 | 18 | 19 | 20 | 21 | 22 | 23 | 24 | 25 | 26 | 27 | 28 | 29 | 30 |
| cIBB1: | | | | | | | | | | | | | | | | | | | | | | | | | | | | | | |
| cIBB5: | | | | | | | | | | | | | | | | | | | | | | | | | | | | | | |
| cIBB6: | | | | | | | | | | | | | | | | | | | | | | | | | | | | | | |
| cIBB3: | | | | | | | | | | | | | | | | | | | | | | | | | | | | | | |
| cIBB4: | | | | | | | | | | | | | | | | | | | | | | | | | | | | | | |
| cIBB2: | | | | | | | | | | | | | | | | | | | | | | | | | | | | | | |
| cIBB7: | | | | | | | | | | | | | | | | | | | | | | | | | | | | | | |
| PISA | 1 | 2 | 3 | 4 | 5 | 6 | 7 | 8 | 9 | 10 | 11 | 12 | 13 | 14 | 15 | 16 | 17 | 18 | 19 | 20 | 21 | 22 | 23 | 24 | 25 | 26 | 27 | 28 | 29 | 30 |
| cIBB1: | | | | | | | | | | | | | | | | | | | | | | | | | | | | | | |
| cIBB5: | | | | | | | | | | | | | | | | | | | | | | | | | | | | | | |
| cIBB6: | | | | | | | | | | | | | | | | | | | | | | | | | | | | | | |
| cIBB3: | | | | | | | | | | | | | | | | | | | | | | | | | | | | | | |
| cIBB4: | | | | | | | | | | | | | | | | | | | | | | | | | | | | | | |
| cIBB2: | | | | | | | | | | | | | | | | | | | | | | | | | | | | | | |
| cIBB7: | | | | | | | | | | | | | | | | | | | | | | | | | | | | | | |

| | 31 | 32 | 33 | 34 | 35 | 36 | 37 | 38 | 39 | 40 | 41 | 42 | 43 | 44 | 45 | 46 | 47 | 48 | 49 | 50 | 51 | 52 | 53 | 54 | 55 | 56 | 57 | 58 | 59 | 60 |
|---|---|---|---|---|---|---|---|---|---|---|---|---|---|---|---|---|---|---|---|---|---|---|---|---|---|---|---|---|---|---|
| cIBB1: | R | E | E | E | G | L | Q | L | R | K | Q | K | R | E | E | Q | L | F | K | R | R | N | V | A | T | A | E | E | E | T |
| cIBB5: | R | E | E | E | G | I | Q | L | R | K | Q | K | R | E | E | Q | L | F | K | R | R | N | V | S | L | P | R | N | D | E |
| cIBB6: | R | E | E | E | G | I | Q | L | R | K | Q | K | R | E | Q | Q | L | F | K | R | R | N | V | E | L | I | N | E | E | A |
| cIBB3: | R | N | E | V | T | V | E | L | R | K | N | K | R | D | E | H | L | L | K | K | R | N | V | P | Q | E | E | S | L | E |
| cIBB4: | R | N | E | V | V | V | E | L | R | K | N | K | R | D | E | H | L | L | K | R | R | N | V | P | H | E | D | I | C | E |
| cIBB2: | R | I | E | V | N | V | E | L | R | K | A | K | K | D | D | Q | M | L | K | R | R | N | V | S | S | F | P | D | D | A |
| cIBB7: | R | I | A | V | S | L | E | L | R | K | A | K | K | D | E | Q | A | L | K | R | R | N | I | T | S | F | S | P | D | P |
| nonBonded | 31 | 32 | 33 | 34 | 35 | 36 | 37 | 38 | 39 | 40 | 41 | 42 | 43 | 44 | 45 | 46 | 47 | 48 | 49 | 50 | 51 | 52 | 53 | 54 | 55 | 56 | 57 | 58 | 59 | 60 |
| cIBB1: | | | | | | | | | | | | | | -9.7 | -12.5 | -17.3 | -8.6 | -21.0 | -24.6 | -28.7 | -24.6 | -27.6 | -16.1 | -6.6 | | | | | | |
| cIBB5: | | | | | | | | | | | | | | -10.9 | -12.4 | -18.6 | -8.7 | -21.3 | -26.6 | -29.1 | -22.1 | -23.5 | -17.4 | -4.7 | | | | | | |
| cIBB6: | | | | | | | | | | | | | | -10.8 | -6.3 | -17.9 | -8.6 | -21.0 | -26.3 | -29.5 | -29.9 | -26.0 | -15.0 | -8.7 | | | | | | |
| cIBB3: | | | | | | | | | | | | | | -4.5 | -15.0 | -16.1 | -8.9 | -18.5 | -25.8 | -23.9 | -25.0 | -26.7 | -13.4 | -4.3 | | | | | | |
| cIBB4: | | | | | | | | | | | | | | -5.0 | -15.1 | -18.1 | -9.3 | -19.5 | -25.7 | -25.2 | -26.2 | -28.0 | -12.0 | -6.8 | | | | | | |
| cIBB2: | | | | | | | | | | | | | | -9.3 | -6.2 | -21.1 | -8.6 | -18.0 | -23.3 | -25.0 | -25.5 | -27.1 | -15.8 | -8.3 | | | | | | |
| cIBB7: | | | | | | | | | | | | | | -8.6 | -14.7 | -18.7 | -7.5 | -18.6 | -24.3 | -25.5 | -23.4 | -24.6 | -18.1 | -7.1 | | | | | | |
| electrostatic | 31 | 32 | 33 | 34 | 35 | 36 | 37 | 38 | 39 | 40 | 41 | 42 | 43 | 44 | 45 | 46 | 47 | 48 | 49 | 50 | 51 | 52 | 53 | 54 | 55 | 56 | 57 | 58 | 59 | 60 |
| cIBB1: | | | | | | | | | | | | | | 2.2 | -5.0 | -13.0 | -5.1 | -2.8 | -10.5 | -22.5 | -13.3 | -15.0 | 2.0 | 0.8 | | | | | | |
| cIBB5: | | | | | | | | | | | | | | -0.5 | -4.5 | -13.3 | -5.0 | -2.5 | -10.1 | -22.3 | -14.1 | -26.1 | -1.3 | -1.6 | | | | | | |
| cIBB6: | | | | | | | | | | | | | | 2.1 | 0.0 | -13.1 | -4.9 | -2.4 | -10.7 | -22.1 | -11.2 | -12.5 | 2.2 | 0.9 | | | | | | |
| cIBB3: | | | | | | | | | | | | | | 2.2 | -4.8 | 4.0 | -4.7 | -4.9 | -11.3 | -20.1 | -11.6 | -10.7 | 1.2 | -3.2 | | | | | | |
| cIBB4: | | | | | | | | | | | | | | 0.6 | -4.7 | 4.1 | -4.1 | -5.1 | -11.8 | -22.3 | -11.0 | -8.9 | -2.4 | 2.8 | | | | | | |
| cIBB2: | | | | | | | | | | | | | | -0.7 | -1.3 | -3.0 | -4.0 | -4.1 | -11.4 | -23.3 | -11.9 | -13.5 | 1.5 | -1.6 | | | | | | |
| cIBB7: | | | | | | | | | | | | | | 1.4 | -4.3 | -11.1 | -4.7 | -4.8 | -11.0 | -22.8 | -13.1 | -25.2 | -2.4 | 19.0 | | | | | | |
| TOTAL | 31 | 32 | 33 | 34 | 35 | 36 | 37 | 38 | 39 | 40 | 41 | 42 | 43 | 44 | 45 | 46 | 47 | 48 | 49 | 50 | 51 | 52 | 53 | 54 | 55 | 56 | 57 | 58 | 59 | 60 |
| cIBB1: | | | | | | | | | | | | | | -7.5 | -17.4 | -30.3 | -13.8 | -23.7 | -35.0 | -51.3 | -37.9 | -42.6 | -14.1 | -5.8 | | | | | | |
| cIBB5: | | | | | | | | | | | | | | -11.5 | -16.8 | -31.8 | -13.7 | -23.8 | -36.7 | -51.4 | -36.2 | -49.6 | -18.7 | -6.2 | | | | | | |
| cIBB6: | | | | | | | | | | | | | | -8.7 | -6.3 | -30.9 | -13.5 | -23.4 | -37.1 | -51.5 | -41.1 | -38.6 | -12.9 | -7.7 | | | | | | |
| cIBB3: | | | | | | | | | | | | | | -2.4 | -19.7 | -12.0 | -13.6 | -23.4 | -37.0 | -44.0 | -36.5 | -37.4 | -12.3 | -7.5 | | | | | | |
| cIBB4: | | | | | | | | | | | | | | -4.4 | -19.8 | -13.9 | -13.4 | -24.6 | -37.5 | -47.4 | -37.2 | -36.9 | -14.4 | -4.0 | | | | | | |
| cIBB2: | | | | | | | | | | | | | | -10.0 | -7.4 | -24.1 | -12.6 | -22.1 | -34.6 | -48.4 | -37.4 | -40.5 | -14.4 | -9.9 | | | | | | |
| cIBB7: | | | | | | | | | | | | | | -7.1 | -19.0 | -29.8 | -12.1 | -23.4 | -35.3 | -48.3 | -36.4 | -49.8 | -20.5 | 14.5 | | | | | | |
| PISA | 31 | 32 | 33 | 34 | 35 | 36 | 37 | 38 | 39 | 40 | 41 | 42 | 43 | 44 | 45 | 46 | 47 | 48 | 49 | 50 | 51 | 52 | 53 | 54 | 55 | 56 | 57 | 58 | 59 | 60 |
| cIBB1: | | | | | | | | | | | | | | -0.7 | -0.3 | -0.7 | -0.1 | 0.4 | 0.0 | -1.7 | -0.8 | -0.4 | 0.4 | -0.1 | | | | | | |
| cIBB5: | | | | | | | | | | | | | | -0.7 | -0.3 | -0.7 | -0.1 | 0.4 | -0.1 | -1.7 | -0.5 | -0.4 | 0.9 | 0.2 | | | | | | |
| cIBB6: | | | | | | | | | | | | | | 0.3 | 0.1 | -0.7 | -0.1 | 0.4 | 0.0 | -1.8 | -0.8 | -0.2 | 0.5 | -0.2 | | | | | | |
| cIBB3: | | | | | | | | | | | | | | -0.3 | -0.5 | 0.5 | -0.1 | 0.5 | -0.1 | -0.5 | -0.8 | -0.4 | 0.3 | 0.7 | | | | | | |
| cIBB4: | | | | | | | | | | | | | | -0.3 | -0.5 | 1.0 | -0.1 | 0.7 | 0.0 | -1.7 | -0.8 | -0.3 | 0.4 | 0.4 | | | | | | |
| cIBB2: | | | | | | | | | | | | | | -0.3 | 0.2 | -0.7 | -0.1 | 0.5 | 0.0 | -1.8 | -0.9 | -0.3 | 0.4 | -0.2 | | | | | | |
| cIBB7: | | | | | | | | | | | | | | -0.1 | -0.5 | -0.7 | -0.1 | 0.5 | 0.0 | -1.7 | -0.7 | -0.4 | 0.6 | 0.2 | | | | | | |

**FIGURE S13 The free energy change for each residue of cIBBs in the autoinhibition form.** The free energy changes of nonBonded interaction, electrostatic interaction, total of them and solvation for each residue of cIBBs due to interaction with ARM in the modeled autoinhibition form were calculated. The energy unit of nonBonded interaction and electrostatic interaction are shown in (KJ / mol), and the energy unit of solvation is shown in (kcal / mol). The Heat map is presented with red (high) and blue (low) with the median in white.

### FIGURE S14

| | I | II | IIa | IIb | III |
|---|---|---|---|---|---|
| cIBB1 | -0.82 | -0.83 | -0.92 | -0.92 | -0.74 |
| cIBB5 | -0.83 | -0.85 | -0.90 | -0.94 | -0.76 |
| cIBB6 | -0.88 | -0.84 | -0.94 | -0.93 | -0.85 |
| cIBB3 | -0.84 | -0.59 | -0.66 | -0.89 | -0.54 |
| cIBB4 | -0.83 | -0.86 | -0.91 | -0.94 | -0.65 |
| cIBB2 | -0.75 | -0.80 | -0.87 | -0.89 | -0.80 |
| cIBB7 | -0.86 | -0.76 | -0.88 | -0.85 | -0.72 |

**FIGURE S14 Correlation between the contact area and the free energy.** The correlation coefficients between the contact area and the sum of the energy changes of the nonBonded and electrostatic interactions due to interaction with each binding partner were calculated for residue of cIBBs at position 13–54, 7–16 and 26–53, 44–54 in Fig S3 in the modeled importin β binding complex (I), nuclear export complex (II), and autoinhibition form (III), respectively. One residue at the N-terminus is excluded from the calculation for cIBB4 in the nuclear export complex structure. CAS (IIa) and ARM (IIb) interactions in the nuclear export complex is distinguished.

### FIGURE S15

## A

| | polarity | hydrophobicity | bulkiness | average flexibility | alpha-helix | beta-sheet | beta-turn | coil |
|---|---|---|---|---|---|---|---|---|
| Ala | 0 | 1.8 | 11.5 | 0.36 | 1.42 | 0.83 | 0.66 | 0.824 |
| Arg | 52 | -4.5 | 14.28 | 0.53 | 0.98 | 0.93 | 0.95 | 0.893 |
| Asn | 3.38 | -3.5 | 12.82 | 0.46 | 0.67 | 0.89 | 1.56 | 1.167 |
| Asp | 49.7 | -3.5 | 11.68 | 0.51 | 1.01 | 0.54 | 1.46 | 1.197 |
| Cys | 1.48 | 2.5 | 13.46 | 0.35 | 0.7 | 1.19 | 1.19 | 0.953 |
| Gln | 3.53 | -3.5 | 14.45 | 0.49 | 1.11 | 1.1 | 0.98 | 0.947 |
| Glu | 49.9 | -3.5 | 13.57 | 0.5 | 1.51 | 0.37 | 0.74 | 0.761 |
| Gly | 0 | -0.4 | 3.4 | 0.54 | 0.57 | 0.75 | 1.56 | 1.251 |
| His | 51.6 | -3.2 | 13.69 | 0.32 | 1 | 0.87 | 0.95 | 1.068 |
| Ile | 0.13 | 4.5 | 21.4 | 0.46 | 1.08 | 1.6 | 0.47 | 0.886 |
| Leu | 0.13 | 3.8 | 21.4 | 0.37 | 1.21 | 1.3 | 0.59 | 0.81 |
| Lys | 49.5 | -3.9 | 15.71 | 0.47 | 1.16 | 0.74 | 1.01 | 0.897 |
| Met | 1.43 | 1.9 | 16.25 | 0.3 | 1.45 | 1.05 | 0.6 | 0.81 |
| Phe | 0.35 | 2.8 | 19.8 | 0.31 | 1.13 | 1.38 | 0.6 | 0.797 |
| Pro | 1.58 | -1.6 | 17.43 | 0.51 | 0.57 | 0.55 | 1.52 | 1.54 |
| Ser | 1.67 | -0.8 | 9.47 | 0.51 | 0.77 | 0.75 | 1.43 | 1.13 |
| Thr | 1.66 | -0.7 | 15.77 | 0.44 | 0.83 | 1.19 | 0.96 | 1.148 |
| Trp | 2.1 | -0.9 | 21.67 | 0.31 | 1.08 | 1.37 | 0.96 | 0.941 |
| Tyr | 1.61 | -1.3 | 18.03 | 0.42 | 0.69 | 1.47 | 1.14 | 1.109 |
| Val | 0.13 | 4.2 | 21.57 | 0.39 | 1.06 | 1.7 | 0.5 | 0.772 |

## B

| | polarity | hydrophobicity | bulkiness | average flexibility | alpha-helix | beta-sheet | beta-turn | coil |
|---|---|---|---|---|---|---|---|---|
| Ala | 4.38 | 5.77 | 4.17 | 4.18 | 6.49 | 4.46 | 4.10 | 4.15 |
| Arg | 6.75 | 3.66 | 4.77 | 6.25 | 4.93 | 4.73 | 4.89 | 4.49 |
| Asn | 4.53 | 3.99 | 4.45 | 5.40 | 3.83 | 4.62 | 6.55 | 5.85 |
| Asp | 6.65 | 3.99 | 4.20 | 6.01 | 5.04 | 3.67 | 6.28 | 6.00 |
| Cys | 4.45 | 6.00 | 4.59 | 4.05 | 3.93 | 5.44 | 5.54 | 4.79 |
| Gln | 4.54 | 3.99 | 4.80 | 5.76 | 5.39 | 5.19 | 4.97 | 4.76 |
| Glu | 6.66 | 3.99 | 4.61 | 5.88 | 6.81 | 3.21 | 4.31 | 3.84 |
| Gly | 4.38 | 5.03 | 2.42 | 6.37 | 3.47 | 4.24 | 6.55 | 6.27 |
| His | 6.73 | 4.09 | 4.64 | 3.69 | 5.00 | 4.57 | 4.89 | 5.36 |
| Ile | 4.39 | 6.67 | 6.30 | 5.40 | 5.28 | 6.55 | 3.58 | 4.46 |
| Leu | 4.39 | 6.44 | 6.30 | 4.30 | 5.75 | 5.74 | 3.91 | 4.08 |
| Lys | 6.64 | 3.86 | 5.07 | 5.52 | 5.57 | 4.22 | 5.05 | 4.51 |
| Met | 4.44 | 5.80 | 5.19 | 3.44 | 6.60 | 5.06 | 3.93 | 4.08 |
| Phe | 4.40 | 6.10 | 5.96 | 3.57 | 5.46 | 5.95 | 3.93 | 4.02 |
| Pro | 4.45 | 4.63 | 5.45 | 6.01 | 3.47 | 3.70 | 6.44 | 7.70 |
| Ser | 4.46 | 4.90 | 3.73 | 6.01 | 4.18 | 4.24 | 6.19 | 5.67 |
| Thr | 4.46 | 4.93 | 5.09 | 5.15 | 4.40 | 5.44 | 4.91 | 5.76 |
| Trp | 4.48 | 4.86 | 6.36 | 3.57 | 5.28 | 5.93 | 4.91 | 4.73 |
| Tyr | 4.45 | 4.73 | 5.57 | 4.91 | 3.90 | 6.20 | 5.40 | 5.56 |
| Val | 4.39 | 6.57 | 6.34 | 4.54 | 5.21 | 6.82 | 3.66 | 3.89 |

**FIGURE S15 Standardization of score index for biochemical/biophysical propensities.** Score index assigned to each amino acid of polarity, hydrophobicity, bulkiness, average flexibility, alpha-helix, beta-sheet, beta-turn, and coil were used to evaluate the biochemical/biophysical propensities of the amino acids that make up the IBB domain. Each score of the index (A) was standardized for each biochemical/biophysical propensity (B).

### FIGURE S16

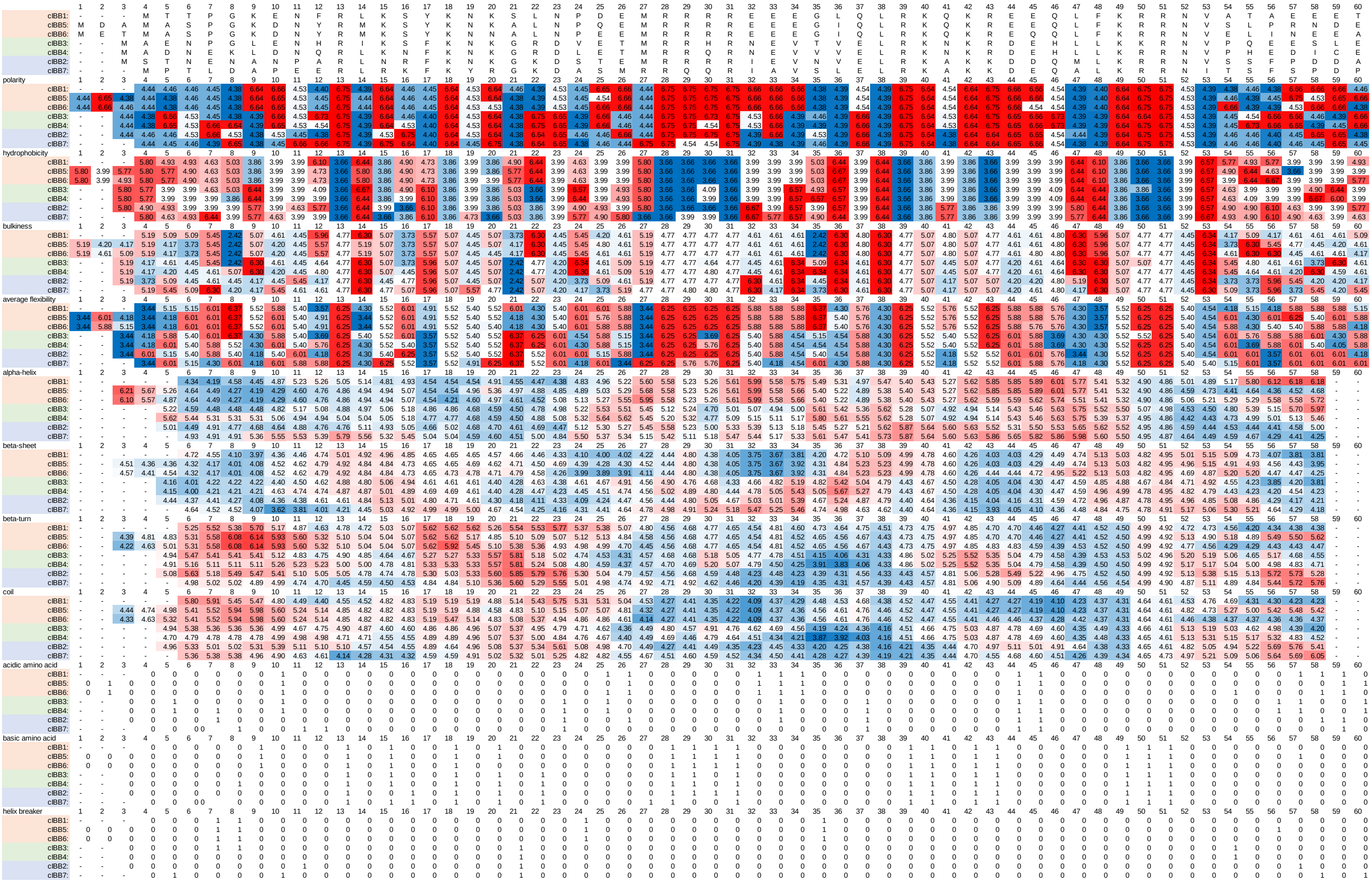

**FIGURE S16 Mapping scores of biochemical propensities to each residue of the IBB domain consensus sequence.** Each amino acid in the IBB domain consensus sequences was scored for polarity, hydrophobicity, bulkiness, average flexibility, alpha-helix, beta-sheet, beta-turn, and coil, using the normalized score index in Figure S5.

### FIGURE S17

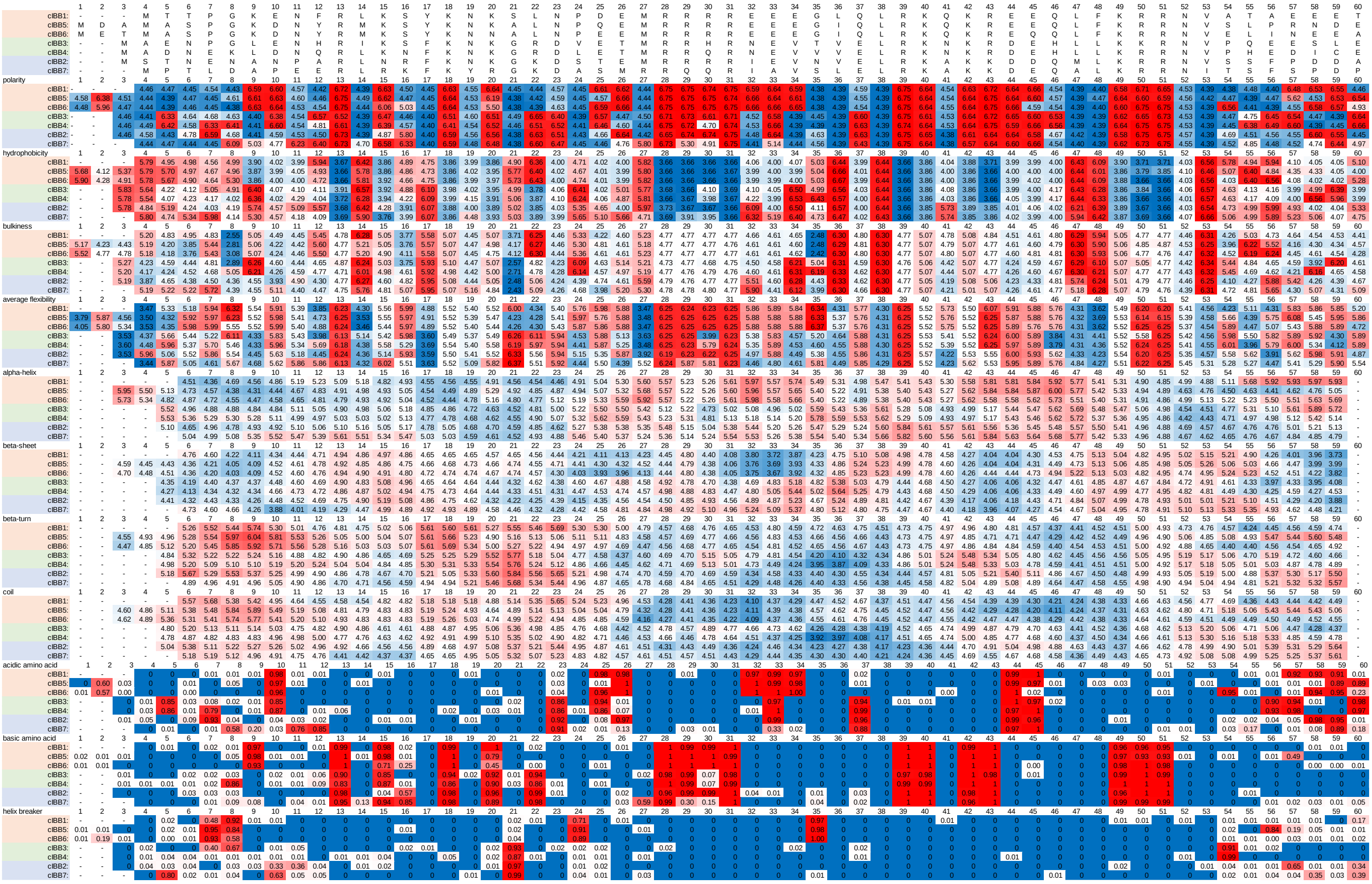

**FIGURE S17 Mapping of average scores of biochemical propensities to each residue of the IBB domain consensus sequence.** Each amino acid in the IBB domain sequence contained in the sequence set of KPNA family members was scored for polarity, hydrophobicity, bulkiness, average flexibility, alpha-helix, beta-sheet, beta-turn, and coil, using the normalized score index in Figure S5.

### FIGURE S18

| | 1 | 2 | 3 | 4 | 5 | 6 | 7 | 8 | 9 | 10 | 11 | 12 | 13 | 14 | 15 | 16 | 17 | 18 | 19 | 20 |
|---|---|---|---|---|---|---|---|---|---|---|---|---|---|---|---|---|---|---|---|---|
| cIBB1: | - | - | - | M | T | T | P | G | K | E | N | F | R | L | K | S | Y | K | N | K |
| cIBB5: | M | D | A | M | A | S | P | G | K | D | N | Y | R | M | K | S | Y | K | N | K |
| cIBB6: | M | E | T | M | A | S | P | G | K | D | N | Y | R | M | K | S | Y | K | N | N |
| cIBB3: | - | - | M | A | E | N | P | G | L | E | N | H | R | I | K | S | F | K | N | K |
| cIBB4: | - | - | M | A | D | N | E | K | L | D | N | Q | R | L | K | N | F | K | N | K |
| cIBB2: | - | - | M | S | T | N | E | N | A | N | P | A | R | L | N | R | F | K | N | K |
| cIBB7: | - | - | - | M | P | T | L | D | A | P | E | E | R | L | R | K | F | K | Y | R |
| polarity | 1 | 2 | 3 | 4 | 5 | 6 | 7 | 8 | 9 | 10 | 11 | 12 | 13 | 14 | 15 | 16 | 17 | 18 | 19 | 20 |
| cIBB1: | - | - | - | 0.18 | 0.17 | 0.02 | 0.39 | 0.31 | 0.35 | 0.35 | 0.29 | 0.26 | 0.25 | 0.01 | 0.17 | 0.30 | 0.01 | 0.16 | 0.17 | 0.01 |
| cIBB5: | 0.55 | 0.73 | 0.46 | 0.01 | 0.02 | 0.18 | 0.01 | 0.68 | 0.26 | 0.17 | 0.39 | 0.19 | 0.00 | 0.31 | 0.25 | 0.19 | 0.00 | 0.00 | 0.01 | 0.87 |
| cIBB6: | 0.36 | 1.03 | 0.15 | 0.01 | 0.02 | 0.14 | 0.00 | 0.01 | 0.14 | 0.15 | 0.01 | 0.43 | 0.01 | 0.01 | 0.94 | 1.00 | 0.01 | 0.00 | 0.00 | 1.05 |
| cIBB3: | - | - | 0.21 | 0.21 | 0.78 | 0.50 | 0.72 | 0.70 | 0.20 | 0.73 | 0.20 | 0.59 | 0.70 | 0.02 | 0.59 | 0.02 | 0.00 | 0.51 | 0.39 | 0.52 |
| cIBB4: | - | - | 0.19 | 0.46 | 0.67 | 0.37 | 0.78 | 0.68 | 0.20 | 0.33 | 0.26 | 0.73 | 0.52 | 0.02 | 0.70 | 0.32 | 0.01 | 0.67 | 0.19 | 0.50 |
| cIBB2: | - | - | 0.17 | 0.50 | 0.04 | 0.72 | 0.36 | 0.58 | 0.21 | 0.52 | 0.48 | 0.48 | 0.24 | 0.01 | 0.83 | 1.09 | 0.01 | 0.31 | 0.22 | 0.41 |
| cIBB7: | - | - | - | 0.00 | 0.23 | 0.03 | 0.33 | 0.96 | 1.02 | 0.77 | 0.87 | 0.72 | 0.23 | 0.78 | 0.54 | 0.76 | 0.01 | 0.30 | 0.04 | 0.69 |
| hydrophobicity | 1 | 2 | 3 | 4 | 5 | 6 | 7 | 8 | 9 | 10 | 11 | 12 | 13 | 14 | 15 | 16 | 17 | 18 | 19 | 20 |
| cIBB1: | - | - | - | 0.18 | 0.26 | 0.28 | 0.34 | 0.18 | 0.33 | 0.18 | 0.01 | 0.39 | 0.10 | 0.20 | 0.09 | 0.20 | 0.19 | 0.01 | 0.00 | 0.02 |
| cIBB5: | 0.58 | 0.45 | 0.73 | 0.15 | 0.27 | 0.34 | 0.26 | 0.43 | 0.18 | 0.01 | 0.22 | 0.50 | 0.00 | 0.31 | 0.16 | 0.22 | 0.00 | 0.00 | 0.22 | 0.41 |
| cIBB6: | 0.62 | 0.53 | 0.18 | 0.24 | 0.28 | 0.18 | 0.05 | 0.37 | 0.06 | 0.11 | 0.17 | 0.16 | 0.02 | 0.07 | 0.22 | 0.67 | 0.15 | 0.00 | 0.00 | 0.26 |
| cIBB3: | - | - | 0.25 | 0.36 | 0.60 | 0.44 | 0.70 | 0.33 | 0.27 | 0.36 | 0.34 | 0.32 | 0.79 | 0.32 | 0.29 | 0.23 | 0.11 | 0.54 | 0.24 | 0.41 |
| cIBB4: | - | - | 0.30 | 0.52 | 0.33 | 0.63 | 0.57 | 0.54 | 0.37 | 0.20 | 0.76 | 0.37 | 0.21 | 0.42 | 0.33 | 0.71 | 0.25 | 0.46 | 0.47 | 0.22 |
| cIBB2: | - | - | 0.24 | 0.24 | 0.50 | 0.55 | 0.28 | 0.47 | 0.52 | 0.66 | 0.61 | 0.54 | 0.23 | 0.20 | 0.57 | 0.50 | 0.20 | 0.23 | 0.07 | 0.22 |
| cIBB7: | - | - | - | 0.00 | 0.34 | 0.71 | 0.74 | 0.46 | 0.89 | 0.42 | 0.48 | 0.35 | 0.15 | 0.90 | 0.23 | 0.56 | 0.25 | 0.02 | 0.35 | 0.64 |
| bulkiness | 1 | 2 | 3 | 4 | 5 | 6 | 7 | 8 | 9 | 10 | 11 | 12 | 13 | 14 | 15 | 16 | 17 | 18 | 19 | 20 |
| cIBB1: | - | - | - | 0.10 | 0.60 | 0.47 | 0.74 | 0.47 | 0.28 | 0.25 | 0.03 | 0.82 | 0.13 | 0.15 | 0.14 | 0.27 | 0.05 | 0.02 | 0.02 | 0.02 |
| cIBB5: | 0.45 | 0.37 | 0.51 | 0.18 | 0.32 | 0.54 | 0.23 | 0.97 | 0.09 | 0.10 | 0.18 | 0.22 | 0.00 | 0.18 | 0.11 | 0.23 | 0.00 | 0.00 | 0.16 | 0.32 |
| cIBB6: | 0.67 | 0.57 | 0.58 | 0.25 | 0.30 | 0.27 | 0.20 | 0.85 | 0.04 | 0.12 | 0.12 | 0.32 | 0.03 | 0.10 | 0.28 | 0.64 | 0.04 | 0.00 | 0.00 | 0.33 |
| cIBB3: | - | - | 0.33 | 0.43 | 0.30 | 0.32 | 0.65 | 0.83 | 0.27 | 0.30 | 0.37 | 0.20 | 0.42 | 0.28 | 0.21 | 0.33 | 0.32 | 0.24 | 0.19 | 0.19 |
| cIBB4: | - | - | 0.37 | 0.31 | 0.41 | 0.57 | 0.38 | 0.35 | 0.43 | 0.23 | 0.58 | 0.14 | 0.26 | 0.79 | 0.42 | 0.50 | 0.29 | 0.57 | 0.31 | 0.32 |
| cIBB2: | - | - | 0.22 | 0.41 | 0.73 | 0.57 | 0.26 | 0.45 | 0.87 | 1.05 | 0.65 | 0.51 | 0.14 | 0.23 | 0.48 | 0.31 | 0.06 | 0.10 | 0.06 | 0.13 |
| cIBB7: | - | - | - | 0.00 | 0.57 | 0.76 | 0.91 | 0.60 | 0.63 | 0.53 | 0.58 | 0.54 | 0.11 | 0.63 | 0.22 | 0.29 | 0.08 | 0.04 | 0.60 | 0.17 |
| average flexibility | 1 | 2 | 3 | 4 | 5 | 6 | 7 | 8 | 9 | 10 | 11 | 12 | 13 | 14 | 15 | 16 | 17 | 18 | 19 | 20 |
| cIBB1: | - | - | - | 0.20 | 0.40 | 0.36 | 0.15 | 0.26 | 0.16 | 0.15 | 0.15 | 0.58 | 0.22 | 0.15 | 0.16 | 0.13 | 0.18 | 0.02 | 0.05 | 0.06 |
| cIBB5: | 0.79 | 0.49 | 0.66 | 0.31 | 0.49 | 0.38 | 0.25 | 0.40 | 0.15 | 0.20 | 0.25 | 0.45 | 0.00 | 0.44 | 0.21 | 0.24 | 0.00 | 0.00 | 0.07 | 0.17 |
| cIBB6: | 0.77 | 0.46 | 0.39 | 0.43 | 0.50 | 0.21 | 0.11 | 1.06 | 0.06 | 0.12 | 0.00 | 0.30 | 0.08 | 0.14 | 0.30 | 0.43 | 0.15 | 0.00 | 0.00 | 0.15 |
| cIBB3: | - | - | 0.33 | 0.53 | 0.67 | 0.29 | 0.88 | 0.64 | 0.18 | 0.21 | 0.29 | 0.77 | 0.46 | 0.53 | 0.45 | 0.18 | 0.27 | 0.21 | 0.26 | 0.27 |
| cIBB4: | - | - | 0.53 | 0.66 | 0.25 | 0.47 | 0.55 | 0.36 | 0.30 | 0.29 | 0.35 | 0.48 | 0.21 | 0.48 | 0.31 | 0.35 | 0.38 | 0.33 | 0.30 | 0.18 |
| cIBB2: | - | - | 0.38 | 0.23 | 0.64 | 0.41 | 0.32 | 0.32 | 0.60 | 0.66 | 0.92 | 0.64 | 0.13 | 0.33 | 0.70 | 0.41 | 0.19 | 0.20 | 0.08 | 0.19 |
| cIBB7: | - | - | - | 0.00 | 0.48 | 0.42 | 0.63 | 0.67 | 0.73 | 0.78 | 0.33 | 0.37 | 0.52 | 0.87 | 0.35 | 0.35 | 0.30 | 0.03 | 0.27 | 0.63 |
| alpha-helix | 1 | 2 | 3 | 4 | 5 | 6 | 7 | 8 | 9 | 10 | 11 | 12 | 13 | 14 | 15 | 16 | 17 | 18 | 19 | 20 |
| cIBB1: | - | - | - | 0.27 | 0.34 | 0.41 | 0.76 | 0.36 | 0.24 | 0.81 | 0.16 | 0.53 | 0.07 | 0.08 | 0.17 | 0.18 | 0.21 | 0.01 | 0.10 | 0.05 |
| cIBB5: | 0.72 | 0.45 | 1.08 | 0.35 | 0.69 | 0.35 | 0.34 | 0.92 | 0.15 | 0.21 | 0.40 | 0.52 | 0.00 | 0.26 | 0.22 | 0.23 | 0.00 | 0.00 | 0.12 | 0.69 |
| cIBB6: | 0.81 | 1.40 | 0.35 | 0.44 | 0.72 | 0.19 | 0.12 | 1.46 | 0.12 | 0.52 | 0.09 | 0.28 | 0.07 | 0.10 | 0.77 | 0.42 | 0.17 | 0.00 | 0.00 | 0.90 |
| cIBB3: | - | - | 0.37 | 0.78 | 0.83 | 0.55 | 1.44 | 0.62 | 0.21 | 0.99 | 0.32 | 0.19 | 0.16 | 0.24 | 0.28 | 0.15 | 0.18 | 0.18 | 0.29 | 0.37 |
| cIBB4: | - | - | 0.55 | 0.87 | 0.58 | 0.68 | 0.91 | 0.36 | 0.29 | 0.36 | 0.56 | 0.35 | 0.26 | 0.53 | 0.48 | 0.51 | 0.37 | 0.51 | 0.30 | 0.41 |
| cIBB2: | - | - | 0.40 | 0.62 | 0.96 | 0.54 | 0.86 | 0.69 | 0.87 | 0.86 | 1.40 | 0.94 | 0.06 | 0.24 | 0.55 | 0.39 | 0.23 | 0.02 | 0.13 | 0.29 |
| cIBB7: | - | - | - | 0.00 | 0.84 | 0.58 | 0.71 | 0.95 | 0.55 | 0.94 | 1.15 | 0.92 | 0.10 | 0.55 | 0.38 | 0.35 | 0.27 | 0.03 | 0.05 | 0.48 |
| beta-sheet | 1 | 2 | 3 | 4 | 5 | 6 | 7 | 8 | 9 | 10 | 11 | 12 | 13 | 14 | 15 | 16 | 17 | 18 | 19 | 20 |
| cIBB1: | - | - | - | 0.16 | 0.58 | 0.41 | 0.58 | 0.10 | 0.30 | 0.28 | 0.11 | 0.70 | 0.11 | 0.07 | 0.11 | 0.21 | 0.03 | 0.08 | 0.07 | 0.04 |
| cIBB5: | 0.44 | 0.48 | 0.51 | 0.17 | 0.27 | 0.41 | 0.32 | 0.50 | 0.12 | 0.16 | 0.20 | 0.20 | 0.00 | 0.28 | 0.13 | 0.11 | 0.00 | 0.00 | 0.19 | 0.34 |
| cIBB6: | 0.72 | 0.68 | 0.53 | 0.23 | 0.25 | 0.22 | 0.23 | 0.12 | 0.13 | 0.14 | 0.12 | 0.42 | 0.06 | 0.11 | 0.20 | 0.38 | 0.03 | 0.00 | 0.00 | 0.31 |
| cIBB3: | - | - | 0.33 | 0.34 | 0.73 | 0.32 | 0.54 | 0.15 | 0.36 | 0.57 | 0.23 | 0.22 | 0.46 | 0.39 | 0.29 | 0.25 | 0.16 | 0.40 | 0.11 | 0.33 |
| cIBB4: | - | - | 0.29 | 0.29 | 0.43 | 0.41 | 0.78 | 0.41 | 0.27 | 0.23 | 0.52 | 0.29 | 0.28 | 0.43 | 0.26 | 0.55 | 0.18 | 0.38 | 0.29 | 0.22 |
| cIBB2: | - | - | 0.27 | 0.28 | 0.65 | 0.44 | 0.43 | 0.39 | 0.77 | 0.50 | 0.59 | 0.47 | 0.16 | 0.27 | 0.44 | 0.41 | 0.04 | 0.20 | 0.10 | 0.15 |
| cIBB7: | - | - | - | 0.00 | 0.31 | 0.67 | 0.73 | 0.65 | 0.86 | 0.59 | 0.58 | 0.52 | 0.08 | 0.47 | 0.24 | 0.43 | 0.08 | 0.14 | 0.76 | 0.34 |
| beta-turn | 1 | 2 | 3 | 4 | 5 | 6 | 7 | 8 | 9 | 10 | 11 | 12 | 13 | 14 | 15 | 16 | 17 | 18 | 19 | 20 |
| cIBB1: | - | - | - | 0.14 | 0.60 | 0.47 | 0.59 | 0.26 | 0.23 | 0.86 | 0.13 | 0.44 | 0.01 | 0.08 | 0.14 | 0.26 | 0.20 | 0.01 | 0.02 | 0.01 |
| cIBB5: | 0.58 | 0.53 | 0.93 | 0.22 | 0.58 | 0.50 | 0.38 | 0.76 | 0.08 | 0.29 | 0.37 | 0.50 | 0.00 | 0.13 | 0.15 | 0.23 | 0.00 | 0.00 | 0.27 | 0.63 |
| cIBB6: | 0.64 | 0.94 | 0.56 | 0.38 | 0.55 | 0.26 | 0.20 | 1.19 | 0.03 | 0.58 | 0.19 | 0.19 | 0.02 | 0.02 | 0.67 | 0.68 | 0.16 | 0.00 | 0.00 | 0.79 |
| cIBB3: | - | - | 0.09 | 0.55 | 0.77 | 0.58 | 1.14 | 0.52 | 0.12 | 0.79 | 0.38 | 0.22 | 0.38 | 0.21 | 0.20 | 0.25 | 0.23 | 0.26 | 0.37 | 0.26 |
| cIBB4: | - | - | 0.42 | 0.71 | 0.58 | 0.78 | 0.67 | 0.33 | 0.27 | 0.42 | 0.85 | 0.29 | 0.46 | 0.61 | 0.34 | 0.82 | 0.39 | 0.42 | 0.45 | 0.32 |
| cIBB2: | - | - | 0.28 | 0.47 | 0.80 | 0.63 | 0.91 | 0.66 | 0.68 | 0.80 | 1.13 | 0.83 | 0.13 | 0.26 | 0.85 | 0.35 | 0.21 | 0.12 | 0.04 | 0.21 |
| cIBB7: | - | - | - | 0.00 | 0.65 | 0.69 | 0.78 | 0.89 | 0.34 | 0.79 | 0.92 | 0.77 | 0.13 | 0.34 | 0.34 | 0.38 | 0.26 | 0.01 | 0.55 | 0.29 |
| coil | 1 | 2 | 3 | 4 | 5 | 6 | 7 | 8 | 9 | 10 | 11 | 12 | 13 | 14 | 15 | 16 | 17 | 18 | 19 | 20 |
| cIBB1: | - | - | - | 0.16 | 0.45 | 0.38 | 1.26 | 0.28 | 0.18 | 0.94 | 0.04 | 0.32 | 0.03 | 0.05 | 0.15 | 0.21 | 0.21 | 0.02 | 0.01 | 0.00 |
| cIBB5: | 0.53 | 0.44 | 0.75 | 0.23 | 0.61 | 0.34 | 0.58 | 0.75 | 0.11 | 0.30 | 0.26 | 0.53 | 0.00 | 0.07 | 0.11 | 0.17 | 0.00 | 0.00 | 0.16 | 0.53 |
| cIBB6: | 0.59 | 1.55 | 0.29 | 0.27 | 0.50 | 0.15 | 0.27 | 1.02 | 0.07 | 0.63 | 0.09 | 0.14 | 0.00 | 0.02 | 0.59 | 0.56 | 0.17 | 0.00 | 0.00 | 0.69 |
| cIBB3: | - | - | 0.05 | 0.61 | 0.83 | 0.42 | 1.70 | 0.44 | 0.16 | 0.83 | 0.42 | 0.27 | 0.17 | 0.21 | 0.29 | 0.19 | 0.20 | 0.06 | 0.24 | 0.42 |
| cIBB4: | - | - | 0.31 | 0.57 | 0.61 | 0.62 | 0.72 | 0.38 | 0.28 | 0.48 | 0.49 | 0.25 | 0.46 | 0.46 | 0.54 | 0.54 | 0.28 | 0.53 | 0.27 | 0.49 |
| cIBB2: | - | - | 0.25 | 0.59 | 0.79 | 0.44 | 1.00 | 0.53 | 0.74 | 0.79 | 1.66 | 0.98 | 0.10 | 0.30 | 0.49 | 0.29 | 0.22 | 0.06 | 0.02 | 0.29 |
| cIBB7: | - | - | - | 0.00 | 1.11 | 0.77 | 0.75 | 0.87 | 0.29 | 1.32 | 0.95 | 0.84 | 0.21 | 0.15 | 0.31 | 0.26 | 0.27 | 0.04 | 0.15 | 0.21 |

| | 21 | 22 | 23 | 24 | 25 | 26 | 27 | 28 | 29 | 30 | 31 | 32 | 33 | 34 | 35 | 36 | 37 | 38 | 39 | 40 |
|---|---|---|---|---|---|---|---|---|---|---|---|---|---|---|---|---|---|---|---|---|
| cIBB1: | S | L | N | P | D | E | M | R | R | R | R | E | E | E | G | L | Q | L | R | K |
| cIBB5: | A | L | N | P | Q | E | M | R | R | R | R | E | E | E | G | I | Q | L | R | K |
| cIBB6: | A | L | N | P | E | E | M | R | R | R | R | E | E | E | G | I | Q | L | R | K |
| cIBB3: | G | R | D | V | E | T | M | R | R | H | R | N | E | V | T | V | E | L | R | K |
| cIBB4: | G | R | D | L | E | T | M | R | R | Q | R | N | E | V | V | V | E | L | R | K |
| cIBB2: | G | K | D | S | T | E | M | R | R | R | R | I | E | V | N | V | E | L | R | K |
| cIBB7: | G | K | D | A | S | M | R | R | Q | Q | R | I | A | V | S | L | E | L | R | K |
| polarity | 21 | 22 | 23 | 24 | 25 | 26 | 27 | 28 | 29 | 30 | 31 | 32 | 33 | 34 | 35 | 36 | 37 | 38 | 39 | 40 |
| cIBB1: | 0.01 | 0.35 | 0.29 | 0.01 | 0.30 | 0.30 | 0.01 | 0.00 | 0.01 | 0.17 | 0.00 | 0.39 | 0.17 | 0.39 | 0.02 | 0.00 | 0.33 | 0.00 | 0.00 | 0.00 |
| cIBB5: | 0.00 | 0.27 | 0.34 | 0.01 | 0.24 | 0.00 | 0.01 | 0.00 | 0.01 | 0.00 | 0.19 | 0.00 | 0.18 | 0.30 | 0.01 | 0.00 | 0.17 | 0.00 | 0.00 | 0.00 |
| cIBB6: | 0.01 | 0.14 | 0.43 | 0.02 | 0.36 | 0.00 | 0.01 | 0.00 | 0.00 | 0.01 | 0.00 | 0.00 | 0.00 | 0.14 | 0.00 | 0.00 | 0.00 | 0.00 | 0.00 | 0.00 |
| cIBB3: | 0.50 | 0.46 | 0.69 | 0.01 | 0.44 | 0.20 | 0.36 | 0.30 | 0.20 | 0.52 | 0.29 | 0.03 | 0.40 | 0.01 | 0.02 | 0.00 | 0.36 | 0.00 | 0.01 | 0.27 |
| cIBB4: | 0.42 | 0.71 | 0.52 | 0.19 | 0.63 | 0.55 | 0.01 | 0.00 | 0.26 | 0.58 | 0.19 | 0.18 | 0.00 | 0.01 | 0.02 | 0.00 | 0.25 | 0.00 | 0.18 | 0.00 |
| cIBB2: | 0.02 | 0.16 | 0.53 | 0.04 | 0.61 | 0.17 | 0.04 | 0.47 | 0.17 | 0.17 | 0.01 | 0.43 | 0.17 | 0.01 | 0.47 | 0.00 | 0.23 | 0.00 | 0.00 | 0.03 |
| cIBB7: | 0.01 | 0.32 | 0.58 | 0.33 | 0.23 | 0.82 | 1.16 | 0.24 | 1.05 | 0.82 | 0.02 | 0.03 | 1.07 | 0.32 | 0.45 | 0.02 | 0.65 | 0.00 | 0.00 | 0.00 |
| hydrophobicity | 21 | 22 | 23 | 24 | 25 | 26 | 27 | 28 | 29 | 30 | 31 | 32 | 33 | 34 | 35 | 36 | 37 | 38 | 39 | 40 |
| cIBB1: | 0.10 | 0.41 | 0.09 | 0.17 | 0.21 | 0.11 | 0.11 | 0.00 | 0.03 | 0.04 | 0.00 | 0.41 | 0.07 | 0.43 | 0.11 | 0.02 | 0.00 | 0.00 | 0.00 | 0.00 |
| cIBB5: | 0.15 | 0.27 | 0.23 | 0.15 | 0.21 | 0.00 | 0.10 | 0.00 | 0.02 | 0.00 | 0.10 | 0.00 | 0.08 | 0.00 | 0.13 | 0.08 | 0.21 | 0.03 | 0.00 | 0.00 |
| cIBB6: | 0.17 | 0.16 | 0.06 | 0.39 | 0.18 | 0.00 | 0.12 | 0.00 | 0.00 | 0.01 | 0.00 | 0.00 | 0.00 | 0.16 | 0.01 | 0.02 | 0.00 | 0.02 | 0.00 | 0.00 |
| cIBB3: | 0.26 | 0.51 | 0.26 | 0.41 | 0.15 | 0.39 | 0.39 | 0.17 | 0.03 | 0.35 | 0.22 | 0.32 | 0.33 | 0.30 | 0.34 | 0.12 | 0.27 | 0.02 | 0.03 | 0.03 |
| cIBB4: | 0.38 | 0.59 | 0.50 | 0.64 | 0.26 | 0.25 | 0.11 | 0.00 | 0.17 | 0.17 | 0.17 | 0.40 | 0.00 | 0.27 | 0.45 | 0.06 | 0.05 | 0.00 | 0.03 | 0.00 |
| cIBB2: | 0.12 | 0.04 | 0.27 | 0.59 | 0.61 | 0.13 | 0.52 | 0.36 | 0.17 | 0.10 | 0.01 | 0.81 | 0.13 | 0.26 | 0.33 | 0.06 | 0.13 | 0.02 | 0.00 | 0.05 |
| cIBB7: | 0.01 | 0.30 | 0.01 | 0.55 | 0.43 | 0.84 | 1.31 | 0.31 | 0.25 | 0.12 | 0.03 | 0.40 | 0.87 | 0.53 | 0.55 | 0.27 | 0.26 | 0.03 | 0.00 | 0.00 |
| bulkiness | 21 | 22 | 23 | 24 | 25 | 26 | 27 | 28 | 29 | 30 | 31 | 32 | 33 | 34 | 35 | 36 | 37 | 38 | 39 | 40 |
| cIBB1: | 0.19 | 0.36 | 0.11 | 0.23 | 0.09 | 0.18 | 0.19 | 0.00 | 0.03 | 0.04 | 0.00 | 0.27 | 0.07 | 0.27 | 0.34 | 0.01 | 0.03 | 0.00 | 0.00 | 0.00 |
| cIBB5: | 0.39 | 0.21 | 0.16 | 0.49 | 0.13 | 0.00 | 0.27 | 0.00 | 0.03 | 0.00 | 0.08 | 0.00 | 0.06 | 0.05 | 0.44 | 0.17 | 0.13 | 0.00 | 0.00 | 0.00 |
| cIBB6: | 0.31 | 0.08 | 0.07 | 0.42 | 0.11 | 0.00 | 0.21 | 0.00 | 0.00 | 0.02 | 0.00 | 0.00 | 0.00 | 0.11 | 0.08 | 0.01 | 0.00 | 0.00 | 0.00 | 0.00 |
| cIBB3: | 0.58 | 0.30 | 0.25 | 0.65 | 0.16 | 0.31 | 0.38 | 0.30 | 0.00 | 0.21 | 0.11 | 0.21 | 0.17 | 0.53 | 0.45 | 0.09 | 0.32 | 0.00 | 0.05 | 0.08 |
| cIBB4: | 0.82 | 0.33 | 0.39 | 0.49 | 0.22 | 0.34 | 0.28 | 0.00 | 0.05 | 0.05 | 0.05 | 0.28 | 0.03 | 0.21 | 0.50 | 0.04 | 0.07 | 0.00 | 0.00 | 0.00 |
| cIBB2: | 0.39 | 0.07 | 0.16 | 0.85 | 0.60 | 0.09 | 0.61 | 0.17 | 0.02 | 0.04 | 0.02 | 0.99 | 0.08 | 0.26 | 0.28 | 0.08 | 0.09 | 0.00 | 0.00 | 0.08 |
| cIBB7: | 0.13 | 0.19 | 0.13 | 0.75 | 0.50 | 0.84 | 0.70 | 0.16 | 0.08 | 0.04 | 0.05 | 0.56 | 0.46 | 0.65 | 0.52 | 0.15 | 0.19 | 0.04 | 0.00 | 0.00 |
| average flexibility | 21 | 22 | 23 | 24 | 25 | 26 | 27 | 28 | 29 | 30 | 31 | 32 | 33 | 34 | 35 | 36 | 37 | 38 | 39 | 40 |
| cIBB1: | 0.16 | 0.25 | 0.09 | 0.40 | 0.21 | 0.08 | 0.15 | 0.00 | 0.06 | 0.11 | 0.00 | 0.16 | 0.01 | 0.25 | 0.22 | 0.04 | 0.02 | 0.00 | 0.00 | 0.00 |
| cIBB5: | 0.31 | 0.10 | 0.10 | 0.24 | 0.10 | 0.00 | 0.26 | 0.00 | 0.06 | 0.00 | 0.02 | 0.00 | 0.06 | 0.07 | 0.31 | 0.17 | 0.10 | 0.13 | 0.00 | 0.00 |
| cIBB6: | 0.41 | 0.09 | 0.13 | 0.47 | 0.17 | 0.00 | 0.16 | 0.00 | 0.00 | 0.05 | 0.00 | 0.00 | 0.00 | 0.09 | 0.02 | 0.16 | 0.00 | 0.10 | 0.00 | 0.00 |
| cIBB3: | 0.52 | 0.48 | 0.25 | 0.26 | 0.07 | 0.22 | 0.60 | 0.07 | 0.04 | 0.79 | 0.20 | 0.16 | 0.34 | 0.34 | 0.27 | 0.42 | 0.14 | 0.10 | 0.02 | 0.08 |
| cIBB4: | 0.58 | 0.68 | 0.31 | 0.40 | 0.18 | 0.28 | 0.27 | 0.00 | 0.17 | 0.18 | 0.17 | 0.13 | 0.01 | 0.15 | 0.26 | 0.17 | 0.01 | 0.00 | 0.04 | 0.00 |
| cIBB2: | 0.28 | 0.16 | 0.29 | 0.82 | 0.39 | 0.14 | 0.61 | 0.28 | 0.25 | 0.22 | 0.05 | 0.64 | 0.13 | 0.29 | 0.38 | 0.15 | 0.14 | 0.11 | 0.00 | 0.18 |
| cIBB7: | 0.04 | 0.26 | 0.28 | 0.60 | 0.78 | 0.80 | 0.98 | 0.09 | 0.40 | 0.28 | 0.13 | 0.83 | 0.81 | 0.40 | 0.42 | 0.34 | 0.15 | 0.07 | 0.00 | 0.00 |
| alpha-helix | 21 | 22 | 23 | 24 | 25 | 26 | 27 | 28 | 29 | 30 | 31 | 32 | 33 | 34 | 35 | 36 | 37 | 38 | 39 | 40 |
| cIBB1: | 0.21 | 0.18 | 0.17 | 0.48 | 0.29 | 0.41 | 0.15 | 0.00 | 0.16 | 0.09 | 0.00 | 0.27 | 0.21 | 0.33 | 0.12 | 0.09 | 0.22 | 0.00 | 0.00 | 0.00 |
| cIBB5: | 0.46 | 0.11 | 0.23 | 0.40 | 0.16 | 0.00 | 0.27 | 0.00 | 0.05 | 0.00 | 0.06 | 0.00 | 0.28 | 0.46 | 0.32 | 0.10 | 0.12 | 0.05 | 0.00 | 0.00 |
| cIBB6: | 0.58 | 0.05 | 0.25 | 0.74 | 0.44 | 0.00 | 0.16 | 0.00 | 0.00 | 0.04 | 0.00 | 0.00 | 0.00 | 0.15 | 0.05 | 0.01 | 0.00 | 0.04 | 0.00 | 0.00 |
| cIBB3: | 0.52 | 0.20 | 0.38 | 0.47 | 0.62 | 0.35 | 0.48 | 0.19 | 0.04 | 0.09 | 0.17 | 0.34 | 0.47 | 0.36 | 0.20 | 0.08 | 0.45 | 0.04 | 0.01 | 0.20 |
| cIBB4: | 0.46 | 0.40 | 0.20 | 0.29 | 0.79 | 0.63 | 0.28 | 0.00 | 0.13 | 0.15 | 0.13 | 0.34 | 0.15 | 0.23 | 0.26 | 0.03 | 0.30 | 0.00 | 0.04 | 0.00 |
| cIBB2: | 0.26 | 0.14 | 0.29 | 1.02 | 0.63 | 0.29 | 0.56 | 0.13 | 0.07 | 0.08 | 0.05 | 0.68 | 0.27 | 0.15 | 0.32 | 0.10 | 0.27 | 0.05 | 0.00 | 0.16 |
| cIBB7: | 0.07 | 0.12 | 0.59 | 0.91 | 1.00 | 0.68 | 0.57 | 0.04 | 0.34 | 0.22 | 0.11 | 0.64 | 0.47 | 0.40 | 0.31 | 0.25 | 0.47 | 0.03 | 0.00 | 0.00 |
| beta-sheet | 21 | 22 | 23 | 24 | 25 | 26 | 27 | 28 | 29 | 30 | 31 | 32 | 33 | 34 | 35 | 36 | 37 | 38 | 39 | 40 |
| cIBB1: | 0.03 | 0.26 | 0.19 | 0.78 | 0.18 | 0.23 | 0.12 | 0.00 | 0.13 | 0.08 | 0.00 | 0.56 | 0.08 | 0.59 | 0.14 | 0.19 | 0.31 | 0.00 | 0.00 | 0.00 |
| cIBB5: | 0.33 | 0.14 | 0.23 | 0.28 | 0.26 | 0.00 | 0.11 | 0.00 | 0.04 | 0.00 | 0.04 | 0.00 | 0.19 | 0.20 | 0.17 | 0.18 | 0.21 | 0.09 | 0.00 | 0.00 |
| cIBB6: | 0.08 | 0.14 | 0.20 | 0.41 | 0.34 | 0.00 | 0.13 | 0.00 | 0.00 | 0.03 | 0.00 | 0.00 | 0.00 | 0.23 | 0.00 | 0.05 | 0.00 | 0.07 | 0.00 | 0.00 |
| cIBB3: | 0.20 | 0.27 | 0.35 | 0.81 | 0.24 | 0.38 | 0.23 | 0.08 | 0.04 | 0.28 | 0.07 | 0.25 | 0.33 | 0.58 | 0.43 | 0.23 | 0.35 | 0.07 | 0.10 | 0.14 |
| cIBB4: | 0.40 | 0.29 | 0.50 | 0.30 | 0.45 | 0.61 | 0.12 | 0.00 | 0.04 | 0.13 | 0.02 | 0.37 | 0.04 | 0.29 | 0.52 | 0.11 | 0.17 | 0.00 | 0.04 | 0.00 |
| cIBB2: | 0.13 | 0.13 | 0.39 | 0.84 | 0.67 | 0.18 | 0.36 | 0.16 | 0.05 | 0.08 | 0.04 | 1.02 | 0.13 | 0.34 | 0.23 | 0.13 | 0.25 | 0.08 | 0.00 | 0.13 |
| cIBB7: | 0.00 | 0.33 | 0.31 | 0.75 | 0.44 | 0.96 | 0.60 | 0.18 | 0.36 | 0.27 | 0.09 | 0.78 | 0.80 | 0.81 | 0.40 | 0.51 | 0.68 | 0.02 | 0.00 | 0.00 |
| beta-turn | 21 | 22 | 23 | 24 | 25 | 26 | 27 | 28 | 29 | 30 | 31 | 32 | 33 | 34 | 35 | 36 | 37 | 38 | 39 | 40 |
| cIBB1: | 0.18 | 0.27 | 0.16 | 0.69 | 0.37 | 0.26 | 0.00 | 0.00 | 0.05 | 0.02 | 0.00 | 0.12 | 0.15 | 0.14 | 0.16 | 0.04 | 0.10 | 0.00 | 0.00 | 0.00 |
| cIBB5: | 0.35 | 0.11 | 0.28 | 0.34 | 0.13 | 0.00 | 0.21 | 0.00 | 0.01 | 0.00 | 0.11 | 0.00 | 0.23 | 0.38 | 0.33 | 0.04 | 0.12 | 0.04 | 0.00 | 0.00 |
| cIBB6: | 0.47 | 0.08 | 0.06 | 0.65 | 0.38 | 0.00 | 0.01 | 0.00 | 0.00 | 0.01 | 0.00 | 0.00 | 0.00 | 0.13 | 0.02 | 0.02 | 0.00 | 0.03 | 0.00 | 0.00 |
| cIBB3: | 0.47 | 0.23 | 0.30 | 0.42 | 0.41 | 0.32 | 0.37 | 0.21 | 0.01 | 0.17 | 0.14 | 0.54 | 0.51 | 0.43 | 0.43 | 0.08 | 0.29 | 0.03 | 0.13 | 0.15 |
| cIBB4: | 0.51 | 0.29 | 0.52 | 0.40 | 0.59 | 0.36 | 0.21 | 0.00 | 0.07 | 0.07 | 0.06 | 0.71 | 0.16 | 0.28 | 0.46 | 0.03 | 0.18 | 0.00 | 0.01 | 0.00 |
| cIBB2: | 0.22 | 0.04 | 0.17 | 1.04 | 0.87 | 0.27 | 0.38 | 0.10 | 0.05 | 0.02 | 0.01 | 0.71 | 0.29 | 0.25 | 0.46 | 0.02 | 0.20 | 0.03 | 0.00 | 0.04 |
| cIBB7: | 0.04 | 0.14 | 0.54 | 0.57 | 0.93 | 0.49 | 0.60 | 0.13 | 0.20 | 0.07 | 0.03 | 0.16 | 0.34 | 0.53 | 0.50 | 0.18 | 0.23 | 0.00 | 0.00 | 0.00 |
| coil | 21 | 22 | 23 | 24 | 25 | 26 | 27 | 28 | 29 | 30 | 31 | 32 | 33 | 34 | 35 | 36 | 37 | 38 | 39 | 40 |
| cIBB1: | 0.15 | 0.18 | 0.03 | 0.92 | 0.35 | 0.29 | 0.00 | 0.00 | 0.05 | 0.02 | 0.00 | 0.10 | 0.14 | 0.08 | 0.14 | 0.03 | 0.14 | 0.00 | 0.00 | 0.00 |
| cIBB5: | 0.30 | 0.15 | 0.12 | 0.68 | 0.13 | 0.00 | 0.18 | 0.00 | 0.00 | 0.00 | 0.10 | 0.00 | 0.29 | 0.37 | 0.25 | 0.09 | 0.10 | 0.04 | 0.00 | 0.00 |
| cIBB6: | 0.48 | 0.03 | 0.03 | 1.01 | 0.45 | 0.00 | 0.00 | 0.00 | 0.00 | 0.00 | 0.00 | 0.00 | 0.00 | 0.14 | 0.04 | 0.11 | 0.00 | 0.03 | 0.00 | 0.00 |
| cIBB3: | 0.40 | 0.17 | 0.24 | 0.56 | 0.67 | 0.44 | 0.25 | 0.22 | 0.02 | 0.27 | 0.11 | 0.20 | 0.56 | 0.44 | 0.38 | 0.22 | 0.31 | 0.03 | 0.14 | 0.14 |
| cIBB4: | 0.53 | 0.31 | 0.38 | 0.39 | 0.69 | 0.50 | 0.18 | 0.00 | 0.04 | 0.09 | 0.03 | 0.17 | 0.18 | 0.21 | 0.50 | 0.08 | 0.32 | 0.00 | 0.02 | 0.00 |
| cIBB2: | 0.20 | 0.02 | 0.19 | 0.81 | 0.48 | 0.29 | 0.24 | 0.23 | 0.07 | 0.11 | 0.00 | 0.44 | 0.32 | 0.27 | 0.22 | 0.08 | 0.21 | 0.04 | 0.00 | 0.00 |
| cIBB7: | 0.06 | 0.07 | 0.56 | 0.87 | 0.68 | 0.48 | 0.61 | 0.00 | 0.24 | 0.15 | 0.00 | 0.23 | 0.34 | 0.40 | 0.35 | 0.17 | 0.28 | 0.01 | 0.00 | 0.00 |

| | 41 | 42 | 43 | 44 | 45 | 46 | 47 | 48 | 49 | 50 | 51 | 52 | 53 | 54 | 55 | 56 | 57 | 58 | 59 | 60 |
|---|---|---|---|---|---|---|---|---|---|---|---|---|---|---|---|---|---|---|---|---|
| cIBB1: | Q | K | R | E | E | Q | L | F | K | R | R | N | V | A | T | A | E | E | E | T |
| cIBB5: | Q | K | R | E | E | Q | L | F | K | R | R | N | V | S | L | P | R | N | D | E |
| cIBB6: | Q | K | R | E | Q | Q | L | F | K | R | R | N | V | E | L | I | N | E | E | A |
| cIBB3: | N | K | R | D | E | H | L | L | K | K | R | N | V | P | Q | E | E | S | L | E |
| cIBB4: | N | K | R | D | E | H | L | L | K | R | R | N | V | P | H | E | D | I | C | E |
| cIBB2: | A | K | K | D | D | Q | M | L | K | R | R | N | V | S | S | F | P | D | D | A |
| cIBB7: | A | K | K | D | E | Q | A | L | K | R | R | N | I | T | S | F | S | P | D | P |
| polarity | 41 | 42 | 43 | 44 | 45 | 46 | 47 | 48 | 49 | 50 | 51 | 52 | 53 | 54 | 55 | 56 | 57 | 58 | 59 | 60 |
| cIBB1: | 0.03 | 0.18 | 0.05 | 0.17 | 0.00 | 0.01 | 0.01 | 0.01 | 0.34 | 0.32 | 0.46 | 0.02 | 0.01 | 0.01 | 0.24 | 0.18 | 0.60 | 0.51 | 0.47 | 0.25 |
| cIBB5: | 0.01 | 0.00 | 0.00 | 0.19 | 0.35 | 0.24 | 0.00 | 0.41 | 0.00 | 0.58 | 0.59 | 0.25 | 0.26 | 0.19 | 0.02 | 0.25 | 1.16 | 0.25 | 0.50 | 0.49 |
| cIBB6: | 0.13 | 0.00 | 0.00 | 0.00 | 0.33 | 0.01 | 0.00 | 0.01 | 0.27 | 0.00 | 0.01 | 0.02 | 0.01 | 0.45 | 0.20 | 0.01 | 0.24 | 0.39 | 0.43 | 0.97 |
| cIBB3: | 0.03 | 0.01 | 0.28 | 0.00 | 0.33 | 0.65 | 0.01 | 0.03 | 0.19 | 0.03 | 0.20 | 0.02 | 0.00 | 0.21 | 0.66 | 0.64 | 0.49 | 0.20 | 0.02 | 0.19 |
| cIBB4: | 0.02 | 0.00 | 0.00 | 0.37 | 0.00 | 0.60 | 0.00 | 0.03 | 0.00 | 0.02 | 0.00 | 0.01 | 0.00 | 0.00 | 0.81 | 0.58 | 0.31 | 0.02 | 0.01 | 0.00 |
| cIBB2: | 0.02 | 0.28 | 0.02 | 0.16 | 0.39 | 0.52 | 0.03 | 0.01 | 0.36 | 0.03 | 0.00 | 0.28 | 0.00 | 0.67 | 0.37 | 0.59 | 0.48 | 0.33 | 0.47 | 0.29 |
| cIBB7: | 0.02 | 0.43 | 0.01 | 0.31 | 0.00 | 0.02 | 0.03 | 0.01 | 0.22 | 0.24 | 0.01 | 0.22 | 0.00 | 0.39 | 0.85 | 0.41 | 0.42 | 0.77 | 0.64 | 0.94 |
| hydrophobicity | 41 | 42 | 43 | 44 | 45 | 46 | 47 | 48 | 49 | 50 | 51 | 52 | 53 | 54 | 55 | 56 | 57 | 58 | 59 | 60 |
| cIBB1: | 0.26 | 0.20 | 0.09 | 0.00 | 0.00 | 0.07 | 0.09 | 0.15 | 0.25 | 0.33 | 0.33 | 0.24 | 0.08 | 0.23 | 0.28 | 0.42 | 0.47 | 0.28 | 0.30 | 0.38 |
| cIBB5: | 0.07 | 0.00 | 0.00 | 0.14 | 0.10 | 0.07 | 0.03 | 0.42 | 0.02 | 0.55 | 0.72 | 0.46 | 0.43 | 0.46 | 0.33 | 0.58 | 0.76 | 0.60 | 0.32 | 0.09 |
| cIBB6: | 0.13 | 0.00 | 0.00 | 0.00 | 0.28 | 0.16 | 0.05 | 0.20 | 0.13 | 0.00 | 0.03 | 0.28 | 0.05 | 0.26 | 0.35 | 0.28 | 0.30 | 0.16 | 0.21 | 0.77 |
| cIBB3: | 0.34 | 0.02 | 0.05 | 0.00 | 0.08 | 0.27 | 0.14 | 0.51 | 0.01 | 0.05 | 0.03 | 0.30 | 0.03 | 0.06 | 0.54 | 0.61 | 0.00 | 0.47 | 0.22 | 0.00 |
| cIBB4: | 0.24 | 0.00 | 0.00 | 0.37 | 0.00 | 0.32 | 0.04 | 0.47 | 0.00 | 0.03 | 0.00 | 0.11 | 0.07 | 0.00 | 0.41 | 0.41 | 0.10 | 0.41 | 0.24 | 0.00 |
| cIBB2: | 0.23 | 0.34 | 0.04 | 0.19 | 0.42 | 0.25 | 0.42 | 0.15 | 0.17 | 0.05 | 0.00 | 0.26 | 0.18 | 0.59 | 0.45 | 0.61 | 0.76 | 0.22 | 0.29 | 0.61 |
| cIBB7: | 0.19 | 0.21 | 0.02 | 0.26 | 0.00 | 0.11 | 0.66 | 0.09 | 0.11 | 0.28 | 0.03 | 0.25 | 0.04 | 0.53 | 1.00 | 0.79 | 0.69 | 0.80 | 0.29 | 0.69 |
| bulkiness | 41 | 42 | 43 | 44 | 45 | 46 | 47 | 48 | 49 | 50 | 51 | 52 | 53 | 54 | 55 | 56 | 57 | 58 | 59 | 60 |
| cIBB1: | 0.21 | 0.10 | 0.13 | 0.18 | 0.00 | 0.08 | 0.13 | 0.26 | 0.24 | 0.24 | 0.18 | 0.21 | 0.20 | 0.52 | 0.40 | 0.94 | 0.33 | 0.25 | 0.21 | 0.98 |
| cIBB5: | 0.09 | 0.00 | 0.00 | 0.04 | 0.10 | 0.09 | 0.01 | 0.31 | 0.08 | 0.31 | 0.39 | 0.32 | 0.38 | 0.63 | 0.38 | 0.29 | 0.96 | 0.53 | 0.32 | 0.21 |
| cIBB6: | 0.05 | 0.00 | 0.00 | 0.06 | 0.17 | 0.10 | 0.07 | 0.27 | 0.12 | 0.00 | 0.01 | 0.18 | 0.16 | 0.29 | 0.44 | 0.33 | 0.34 | 0.13 | 0.22 | 0.36 |
| cIBB3: | 0.24 | 0.03 | 0.03 | 0.12 | 0.13 | 0.15 | 0.11 | 0.64 | 0.06 | 0.08 | 0.00 | 0.24 | 0.01 | 0.08 | 0.40 | 0.54 | 0.12 | 0.64 | 0.39 | 0.05 |
| cIBB4: | 0.10 | 0.00 | 0.00 | 0.35 | 0.07 | 0.19 | 0.03 | 0.40 | 0.00 | 0.04 | 0.00 | 0.18 | 0.18 | 0.00 | 0.28 | 0.18 | 0.12 | 0.56 | 0.26 | 0.11 |
| cIBB2: | 0.15 | 0.16 | 0.07 | 0.17 | 0.37 | 0.16 | 0.56 | 0.27 | 0.39 | 0.08 | 0.00 | 0.15 | 0.45 | 0.65 | 0.75 | 0.48 | 0.59 | 0.29 | 0.32 | 0.67 |
| cIBB7: | 0.27 | 0.15 | 0.03 | 0.24 | 0.04 | 0.25 | 1.01 | 0.13 | 0.00 | 0.16 | 0.06 | 0.20 | 0.01 | 0.74 | 0.95 | 0.88 | 0.94 | 0.86 | 0.41 | 0.61 |
| average flexibility | 41 | 42 | 43 | 44 | 45 | 46 | 47 | 48 | 49 | 50 | 51 | 52 | 53 | 54 | 55 | 56 | 57 | 58 | 59 | 60 |
| cIBB1: | 0.23 | 0.17 | 0.31 | 0.07 | 0.00 | 0.02 | 0.19 | 0.30 | 0.28 | 0.31 | 0.22 | 0.14 | 0.19 | 0.31 | 0.39 | 0.31 | 0.33 | 0.30 | 0.29 | 0.70 |
| cIBB5: | 0.04 | 0.00 | 0.00 | 0.14 | 0.04 | 0.04 | 0.10 | 0.49 | 0.07 | 0.39 | 0.41 | 0.09 | 0.25 | 0.78 | 0.32 | 0.65 | 0.47 | 0.45 | 0.18 | 0.21 |
| cIBB6: | 0.11 | 0.00 | 0.00 | 0.02 | 0.11 | 0.08 | 0.15 | 0.34 | 0.05 | 0.00 | 0.03 | 0.20 | 0.07 | 0.16 | 0.49 | 0.50 | 0.27 | 0.12 | 0.16 | 0.80 |
| cIBB3: | 0.20 | 0.07 | 0.09 | 0.03 | 0.05 | 0.47 | 0.12 | 0.47 | 0.01 | 0.20 | 0.04 | 0.17 | 0.16 | 0.23 | 0.67 | 0.33 | 0.05 | 0.31 | 0.17 | 0.02 |
| cIBB4: | 0.17 | 0.00 | 0.00 | 0.21 | 0.02 | 0.38 | 0.14 | 0.35 | 0.00 | 0.10 | 0.00 | 0.09 | 0.08 | 0.00 | 0.66 | 0.41 | 0.10 | 0.35 | 0.26 | 0.03 |
| cIBB2: | 0.25 | 0.18 | 0.16 | 0.11 | 0.29 | 0.51 | 0.94 | 0.25 | 0.13 | 0.18 | 0.00 | 0.26 | 0.25 | 0.79 | 0.57 | 0.65 | 0.71 | 0.12 | 0.27 | 0.89 |
| cIBB7: | 0.25 | 0.30 | 0.07 | 0.28 | 0.01 | 0.08 | 0.65 | 0.15 | 0.04 | 0.22 | 0.02 | 0.18 | 0.26 | 0.59 | 0.86 | 0.86 | 0.88 | 0.96 | 0.39 | 0.71 |
| alpha-helix | 41 | 42 | 43 | 44 | 45 | 46 | 47 | 48 | 49 | 50 | 51 | 52 | 53 | 54 | 55 | 56 | 57 | 58 | 59 | 60 |
| cIBB1: | 0.23 | 0.05 | 0.27 | 0.79 | 0.00 | 0.10 | 0.14 | 0.15 | 0.23 | 0.15 | 0.25 | 0.16 | 0.16 | 0.40 | 0.38 | 0.61 | 0.68 | 0.82 | 0.74 | 0.938 |
| cIBB5: | 0.16 | 0.00 | 0.00 | 0.03 | 0.40 | 0.23 | 0.12 | 0.33 | 0.22 | 0.21 | 0.16 | 0.30 | 0.23 | 0.56 | 0.38 | 0.74 | 0.67 | 0.67 | 0.78 | 0.724 |
| cIBB6: | 0.13 | 0.00 | 0.00 | 0.27 | 0.18 | 0.01 | 0.09 | 0.18 | 0.16 | 0.00 | 0.17 | 0.30 | 0.12 | 0.88 | 0.39 | 0.31 | 0.34 | 0.49 | 0.83 | 0.726 |
| cIBB3: | 0.28 | 0.06 | 0.08 | 0.51 | 0.51 | 0.33 | 0.13 | 0.38 | 0.16 | 0.17 | 0.04 | 0.19 | 0.09 | 0.17 | 0.26 | 0.70 | 0.57 | 0.35 | 0.42 | 0.257 |
| cIBB4: | 0.28 | 0.00 | 0.00 | 0.42 | 0.29 | 0.17 | 0.07 | 0.19 | 0.00 | 0.09 | 0.00 | 0.04 | 0.11 | 0.00 | 0.30 | 0.40 | 0.15 | 0.27 | 0.23 | 0.492 |
| cIBB2: | 0.36 | 0.12 | 0.14 | 0.29 | 0.64 | 0.15 | 0.64 | 0.23 | 0.35 | 0.16 | 0.00 | 0.34 | 0.25 | 0.69 | 0.46 | 0.47 | 1.13 | 0.68 | 0.93 | 1.43 |
| cIBB7: | 0.45 | 0.30 | 0.07 | 0.45 | 0.18 | 0.29 | 0.83 | 0.10 | 0.12 | 0.12 | 0.01 | 0.15 | 0.05 | 0.52 | 0.69 | 0.66 | 0.66 | 1.28 | 0.86 | 1.307 |
| beta-sheet | 41 | 42 | 43 | 44 | 45 | 46 | 47 | 48 | 49 | 50 | 51 | 52 | 53 | 54 | 55 | 56 | 57 | 58 | 59 | 60 |
| cIBB1: | 0.13 | 0.12 | 0.22 | 0.22 | 0.00 | 0.07 | 0.20 | 0.20 | 0.10 | 0.19 | 0.20 | 0.14 | 0.21 | 0.37 | 0.33 | 1.00 | 0.64 | 0.38 | 0.36 | 0.561 |
| cIBB5: | 0.09 | 0.00 | 0.00 | 0.10 | 0.23 | 0.24 | 0.24 | 0.51 | 0.18 | 0.46 | 0.39 | 0.35 | 0.53 | 0.52 | 0.49 | 0.71 | 0.46 | 0.33 | 0.43 | 0.395 |
| cIBB6: | 0.13 | 0.00 | 0.00 | 0.07 | 0.30 | 0.10 | 0.13 | 0.21 | 0.11 | 0.00 | 0.14 | 0.12 | 0.16 | 0.43 | 0.54 | 0.49 | 0.26 | 0.27 | 0.38 | 0.573 |
| cIBB3: | 0.22 | 0.05 | 0.07 | 0.13 | 0.29 | 0.32 | 0.08 | 0.36 | 0.04 | 0.14 | 0.04 | 0.13 | 0.20 | 0.18 | 0.27 | 0.85 | 0.47 | 0.60 | 0.28 | 0.187 |
| cIBB4: | 0.10 | 0.00 | 0.00 | 0.46 | 0.07 | 0.28 | 0.13 | 0.20 | 0.00 | 0.07 | 0.00 | 0.04 | 0.19 | 0.00 | 0.38 | 0.51 | 0.23 | 0.52 | 0.15 | 0.128 |
| cIBB2: | 0.16 | 0.25 | 0.11 | 0.24 | 0.55 | 0.16 | 0.72 | 0.13 | 0.11 | 0.13 | 0.00 | 0.09 | 0.44 | 0.52 | 0.68 | 0.73 | 0.84 | 0.30 | 0.41 | 0.493 |
| cIBB7: | 0.29 | 0.22 | 0.05 | 0.37 | 0.05 | 0.13 | 0.98 | 0.08 | 0.12 | 0.13 | 0.11 | 0.14 | 0.11 | 0.66 | 0.95 | 0.91 | 0.69 | 0.81 | 0.48 | 0.853 |
| beta-turn | 41 | 42 | 43 | 44 | 45 | 46 | 47 | 48 | 49 | 50 | 51 | 52 | 53 | 54 | 55 | 56 | 57 | 58 | 59 | 60 |
| cIBB1: | 0.17 | 0.09 | 0.07 | 0.86 | 0.00 | 0.10 | 0.06 | 0.26 | 0.19 | 0.18 | 0.32 | 0.21 | 0.21 | 0.29 | 0.40 | 0.38 | 0.57 | 0.77 | 0.75 | 0.794 |
| cIBB5: | 0.16 | 0.00 | 0.00 | 0.02 | 0.29 | 0.18 | 0.06 | 0.26 | 0.13 | 0.24 | 0.28 | 0.59 | 0.37 | 0.63 | 0.35 | 0.84 | 0.80 | 0.67 | 0.86 | 0.616 |
| cIBB6: | 0.07 | 0.00 | 0.00 | 0.30 | 0.18 | 0.08 | 0.04 | 0.30 | 0.10 | 0.00 | 0.05 | 0.35 | 0.17 | 0.88 | 0.29 | 0.30 | 0.40 | 0.36 | 0.83 | 0.628 |
| cIBB3: | 0.34 | 0.01 | 0.02 | 0.56 | 0.49 | 0.11 | 0.10 | 0.55 | 0.13 | 0.04 | 0.01 | 0.25 | 0.05 | 0.20 | 0.32 | 0.49 | 0.54 | 0.66 | 0.35 | 0.254 |
| cIBB4: | 0.28 | 0.00 | 0.00 | 0.57 | 0.32 | 0.16 | 0.04 | 0.32 | 0.00 | 0.02 | 0.00 | 0.03 | 0.04 | 0.00 | 0.36 | 0.27 | 0.08 | 0.56 | 0.29 | 0.542 |
| cIBB2: | 0.21 | 0.16 | 0.03 | 0.37 | 0.80 | 0.16 | 0.17 | 0.24 | 0.25 | 0.04 | 0.00 | 0.38 | 0.22 | 0.71 | 0.76 | 0.42 | 1.06 | 0.73 | 0.99 | 1.136 |
| cIBB7: | 0.26 | 0.19 | 0.02 | 0.55 | 0.20 | 0.27 | 0.69 | 0.01 | 0.01 | 0.10 | 0.14 | 0.10 | 0.04 | 0.72 | 1.15 | 0.97 | 0.84 | 1.13 | 0.83 | 1.029 |
| coil | 41 | 42 | 43 | 44 | 45 | 46 | 47 | 48 | 49 | 50 | 51 | 52 | 53 | 54 | 55 | 56 | 57 | 58 | 59 | 60 |
| cIBB1: | 0.16 | 0.07 | 0.01 | 0.94 | 0.00 | 0.07 | 0.06 | 0.19 | 0.18 | 0.16 | 0.24 | 0.25 | 0.15 | 0.24 | 0.33 | 0.37 | 0.63 | 0.88 | 0.82 | 0.703 |
| cIBB5: | 0.12 | 0.00 | 0.00 | 0.03 | 0.27 | 0.16 | 0.05 | 0.23 | 0.12 | 0.22 | 0.10 | 0.27 | 0.40 | 0.49 | 0.28 | 1.23 | 0.81 | 0.49 | 0.92 | 0.652 |
| cIBB6: | 0.09 | 0.00 | 0.00 | 0.33 | 0.15 | 0.05 | 0.05 | 0.22 | 0.15 | 0.00 | 0.06 | 0.22 | 0.09 | 0.97 | 0.34 | 0.30 | 0.37 | 0.49 | 0.93 | 0.657 |
| cIBB3: | 0.23 | 0.00 | 0.03 | 0.61 | 0.53 | 0.25 | 0.15 | 0.38 | 0.12 | 0.01 | 0.02 | 0.16 | 0.10 | 0.28 | 0.28 | 0.49 | 0.61 | 0.38 | 0.26 | 0.285 |
| cIBB4: | 0.21 | 0.00 | 0.00 | 0.56 | 0.35 | 0.16 | 0.05 | 0.22 | 0.00 | 0.00 | 0.00 | 0.04 | 0.05 | 0.00 | 0.26 | 0.31 | 0.05 | 0.31 | 0.23 | 0.597 |
| cIBB2: | 0.26 | 0.04 | 0.00 | 0.38 | 0.82 | 0.18 | 0.19 | 0.14 | 0.28 | 0.00 | 0.00 | 0.22 | 0.21 | 0.53 | 0.42 | 0.42 | 1.61 | 0.79 | 1.07 | 1.67 |
| cIBB7: | 0.32 | 0.15 | 0.00 | 0.56 | 0.22 | 0.22 | 0.63 | 0.01 | 0.13 | 0.04 | 0.15 | 0.05 | 0.16 | 0.51 | 0.81 | 0.94 | 0.74 | 1.59 | 0.92 | 1.602 |

**FIGURE S18 Fluctuation of biochemical propensities of each residue of the IBB domain consensus sequence.** The standard deviation of the score for each biochemical propensity among organisms is shown for each position in the IBB consensus sequence.

### FIGURE S19

#### polarity

| Euclidean distance | |
|---|---|
| cIBB1: cIBB5 | 13.83 |
| cIBB3: cIBB4 | 24.43 |
| cIBB1: cIBB6 | 29.70 |
| cIBB2: cIBB7 | 54.67 |
| cIBB3: cIBB2 | 107.62 |
| cIBB1: cIBB3 | 169.78 |

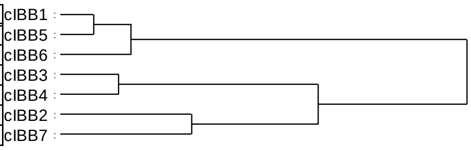


#### acidic amino acid

| Euclidean distance | |
|---|---|
| cIBB3: cIBB4 | 1.00 |
| cIBB1: cIBB6 | 3.00 |
| cIBB1: cIBB5 | 5.00 |
| cIBB2: cIBB7 | 7.00 |
| cIBB3: cIBB2 | 17.00 |
| cIBB1: cIBB3 | 22.43 |

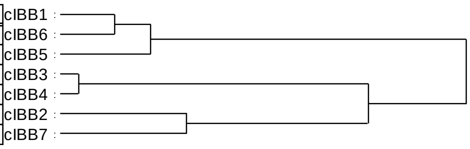


#### hydrophobicity

| Euclidean distance | |
|---|---|
| cIBB5: cIBB6 | 8.27 |
| cIBB3: cIBB4 | 8.77 |
| cIBB1: cIBB5 | 10.49 |
| cIBB2: cIBB7 | 27.06 |
| cIBB3: cIBB2 | 70.48 |
| cIBB1: cIBB3 | 124.12 |

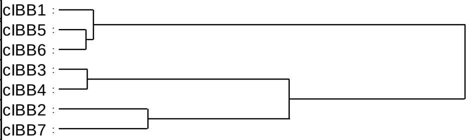


#### basic amino acid

| Euclidean distance | |
|---|---|
| cIBB1: cIBB5 | 1.00 |
| cIBB3: cIBB4 | 1.00 |
| cIBB1: cIBB6 | 1.67 |
| cIBB2: cIBB7 | 4.00 |
| cIBB3: cIBB2 | 4.50 |
| cIBB1: cIBB3 | 11.26 |

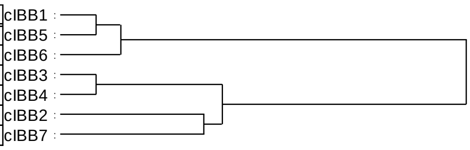


#### bulkiness

| Euclidean distance | |
|---|---|
| cIBB5: cIBB6 | 2.47 |
| cIBB1: cIBB5 | 12.97 |
| cIBB3: cIBB4 | 20.09 |
| cIBB2: cIBB7 | 21.51 |
| cIBB3: cIBB2 | 56.80 |
| cIBB1: cIBB3 | 89.92 |

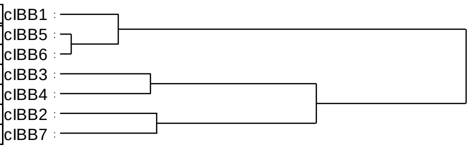


#### helix breaker

| Euclidean distance | |
|---|---|
| cIBB1: cIBB6 | 0.00 |
| cIBB1: cIBB5 | 1.33 |
| cIBB3: cIBB4 | 2.00 |
| cIBB3: cIBB2 | 4.67 |
| cIBB3: cIBB7 | 7.33 |
| cIBB1: cIBB3 | 16.67 |

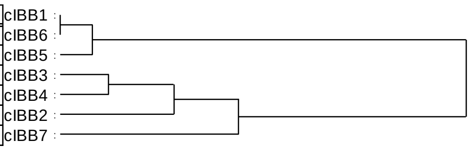


#### average flexibility

| Euclidean distance | |
|---|---|
| cIBB5: cIBB6 | 4.33 |
| cIBB3: cIBB4 | 16.68 |
| cIBB1: cIBB5 | 20.13 |
| cIBB2: cIBB7 | 42.70 |
| cIBB3: cIBB2 | 53.92 |
| cIBB1: cIBB3 | 87.85 |

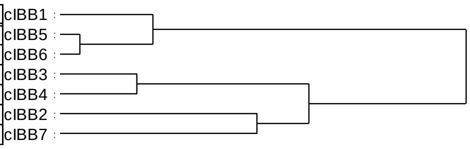


#### alpha-helix

| Euclidean distance | |
|---|---|
| cIBB3: cIBB4 | 4.03 |
| cIBB1: cIBB6 | 4.15 |
| cIBB5: cIBB2 | 4.79 |
| cIBB5: cIBB7 | 9.50 |
| cIBB5: cIBB3 | 15.65 |
| cIBB1: cIBB5 | 21.62 |

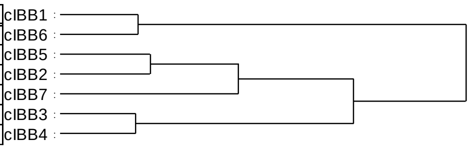


#### beta-sheet

| Euclidean distance | |
|---|---|
| cIBB3: cIBB4 | 1.13 |
| cIBB1: cIBB5 | 1.70 |
| cIBB1: cIBB6 | 2.79 |
| cIBB3: cIBB2 | 3.81 |
| cIBB3: cIBB7 | 9.59 |
| cIBB1: cIBB3 | 32.63 |

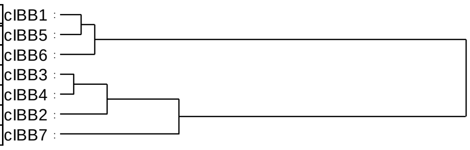


#### beta-turn

| Euclidean distance | |
|---|---|
| cIBB3: cIBB4 | 1.39 |
| cIBB1: cIBB6 | 4.20 |
| cIBB2: cIBB7 | 4.20 |
| cIBB1: cIBB5 | 8.41 |
| cIBB3: cIBB2 | 12.12 |
| cIBB1: cIBB3 | 20.40 |

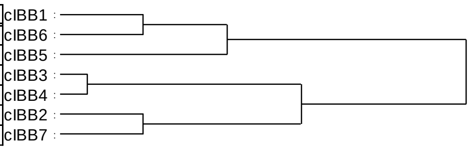


#### coil

| Euclidean distance | |
|---|---|
| cIBB3: cIBB4 | 2.97 |
| cIBB2: cIBB7 | 4.24 |
| cIBB1: cIBB6 | 4.73 |
| cIBB1: cIBB5 | 8.13 |
| cIBB3: cIBB2 | 10.97 |
| cIBB1: cIBB3 | 22.54 |

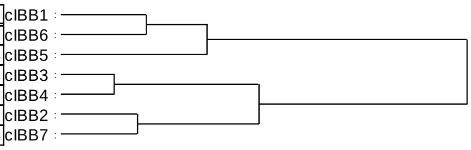


**FIGURE S19 Clustering of the KPNA family proteins based on the score of biochemical/biophysical propensities of the IBB domain.** Euclidean distance among the IBB consensus sequences obtained from the contact area at each position in each family. A dendrogram was drawn based on this Euclidean distance.

### FIGURE S20

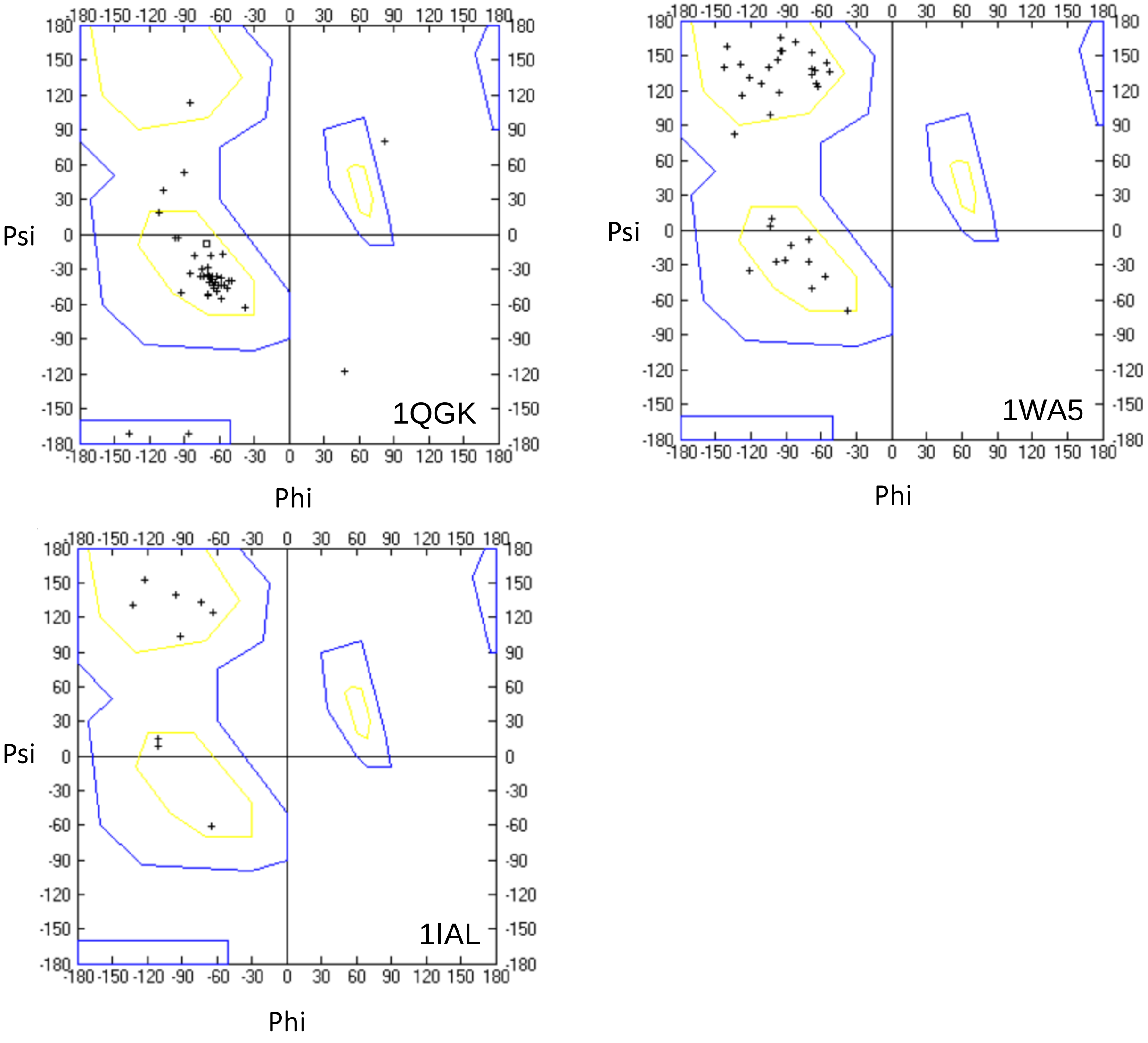


**FIGURE S20 Ramachandran plot of IBB domain residues.** Ramachandran plot of IBB domain residues in the crystal structures in 1QGK (importin β binding form), 1WA5 (nuclear export complex), 1IAL (autoinhibition form). The residues corresponding to position 13–54 for 1QGK, 7–16 and 26–53 for1WA5, 44–54 for 1IAL in Fig S3 were plotted.

### FIGURE S21

| | | | | Residues in close proximity in a binding partner in the modeled structure | | | | | | | Conservation of amino acid in tested organisms | | | | | | |
|---|---|---|---|---|---|---|---|---|---|---|---|---|---|---|---|---|---|
| Criteria | Position | Residues in close proximity in a binding partner previously reported | MAX/MIN | cIBB1 | cIBB5 | cIBB6 | cIBB3 | cIBB4 | cIBB2 | cIBB7 | cIBB1 | cIBB5 | cIBB6 | cIBB3 | cIBB4 | cIBB2 | cIBB7 |
| ● | 13 | E281 D288 W342 V350 | 1.17 | E281 S284 N285 D288 W342 V350 L354 | E281 S284 N285 D288 W342 V350 L354 | E281 S284 N285 D288 W342 V350 L354 | E281 S284 N285 D288 W342 V350 L354 | E281 S284 N285 D288 W342 V350 L354 | E281 S284 N285 D288 W342 V350 L354 | E281 S284 N285 D288 W342 V350 L354 | 98.1 | 100 | 98.8 | 88.7 | 81.1 | 97.9 | 93.8 |
| | 18 | D426 T427 M388 W430 | 1.04 | M388 D426 T427 W430 | M388 D426 T427 W430 | M388 D426 T427 W430 | M388 D426 T427 W430 | M388 D426 T427 W430 | M388 D426 T427 W430 | M388 D426 T427 W430 | 99.4 | 100 | 100 | 94.4 | 85.8 | 97.9 | 97.9 |
| | 28 | G623 D627 | 1.11 | S582 G623 E626 D627 Q665 | S582 G623 E626 D627 Q665 | S582 G623 E626 D627 Q665 | S582 G623 E626 D627 Q665 | S582 G623 E626 D627 Q665 | S582 G623 E626 D627 Q665 | S582 G623 E626 D627 Q665 | 100 | 100 | 100 | 96.8 | 100 | 95.7 | 99.0 |
| | 31 | E530 | 1.13 | E530 A586 Q589 D627 | E530 A586 Q589 D627 | E530 A586 Q589 D627 | E530 A586 Q589 D627 | E530 A586 Q589 D627 | E530 A586 Q589 D627 | E530 A586 Q589 D627 | 100 | 98.7 | 100 | 97.6 | 99.3 | 99.5 | 96.9 |
| | 39 | G672 D676 | 1.13 | S633 G672 D676 R679 | S633 G672 D676 | S633 G672 D676 | S633 G672 D676 R679 | G672 D676 R679 | S633 G672 D676 | S633 G672 D676 | 100 | 100 | 100 | 96.8 | 99.3 | 100 | 100 |
| | 40 | E763 | 1.10 | E763 | E763 | E763 | E763 | E763 | E763 | E763 | 100 | 100 | 100 | 96.8 | 99.3 | 93.6 | 100 |
| | 51 | D824 W864 | 1.13 | G820 D824 L861 W864 | T770 G820 D824 L861 W864 | T770 G820 D824 L861 W864 | T770 G820 D824 L861 W864 | T770 G820 D824 L861 W864 | T770 G820 D824 L861 W864 | T770 G820 D824 L861 W864 | 95.0 | 93.3 | 98.4 | 99.2 | 100 | 100 | 99.0 |
| | 14 | W342 | 1.26 | D339 D340 D341 W342 | D340 W342 | D339 D340 W342 | D340 W342 | D339 D340 D341 W342 | D339 D340 D341 W342 | D339 D340 D341 W342 | 98.1 | 94.7 | 95.6 | 77.4 | 81.8 | 91.4 | 51.0 |
| | 17 | K346 | 1.26 | K346 V350 M388 | K346 V350 M388 | K346 V350 M388 | K346 | K346 V350 | K346 V350 | K346 V350 | 98.1 | 100 | 98.8 | 96.8 | 87.2 | 97.9 | 94.8 |
| | 43 | D719 E767 | 1.47 | D719 L722 E767 G771 | D719 L722 E767 G771 | D719 L722 E767 G771 | D719 L722 E767 G771 | D719 L722 E767 G771 | R679 S715 D719 E767 | R679 S715 D719 E767 | 75.8 | 100 | 100 | 97.6 | 100 | 95.2 | 99.0 |
| | 50 | L722 | 1.33 | L722 Q774 | L722 Q774 | L722 Q774 | L722 Q774 | L722 Q774 | L722 Q774 | L722 Q774 | 95.0 | 92.0 | 100 | 91.9 | 98.0 | 93.6 | 96.9 |
| | 53 | W864 | 1.12 | T860 W864 | T860 W864 | T860 W864 | T860 W864 | T860 W864 | T860 W864 | T860 L861 W864 | 93.2 | 90.7 | 98.0 | 93.5 | 98.6 | 90.4 | 90.6 |
| ○ | 16 | V350 | 2.29 | V350 M353 | V350 M353 | V350 M353 | V350 M353 | V350 M353 | V350 M353 L354 | D288 V350 M353 | 96.3 | 92.7 | 69.1 | 87.9 | 81.8 | 49.2 | 80.2 |
| | 22 | W472 | 2.23 | W472 | W472 | W472 | E437 W472 | E437 W472 | W472 | W472 | 96.9 | 97.3 | 98.4 | 88.7 | 80.4 | 94.7 | 94.8 |
| | 24 | - | 2.28 | S526 | S526 | S526 | S526 | D579 | - | - | 71.4 | 90.0 | 88.8 | 78.2 | 89.2 | 42.2 | 60.4 |
| | 34 | - | 2.57 | K537 | K537 | K537 | R593 | R593 | R593 | R593 | 96.9 | 96.7 | 99.2 | 89.5 | 96.6 | 89.8 | 86.5 |
| | 35 | R593 | 5.88 | - | - | - | R593 | R593 | R593 M630 | R593 | 96.9 | 98.0 | 99.6 | 85.5 | 90.5 | 84.0 | 68.8 |
| | 48 | - | 2.71 | K857 L861 | K857 L861 | K857 L861 | - | - | - | - | 92.5 | 92.7 | 96.0 | 89.5 | 93.2 | 88.8 | 95.8 |
| | 54 | - | 2.32 | T860 | T860 | T860 | T860 | T860 | T860 | T860 | 91.9 | 78.0 | 72.3 | 91.1 | 99.3 | 52.9 | 55.2 |

**FIGURE S21 Screening and classification of residues of the IBB present in the interface for importin β binding.** The position of the residue which is important for the interaction between the IBB and importin β in the model structures of the importin β binding form were screened. ●: The positions where all family had commonly large contact area. ○: Positions where at least one family had large contact area and difference among family was large. The ratio of maximum value to minimum value among the family is also shown to distinguish commonly important residues among the family and specifically important residues for only some family members. The average value of the contact area per residue were 51.2 angstrom squared, and median of the ratio of maximum and minimum values was 1.49. The residue–residue interactions between IBB domain and importin β at the corresponding positions in the template structure are shown with polar interaction (red) and hydrophobic interaction (purple) indicated. The proximal residues of the importin β present within 4 Å from the residues of IBB domain at each position in modeled structure were detected with PyMol.

### FIGURE S22

| | | | | | Residues in close proximity in a binding partner in the modeled structure | | | | | | | Conservation of amino acid in tested organisms | | | | | | |
|---|---|---|---|---|---|---|---|---|---|---|---|---|---|---|---|---|---|---|
| Criteria | Position | binding molecule | Residues in close proximity in a binding partner previously reported | MAX/MIN | cIBB1 | cIBB5 | cIBB6 | cIBB3 | cIBB4 | cIBB2 | cIBB7 | cIBB1 | cIBB5 | cIBB6 | cIBB3 | cIBB4 | cIBB2 | cIBB7 |
| | 13 | CAS | F117 L157 E166 | 1.411 | R120 F123 P124 E163 E172 | F123 P124 E163 E172 | R120 F123 P124 E163 E172 | R120 F123 P124 E163 E172 | R120 F123 P124 E163 E172 | R120 F123 P124 E163 E172 | R120 F123 P124 E163 E172 | 98.1 | 100 | 98.8 | 88.7 | 81.1 | 97.9 | 93.8 |
| | 28 | ARM repeat | S412 A370 | 1.055 | V324 T325 G326 D328 T331 N364 A367 G368 Q372 | V325 T326 G327 D329 T332 N365 A368 G369 Q373 | V322 T323 G324 D326 T329 N362 A365 G366 Q370 | V312 T313 G314 D316 T319 N352 A355 G356 Q360 | V312 T313 G314 D316 T319 N352 A355 G356 Q360 | V319 T320 G321 D323 T326 N359 A362 G363 Q367 | V310 T311 G312 D314 T317 N350 A353 G354 | 100 | 100 | 100 | 96.8 | 100 | 95.7 | 99.0 |
| | 31 | ARM repeat | G287 | 1.074 | D283 R318 N322 E357 W360 | D284 R319 N323 E358 W361 | D281 R316 N320 E355 W358 | D271 R306 N310 E345 W348 | D271 R306 N310 E345 W348 | D278 R313 N317 E352 W355 | D269 R304 N308 E343 W346 | 100 | 98.7 | 100 | 97.6 | 99.3 | 99.5 | 96.9 |
| | 38 | CAS | - | 1.092 | W170 I173 K174 L177 P228 E229 | W170 I173 K174 L177 P228 E229 | W170 I173 K174 L177 P228 E229 | W170 I173 K174 L177 P228 E229 | W170 I173 K174 L177 P228 E229 | W170 I173 K174 P228 E229 | W170 I173 K174 L177 P228 E229 | 100 | 98.7 | 99.2 | 97.6 | 100 | 98.9 | 99.0 |
| | 39 | CAS | F164 I167 E223 | 1.251 | D226 L227 E232 Y282 E285 | D226 L227 Y282 E284 E285 | D226 L227 Y282 E284 E285 | D226 L227 Y282 E285 | D226 L227 Y282 E285 | D226 L227 E232 Y282 E285 | D226 L227 Y282 E284 E285 | 100 | 100 | 100 | 96.8 | 99.3 | 100 | 100 |
| | | ARM repeat | - | | - | - | - | - | - | K390 | K381 | | | | | | | |
| | 40 | CAS | I167 D220 D279 | 1.033 | E229 E232 D233 | E229 E232 D233 | E229 E232 D233 | E229 E232 D233 | E229 E232 D233 | E229 E232 D233 | E229 E232 D233 | 100 | 100 | 100 | 96.8 | 99.3 | 93.6 | 100 |
| | 49 | ARM repeat | N241 W237 | 1.020 | A155 S156 G157 N158 T162 N195 D199 W234 | A156 G158 T159 T163 N196 D200 W235 | A156 G157 T158 T163 N196 D200 W235 | A143 S144 G145 T146 T150N183 D187 | A143 S144 G145 T146 T150 N183 D187 | A146 S147 G148 T149 T153 N186 D190 W229 | A141 S142 G143 T144 T148 N181 D185 W220 | 96.3 | 96.7 | 98.4 | 99.2 | 100 | 96.3 | 99.0 |
| ● | 51 | ARM repeat | D276 W237 N199 W195 L177 | 1.201 | L110 P114 P116 N153 S156 G157 W191 | L111 P115 P117 N154 S157 W192 | L111 P115 P117 N154 S157 W192 | L99 P105 N141 S144 G145 W179 | L99 P105 N141 S143 W179 | L102 P108 N144 S147 G148 W182 | L97 Q99 P103 N139 S142 W177 | 95.0 | 93.3 | 98.4 | 99.2 | 100 | 100 | 99.0 |
| | 52 | ARM repeat | S160 N157 I123 P121 R117 L115 | 1.148 | S111 W149 N153 Q188 W191 | S112 W150 N154 Q189 W192 | A153 S154 G155 T156 T160 N193 D197 W232 | S100 W137 N141 Q176 W179 | S100 W137 N141 Q176 W179 | S103 W140 N144 Q179 W182 | S98 W135 N139 Q174 W177 | 96.3 | 92.0 | 98.0 | 93.5 | 98.6 | 96.3 | 90.6 |
| | 15 | CAS | F158 | 1.412 | H162 E163 | H162 E163 | H162 E163 | H162 E163 | H162 E163 | H162 F164 | H162 E163 F164 | 93.2 | 92.0 | 71.1 | 84.7 | 81.8 | 58.8 | 68.8 |
| | | ARM repeat | K496 D495 N494 | | R10 Y14 K15 Q490 | R13 Y17 K18 N19 N490 Q491 | R13 Y17 K18 N19 N487 Q488 | R11 F15 K16 N17 N470 E471 D472 | R11 F15 K16 N17 N470 E471 D472 | R11 F15 K16 N17 N479 S481 | E9 R10 F14 K15 Y16 N470 | | | | | | | |
| | 27 | ARM repeat | - | 1.398 | P21 D22 R26 A367 N406 S409 | Q25 R29 A368 N407 S410 | E25 R29 A365 N404 S407 | V22 E23 R27 A355 W390 N394 I397 | L22 E23 R27 A355 W390 N394 I397 | T23 R27 A362 N401 S404 | A21 S22 Q26 A353 N392 T395 D431 | 96.9 | 97.3 | 96.4 | 87.9 | 97.3 | 53.5 | 59.4 |
| | 36 | CAS | - | 1.247 | N167 W170 T171 | N167 W170 T171 | N167 W170 T171 | N167 W170 T171 | N167 W170 T171 K174 | N167 W170 K174 | N167 W170 T171 | 96.9 | 96.7 | 96.4 | 75.0 | 96.6 | 97.3 | 60.4 |
| | 44 | ARM repeat | - | 1.204 | K39 W276 S353 | K42 Q46 W277 S354 | K42 W274 S351 | K40 H44 W264 R306 K341 | K40 W264 R306 K341 | K41 Q44 W271 T308 R313 N348 | K39 Q43 W262 T300 R304 | 74.5 | 99.3 | 97.6 | 91.1 | 91.9 | 96.8 | 90.6 |
| | 45 | ARM repeat | - | 1.315 | R40 L44 R241 W276 Y280 | R43 L47 R242 W277 Y281 | R43 L47 R239 W274 Y278 | R41 L45 R229 W264 Y268 | R41 L45 R229 W264 Y268 | K41 M45 R236 W271 | K40 A44 R227 W262 Y266 | 100 | 97.3 | 96.4 | 91.1 | 97.3 | 81.3 | 99.0 |
| | 50 | ARM repeat | D203 T166 | 1.419 | Q43 S156 W191 N195 W234 D273 | Q46 N52 S157 W192 N196 W235 D274 | Q46 S154 W189 N193 W232 D271 | N50 S144 W179 N183 N219 W222 | H44 N50 S144 W179 N183 W222 D261 | Q44 S147 W182 N186 W229 D268 | N49 S142 W177 N181 W220 D259 | 95.0 | 92.0 | 100 | 91.9 | 98.0 | 93.6 | 96.9 |
| | 8 | CAS | E72 N73 G74 | 4.470 | R75 R120 E121 | R75 R120 E121 | R75 E121 | R75 E121 | R75 I76 V77 E78 D79 E80 R120 E121 | R75 I76 V77 E78 D79 R120 E121 | R75 I76 V77 E78 D79 E80 E121 | 91.9 | 84.0 | 57.8 | 66.9 | 85.1 | 70.6 | 41.7 |
| | 12 | CAS | E162 L163 E166 | 2.218 | S166 E168 | P124 E163 S166 E168 L169 E172 | P124 E163 S166 E168 L169 E172 | E163 S166 E168 L169 | E163 S166 E168 | E163 S166 E168 | S166 E168 L169 | 70.2 | 84.0 | 92.8 | 83.9 | 80.4 | 46.0 | 74.0 |
| ○ | 29 | ARM repeat | T334 D331 W327 | 2.389 | M24 E30 T325 W360 S363 N364 E399 W402 | M27 E33 T326 W361 S364 N365 E400 W403 | M27 E33 T323 W358 S361 N362 E397 W400 | M25 E31 T313 W348 S351 N352 E387 W390 | M25 E31 T313 W348 S351 N352 E387 W390 | M25 E31 T320 W355 S358 N359 E394 W397 | R24 R28 A30 T311 W346 N350 | 98.8 | 99.3 | 100 | 99.2 | 98.6 | 98.9 | 64.6 |
| | 34 | CAS | - | 2.032 | K165 S166 N167 | K165 S166 N167 | K165 S166 N167 | N167 | N167 | N167 | N167 | 96.9 | 96.7 | 99.2 | 89.5 | 96.6 | 89.8 | 86.5 |
| | | ARM repeat | - | | E29 L33 Q34 K438 | E32 I36 Q37 K439 | E32 I36 Q37 K436 | N30 V34 | N30 V34 | I30 V34 | I29 L33 | | | | | | | |

**FIGURE S22 Screening and classification of residues of the IBB present in the interface for nuclear export complex.** The position of the residue which is important for the interaction between the IBB and CAS and/or ARM repeat in the model structures of the nuclear export complex were screened. ●: The positions where all family had commonly large contact area. ○: Positions where at least one family had large contact area and difference among family was large. The ratio of maximum value to minimum value among the family is also shown to distinguish commonly important residues among the family and specifically important residues for only some family members. The average value of the contact area per residue were 86.1 angstrom squared, and median of the ratio of maximum and minimum values was 1.42. The residue–residue interactions between IBB domain and CAS or ARM at the corresponding positions in the template structure are shown with polar interaction (red), hydrophobic interaction (purple) and both (blue) indicated. The proximal residues of the CAS and ARM present within 4 Å from the residues of IBB domain at each position in modeled structure were detected with PyMol.

### FIGURE S23

| | | | | Residues in close proximity in a binding partner in the modeled structure | | | | | | | Conservation of amino acid in tested organisms | | | | | | |
|---|---|---|---|---|---|---|---|---|---|---|---|---|---|---|---|---|---|
| Criteria | Position | Residues in close proximity in a binding partner previously reported | MAX/MIN | cIBB1 | cIBB5 | cIBB6 | cIBB3 | cIBB4 | cIBB2 | cIBB7 | cIBB1 | cIBB5 | cIBB6 | cIBB3 | cIBB4 | cIBB2 | cIBB7 |
| ● | 49 | G150 T151 D192 | 1.0597898 | A155 S156 G157 S158 T162 N195 D199 W234 | A156 S157 G158 T159 F160 T163 | A153 S154 G155 T156 S157 T160 N193 D197 W232 | A143 S144 G145 T146 S147 T150 N183 D187 W222 | A143 S144 G145 T146 S147 T150 N183 D187 W222 | A146 S147 G148 T149 S150 T153 N186 D190 | A153 S154 G155 T156 S157 T160 N193 D197 W232 | 96.3 | 96.7 | 98.4 | 99.2 | 100 | 96.3 | 99.0 |
| | 51 | L104 R106 E107 | 1.1462509 | L110 S111 K112 E113 P116 N153 S156 W191 | L111 K113 E114 P117 N154 S157 W192 | L108 S109 K110 E111 P114 N151 S154 W189 | L99 S100 S101 D102 P105 N141 S144W179 | L99 S100 S101 D102 P105 N141 S144 W179 | L102 S103 R104 E105 P108 N144 S147 W182 | L108 S109 K110 E111 P114 N151 S154 | 95.0 | 93.3 | 98.4 | 99.2 | 100 | 100 | 99.0 |
| | 52 | S105 N146 | 1.2119613 | S111 W149 N153 Q188 W191 | S112 W150 N154 Q189 W192 | S99 W147 N151 Q186 W189 | S60 S100 W137 N141 Q176 W179 | S100 W137 N141 Q176 W179 | S103 W140 N144 Q179 W182 | S109 W147 N151 Q186 W189 | 96.3 | 92.0 | 98.0 | 93.5 | 98.6 | 96.3 | 90.6 |
| | 44 | - | 1.382716 | Q43 W276 S314 R318 S353 | Q46 W277 S315 R319 S354 | Q46 W274 R316 | H44 W264 R306 K341 | H44 R306 K341 | Q44 W271 R313 | Q43 W262 R304 | 74.5 | 99.3 | 97.6 | 91.1 | 91.9 | 96.8 | 90.6 |
| | 46 | R238 S234 D270 Y277 | 1.1538974 | E41 F45 S237 R241 D273 W276 Y280 | E44 F48 S238 R242 D274 W277 Y281 | E44 F48 S235 R239 D271 W274 Y278 | D42 L46 W222 V225 R229 D261 W264 Y268 | D42 L46 W222 V225 R229 D261 W264 Y268 | D42 L46 W229 S232 R236 D268 W271 Y275 | D41 L45 S223 R227 D259 W262 Y266 | 99.4 | 97.3 | 99.6 | 89.5 | 92.6 | 92.5 | 96.9 |
| ○ | 45 | - | 9.1517857 | L44 W276 S314 | L47 W277 S315 | L47 | L45 W264 T302 K341 | L45 W264 T302 K341 | M45 | A44 W262 N297 T300 | 100 | 97.3 | 96.4 | 91.1 | 97.3 | 81.3 | 99.0 |

**FIGURE S23 Screening and classification of residues of the IBB present in the interface for autoinhibition form.** The position of the residue which is important for the interaction between the IBB and ARM repeat in the model structures of the autoinhibition form were screened. ●: The positions where all family had commonly large contact area. ○: Positions where at least one family had large contact area and difference among family was large. The ratio of maximum value to minimum value among the family is also shown to distinguish commonly important residues among the family and specifically important residues for only some family members. The average value of the contact area per residue were 98.8 angstrom squared, and median of the ratio of maximum and minimum values was 1.38. The residue–residue interactions between IBB domain and ARM at the corresponding positions in the template structure are shown with polar interaction (red) indicated. The proximal residues of the ARM present within 4 Å from the residues of IBB domain at each position in modeled structure were detected with PyMol.
