## Supporting file legends for "Biochemical propensity mapping for structural and functional anatomy of importin α IBB domain"

**Supporting file 1**

The accession number (AC), gene name (GN), class, organism of the entries included in sequence sets of KPNA1-7 are listed.

**Supporting file 2**

Multiple sequence alignment of IBB domain sequences of each importin α family member (listed in Supporitng file 1). Arbitrary numbers are assigned on each multiple alignment result. The most conserved amino acids or gaps at each position are shown at the bottom.

**Supporting file 3**

The multiple alignment of the full-length consensus sequences of the importin α family.

**Supporting file 4**

The number of each amino acid organisms or gap that appears at a position in the aligned IBB domain sequence included in the sequence set for each 'KPNA'. It corresponds to the number of IBB domain sequences of organisms that have the amino acid or gap in that position when aligned. The residue position numbers corresponding to Supporting file 2 are shown at the top. The number of amino acid species that appear at that position is shown as an amino acid variation. The most emerging amino acid species and their preservation at that location are also shown.

**Supporting file 5**

The alignment of the target sequence and template sequence in homology modeling. Arbitrary numbers are assigned on each multiple alignment result.
