## Supporting file 1 for "Biochemical propensity mapping for structural and functional anatomy of importin α IBB domain"

| AC | GN | Class | Organism |
|---|---|---|---|
| A0A2S2R7W1 | KPNA1 | Insecta | Sipha flava (yellow sugarcane aphid) |
| A0A6F9DGR6 | KPNA1 | Ascidiacea | Phallusia mammillata |
| A0A1A8RH86 | KPNA1 | Actinopterygii | Nothobranchius rachovii (bluefin notho) |
| A0A6P3VH89 | KPNA1 | Actinopterygii | Clupea harengus (Atlantic herring) |
| A0A4W4H6M5 | KPNA1 | Actinopterygii | Electrophorus electricus (Electric eel) (Gymnotus electricus) |
| A0A4W4GZA4 | KPNA1 | Actinopterygii | Electrophorus electricus (Electric eel) (Gymnotus electricus) |
| A0A4W4H6H5 | KPNA1 | Actinopterygii | Electrophorus electricus (Electric eel) (Gymnotus electricus) |
| A0A1A8LEE8 | KPNA1 | Actinopterygii | Nothobranchius pienaari |
| A0A671WDU1 | KPNA1 | Actinopterygii | Sparus aurata (Gilthead sea bream) |
| A0A671WDK5 | KPNA1 | Actinopterygii | Sparus aurata (Gilthead sea bream) |
| A0A671WDF6 | KPNA1 | Actinopterygii | Sparus aurata (Gilthead sea bream) |
| A0A671W6F8 | KPNA1 | Actinopterygii | Sparus aurata (Gilthead sea bream) |
| A0A4W6DL04 | KPNA1 | Actinopterygii | Lates calcarifer (Barramundi) (Holocentrus calcarifer) |
| W5U9X8 | KPNA1 | Actinopterygii | Ictalurus punctatus (Channel catfish) (Silurus punctatus) |
| A0A1A8JDE8 | KPNA1 | Actinopterygii | Nothobranchius kuhntae (Beira killifish) |
| A0A1A7Y2I3 | KPNA1 | Actinopterygii | Iconisemion striatum |
| A0A1A8A5F9 | KPNA1 | Actinopterygii | Nothobranchius furzeri (Turquoise killifish) |
| A0A671W6K4 | KPNA1 | Actinopterygii | Sparus aurata (Gilthead sea bream) |
| E6ZIU3 | KPNA1 | Actinopterygii | Dicentrarchus labrax (European seabass) (Morone labrax) |
| A0A4W6DJI5 | KPNA1 | Actinopterygii | Lates calcarifer (Barramundi) (Holocentrus calcarifer) |
| A0A4W6DL44 | KPNA1 | Actinopterygii | Lates calcarifer (Barramundi) (Holocentrus calcarifer) |
| A0A4W6DKG1 | KPNA1 | Actinopterygii | Lates calcarifer (Barramundi) (Holocentrus calcarifer) |
| A0A4W6DJC0 | KPNA1 | Actinopterygii | Lates calcarifer (Barramundi) (Holocentrus calcarifer) |
| A0A4W6DJY6 | KPNA1 | Actinopterygii | Lates calcarifer (Barramundi) (Holocentrus calcarifer) |
| A0A4W6DLH5 | KPNA1 | Actinopterygii | Lates calcarifer (Barramundi) (Holocentrus calcarifer) |
| A0A4W6DL87 | KPNA1 | Actinopterygii | Lates calcarifer (Barramundi) (Holocentrus calcarifer) |
| F1QB53 | KPNA1 | Actinopterygii | Danio rerio (Zebrafish) (Brachydanio rerio) |
| A0A6J2UXX7 | KPNA1 | Actinopterygii | Chanos chanos (Milkfish) (Mugil chanos) |
| A0A4W4H6S8 | KPNA1 | Actinopterygii | Electrophorus electricus (Electric eel) (Gymnotus electricus) |
| A0A4W4H6S2 | KPNA1 | Actinopterygii | Electrophorus electricus (Electric eel) (Gymnotus electricus) |
| A0A6P8SZR2 | KPNA1 | Actinopterygii | Gymnodraco acuticeps (Antarctic dragonfish) |
| A0A4Z2JG54 | KPNA1 | Actinopterygii | Liparis tanakae (Tanaka's snailfish) |
| A0A2I4AQA5 | KPNA1 | Actinopterygii | Austrofundulus limnaeus |
| A0A4Z2HHP2 | KPNA1 | Actinopterygii | Liparis tanakae (Tanaka's snailfish) |
| A0A6P7KJE0 | KPNA1 | Actinopterygii | Parambassis ranga (Indian glassy fish) |
| A0A6J2RVP8 | KPNA1 | Actinopterygii | Cottoperca gobio (Frogmouth) (Aphritis gobio) |
| A0A1A8EKN4 | KPNA1 | Actinopterygii | Nothobranchius korthausae |
| A0A1A8C032 | KPNA1 | Actinopterygii | Nothobranchius kadleci |
| H2LZX3 | KPNA1 | Actinopterygii | Oryzias latipes (Japanese rice fish) (Japanese killifish) |
| H3A826 | KPNA1 | Sarcopterygii | Latimeria chalumnae (Coelacanth) |
| A0A6I8PYZ6 | KPNA1 | Amphibia | Xenopus tropicalis (Western clawed frog) (Silurana tropicalis) |
| A0A1L8H5K7 | KPNA1 | Amphibia | Xenopus laevis (African clawed frog) |
| A0A6P7XWN2 | KPNA1 | Amphibia | Microcaecilia unicolor |
| A0A6I8RJ96 | KPNA1 | Amphibia | Xenopus tropicalis (Western clawed frog) (Silurana tropicalis) |
| F6TUG5 | KPNA1 | Amphibia | Xenopus tropicalis (Western clawed frog) (Silurana tropicalis) |
| A0A6I8RA27 | KPNA1 | Amphibia | Xenopus tropicalis (Western clawed frog) (Silurana tropicalis) |
| A0A6P8RC89 | KPNA1 | Amphibia | Geotrypetes seraphini (Gaboon caecilian) (Caecilia seraphini) |
| P52170 | KPNA1 | Amphibia | Xenopus laevis (African clawed frog) |
| A0A669QB17 | KPNA1 | Reptilia | Phasianus colchicus (Common pheasant) |
| A0A674I969 | KPNA1 | Reptilia | Terrapene carolina triunguis (Three-toed box turtle) |
| A0A3Q0FNG3 | KPNA1 | Reptilia | Alligator sinensis (Chinese alligator) |
| A0A663LTN7 | KPNA1 | Reptilia | Athene cunicularia (Burrowing owl) (Speotyto cunicularia) |
| A0A6J1URU5 | KPNA1 | Reptilia | Notechis scutatus (mainland tiger snake) |
| A0A674GRG3 | KPNA1 | Reptilia | Taeniopygia guttata (Zebra finch) (Poephila guttata) |
| A0A670JR46 | KPNA1 | Reptilia | Podarcis muralis (Wall lizard) (Lacerta muralis) |
| A0A452I0C3 | KPNA1 | Reptilia | Gopherus agassizii (Agassiz's desert tortoise) |
| A0A452J830 | KPNA1 | Reptilia | Gallus gallus (Chicken) |
| A0A2I0ML81 | KPNA1 | Reptilia | Columba livia (Rock dove) |
| A0A6J0H909 | KPNA1 | Reptilia | Lepidothrix coronata (blue-crowned manakin) |
| A0A6J2HDZ7 | KPNA1 | Reptilia | Pipra filicauda (Wire-tailed manakin) |
| H9GBL6 | KPNA1 | Reptilia | Anolis carolinensis (Green anole) (American chameleon) |
| A0A6I9XTZ4 | KPNA1 | Reptilia | Thamnophis sirtalis |
| A0A6I9HG44 | KPNA1 | Reptilia | Geospiza fortis (Medium ground-finch) |
| U3JQT0 | KPNA1 | Reptilia | Ficedula albicollis (Collared flycatcher) (Muscicapa albicollis) |
| U3IB40 | KPNA1 | Reptilia | Anas platyrhynchos platyrhynchos (Northern mallard) |
| K7GFN3 | KPNA1 | Reptilia | Pelodiscus sinensis (Chinese softshell turtle) (Trionyx sinensis) |
| A0A670ZFW8 | KPNA1 | Reptilia | Pseudonaja textilis (Eastern brown snake) |
| A0A218UWZ4 | KPNA1 | Reptilia | Lonchura striata domestica (Bengalese finch) |
| A0A218VEF9 | KPNA1 | Reptilia | Lonchura striata domestica (Bengalese finch) |
| A0A452I0D9 | KPNA1 | Reptilia | Gopherus agassizii (Agassiz's desert tortoise) |
| V8P5N0 | KPNA1 | Reptilia | Ophiophagus hannah (King cobra) (Naja hannah) |
| A0A091PWE7 | KPNA1 | Reptilia | Haliaeetus albicilla (White-tailed sea-eagle) (Falco albicilla) |
| A0A672U9J9 | KPNA1 | Reptilia | Strigops habroptila (Kakapo) |
| A0A6P9BB21 | KPNA1 | Reptilia | Pantherophis guttatus (Corn snake) (Elaphe guttata) |
| A0A6J3DK49 | KPNA1 | Reptilia | Aythya fuligula (Tufted duck) (Anas fuligula) |
| G1NMJ6 | KPNA1 | Reptilia | Meleagris gallopavo (Wild turkey) |
| A0A1V4JB82 | KPNA1 | Reptilia | Patagioenas fasciata monilis |
| A0A663FFE2 | KPNA1 | Reptilia | Aquila chrysaetos chrysaetos |
| A0A6J0TGA2 | KPNA1 | Reptilia | Pogona vitticeps (central bearded dragon) |
| P83953 | KPNA1 | Mammalia | Rattus norvegicus (Rat) |
| P52294 | KPNA1 | Mammalia | Homo sapiens (Human) |
| Q60960 | KPNA1 | Mammalia | Mus musculus (Mouse) |
| A2VE08 | KPNA1 | Mammalia | Bos taurus (Bovine) |

| AC | GN | Class | Organism |
|---|---|---|---|
| Q5R909 | KPNA1 | Mammalia | Pongo abelii (Sumatran orangutan) (Pongo pygmaeus abelii) |
| A0A6P6DCY3 | KPNA1 | Mammalia | Octodon degus (Degu) (Sciurus degus) |
| I3N7W7 | KPNA1 | Mammalia | Ictidomys tridecemlineatus (Thirteen-lined ground squirrel) (Spermophilus tridecemlineatus) |
| A0A6P5RCS7 | KPNA1 | Mammalia | Mus caroli (Ryukyu mouse) (Ricefield mouse) |
| H0VB68 | KPNA1 | Mammalia | Cavia porcellus (Guinea pig) |
| A0A3Q0CSU4 | KPNA1 | Mammalia | Mesocricetus auratus (Golden hamster) |
| A0A1S3FE51 | KPNA1 | Mammalia | Dipodomys ordii (Ord's kangaroo rat) |
| A0A1S3AD74 | KPNA1 | Mammalia | Erinaceus europaeus (Western European hedgehog) |
| A0A6I9K6T6 | KPNA1 | Mammalia | Chrysochloris asiatica (Cape golden mole) |
| I3L9F7 | KPNA1 | Mammalia | Sus scrofa (Pig) |
| A0A452G7U7 | KPNA1 | Mammalia | Capra hircus (Goat) |
| A0A2I3HG58 | KPNA1 | Mammalia | Nomascus leucogenys (Northern white-cheeked gibbon) (Hylobates leucogenys) |
| A0A5G2QTJ8 | KPNA1 | Mammalia | Sus scrofa (Pig) |
| A0A0B8RTI9 | KPNA1 | Mammalia | Sus scrofa domesticus (domestic pig) |
| A0A341DEI5 | KPNA1 | Mammalia | Neophocaena asiaeorientalis asiaeorientalis (Yangtze finless porpoise) (Neophocaena phocaenoides subsp. asiaeorientalis) |
| A0A667HMJ9 | KPNA1 | Mammalia | Lynx canadensis (Canada lynx) |
| A0A2I2YX55 | KPNA1 | Mammalia | Gorilla gorilla gorilla (Western lowland gorilla) |
| G1M736 | KPNA1 | Mammalia | Ailuropoda melanoleuca (Giant panda) |
| A0A6G1ATK9 | KPNA1 | Mammalia | Crocuta crocuta (Spotted hyena) |
| F1PFK6 | KPNA1 | Mammalia | Canis lupus familiaris (Dog) (Canis familiaris) |
| G1SI36 | KPNA1 | Mammalia | Oryctolagus cuniculus (Rabbit) |
| A0A2K5WRD1 | KPNA1 | Mammalia | Macaca fascicularis (Crab-eating macaque) (Cynomolgus monkey) |
| A0A6P6H2Z3 | KPNA1 | Mammalia | Puma concolor (Mountain lion) |
| G1NUG8 | KPNA1 | Mammalia | Myotis lucifugus (Little brown bat) |
| A0A6P5B348 | KPNA1 | Mammalia | Bos indicus (Zebu) |
| A0A6J2L0N9 | KPNA1 | Mammalia | Phyllostomus discolor (pale spear-nosed bat) |
| W5QFX7 | KPNA1 | Mammalia | Ovis aries (Sheep) |
| A0A1D5QJL1 | KPNA1 | Mammalia | Macaca mulatta (Rhesus macaque) |
| A0A6P3H099 | KPNA1 | Mammalia | Bison bison bison |
| F7ISW6 | KPNA1 | Mammalia | Callithrix jacchus (White-tufted-ear marmoset) |
| A0A2K6NKC7 | KPNA1 | Mammalia | Rhinopithecus roxellana (Golden snub-nosed monkey) (Pygathrix roxellana) |
| A0A3Q7RJW3 | KPNA1 | Mammalia | Vulpes vulpes (Red fox) |
| A0A6J0Z446 | KPNA1 | Mammalia | Odocoileus virginianus texanus |
| A0A3Q1LXF0 | KPNA1 | Mammalia | Bos taurus (Bovine) |
| A0A2K5RVS8 | KPNA1 | Mammalia | Cebus imitator (Panamanian white-faced capuchin) (Cebus capucinus imitator) |
| A0A6J1ZKI4 | KPNA1 | Mammalia | Acinonyx jubatus (Cheetah) |
| A0A2U3YI58 | KPNA1 | Mammalia | Leptonychotes weddellii (Weddell seal) (Otaria weddellii) |
| H2QN78 | KPNA1 | Mammalia | Pan troglodytes (Chimpanzee) |
| A0A6J2C9M7 | KPNA1 | Mammalia | Zalophus californianus (California sealion) |
| A0A337S696 | KPNA1 | Mammalia | Felis catus (Cat) (Felis silvestris catus) |
| F6PVH5 | KPNA1 | Mammalia | Monodelphis domestica (Gray short-tailed opossum) |
| A0A2Y9SJ29 | KPNA1 | Mammalia | Physeter macrocephalus (Sperm whale) (Physeter catodon) |
| A0A2K5N988 | KPNA1 | Mammalia | Cercocebus atys (Sooty mangabey) (Cercocebus torquatus atys) |
| A0A2Y9HG96 | KPNA1 | Mammalia | Neomonachus schauinslandi (Hawaiian monk seal) (Monachus schauinslandi) |
| A0A4W2FGB4 | KPNA1 | Mammalia | Bos indicus x Bos taurus (Hybrid cattle) |
| A0A2K5DU08 | KPNA1 | Mammalia | Aotus nancymaae (Ma's night monkey) |
| A0A2I3MWI4 | KPNA1 | Mammalia | Papio anubis (Olive baboon) |
| A0A4W2FGM4 | KPNA1 | Mammalia | Bos indicus x Bos taurus (Hybrid cattle) |
| A0A384DH80 | KPNA1 | Mammalia | Ursus maritimus (Polar bear) (Thalarctos maritimus) |
| A0A671DWC4 | KPNA1 | Mammalia | Rhinolophus ferrumequinum (Greater horseshoe bat) |
| A0A6J3H055 | KPNA1 | Mammalia | Sapajus apella (Brown-capped capuchin) (Cebus apella) |
| A0A6P5K899 | KPNA1 | Mammalia | Phascolarctos cinereus (Koala) |
| A0A2K5Y941 | KPNA1 | Mammalia | Mandrillus leucophaeus (Drill) (Papio leucophaeus) |
| F6QFP5 | KPNA1 | Mammalia | Equus caballus (Horse) |
| A0A0D9R4X1 | KPNA1 | Mammalia | Chlorocebus sabaeus (Green monkey) (Cercopithecus sabaeus) |
| G3UJ13 | KPNA1 | Mammalia | Loxodonta africana (African elephant) |
| A0A673T529 | KPNA1 | Mammalia | Suricata suricatta (Meerkat) |
| A0A3Q7PAX2 | KPNA1 | Mammalia | Callorhinus ursinus (Northern fur seal) |
| A0A2I3TP22 | KPNA1 | Mammalia | Pan troglodytes (Chimpanzee) |
| A0A2R9BX67 | KPNA1 | Mammalia | Pan paniscus (Pygmy chimpanzee) (Bonobo) |
| A0A340WM35 | KPNA1 | Mammalia | Lipotes vexillifer (Yangtze river dolphin) |
| A0A6P3R5V2 | KPNA1 | Mammalia | Pteropus vampyrus (Large flying fox) |
| A0A2U4AAL0 | KPNA1 | Mammalia | Tursiops truncatus (Atlantic bottle-nosed dolphin) (Delphinus truncatus) |
| A0A383ZKK2 | KPNA1 | Mammalia | Balaenoptera acutorostrata scammoni (North Pacific minke whale) (Balaenoptera davidsoni) |
| A0A2U3VBV0 | KPNA1 | Mammalia | Odobenus rosmarus divergens (Pacific walrus) |
| A0A2Y9MUZ0 | KPNA1 | Mammalia | Delphinapterus leucas (Beluga whale) |
| A0A2K5IUS9 | KPNA1 | Mammalia | Colobus angolensis palliatus (Peters' Angolan colobus) |
| A0A6I9IM75 | KPNA1 | Mammalia | Vicugna pacos (Alpaca) (Lama pacos) |
| A0A2K6TSQ2 | KPNA1 | Mammalia | Saimiri boliviensis boliviensis (Bolivian squirrel monkey) |
| M3Y068 | KPNA1 | Mammalia | Mustela putorius furo (European domestic ferret) (Mustela furo) |
| A0A452RY52 | KPNA1 | Mammalia | Ursus americanus (American black bear) (Euarctos americanus) |
| A0A2K6EIA0 | KPNA1 | Mammalia | Propithecus coquereli (Coquerel's sifaka) (Propithecus verreauxi coquereli) |
| A0A4X2KDM2 | KPNA1 | Mammalia | Vombatus ursinus (Common wombat) |
| A0A384DH77 | KPNA1 | Mammalia | Ursus maritimus (Polar bear) (Thalarctos maritimus) |
| A0A2K6LM96 | KPNA1 | Mammalia | Rhinopithecus bieti (Black snub-nosed monkey) (Pygathrix bieti) |
| A0A2K6DHG5 | KPNA1 | Mammalia | Macaca nemestrina (Pig-tailed macaque) |
| A0A6P4VZJ1 | KPNA1 | Mammalia | Panthera pardus (Leopard) (Felis pardus) |
| A0A3Q7WJB8 | KPNA1 | Mammalia | Ursus arctos horribilis |
| A0A2B4RY91 | KPNA2 | Anthozoa | Stylophora pistillata (Smooth cauliflower coral) |
| T2MFS8 | KPNA2 | Hydrozoa | Hydra vulgaris (Hydra) (Hydra attenuata) |
| A0A4Y2GZE4 | KPNA2 | Arachnida | Araneus ventricosus (Orbweaver spider) (Epeira ventricosa) |
| A0A5N5TKN5 | KPNA2 | Malacostraca | Armadillidium nasatum |
| A0A5B7IN23 | KPNA2 | Malacostraca | Portunus trituberculatus (Swimming crab) (Neptunus trituberculatus) |

| AC | GN | Class | Organism |
|---|---|---|---|
| A0A2H8THT7 | KPNA2 | Insecta | Melanaphis sacchari |
| A0A2J7Q2F0 | KPNA2 | Insecta | Cryptotermes secundus |
| A0A0A9Y1I6 | KPNA2 | Insecta | Lygus hesperus (Western plant bug) |
| A0A2P8XZQ8 | KPNA2 | Insecta | Blattella germanica (German cockroach) (Blatta germanica) |
| A0A2S2NCU0 | KPNA2 | Insecta | Schizaphis graminum (Green bug aphid) |
| A0A6A4VAT7 | KPNA2 | Hexanauplia | Amphibalanus amphitrite (Striped barnacle) (Balanus amphitrite) |
| A0A4W3KKI1 | KPNA2 | Chondrichthyes | Callorhinchus milii (Ghost shark) |
| A0A668SPP6 | KPNA2 | Actinopterygii | Oreochromis aureus (Israeli tilapia) (Chromis aureus) |
| A0A3P9AX06 | KPNA2 | Actinopterygii | Maylandia zebra (zebra mbuna) |
| A0A6P7J9M6 | KPNA2 | Actinopterygii | Parambassis ranga (Indian glassy fish) |
| A0A1A7ZW05 | KPNA2 | Actinopterygii | Nothobranchius furzeri (Turquoise killifish) |
| A0A1A7XN45 | KPNA2 | Actinopterygii | Iconisemion striatum |
| A0A1A8SK23 | KPNA2 | Actinopterygii | Nothobranchius rachovii (bluefin notho) |
| A0A1A8QNZ3 | KPNA2 | Actinopterygii | Nothobranchius rachovii (bluefin notho) |
| A0A668SNT1 | KPNA2 | Actinopterygii | Oreochromis aureus (Israeli tilapia) (Chromis aureus) |
| A0A3B4WCW0 | KPNA2 | Actinopterygii | Seriola lalandi dorsalis |
| A0A665VN51 | KPNA2 | Actinopterygii | Echeneis naucrates (Live sharksucker) |
| A0A3B4EME2 | KPNA2 | Actinopterygii | Pygocentrus nattereri (Red-bellied piranha) |
| A0A3Q3VX72 | KPNA2 | Actinopterygii | Mola mola (Ocean sunfish) (Tetraodon mola) |
| A0A665VN42 | KPNA2 | Actinopterygii | Echeneis naucrates (Live sharksucker) |
| A0A1A7YRI1 | KPNA2 | Actinopterygii | Iconisemion striatum |
| A0A3Q2PPK5 | KPNA2 | Actinopterygii | Fundulus heteroclitus (Killifish) (Mummichog) |
| A0A665VNE5 | KPNA2 | Actinopterygii | Echeneis naucrates (Live sharksucker) |
| A0A3Q3F6G4 | KPNA2 | Actinopterygii | Labrus bergylta (ballan wrasse) |
| A0A3P9MSI2 | KPNA2 | Actinopterygii | Poecilia reticulata (Guppy) (Acanthophacelus reticulatus) |
| Q6DI01 | KPNA2 | Actinopterygii | Danio rerio (Zebrafish) (Brachydanio rerio) |
| A0A1A8DGG9 | KPNA2 | Actinopterygii | Nothobranchius kadleci |
| A0A1A8PWJ9 | KPNA2 | Actinopterygii | Nothobranchius pienaari |
| A0A3P8YDR8 | KPNA2 | Actinopterygii | Esox lucius (Northern pike) |
| A0A671TR35 | KPNA2 | Actinopterygii | Sparus aurata (Gilthead sea bream) |
| A0A671TQX5 | KPNA2 | Actinopterygii | Sparus aurata (Gilthead sea bream) |
| A0A3B3ZC03 | KPNA2 | Actinopterygii | Periophthalmus magnuspinnatus |
| A0A3B3ZD83 | KPNA2 | Actinopterygii | Periophthalmus magnuspinnatus |
| I3JGZ1 | KPNA2 | Actinopterygii | Oreochromis niloticus (Nile tilapia) (Tilapia nilotica) |
| A0A671TQV7 | KPNA2 | Actinopterygii | Sparus aurata (Gilthead sea bream) |
| A0A1A8J5N3 | KPNA2 | Actinopterygii | Nothobranchius kuhntae (Beira killifish) |
| H2MF99 | KPNA2 | Actinopterygii | Oryzias latipes (Japanese rice fish) (Japanese killifish) |
| A0A3P9MPA0 | KPNA2 | Actinopterygii | Oryzias latipes (Japanese rice fish) (Japanese killifish) |
| A0A673MQV0 | KPNA2 | Actinopterygii | Sinocyclocheilus rhinocerous |
| A0A1A8IJ41 | KPNA2 | Actinopterygii | Nothobranchius kuhntae (Beira killifish) |
| A0A673MIX7 | KPNA2 | Actinopterygii | Sinocyclocheilus rhinocerous |
| A0A669F8L0 | KPNA2 | Actinopterygii | Oreochromis niloticus (Nile tilapia) (Tilapia nilotica) |
| A0A3P8NG90 | KPNA2 | Actinopterygii | Astatotilapia calliptera (Eastern happy) (Chromis callipterus) |
| A0A3Q1HQH3 | KPNA2 | Actinopterygii | Anabas testudineus (Climbing perch) (Anthias testudineus) |
| A0A1A8L5M0 | KPNA2 | Actinopterygii | Nothobranchius pienaari |
| A0A1A8U0E9 | KPNA2 | Actinopterygii | Nothobranchius furzeri (Turquoise killifish) |
| A0A671P8Z3 | KPNA2 | Actinopterygii | Sinocyclocheilus anshuiensis |
| A0A3B3YGX7 | KPNA2 | Actinopterygii | Poecilia mexicana |
| A0A1A8EWU9 | KPNA2 | Actinopterygii | Nothobranchius korthausae |
| A0A3Q3JK14 | KPNA2 | Actinopterygii | Monopterus albus (Swamp eel) |
| A0A3B4V3Q9 | KPNA2 | Actinopterygii | Seriola dumerili (Greater amberjack) (Caranx dumerili) |
| A0A3Q3B1Q0 | KPNA2 | Actinopterygii | Kryptolebias marmoratus (Mangrove killifish) (Rivulus marmoratus) |
| A0A6Q2WUF4 | KPNA2 | Actinopterygii | Esox lucius (Northern pike) |
| A0A3B3TJ61 | KPNA2 | Actinopterygii | Poecilia latipinna (sailfin molly) |
| A0A3Q0RB15 | KPNA2 | Actinopterygii | Amphilophus citrinellus (Midas cichlid) (Cichlasoma citrinellum) |
| A0A673BHL1 | KPNA2 | Actinopterygii | Sphaeramia orbicularis (orbiculate cardinalfish) |
| A0A668AE89 | KPNA2 | Actinopterygii | Myripristis murdjan (pinecone soldierfish) |
| A0A286R5L0 | KPNA2 | Actinopterygii | Channa punctata (Spotted snakehead) (Ophicephalus punctatus) |
| A0A674D460 | KPNA2 | Actinopterygii | Salmo trutta (Brown trout) |
| A0A668A4Z8 | KPNA2 | Actinopterygii | Myripristis murdjan (pinecone soldierfish) |
| A0A3B3Z282 | KPNA2 | Actinopterygii | Poecilia mexicana |
| A0A3B4GL06 | KPNA2 | Actinopterygii | Pundamilia nyererei |
| A0A1A8C3Z5 | KPNA2 | Actinopterygii | Nothobranchius kadleci |
| A0A4W6FTT1 | KPNA2 | Actinopterygii | Lates calcarifer (Barramundi) (Holocentrus calcarifer) |
| A0A672HYS4 | KPNA2 | Actinopterygii | Salarias fasciatus (Jewelled blenny) (Blennius fasciatus) |
| A0A672T5X6 | KPNA2 | Actinopterygii | Sinocyclocheilus grahami (Dianchi golden-line fish) (Barbus grahami) |
| M4AGR3 | KPNA2 | Actinopterygii | Xiphophorus maculatus (Southern platyfish) (Platypoecilus maculatus) |
| A0A4Z2FV23 | KPNA2 | Actinopterygii | Liparis tanakae (Tanaka's snailfish) |
| A0A6P8W0W9 | KPNA2 | Actinopterygii | Gymnodraco acuticeps (Antarctic dragonfish) |
| A0A3Q3MAK0 | KPNA2 | Actinopterygii | Mastacembelus armatus (zig-zag eel) |
| A0A673BLF7 | KPNA2 | Actinopterygii | Sphaeramia orbicularis (orbiculate cardinalfish) |
| A0A2D0S4E7 | KPNA2 | Actinopterygii | Ictalurus punctatus (Channel catfish) (Silurus punctatus) |
| A0A674BIA8 | KPNA2 | Actinopterygii | Salmo trutta (Brown trout) |
| A0A3B3RWN1 | KPNA2 | Actinopterygii | Paramormyrops kingsleyae |
| A0A087Y8H4 | KPNA2 | Actinopterygii | Poecilia formosa (Amazon molly) (Limia formosa) |
| A0A3Q1B5I6 | KPNA2 | Actinopterygii | Amphiprion ocellaris (Clown anemonefish) |
| G3Q3L2 | KPNA2 | Actinopterygii | Gasterosteus aculeatus (Three-spined stickleback) |
| A0A3Q4G708 | KPNA2 | Actinopterygii | Neolamprologus brichardi (Fairy cichlid) (Lamprologus brichardi) |
| A0A3B3RXW3 | KPNA2 | Actinopterygii | Paramormyrops kingsleyae |
| A0A3Q2VX29 | KPNA2 | Actinopterygii | Haplochromis burtoni (Burton's mouthbrooder) (Chromis burtoni) |
| A0A4Z2G449 | KPNA2 | Actinopterygii | Liparis tanakae (Tanaka's snailfish) |
| A0A668SNT5 | KPNA2 | Actinopterygii | Oreochromis aureus (Israeli tilapia) (Chromis aureus) |
| W5L7R7 | KPNA2 | Actinopterygii | Astyanax mexicanus (Blind cave fish) (Astyanax fasciatus mexicanus) |

| AC | GN | Class | Organism |
|---|---|---|---|
| A0A671PG44 | KPNA2 | Actinopterygii | Sinocyclocheilus anshuiensis |
| A0A3B5BN85 | KPNA2 | Actinopterygii | Stegastes partitus (bicolor damselfish) |
| A0A3P8RZS7 | KPNA2 | Actinopterygii | Amphiprion percula (Orange clownfish) (Lutjanus percula) |
| A0A3B3BT30 | KPNA2 | Actinopterygii | Oryzias melastigma (Marine medaka) |
| A0A2I4CGJ0 | KPNA2 | Actinopterygii | Austrofundulus limnaeus |
| A0A1A8GM44 | KPNA2 | Actinopterygii | Nothobranchius korthausae |
| A0A3P8VN82 | KPNA2 | Actinopterygii | Cynoglossus semilaevis (Tongue sole) |
| H3B358 | KPNA2 | Sarcopterygii | Latimeria chalumnae (Coelacanth) |
| A0A6I8QW41 | KPNA2 | Amphibia | Xenopus tropicalis (Western clawed frog) (Silurana tropicalis) |
| Q66IF6 | KPNA2 | Amphibia | Xenopus tropicalis (Western clawed frog) (Silurana tropicalis) |
| A0A6P7YMD2 | KPNA2 | Amphibia | Microcaecilia unicolor |
| Q7ZX00 | KPNA2 | Amphibia | Xenopus laevis (African clawed frog) |
| Q7ZYI0 | KPNA2 | Amphibia | Xenopus laevis (African clawed frog) |
| A0A6P8SSA5 | KPNA2 | Amphibia | Geotrypetes seraphini (Gaboon caecilian) (Caecilia seraphini) |
| P52171 | KPNA2 | Amphibia | Xenopus laevis (African clawed frog) |
| A0A674H5U9 | KPNA2 | Reptilia | Taeniopygia guttata (Zebra finch) (Poephila guttata) |
| A0A669Q9B1 | KPNA2 | Reptilia | Phasianus colchicus (Common pheasant) |
| A0A670HXI0 | KPNA2 | Reptilia | Podarcis muralis (Wall lizard) (Lacerta muralis) |
| A0A6J0GYH6 | KPNA2 | Reptilia | Lepidothrix coronata (blue-crowned manakin) |
| A0A663M287 | KPNA2 | Reptilia | Athene cunicularia (Burrowing owl) (Speotyto cunicularia) |
| A0A674IKE5 | KPNA2 | Reptilia | Terrapene carolina triunguis (Three-toed box turtle) |
| A0A6J2IZT9 | KPNA2 | Reptilia | Pipra filicauda (Wire-tailed manakin) |
| H0Z2A4 | KPNA2 | Reptilia | Taeniopygia guttata (Zebra finch) (Poephila guttata) |
| A0A452I6U0 | KPNA2 | Reptilia | Gopherus agassizii (Agassiz's desert tortoise) |
| U3JI37 | KPNA2 | Reptilia | Ficedula albicollis (Collared flycatcher) (Muscicapa albicollis) |
| H9G3N8 | KPNA2 | Reptilia | Anolis carolinensis (Green anole) (American chameleon) |
| A0A6J0SWD6 | KPNA2 | Reptilia | Pogona vitticeps (central bearded dragon) |
| V8PBZ1 | KPNA2 | Reptilia | Ophiophagus hannah (King cobra) (Naja hannah) |
| A0A1L1RX89 | KPNA2 | Reptilia | Gallus gallus (Chicken) |
| K7FQJ3 | KPNA2 | Reptilia | Pelodiscus sinensis (Chinese softshell turtle) (Trionyx sinensis) |
| A0A091QRY2 | KPNA2 | Reptilia | Haliaeetus albicilla (White-tailed sea-eagle) (Falco albicilla) |
| A0A670YIX9 | KPNA2 | Reptilia | Pseudonaja textilis (Eastern brown snake) |
| U3IRY8 | KPNA2 | Reptilia | Anas platyrhynchos platyrhynchos (Northern mallard) |
| A0A6P9BCB7 | KPNA2 | Reptilia | Pantherophis guttatus (Corn snake) (Elaphe guttata) |
| A0A6I9YEM7 | KPNA2 | Reptilia | Thamnophis sirtalis |
| A0A6J1UUA8 | KPNA2 | Reptilia | Notechis scutatus (mainland tiger snake) |
| A0A6J3E0U5 | KPNA2 | Reptilia | Aythya fuligula (Tufted duck) (Anas fuligula) |
| A0A663E939 | KPNA2 | Reptilia | Aquila chrysaetos chrysaetos |
| G1N6L6 | KPNA2 | Reptilia | Meleagris gallopavo (Wild turkey) |
| A0A672VFD3 | KPNA2 | Reptilia | Strigops habroptila (Kakapo) |
| J3KS65 | KPNA2 | Mammalia | Homo sapiens (Human) |
| Q6P6T9 | KPNA2 | Mammalia | Rattus norvegicus (Rat) |
| I3NFF1 | KPNA2 | Mammalia | Ictidomys tridecemlineatus (Thirteen-lined ground squirrel) (Spermophilus tridecemlineatus) |
| A0A250YJD2 | KPNA2 | Mammalia | Castor canadensis (American beaver) |
| A0A3Q0CQD4 | KPNA2 | Mammalia | Mesocricetus auratus (Golden hamster) |
| A0A6I9KZ87 | KPNA2 | Mammalia | Peromyscus maniculatus bairdii (Prairie deer mouse) |
| A6PW68 | KPNA2 | Mammalia | Mus musculus (Mouse) |
| A0A1S3H041 | KPNA2 | Mammalia | Dipodomys ordii (Ord's kangaroo rat) |
| A0A6P5QND8 | KPNA2 | Mammalia | Mus caroli (Ryukyu mouse) (Ricefield mouse) |
| A0A0P6IWF6 | KPNA2 | Mammalia | Heterocephalus glaber (Naked mole rat) |
| A0A1S3A179 | KPNA2 | Mammalia | Erinaceus europaeus (Western European hedgehog) |
| F6RK29 | KPNA2 | Mammalia | Ornithorhynchus anatinus (Duckbill platypus) |
| A0A287AHI1 | KPNA2 | Mammalia | Sus scrofa (Pig) |
| A0A452G8T5 | KPNA2 | Mammalia | Capra hircus (Goat) |
| G3TP33 | KPNA2 | Mammalia | Loxodonta africana (African elephant) |
| A0A6P5DHY5 | KPNA2 | Mammalia | Bos indicus (Zebu) |
| A0A6G1ACR9 | KPNA2 | Mammalia | Crocuta crocuta (Spotted hyena) |
| A0A2I3H982 | KPNA2 | Mammalia | Nomascus leucogenys (Northern white-cheeked gibbon) (Hylobates leucogenys) |
| A0A2I2ZHY4 | KPNA2 | Mammalia | Gorilla gorilla gorilla (Western lowland gorilla) |
| A0A667I0F1 | KPNA2 | Mammalia | Lynx canadensis (Canada lynx) |
| A0A6J3EXS8 | KPNA2 | Mammalia | Sapajus apella (Brown-capped capuchin) (Cebus apella) |
| A0A6P5LDL0 | KPNA2 | Mammalia | Phascolarctos cinereus (Koala) |
| G1QDN1 | KPNA2 | Mammalia | Myotis lucifugus (Little brown bat) |
| E2R6L9 | KPNA2 | Mammalia | Canis lupus familiaris (Dog) (Canis familiaris) |
| G1MJD5 | KPNA2 | Mammalia | Ailuropoda melanoleuca (Giant panda) |
| A0A6J2MAG7 | KPNA2 | Mammalia | Phyllostomus discolor (pale spear-nosed bat) |
| F6VLU3 | KPNA2 | Mammalia | Macaca mulatta (Rhesus macaque) |
| U3EY78 | KPNA2 | Mammalia | Callithrix jacchus (White-tufted-ear marmoset) |
| A0A6J0YM57 | KPNA2 | Mammalia | Odocoileus virginianus texanus |
| W5Q181 | KPNA2 | Mammalia | Ovis aries (Sheep) |
| U3EQN0 | KPNA2 | Mammalia | Callithrix jacchus (White-tufted-ear marmoset) |
| A0A6P9F491 | KPNA2 | Mammalia | Zalophus californianus (California sealion) |
| K7C276 | KPNA2 | Mammalia | Pan troglodytes (Chimpanzee) |
| Q3SYV6 | KPNA2 | Mammalia | Bos taurus (Bovine) |
| A0A3Q7WT66 | KPNA2 | Mammalia | Ursus arctos horribilis |
| F7D853 | KPNA2 | Mammalia | Monodelphis domestica (Gray short-tailed opossum) |
| A0A2Y9G4Z0 | KPNA2 | Mammalia | Neomonachus schauinslandi (Hawaiian monk seal) (Monachus schauinslandi) |
| A0A2U3XDN0 | KPNA2 | Mammalia | Leptonychotes weddellii (Weddell seal) (Otaria weddellii) |
| H2NUI7 | KPNA2 | Mammalia | Pongo abelii (Sumatran orangutan) (Pongo pygmaeus abelii) |
| A0A3Q7S7L5 | KPNA2 | Mammalia | Vulpes vulpes (Red fox) |
| A0A6J1YFZ8 | KPNA2 | Mammalia | Acinonyx jubatus (Cheetah) |
| A0A6P3IU58 | KPNA2 | Mammalia | Bison bison bison |
| A0A6P4X1F9 | KPNA2 | Mammalia | Panthera pardus (Leopard) (Felis pardus) |

| AC | GN | Class | Organism |
|---|---|---|---|
| A0A4W2E8W0 | KPNA2 | Mammalia | Bos indicus x Bos taurus (Hybrid cattle) |
| A0A341CRR5 | KPNA2 | Mammalia | Neophocaena asiaeorientalis asiaeorientalis (Yangtze finless porpoise) (Neophocaena phocaenoides subsp. asiaeorientalis) |
| A0A3Q2H8G1 | KPNA2 | Mammalia | Equus caballus (Horse) |
| A0A2R9CTC7 | KPNA2 | Mammalia | Pan paniscus (Pygmy chimpanzee) (Bonobo) |
| A0A0D9QW15 | KPNA2 | Mammalia | Chlorocebus sabaeus (Green monkey) (Cercopithecus sabaeus) |
| A0A340XCA9 | KPNA2 | Mammalia | Lipotes vexillifer (Yangtze river dolphin) |
| A0A6P3QVC4 | KPNA2 | Mammalia | Pteropus vampyrus (Large flying fox) |
| A0A2U4C3S8 | KPNA2 | Mammalia | Tursiops truncatus (Atlantic bottle-nosed dolphin) (Delphinus truncatus) |
| A0A384AFI2 | KPNA2 | Mammalia | Balaenoptera acutorostrata scammoni (North Pacific minke whale) (Balaenoptera davidsoni) |
| A0A6I9I0T4 | KPNA2 | Mammalia | Vicugna pacos (Alpaca) (Lama pacos) |
| A0A2Y9PDA8 | KPNA2 | Mammalia | Delphinapterus leucas (Beluga whale) |
| A0A2U3ZL89 | KPNA2 | Mammalia | Odobenus rosmarus divergens (Pacific walrus) |
| A0A452UTX9 | KPNA2 | Mammalia | Ursus maritimus (Polar bear) (Thalarctos maritimus) |
| A0A4X2LEJ2 | KPNA2 | Mammalia | Vombatus ursinus (Common wombat) |
| A0A6P6I5W2 | KPNA2 | Mammalia | Puma concolor (Mountain lion) |
| H0VVB1 | KPNA2 | Mammalia | Cavia porcellus (Guinea pig) |
| Q71VM5 | KPNA3 | Hydrozoa | Hydra vulgaris (Hydra) (Hydra attenuata) |
| G7YSA1 | KPNA3 | Trematoda | Clonorchis sinensis (Chinese liver fluke) |
| A0A0V1J5Y5 | KPNA3 | Enoplea | Trichinella pseudospiralis (Parasitic roundworm) |
| A0A0V1HFN1 | KPNA3 | Enoplea | Trichinella zimbabwensis |
| A0A0V1BD23 | KPNA3 | Enoplea | Trichinella spiralis (Trichina worm) |
| A0A0V0TWQ0 | KPNA3 | Enoplea | Trichinella murrelli |
| A0A0V1P443 | KPNA3 | Enoplea | Trichinella sp. T8 |
| A0A0V1FJA8 | KPNA3 | Enoplea | Trichinella pseudospiralis (Parasitic roundworm) |
| A0A0V1N4W6 | KPNA3 | Enoplea | Trichinella papuae |
| A0A4C1V8D9 | KPNA3 | Insecta | Eumeta variegata (Bagworm moth) (Eumeta japonica) |
| A0A2S2R6K7 | KPNA3 | Insecta | Sipha flava (yellow sugarcane aphid) |
| A0A2S2NHS8 | KPNA3 | Insecta | Schizaphis graminum (Green bug aphid) |
| A0A6J2RYW0 | KPNA3 | Actinopterygii | Cottoperca gobio (Frogmouth) (Aphritis gobio) |
| A0A6P7HJQ6 | KPNA3 | Actinopterygii | Parambassis ranga (Indian glassy fish) |
| A0A6P7LAX0 | KPNA3 | Actinopterygii | Betta splendens (Siamese fighting fish) |
| A0A1A8Q8V8 | KPNA3 | Actinopterygii | Nothobranchius rachovii (bluefin notho) |
| A0A1A7ZEA1 | KPNA3 | Actinopterygii | Nothobranchius furzeri (Turquoise killifish) |
| A0A1A8VCM4 | KPNA3 | Actinopterygii | Nothobranchius furzeri (Turquoise killifish) |
| A0A6I9NPM0 | KPNA3 | Actinopterygii | Notothenia coriiceps (black rockcod) |
| A0A665V3R5 | KPNA3 | Actinopterygii | Echeneis naucrates (Live sharksucker) |
| A0A4W4HIJ1 | KPNA3 | Actinopterygii | Electrophorus electricus (Electric eel) (Gymnotus electricus) |
| A0A6P3WEA8 | KPNA3 | Actinopterygii | Clupea harengus (Atlantic herring) |
| A0A674N306 | KPNA3 | Actinopterygii | Takifugu rubripes (Japanese pufferfish) (Fugu rubripes) |
| A0A674NY42 | KPNA3 | Actinopterygii | Takifugu rubripes (Japanese pufferfish) (Fugu rubripes) |
| A0A1A8PEW1 | KPNA3 | Actinopterygii | Nothobranchius pienaari |
| A0A1A8LI71 | KPNA3 | Actinopterygii | Nothobranchius pienaari |
| A0A4W5L687 | KPNA3 | Actinopterygii | Hucho hucho (huchen) |
| A0A4W5PPZ0 | KPNA3 | Actinopterygii | Hucho hucho (huchen) |
| A0A1A8JPR2 | KPNA3 | Actinopterygii | Nothobranchius kuhntae (Beira killifish) |
| A0A1A8ILH3 | KPNA3 | Actinopterygii | Nothobranchius kuhntae (Beira killifish) |
| Q8JFT7 | KPNA3 | Actinopterygii | Danio rerio (Zebrafish) (Brachydanio rerio) |
| A0A1A7X5R1 | KPNA3 | Actinopterygii | Iconisemion striatum |
| A0A671MPH1 | KPNA3 | Actinopterygii | Sinocyclocheilus anshuiensis |
| A0A1A8F687 | KPNA3 | Actinopterygii | Nothobranchius korthausae |
| A0A4W5L543 | KPNA3 | Actinopterygii | Hucho hucho (huchen) |
| A0A672Z3N3 | KPNA3 | Actinopterygii | Sphaeramia orbicularis (orbiculate cardinalfish) |
| A0A667Z8B4 | KPNA3 | Actinopterygii | Myripristis murdjan (pinecone soldierfish) |
| A0A1A8BPJ4 | KPNA3 | Actinopterygii | Nothobranchius kadleci |
| W5UB95 | KPNA3 | Actinopterygii | Ictalurus punctatus (Channel catfish) (Silurus punctatus) |
| A0A6P8UDL6 | KPNA3 | Actinopterygii | Gymnodraco acuticeps (Antarctic dragonfish) |
| A0A672HF09 | KPNA3 | Actinopterygii | Salarias fasciatus (Jewelled blenny) (Blennius fasciatus) |
| A0A672HEZ1 | KPNA3 | Actinopterygii | Salarias fasciatus (Jewelled blenny) (Blennius fasciatus) |
| A0A672Z3A8 | KPNA3 | Actinopterygii | Sphaeramia orbicularis (orbiculate cardinalfish) |
| A0A672PQJ6 | KPNA3 | Actinopterygii | Sinocyclocheilus grahami (Dianchi golden-line fish) (Barbus grahami) |
| A0A6J2VU04 | KPNA3 | Actinopterygii | Chanos chanos (Milkfish) (Mugil chanos) |
| A0A4Z2GJN9 | KPNA3 | Actinopterygii | Liparis tanakae (Tanaka's snailfish) |
| A0A2I4BHC9 | KPNA3 | Actinopterygii | Austrofundulus limnaeus |
| H3AV75 | KPNA3 | Sarcopterygii | Latimeria chalumnae (Coelacanth) |
| Q70PC6 | KPNA3 | Amphibia | Xenopus laevis (African clawed frog) |
| A0A6P7Y0L1 | KPNA3 | Amphibia | Microcaecilia unicolor |
| Q6IRB7 | KPNA3 | Amphibia | Xenopus laevis (African clawed frog) |
| A0A6I8QJZ7 | KPNA3 | Amphibia | Xenopus tropicalis (Western clawed frog) (Silurana tropicalis) |
| A0A6P8RJ78 | KPNA3 | Amphibia | Geotrypetes seraphini (Gaboon caecilian) (Caecilia seraphini) |
| A0A670I6K3 | KPNA3 | Reptilia | Podarcis muralis (Wall lizard) (Lacerta muralis) |
| A0A6P9CHE3 | KPNA3 | Reptilia | Pantherophis guttatus (Corn snake) (Elaphe guttata) |
| A0A6J1V4X7 | KPNA3 | Reptilia | Notechis scutatus (mainland tiger snake) |
| A0A6J0HA95 | KPNA3 | Reptilia | Lepidothrix coronata (blue-crowned manakin) |
| A0A151NB90 | KPNA3 | Reptilia | Alligator mississippiensis (American alligator) |
| A0A674HEZ2 | KPNA3 | Reptilia | Taeniopygia guttata (Zebra finch) (Poephila guttata) |
| A0A669QP80 | KPNA3 | Reptilia | Phasianus colchicus (Common pheasant) |
| A0A452IFV5 | KPNA3 | Reptilia | Gopherus agassizii (Agassiz's desert tortoise) |
| F1NV68 | KPNA3 | Reptilia | Gallus gallus (Chicken) |
| G1KPC3 | KPNA3 | Reptilia | Anolis carolinensis (Green anole) (American chameleon) |
| A0A6J3DF17 | KPNA3 | Reptilia | Aythya fuligula (Tufted duck) (Anas fuligula) |
| A0A6J2GPV2 | KPNA3 | Reptilia | Pipra filicauda (Wire-tailed manakin) |
| U3IJT7 | KPNA3 | Reptilia | Anas platyrhynchos platyrhynchos (Northern mallard) |
| A0A670Y0Q9 | KPNA3 | Reptilia | Pseudonaja textilis (Eastern brown snake) |

| AC | GN | Class | Organism |
|---|---|---|---|
| A0A6I9X8Y6 | KPNA3 | Reptilia | Thamnophis sirtalis |
| A0A218UQY7 | KPNA3 | Reptilia | Lonchura striata domestica (Bengalese finch) |
| A0A663F212 | KPNA3 | Reptilia | Aquila chrysaetos chrysaetos |
| A0A1V4J4R5 | KPNA3 | Reptilia | Patagioenas fasciata monilis |
| A0A672TI99 | KPNA3 | Reptilia | Strigops habroptila (Kakapo) |
| H0V3K6 | KPNA3 | Mammalia | Cavia porcellus (Guinea pig) |
| Q56R18 | KPNA3 | Mammalia | Rattus norvegicus (Rat) |
| A0A3Q0D085 | KPNA3 | Mammalia | Mesocricetus auratus (Golden hamster) |
| A0A6P5QYT5 | KPNA3 | Mammalia | Mus caroli (Ryukyu mouse) (Ricefield mouse) |
| Q9CT07 | KPNA3 | Mammalia | Mus musculus (Mouse) |
| A0A250Y2A9 | KPNA3 | Mammalia | Castor canadensis (American beaver) |
| A0A6P6EKZ5 | KPNA3 | Mammalia | Octodon degus (Degu) (Sciurus degus) |
| A0A0P6JDA1 | KPNA3 | Mammalia | Heterocephalus glaber (Naked mole rat) |
| A0A1S3AHI3 | KPNA3 | Mammalia | Erinaceus europaeus (Western European hedgehog) |
| A0A452FEC0 | KPNA3 | Mammalia | Capra hircus (Goat) |
| A0A5G2R9E6 | KPNA3 | Mammalia | Sus scrofa (Pig) |
| A0A6P5CMI9 | KPNA3 | Mammalia | Bos indicus (Zebu) |
| A0A6P5KU41 | KPNA3 | Mammalia | Phascolarctos cinereus (Koala) |
| A0A667GNB9 | KPNA3 | Mammalia | Lynx canadensis (Canada lynx) |
| A0A2K6R1J6 | KPNA3 | Mammalia | Rhinopithecus roxellana (Golden snub-nosed monkey) (Pygathrix roxellana) |
| A0A2K5RYQ6 | KPNA3 | Mammalia | Cebus imitator (Panamanian white-faced capuchin) (Cebus capucinus imitator) |
| A0A5F8A5V2 | KPNA3 | Mammalia | Macaca mulatta (Rhesus macaque) |
| A0A3Q1MUS6 | KPNA3 | Mammalia | Bos taurus (Bovine) |
| A0A6P3YND9 | KPNA3 | Mammalia | Ovis aries (Sheep) |
| H0WKJ7 | KPNA3 | Mammalia | Otolemur garnettii (Small-eared galago) (Garnett's greater bushbaby) |
| A0A4W2E6Z1 | KPNA3 | Mammalia | Bos indicus x Bos taurus (Hybrid cattle) |
| A0A2Y9H0K2 | KPNA3 | Mammalia | Neomonachus schauinslandi (Hawaiian monk seal) (Monachus schauinslandi) |
| A0A2K5MEL9 | KPNA3 | Mammalia | Cercocebus atys (Sooty mangabey) (Cercocebus torquatus atys) |
| A0A5F8GJT3 | KPNA3 | Mammalia | Monodelphis domestica (Gray short-tailed opossum) |
| M3WQJ0 | KPNA3 | Mammalia | Felis catus (Cat) (Felis silvestris catus) |
| A0A2R9AVB4 | KPNA3 | Mammalia | Pan paniscus (Pygmy chimpanzee) (Bonobo) |
| A0A6J1ZVC0 | KPNA3 | Mammalia | Acinonyx jubatus (Cheetah) |
| H2Q7K7 | KPNA3 | Mammalia | Pan troglodytes (Chimpanzee) |
| I7GIR2 | KPNA3 | Mammalia | Macaca fascicularis (Crab-eating macaque) (Cynomolgus monkey) |
| A0A2Y9EMI4 | KPNA3 | Mammalia | Physeter macrocephalus (Sperm whale) (Physeter catodon) |
| U3BKZ4 | KPNA3 | Mammalia | Callithrix jacchus (White-tufted-ear marmoset) |
| A0A0D9RYP1 | KPNA3 | Mammalia | Chlorocebus sabaeus (Green monkey) (Cercopithecus sabaeus) |
| A0A2U4BGI2 | KPNA3 | Mammalia | Tursiops truncatus (Atlantic bottle-nosed dolphin) (Delphinus truncatus) |
| A0A340Y4X1 | KPNA3 | Mammalia | Lipotes vexillifer (Yangtze river dolphin) |
| A0A2K5DTK7 | KPNA3 | Mammalia | Aotus nancymaae (Ma's night monkey) |
| A0A6J2CSB0 | KPNA3 | Mammalia | Zalophus californianus (California sealion) |
| A0A096NP23 | KPNA3 | Mammalia | Papio anubis (Olive baboon) |
| A0A3Q7PA97 | KPNA3 | Mammalia | Callorhinus ursinus (Northern fur seal) |
| A0A024RDV7 | KPNA3 | Mammalia | Homo sapiens (Human) |
| A0A2Y9NMF0 | KPNA3 | Mammalia | Delphinapterus leucas (Beluga whale) |
| A0A2U3VVQ1 | KPNA3 | Mammalia | Odobenus rosmarus divergens (Pacific walrus) |
| A0A6I8P1J4 | KPNA3 | Mammalia | Ornithorhynchus anatinus (Duckbill platypus) |
| A0A6I8PME1 | KPNA3 | Mammalia | Ornithorhynchus anatinus (Duckbill platypus) |
| A0A2K6FG80 | KPNA3 | Mammalia | Propithecus coquereli (Coquerel's sifaka) (Propithecus verreauxi coquereli) |
| A0A4X2M2X5 | KPNA3 | Mammalia | Vombatus ursinus (Common wombat) |
| A0A2K6MWD3 | KPNA3 | Mammalia | Rhinopithecus bieti (Black snub-nosed monkey) (Pygathrix bieti) |
| A0A6J3J7F8 | KPNA3 | Mammalia | Sapajus apella (Brown-capped capuchin) (Cebus apella) |
| A0A6J3J6G8 | KPNA3 | Mammalia | Sapajus apella (Brown-capped capuchin) (Cebus apella) |
| A0A2K6DCA2 | KPNA3 | Mammalia | Macaca nemestrina (Pig-tailed macaque) |
| A0A6P4VPZ6 | KPNA3 | Mammalia | Panthera pardus (Leopard) (Felis pardus) |
| A0A5F9CSY8 | KPNA3 | Mammalia | Oryctolagus cuniculus (Rabbit) |
| A0A3Q7UKJ0 | KPNA3 | Mammalia | Ursus arctos horribilis |
| A0A2B4S0M6 | KPNA4 | Anthozoa | Stylophora pistillata (Smooth cauliflower coral) |
| A0A0V0RTF5 | KPNA4 | Enoplea | Trichinella nelsoni |
| A0A4Y2Q2Z3 | KPNA4 | Arachnida | Araneus ventricosus (Orbweaver spider) (Epeira ventricosa) |
| A0A6G1S6S5 | KPNA4 | Arachnida | Aceria tosichella (wheat curl mite) |
| A0A5N5SW62 | KPNA4 | Malacostraca | Armadillidium nasatum |
| A0A2J7RBM0 | KPNA4 | Insecta | Cryptotermes secundus |
| A0A0C9RV98 | KPNA4 | Insecta | Fopius arisanus |
| A0A0A9WQY7 | KPNA4 | Insecta | Lygus hesperus (Western plant bug) |
| A0A0K8WEE2 | KPNA4 | Insecta | Bactrocera latifrons (Malaysian fruit fly) (Chaetodacus latifrons) |
| A0A0A1XJB6 | KPNA4 | Insecta | Zeugodacus cucurbitae (Melon fruit fly) (Bactrocera cucurbitae) |
| A0A2P8ZE70 | KPNA4 | Insecta | Blattella germanica (German cockroach) (Blatta germanica) |
| A0A6F9DGB8 | KPNA4 | Ascidiacea | Phallusia mammillata |
| A0A4W3JQE6 | KPNA4 | Chondrichthyes | Callorhinchus milii (Ghost shark) |
| A0A4W3JN51 | KPNA4 | Chondrichthyes | Callorhinchus milii (Ghost shark) |
| A0A4W3K100 | KPNA4 | Chondrichthyes | Callorhinchus milii (Ghost shark) |
| A0A6J2QXP4 | KPNA4 | Actinopterygii | Cottoperca gobio (Frogmouth) (Aphritis gobio) |
| A0A6P7PBN0 | KPNA4 | Actinopterygii | Betta splendens (Siamese fighting fish) |
| A0A2Z2GP46 | KPNA4 | Actinopterygii | Synbranchidae sp. HB-2017 |
| A0A4W4FLE0 | KPNA4 | Actinopterygii | Electrophorus electricus (Electric eel) (Gymnotus electricus) |
| A0A665UA99 | KPNA4 | Actinopterygii | Echeneis naucrates (Live sharksucker) |
| A0A665UA62 | KPNA4 | Actinopterygii | Echeneis naucrates (Live sharksucker) |
| A0A6I9NX86 | KPNA4 | Actinopterygii | Notothenia coriiceps (black rockcod) |
| A0A1S3T0D1 | KPNA4 | Actinopterygii | Salmo salar (Atlantic salmon) |
| A0A6P3WAK5 | KPNA4 | Actinopterygii | Clupea harengus (Atlantic herring) |
| A0A674PLW0 | KPNA4 | Actinopterygii | Takifugu rubripes (Japanese pufferfish) (Fugu rubripes) |
| I3J6S5 | KPNA4 | Actinopterygii | Oreochromis niloticus (Nile tilapia) (Tilapia nilotica) |

| AC | GN | Class | Organism |
| --- | --- | --- | --- |
| F1QEW3 | KPNA4 | Actinopterygii | Danio rerio (Zebrafish) (Brachydanio rerio) |
| A0A1A7YVU0 | KPNA4 | Actinopterygii | Iconisemion striatum |
| A0A1A8IH68 | KPNA4 | Actinopterygii | Nothobranchius kuhntae (Beira killifish) |
| A0A1A8QHT5 | KPNA4 | Actinopterygii | Nothobranchius rachovii (bluefin notho) |
| A0A1A8MER6 | KPNA4 | Actinopterygii | Nothobranchius pienaari |
| W5UA83 | KPNA4 | Actinopterygii | Ictalurus punctatus (Channel catfish) (Silurus punctatus) |
| A0A1A8U2Q0 | KPNA4 | Actinopterygii | Nothobranchius furzeri (Turquoise killifish) |
| A0A667XPZ3 | KPNA4 | Actinopterygii | Myripristis murdjan (pinecone soldierfish) |
| A0A4W6BU17 | KPNA4 | Actinopterygii | Lates calcarifer (Barramundi) (Holocentrus calcarifer) |
| A0A1A8FJ46 | KPNA4 | Actinopterygii | Nothobranchius korthausae |
| A0A1A8DEU3 | KPNA4 | Actinopterygii | Nothobranchius kadleci |
| A0A672FRF4 | KPNA4 | Actinopterygii | Salarias fasciatus (Jewelled blenny) (Blennius fasciatus) |
| A0A6P8WCL6 | KPNA4 | Actinopterygii | Gymnodraco acuticeps (Antarctic dragonfish) |
| A0A4Z2IS62 | KPNA4 | Actinopterygii | Liparis tanakae (Tanaka's snailfish) |
| A0A673ZKB1 | KPNA4 | Actinopterygii | Salmo trutta (Brown trout) |
| A0A673BC87 | KPNA4 | Actinopterygii | Sphaeramia orbicularis (orbiculate cardinalfish) |
| A0A673BBF6 | KPNA4 | Actinopterygii | Sphaeramia orbicularis (orbiculate cardinalfish) |
| A0A6J2VJW1 | KPNA4 | Actinopterygii | Chanos chanos (Milkfish) (Mugil chanos) |
| A0A673B6V3 | KPNA4 | Actinopterygii | Sphaeramia orbicularis (orbiculate cardinalfish) |
| A0A2I4CTN3 | KPNA4 | Actinopterygii | Austrofundulus limnaeus |
| A0A6P7JGS9 | KPNA4 | Actinopterygii | Parambassis ranga (Indian glassy fish) |
| H2LP51 | KPNA4 | Actinopterygii | Oryzias latipes (Japanese rice fish) (Japanese killifish) |
| H3AN54 | KPNA4 | Sarcopterygii | Latimeria chalumnae (Coelacanth) |
| F6TB69 | KPNA4 | Amphibia | Xenopus tropicalis (Western clawed frog) (Silurana tropicalis) |
| A0A6P7Z784 | KPNA4 | Amphibia | Microcaecilia unicolor |
| A0A1L8GA53 | KPNA4 | Amphibia | Xenopus laevis (African clawed frog) |
| A0A6P8S910 | KPNA4 | Amphibia | Geotrypetes seraphini (Gaboon caecilian) (Caecilia seraphini) |
| A0A674ILH1 | KPNA4 | Reptilia | Terrapene carolina triunguis (Three-toed box turtle) |
| A0A452HUT1 | KPNA4 | Reptilia | Gopherus agassizii (Agassiz's desert tortoise) |
| A0A674G8H0 | KPNA4 | Reptilia | Taeniopygia guttata (Zebra finch) (Poephila guttata) |
| A0A1U8DF45 | KPNA4 | Reptilia | Alligator sinensis (Chinese alligator) |
| A0A6P9DVB1 | KPNA4 | Reptilia | Pantherophis guttatus (Corn snake) (Elaphe guttata) |
| A0A6J1U177 | KPNA4 | Reptilia | Notechis scutatus (mainland tiger snake) |
| A0A6J2IMM6 | KPNA4 | Reptilia | Pipra filicauda (Wire-tailed manakin) |
| A0A669Q581 | KPNA4 | Reptilia | Phasianus colchicus (Common pheasant) |
| A0A670I2H4 | KPNA4 | Reptilia | Podarcis muralis (Wall lizard) (Lacerta muralis) |
| U3JFA4 | KPNA4 | Reptilia | Ficedula albicollis (Collared flycatcher) (Muscicapa albicollis) |
| A0A1V4KM60 | KPNA4 | Reptilia | Patagioenas fasciata monilis |
| V8PFW4 | KPNA4 | Reptilia | Ophiophagus hannah (King cobra) (Naja hannah) |
| A0A6I9YGU7 | KPNA4 | Reptilia | Thamnophis sirtalis |
| A0A3Q0GSB0 | KPNA4 | Reptilia | Alligator sinensis (Chinese alligator) |
| U3IJM3 | KPNA4 | Reptilia | Anas platyrhynchos platyrhynchos (Northern mallard) |
| A0A670Z644 | KPNA4 | Reptilia | Pseudonaja textilis (Eastern brown snake) |
| K7FLG7 | KPNA4 | Reptilia | Pelodiscus sinensis (Chinese softshell turtle) (Trionyx sinensis) |
| A0A663DS62 | KPNA4 | Reptilia | Aquila chrysaetos chrysaetos |
| Q5ZLF7 | KPNA4 | Reptilia | Gallus gallus (Chicken) |
| A0A672UI36 | KPNA4 | Reptilia | Strigops habroptila (Kakapo) |
| A0A663DS72 | KPNA4 | Reptilia | Aquila chrysaetos chrysaetos |
| A0A6J3DFR9 | KPNA4 | Reptilia | Aythya fuligula (Tufted duck) (Anas fuligula) |
| A0A6J0TFY7 | KPNA4 | Reptilia | Pogona vitticeps (central bearded dragon) |
| O00629 | KPNA4 | Mammalia | Homo sapiens (Human) |
| H0VHE6 | KPNA4 | Mammalia | Cavia porcellus (Guinea pig) |
| Q56R17 | KPNA4 | Mammalia | Rattus norvegicus (Rat) |
| A0A0B4J1E7 | KPNA4 | Mammalia | Mus musculus (Mouse) |
| A0A250YJK4 | KPNA4 | Mammalia | Castor canadensis (American beaver) |
| I3M4E9 | KPNA4 | Mammalia | Ictidomys tridecemlineatus (Thirteen-lined ground squirrel) (Spermophilus tridecemlineatus) |
| A0A6I9KX70 | KPNA4 | Mammalia | Peromyscus maniculatus bairdii (Prairie deer mouse) |
| A0A3Q0DIG1 | KPNA4 | Mammalia | Mesocricetus auratus (Golden hamster) |
| A0A6P7QP70 | KPNA4 | Mammalia | Mus caroli (Ryukyu mouse) (Ricefield mouse) |
| A0A6P5PG61 | KPNA4 | Mammalia | Mus caroli (Ryukyu mouse) (Ricefield mouse) |
| A0A6P6D5V3 | KPNA4 | Mammalia | Octodon degus (Degu) (Sciurus degus) |
| A0A2U3WCB1 | KPNA4 | Mammalia | Odobenus rosmarus divergens (Pacific walrus) |
| A0A1U7T2Y1 | KPNA4 | Mammalia | Carlito syrichta (Philippine tarsier) (Tarsius syrichta) |
| M3XV10 | KPNA4 | Mammalia | Mustela putorius furo (European domestic ferret) (Mustela furo) |
| A0A452GBK4 | KPNA4 | Mammalia | Capra hircus (Goat) |
| A0A3Q2I6U1 | KPNA4 | Mammalia | Equus caballus (Horse) |
| A0A5G2R9L2 | KPNA4 | Mammalia | Sus scrofa (Pig) |
| J9PAW2 | KPNA4 | Mammalia | Canis lupus familiaris (Dog) (Canis familiaris) |
| G1QXC4 | KPNA4 | Mammalia | Nomascus leucogenys (Northern white-cheeked gibbon) (Hylobates leucogenys) |
| A0A667HGJ4 | KPNA4 | Mammalia | Lynx canadensis (Canada lynx) |
| A0A6P5BS84 | KPNA4 | Mammalia | Bos indicus (Zebu) |
| A0A5F9DRC8 | KPNA4 | Mammalia | Oryctolagus cuniculus (Rabbit) |
| A0A0B8RSP3 | KPNA4 | Mammalia | Sus scrofa domesticus (domestic pig) |
| A0A6J2L227 | KPNA4 | Mammalia | Phyllostomus discolor (pale spear-nosed bat) |
| A0A6P5KG24 | KPNA4 | Mammalia | Phascolarctos cinereus (Koala) |
| A0A7E6D5F7 | KPNA4 | Mammalia | Phyllostomus discolor (pale spear-nosed bat) |
| A0A2K5WX73 | KPNA4 | Mammalia | Macaca fascicularis (Crab-eating macaque) (Cynomolgus monkey) |
| A0A2K5PA95 | KPNA4 | Mammalia | Cebus imitator (Panamanian white-faced capuchin) (Cebus capucinus imitator) |
| A0A3S5ZPE0 | KPNA4 | Mammalia | Bos taurus (Bovine) |
| F6VIF8 | KPNA4 | Mammalia | Callithrix jacchus (White-tufted-ear marmoset) |
| H9EQP0 | KPNA4 | Mammalia | Macaca mulatta (Rhesus macaque) |
| A0A5F4W1H4 | KPNA4 | Mammalia | Callithrix jacchus (White-tufted-ear marmoset) |
| A0A2K6NFI6 | KPNA4 | Mammalia | Rhinopithecus roxellana (Golden snub-nosed monkey) (Pygathrix roxellana) |

| AC | GN | Class | Organism |
|---|---|---|---|
| A0A671DSE8 | KPNA4 | Mammalia | Rhinolophus ferrumequinum (Greater horseshoe bat) |
| A0A6P3YIU1 | KPNA4 | Mammalia | Ovis aries (Sheep) |
| H0X6W3 | KPNA4 | Mammalia | Otolemur garnettii (Small-eared galago) (Garnett's greater bushbaby) |
| H2QNN5 | KPNA4 | Mammalia | Pan troglodytes (Chimpanzee) |
| A0A6J1ZKW3 | KPNA4 | Mammalia | Acinonyx jubatus (Cheetah) |
| A0A6J0Z606 | KPNA4 | Mammalia | Odocoileus virginianus texanus |
| A0A2Y9I3B6 | KPNA4 | Mammalia | Neomonachus schauinslandi (Hawaiian monk seal) (Monachus schauinslandi) |
| A0A3Q7T5I3 | KPNA4 | Mammalia | Vulpes vulpes (Red fox) |
| A0A7F8RJW8 | KPNA4 | Mammalia | Leptonychotes weddellii (Weddell seal) (Otaria weddellii) |
| A0A2K5NJH9 | KPNA4 | Mammalia | Cercocebus atys (Sooty mangabey) (Cercocebus torquatus atys) |
| A0A2Y9SU99 | KPNA4 | Mammalia | Physeter macrocephalus (Sperm whale) (Physeter catodon) |
| A0A096MXT2 | KPNA4 | Mammalia | Papio anubis (Olive baboon) |
| F7FL50 | KPNA4 | Mammalia | Monodelphis domestica (Gray short-tailed opossum) |
| M3WRK2 | KPNA4 | Mammalia | Felis catus (Cat) (Felis silvestris catus) |
| A0A2K5DHE6 | KPNA4 | Mammalia | Aotus nancymaae (Ma's night monkey) |
| A0A6J1ZM48 | KPNA4 | Mammalia | Acinonyx jubatus (Cheetah) |
| A0A4W2HRN9 | KPNA4 | Mammalia | Bos indicus x Bos taurus (Hybrid cattle) |
| A0A6I9JJK3 | KPNA4 | Mammalia | Chrysochloris asiatica (Cape golden mole) |
| A0A6J2CG34 | KPNA4 | Mammalia | Zalophus californianus (California sealion) |
| A0A6J2CAN4 | KPNA4 | Mammalia | Zalophus californianus (California sealion) |
| A0A2R8ZEY4 | KPNA4 | Mammalia | Pan paniscus (Pygmy chimpanzee) (Bonobo) |
| A0A6P6BYU9 | KPNA4 | Mammalia | Pteropus vampyrus (Large flying fox) |
| A0A6P3RQQ3 | KPNA4 | Mammalia | Pteropus vampyrus (Large flying fox) |
| A0A340X538 | KPNA4 | Mammalia | Lipotes vexillifer (Yangtze river dolphin) |
| A0A6J3RA32 | KPNA4 | Mammalia | Tursiops truncatus (Atlantic bottle-nosed dolphin) (Delphinus truncatus) |
| A0A3Q7PNZ1 | KPNA4 | Mammalia | Callorhinus ursinus (Northern fur seal) |
| A0A6I9I3N0 | KPNA4 | Mammalia | Vicugna pacos (Alpaca) (Lama pacos) |
| A0A1S2ZT18 | KPNA4 | Mammalia | Erinaceus europaeus (Western European hedgehog) |
| A0A2Y9NIB7 | KPNA4 | Mammalia | Delphinapterus leucas (Beluga whale) |
| A0A6I8PKN1 | KPNA4 | Mammalia | Ornithorhynchus anatinus (Duckbill platypus) |
| A0A3Q7UWJ8 | KPNA4 | Mammalia | Ursus arctos horribilis |
| A0A452SUQ2 | KPNA4 | Mammalia | Ursus americanus (American black bear) (Euarctos americanus) |
| G1SG68 | KPNA4 | Mammalia | Oryctolagus cuniculus (Rabbit) |
| A0A2K6DJI1 | KPNA4 | Mammalia | Macaca nemestrina (Pig-tailed macaque) |
| A0A6P4VF85 | KPNA4 | Mammalia | Panthera pardus (Leopard) (Felis pardus) |
| A0A2K6K7P6 | KPNA4 | Mammalia | Rhinopithecus bieti (Black snub-nosed monkey) (Pygathrix bieti) |
| A0A6P4VIN5 | KPNA4 | Mammalia | Panthera pardus (Leopard) (Felis pardus) |
| A0A6J3GID2 | KPNA4 | Mammalia | Sapajus apella (Brown-capped capuchin) (Cebus apella) |
| G1PMS5 | KPNA4 | Mammalia | Myotis lucifugus (Little brown bat) |
| A0A0B2V5K4 | KPNA5 | Chromadorea | Toxocara canis (Canine roundworm) |
| A0A4Y2LEL5 | KPNA5 | Arachnida | Araneus ventricosus (Orbweaver spider) (Epeira ventricosa) |
| A0A2P8YN66 | KPNA5 | Insecta | Blattella germanica (German cockroach) (Blatta germanica) |
| A0A4W3HMB2 | KPNA5 | Chondrichthyes | Callorhinchus milii (Ghost shark) |
| A0A4W3HM51 | KPNA5 | Chondrichthyes | Callorhinchus milii (Ghost shark) |
| A0A4W3HQ66 | KPNA5 | Chondrichthyes | Callorhinchus milii (Ghost shark) |
| A0A6P7JY08 | KPNA5 | Actinopterygii | Parambassis ranga (Indian glassy fish) |
| A0A1A8AG92 | KPNA5 | Actinopterygii | Nothobranchius furzeri (Turquoise killifish) |
| A0A6P3VTJ8 | KPNA5 | Actinopterygii | Clupea harengus (Atlantic herring) |
| A0A1A8S747 | KPNA5 | Actinopterygii | Nothobranchius rachovii (bluefin notho) |
| A0A4W4FAY0 | KPNA5 | Actinopterygii | Electrophorus electricus (Electric eel) (Gymnotus electricus) |
| A0A4W4FB30 | KPNA5 | Actinopterygii | Electrophorus electricus (Electric eel) (Gymnotus electricus) |
| A0A665TQL2 | KPNA5 | Actinopterygii | Echeneis naucrates (Live sharksucker) |
| A0A665T7A5 | KPNA5 | Actinopterygii | Echeneis naucrates (Live sharksucker) |
| A0A4W4FCA9 | KPNA5 | Actinopterygii | Electrophorus electricus (Electric eel) (Gymnotus electricus) |
| A0A4W4FCE0 | KPNA5 | Actinopterygii | Electrophorus electricus (Electric eel) (Gymnotus electricus) |
| A0A665TF58 | KPNA5 | Actinopterygii | Echeneis naucrates (Live sharksucker) |
| A0A665T652 | KPNA5 | Actinopterygii | Echeneis naucrates (Live sharksucker) |
| A0A1A8CN21 | KPNA5 | Actinopterygii | Nothobranchius kadleci |
| A0A1A8NLB0 | KPNA5 | Actinopterygii | Nothobranchius pienaari |
| A0A0R4I9E5 | KPNA5 | Actinopterygii | Danio rerio (Zebrafish) (Brachydanio rerio) |
| A0A1A8KCW0 | KPNA5 | Actinopterygii | Nothobranchius kuhntae (Beira killifish) |
| A0A1A7XYL4 | KPNA5 | Actinopterygii | Iconisemion striatum |
| A0A6P7KQ24 | KPNA5 | Actinopterygii | Betta splendens (Siamese fighting fish) |
| A0A1A8H282 | KPNA5 | Actinopterygii | Nothobranchius korthausae |
| A0A6P8W9H9 | KPNA5 | Actinopterygii | Gymnodraco acuticeps (Antarctic dragonfish) |
| A0A4W4F9E2 | KPNA5 | Actinopterygii | Electrophorus electricus (Electric eel) (Gymnotus electricus) |
| A0A4W4FB35 | KPNA5 | Actinopterygii | Electrophorus electricus (Electric eel) (Gymnotus electricus) |
| A0A4W4F8U5 | KPNA5 | Actinopterygii | Electrophorus electricus (Electric eel) (Gymnotus electricus) |
| A0A2D0T8X7 | KPNA5 | Actinopterygii | Ictalurus punctatus (Channel catfish) (Silurus punctatus) |
| A0A6J2W6F1 | KPNA5 | Actinopterygii | Chanos chanos (Milkfish) (Mugil chanos) |
| A0A673B4D3 | KPNA5 | Actinopterygii | Sphaeramia orbicularis (orbiculate cardinalfish) |
| A0A673B1Y4 | KPNA5 | Actinopterygii | Sphaeramia orbicularis (orbiculate cardinalfish) |
| W5U8I8 | KPNA5 | Actinopterygii | Ictalurus punctatus (Channel catfish) (Silurus punctatus) |
| H3ASY1 | KPNA5 | Sarcopterygii | Latimeria chalumnae (Coelacanth) |
| A0A1U7RMT4 | KPNA5 | Reptilia | Alligator sinensis (Chinese alligator) |
| A0A674I1L9 | KPNA5 | Reptilia | Terrapene carolina triunguis (Three-toed box turtle) |
| A0A6P9C420 | KPNA5 | Reptilia | Pantherophis guttatus (Corn snake) (Elaphe guttata) |
| A0A663LZJ9 | KPNA5 | Reptilia | Athene cunicularia (Burrowing owl) (Speotyto cunicularia) |
| A0A6J0IEC5 | KPNA5 | Reptilia | Lepidothrix coronata (blue-crowned manakin) |
| A0A151M309 | KPNA5 | Reptilia | Alligator mississippiensis (American alligator) |
| A0A6J1TT36 | KPNA5 | Reptilia | Notechis scutatus (mainland tiger snake) |
| A0A6J2ILX5 | KPNA5 | Reptilia | Pipra filicauda (Wire-tailed manakin) |
| A0A669Q858 | KPNA5 | Reptilia | Phasianus colchicus (Common pheasant) |

| AC | GN | Class | Organism |
|---|---|---|---|
| U3K917 | KPNA5 | Reptilia | Ficedula albicollis (Collared flycatcher) (Muscicapa albicollis) |
| G1KPI9 | KPNA5 | Reptilia | Anolis carolinensis (Green anole) (American chameleon) |
| A0A6J0UAQ1 | KPNA5 | Reptilia | Pogona vitticeps (central bearded dragon) |
| A0A2I0MVK9 | KPNA5 | Reptilia | Columba livia (Rock dove) |
| E1C4J0 | KPNA5 | Reptilia | Gallus gallus (Chicken) |
| A0A6I9X684 | KPNA5 | Reptilia | Thamnophis sirtalis |
| A0A6I9XJ79 | KPNA5 | Reptilia | Thamnophis sirtalis |
| K7FKR6 | KPNA5 | Reptilia | Pelodiscus sinensis (Chinese softshell turtle) (Trionyx sinensis) |
| A0A6I9HDF7 | KPNA5 | Reptilia | Geospiza fortis (Medium ground-finch) |
| U3I6P6 | KPNA5 | Reptilia | Anas platyrhynchos platyrhynchos (Northern mallard) |
| A0A670Y8L6 | KPNA5 | Reptilia | Pseudonaja textilis (Eastern brown snake) |
| A0A218UX91 | KPNA5 | Reptilia | Lonchura striata domestica (Bengalese finch) |
| A0A672TRT0 | KPNA5 | Reptilia | Strigops habroptila (Kakapo) |
| A0A672TRP4 | KPNA5 | Reptilia | Strigops habroptila (Kakapo) |
| A0A1V4JN39 | KPNA5 | Reptilia | Patagioenas fasciata monilis |
| V8NNB7 | KPNA5 | Reptilia | Ophiophagus hannah (King cobra) (Naja hannah) |
| A0A663ECT7 | KPNA5 | Reptilia | Aquila chrysaetos chrysaetos |
| G1NKY4 | KPNA5 | Reptilia | Meleagris gallopavo (Wild turkey) |
| A0A6J3CN91 | KPNA5 | Reptilia | Aythya fuligula (Tufted duck) (Anas fuligula) |
| Q5TD90 | KPNA5 | Mammalia | Homo sapiens (Human) |
| A0A286Y281 | KPNA5 | Mammalia | Cavia porcellus (Guinea pig) |
| A0A6I9M1J8 | KPNA5 | Mammalia | Peromyscus maniculatus bairdii (Prairie deer mouse) |
| A0A0P6JBC4 | KPNA5 | Mammalia | Heterocephalus glaber (Naked mole rat) |
| A0A1U7Q8Z8 | KPNA5 | Mammalia | Mesocricetus auratus (Golden hamster) |
| A0A0H2UHW5 | KPNA5 | Mammalia | Rattus norvegicus (Rat) |
| A0A1S2ZL74 | KPNA5 | Mammalia | Erinaceus europaeus (Western European hedgehog) |
| A0A452G264 | KPNA5 | Mammalia | Capra hircus (Goat) |
| A0A4X1TCK2 | KPNA5 | Mammalia | Sus scrofa (Pig) |
| A0A6J2DZ74 | KPNA5 | Mammalia | Zalophus californianus (California sealion) |
| A0A2I3GNS8 | KPNA5 | Mammalia | Nomascus leucogenys (Northern white-cheeked gibbon) (Hylobates leucogenys) |
| A0A6P5CAC8 | KPNA5 | Mammalia | Bos indicus (Zebu) |
| A0A341CCE6 | KPNA5 | Mammalia | Neophocaena asiaeorientalis asiaeorientalis (Yangtze finless porpoise) (Neophocaena phocaenoides subsp. asiaeorientalis) |
| A0A4X1TD58 | KPNA5 | Mammalia | Sus scrofa (Pig) |
| A0A6J2LE28 | KPNA5 | Mammalia | Phyllostomus discolor (pale spear-nosed bat) |
| A0A6P5J864 | KPNA5 | Mammalia | Phascolarctos cinereus (Koala) |
| A0A6J3HK00 | KPNA5 | Mammalia | Sapajus apella (Brown-capped capuchin) (Cebus apella) |
| G1PF61 | KPNA5 | Mammalia | Myotis lucifugus (Little brown bat) |
| A0A2I2Y6F3 | KPNA5 | Mammalia | Gorilla gorilla gorilla (Western lowland gorilla) |
| A0A2K5TMX7 | KPNA5 | Mammalia | Macaca fascicularis (Crab-eating macaque) (Cynomolgus monkey) |
| A0A5F4WLV0 | KPNA5 | Mammalia | Callithrix jacchus (White-tufted-ear marmoset) |
| F1N1K5 | KPNA5 | Mammalia | Bos taurus (Bovine) |
| A0A5F4W540 | KPNA5 | Mammalia | Callithrix jacchus (White-tufted-ear marmoset) |
| W5PGW2 | KPNA5 | Mammalia | Ovis aries (Sheep) |
| F7A9W1 | KPNA5 | Mammalia | Callithrix jacchus (White-tufted-ear marmoset) |
| A0A1D5QVJ1 | KPNA5 | Mammalia | Macaca mulatta (Rhesus macaque) |
| A0A2K6NC93 | KPNA5 | Mammalia | Rhinopithecus roxellana (Golden snub-nosed monkey) (Pygathrix roxellana) |
| A0A2K5Q2A3 | KPNA5 | Mammalia | Cebus imitator (Panamanian white-faced capuchin) (Cebus capucinus imitator) |
| A0A6P3J9Q1 | KPNA5 | Mammalia | Bison bison bison |
| A0A671FXJ8 | KPNA5 | Mammalia | Rhinolophus ferrumequinum (Greater horseshoe bat) |
| H2QTM3 | KPNA5 | Mammalia | Pan troglodytes (Chimpanzee) |
| A0A6J0Y751 | KPNA5 | Mammalia | Odocoileus virginianus texanus |
| A0A6P3QTU4 | KPNA5 | Mammalia | Pteropus vampyrus (Large flying fox) |
| D2HNW3 | KPNA5 | Mammalia | Ailuropoda melanoleuca (Giant panda) |
| A0A6I9ZEY2 | KPNA5 | Mammalia | Acinonyx jubatus (Cheetah) |
| A0A2K5MAA4 | KPNA5 | Mammalia | Cercocebus atys (Sooty mangabey) (Cercocebus torquatus atys) |
| A0A2K5MAE7 | KPNA5 | Mammalia | Cercocebus atys (Sooty mangabey) (Cercocebus torquatus atys) |
| M3WVA3 | KPNA5 | Mammalia | Felis catus (Cat) (Felis silvestris catus) |
| H0X5T9 | KPNA5 | Mammalia | Otolemur garnettii (Small-eared galago) (Garnett's greater bushbaby) |
| A0A2U3XT32 | KPNA5 | Mammalia | Leptonychotes weddellii (Weddell seal) (Otaria weddellii) |
| A0A455BMW6 | KPNA5 | Mammalia | Physeter macrocephalus (Sperm whale) (Physeter catodon) |
| A0A2Y9HI85 | KPNA5 | Mammalia | Neomonachus schauinslandi (Hawaiian monk seal) (Monachus schauinslandi) |
| H2PK58 | KPNA5 | Mammalia | Pongo abelii (Sumatran orangutan) (Pongo pygmaeus abelii) |
| A0A3Q7TSK3 | KPNA5 | Mammalia | Vulpes vulpes (Red fox) |
| A0A2Y9LPW6 | KPNA5 | Mammalia | Delphinapterus leucas (Beluga whale) |
| H9EPP2 | KPNA5 | Mammalia | Macaca mulatta (Rhesus macaque) |
| A0A4W2FFP7 | KPNA5 | Mammalia | Bos indicus x Bos taurus (Hybrid cattle) |
| A0A384AUJ5 | KPNA5 | Mammalia | Balaenoptera acutorostrata scammoni (North Pacific minke whale) (Balaenoptera davidsoni) |
| A0A2K6A233 | KPNA5 | Mammalia | Mandrillus leucophaeus (Drill) (Papio leucophaeus) |
| A0A0D9RTD2 | KPNA5 | Mammalia | Chlorocebus sabaeus (Green monkey) (Cercopithecus sabaeus) |
| A0A3Q7NQD1 | KPNA5 | Mammalia | Callorhinus ursinus (Northern fur seal) |
| A0A340X6R4 | KPNA5 | Mammalia | Lipotes vexillifer (Yangtze river dolphin) |
| G3ST52 | KPNA5 | Mammalia | Loxodonta africana (African elephant) |
| A0A2I3TFK8 | KPNA5 | Mammalia | Pan troglodytes (Chimpanzee) |
| A0A2K5BXN1 | KPNA5 | Mammalia | Aotus nancymaae (Ma's night monkey) |
| A0A2U4AW54 | KPNA5 | Mammalia | Tursiops truncatus (Atlantic bottle-nosed dolphin) (Delphinus truncatus) |
| F7DMT7 | KPNA5 | Mammalia | Equus caballus (Horse) |
| A0A6I9JBC6 | KPNA5 | Mammalia | Chrysochloris asiatica (Cape golden mole) |
| A0A673UVH5 | KPNA5 | Mammalia | Suricata suricatta (Meerkat) |
| A0A2R9AVQ0 | KPNA5 | Mammalia | Pan paniscus (Pygmy chimpanzee) (Bonobo) |
| A0A096NFQ3 | KPNA5 | Mammalia | Papio anubis (Olive baboon) |
| A0A3Q7NFQ8 | KPNA5 | Mammalia | Callorhinus ursinus (Northern fur seal) |
| F1PQD8 | KPNA5 | Mammalia | Canis lupus familiaris (Dog) (Canis familiaris) |
| A0A2K6FAD0 | KPNA5 | Mammalia | Propithecus coquereli (Coquerel's sifaka) (Propithecus verreauxi coquereli) |

| AC | GN | Class | Organism |
|---|---|---|---|
| A0A2K6FAB4 | KPNA5 | Mammalia | Propithecus coquereli (Coquerel's sifaka) (Propithecus verreauxi coquereli) |
| A0A2K6KIL0 | KPNA5 | Mammalia | Rhinopithecus bieti (Black snub-nosed monkey) (Pygathrix bieti) |
| A0A452RB17 | KPNA5 | Mammalia | Ursus americanus (American black bear) (Euarctos americanus) |
| A0A2U3VS73 | KPNA5 | Mammalia | Odobenus rosmarus divergens (Pacific walrus) |
| M3YS23 | KPNA5 | Mammalia | Mustela putorius furo (European domestic ferret) (Mustela furo) |
| A0A6I9IAP3 | KPNA5 | Mammalia | Vicugna pacos (Alpaca) (Lama pacos) |
| A0A1U7UBB2 | KPNA5 | Mammalia | Carlito syrichta (Philippine tarsier) (Tarsius syrichta) |
| A0A2K6SJ59 | KPNA5 | Mammalia | Saimiri boliviensis boliviensis (Bolivian squirrel monkey) |
| A0A452RB58 | KPNA5 | Mammalia | Ursus americanus (American black bear) (Euarctos americanus) |
| A0A2K6FAA6 | KPNA5 | Mammalia | Propithecus coquereli (Coquerel's sifaka) (Propithecus verreauxi coquereli) |
| A0A2K6FAC6 | KPNA5 | Mammalia | Propithecus coquereli (Coquerel's sifaka) (Propithecus verreauxi coquereli) |
| A0A4X2M1C7 | KPNA5 | Mammalia | Vombatus ursinus (Common wombat) |
| A0A2K5HJB4 | KPNA5 | Mammalia | Colobus angolensis palliatus (Peters' Angolan colobus) |
| G3R8B6 | KPNA5 | Mammalia | Gorilla gorilla gorilla (Western lowland gorilla) |
| A0A6P6HZV5 | KPNA5 | Mammalia | Puma concolor (Mountain lion) |
| A0A2K6C6R2 | KPNA5 | Mammalia | Macaca nemestrina (Pig-tailed macaque) |
| A0A384C3E9 | KPNA5 | Mammalia | Ursus maritimus (Polar bear) (Thalarctos maritimus) |
| G1STV1 | KPNA5 | Mammalia | Oryctolagus cuniculus (Rabbit) |
| A0A667FL52 | KPNA5 | Mammalia | Lynx canadensis (Canada lynx) |
| A0A6P4T529 | KPNA5 | Mammalia | Panthera pardus (Leopard) (Felis pardus) |
| A0A452TA68 | KPNA5 | Mammalia | Ursus maritimus (Polar bear) (Thalarctos maritimus) |
| A0A5B6UCM2 | KPNA5 | Magnoliopsida | Gossypium australe |
| A0A5B6VL98 | KPNA5 | Magnoliopsida | Gossypium australe |
| A0A2B4SYN2 | KPNA6 | Anthozoa | Stylophora pistillata (Smooth cauliflower coral) |
| T2MID0 | KPNA6 | Hydrozoa | Hydra vulgaris (Hydra) (Hydra attenuata) |
| A0A6G1S6W7 | KPNA6 | Arachnida | Aceria tosichella (wheat curl mite) |
| A0A4C1U0C7 | KPNA6 | Insecta | Eumeta variegata (Bagworm moth) (Eumeta japonica) |
| A0A0A1XSA1 | KPNA6 | Insecta | Zeugodacus cucurbitae (Melon fruit fly) (Bactrocera cucurbitae) |
| A0A2S2R7M9 | KPNA6 | Insecta | Sipha flava (yellow sugarcane aphid) |
| A0A0K8VAU4 | KPNA6 | Insecta | Bactrocera latifrons (Malaysian fruit fly) (Chaetodacus latifrons) |
| A0A2H8U0D0 | KPNA6 | Insecta | Melanaphis sacchari |
| A0A0A9XP06 | KPNA6 | Insecta | Lygus hesperus (Western plant bug) |
| A0A0C9QJY4 | KPNA6 | Insecta | Fopius arisanus |
| A0A6A4WGB8 | KPNA6 | Hexanauplia | Amphibalanus amphitrite (Striped barnacle) (Balanus amphitrite) |
| A0A6F9DGL9 | KPNA6 | Ascidiacea | Phallusia mammillata |
| A0A4W3ITE9 | KPNA6 | Chondrichthyes | Callorhinchus milii (Ghost shark) |
| A0A4W3ITH2 | KPNA6 | Chondrichthyes | Callorhinchus milii (Ghost shark) |
| A0A6P7NWW0 | KPNA6 | Actinopterygii | Betta splendens (Siamese fighting fish) |
| A0A6P7NY74 | KPNA6 | Actinopterygii | Betta splendens (Siamese fighting fish) |
| A0A1A7WS65 | KPNA6 | Actinopterygii | Iconisemion striatum |
| A0A6J2QLQ7 | KPNA6 | Actinopterygii | Cottoperca gobio (Frogmouth) (Aphritis gobio) |
| A0A6P3VJS0 | KPNA6 | Actinopterygii | Clupea harengus (Atlantic herring) |
| A0A665UTB0 | KPNA6 | Actinopterygii | Echeneis naucrates (Live sharksucker) |
| A0A665USW8 | KPNA6 | Actinopterygii | Echeneis naucrates (Live sharksucker) |
| A0A665UT43 | KPNA6 | Actinopterygii | Echeneis naucrates (Live sharksucker) |
| A0A674N9Z8 | KPNA6 | Actinopterygii | Takifugu rubripes (Japanese pufferfish) (Fugu rubripes) |
| A0A674MZC1 | KPNA6 | Actinopterygii | Takifugu rubripes (Japanese pufferfish) (Fugu rubripes) |
| A0A665URM1 | KPNA6 | Actinopterygii | Echeneis naucrates (Live sharksucker) |
| A0A674P4V4 | KPNA6 | Actinopterygii | Takifugu rubripes (Japanese pufferfish) (Fugu rubripes) |
| A0A665USJ2 | KPNA6 | Actinopterygii | Echeneis naucrates (Live sharksucker) |
| A0A674PHN0 | KPNA6 | Actinopterygii | Takifugu rubripes (Japanese pufferfish) (Fugu rubripes) |
| A0A674N3Q7 | KPNA6 | Actinopterygii | Takifugu rubripes (Japanese pufferfish) (Fugu rubripes) |
| A0A674PL91 | KPNA6 | Actinopterygii | Takifugu rubripes (Japanese pufferfish) (Fugu rubripes) |
| A0A665USW9 | KPNA6 | Actinopterygii | Echeneis naucrates (Live sharksucker) |
| A0A665USF0 | KPNA6 | Actinopterygii | Echeneis naucrates (Live sharksucker) |
| A0A674NDE2 | KPNA6 | Actinopterygii | Takifugu rubripes (Japanese pufferfish) (Fugu rubripes) |
| H2TLL4 | KPNA6 | Actinopterygii | Takifugu rubripes (Japanese pufferfish) (Fugu rubripes) |
| A0A669E488 | KPNA6 | Actinopterygii | Oreochromis niloticus (Nile tilapia) (Tilapia nilotica) |
| A0A669F393 | KPNA6 | Actinopterygii | Oreochromis niloticus (Nile tilapia) (Tilapia nilotica) |
| A0A669CXP1 | KPNA6 | Actinopterygii | Oreochromis niloticus (Nile tilapia) (Tilapia nilotica) |
| A0A669C9I5 | KPNA6 | Actinopterygii | Oreochromis niloticus (Nile tilapia) (Tilapia nilotica) |
| A0A669ENE7 | KPNA6 | Actinopterygii | Oreochromis niloticus (Nile tilapia) (Tilapia nilotica) |
| A0A669C1X0 | KPNA6 | Actinopterygii | Oreochromis niloticus (Nile tilapia) (Tilapia nilotica) |
| E7F7U9 | KPNA6 | Actinopterygii | Danio rerio (Zebrafish) (Brachydanio rerio) |
| H2MGF0 | KPNA6 | Actinopterygii | Oryzias latipes (Japanese rice fish) (Japanese killifish) |
| A0A669BJX6 | KPNA6 | Actinopterygii | Oreochromis niloticus (Nile tilapia) (Tilapia nilotica) |
| I3JLH4 | KPNA6 | Actinopterygii | Oreochromis niloticus (Nile tilapia) (Tilapia nilotica) |
| A0A1A8RFX3 | KPNA6 | Actinopterygii | Nothobranchius rachovii (bluefin notho) |
| A0A1A8KQY4 | KPNA6 | Actinopterygii | Nothobranchius kuhntae (Beira killifish) |
| A0A1A8MC56 | KPNA6 | Actinopterygii | Nothobranchius pienaari |
| A0A1A8AWK2 | KPNA6 | Actinopterygii | Nothobranchius furzeri (Turquoise killifish) |
| A0A1A8D3J2 | KPNA6 | Actinopterygii | Nothobranchius kadleci |
| A0A673AFG9 | KPNA6 | Actinopterygii | Sphaeramia orbicularis (orbiculate cardinalfish) |
| A0A673AGP4 | KPNA6 | Actinopterygii | Sphaeramia orbicularis (orbiculate cardinalfish) |
| A0A667ZAA2 | KPNA6 | Actinopterygii | Myripristis murdjan (pinecone soldierfish) |
| A0A667Z1Z9 | KPNA6 | Actinopterygii | Myripristis murdjan (pinecone soldierfish) |
| A0A667Z113 | KPNA6 | Actinopterygii | Myripristis murdjan (pinecone soldierfish) |
| A0A667YPD2 | KPNA6 | Actinopterygii | Myripristis murdjan (pinecone soldierfish) |
| A0A2I4CRW0 | KPNA6 | Actinopterygii | Austrofundulus limnaeus |
| A0A2D0S2W3 | KPNA6 | Actinopterygii | Ictalurus punctatus (Channel catfish) (Silurus punctatus) |
| A0A667ZA63 | KPNA6 | Actinopterygii | Myripristis murdjan (pinecone soldierfish) |
| A0A667Z1G5 | KPNA6 | Actinopterygii | Myripristis murdjan (pinecone soldierfish) |
| A0A673AGD2 | KPNA6 | Actinopterygii | Sphaeramia orbicularis (orbiculate cardinalfish) |

| AC | GN | Class | Organism |
|---|---|---|---|
| A0A673AGC1 | KPNA6 | Actinopterygii | Sphaeramia orbicularis (orbiculate cardinalfish) |
| A0A673AJ84 | KPNA6 | Actinopterygii | Sphaeramia orbicularis (orbiculate cardinalfish) |
| A0A673AJD9 | KPNA6 | Actinopterygii | Sphaeramia orbicularis (orbiculate cardinalfish) |
| A0A6J2WLL7 | KPNA6 | Actinopterygii | Chanos chanos (Milkfish) (Mugil chanos) |
| A0A673AID3 | KPNA6 | Actinopterygii | Sphaeramia orbicularis (orbiculate cardinalfish) |
| A0A673AFB1 | KPNA6 | Actinopterygii | Sphaeramia orbicularis (orbiculate cardinalfish) |
| A0A4Z2H8L4 | KPNA6 | Actinopterygii | Liparis tanakae (Tanaka's snailfish) |
| A0A667YPY9 | KPNA6 | Actinopterygii | Myripristis murdjan (pinecone soldierfish) |
| A0A667YHP8 | KPNA6 | Actinopterygii | Myripristis murdjan (pinecone soldierfish) |
| A0A674PD67 | KPNA6 | Actinopterygii | Takifugu rubripes (Japanese pufferfish) (Fugu rubripes) |
| A0A6P7JZ47 | KPNA6 | Actinopterygii | Parambassis ranga (Indian glassy fish) |
| A0A6P7JV58 | KPNA6 | Actinopterygii | Parambassis ranga (Indian glassy fish) |
| A0A1A8FMT4 | KPNA6 | Actinopterygii | Nothobranchius korthausae |
| A0A3B3IGL5 | KPNA6 | Actinopterygii | Oryzias latipes (Japanese rice fish) (Japanese killifish) |
| Q6IP69 | KPNA6 | Amphibia | Xenopus laevis (African clawed frog) |
| A0A6I8SII8 | KPNA6 | Amphibia | Xenopus tropicalis (Western clawed frog) (Silurana tropicalis) |
| A0A6I8SHV4 | KPNA6 | Amphibia | Xenopus tropicalis (Western clawed frog) (Silurana tropicalis) |
| A0A6I8T171 | KPNA6 | Amphibia | Xenopus tropicalis (Western clawed frog) (Silurana tropicalis) |
| Q70PC4 | KPNA6 | Amphibia | Xenopus laevis (African clawed frog) |
| A0A6P8S1I7 | KPNA6 | Amphibia | Geotrypetes seraphini (Gaboon caecilian) (Caecilia seraphini) |
| A0A6P7ZKX7 | KPNA6 | Amphibia | Microcaecilia unicolor |
| H0YT19 | KPNA6 | Reptilia | Taeniopygia guttata (Zebra finch) (Poephila guttata) |
| A0A6P9BDZ8 | KPNA6 | Reptilia | Pantherophis guttatus (Corn snake) (Elaphe guttata) |
| A0A663MVT2 | KPNA6 | Reptilia | Athene cunicularia (Burrowing owl) (Speotyto cunicularia) |
| A0A6J0IB21 | KPNA6 | Reptilia | Lepidothrix coronata (blue-crowned manakin) |
| A0A669Q4K7 | KPNA6 | Reptilia | Phasianus colchicus (Common pheasant) |
| A0A674I4Z4 | KPNA6 | Reptilia | Terrapene carolina triunguis (Three-toed box turtle) |
| A0A452IZ11 | KPNA6 | Reptilia | Gopherus agassizii (Agassiz's desert tortoise) |
| A0A3Q0G435 | KPNA6 | Reptilia | Alligator sinensis (Chinese alligator) |
| A0A669PNE7 | KPNA6 | Reptilia | Phasianus colchicus (Common pheasant) |
| H9G9G7 | KPNA6 | Reptilia | Anolis carolinensis (Green anole) (American chameleon) |
| F7B5S0 | KPNA6 | Reptilia | Gallus gallus (Chicken) |
| A0A6J0U645 | KPNA6 | Reptilia | Pogona vitticeps (central bearded dragon) |
| A0A6J0U8D3 | KPNA6 | Reptilia | Pogona vitticeps (central bearded dragon) |
| A0A2I0LNF2 | KPNA6 | Reptilia | Columba livia (Rock dove) |
| A0A6J0IB16 | KPNA6 | Reptilia | Lepidothrix coronata (blue-crowned manakin) |
| A0A6I9Y5G6 | KPNA6 | Reptilia | Thamnophis sirtalis |
| K7FQU4 | KPNA6 | Reptilia | Pelodiscus sinensis (Chinese softshell turtle) (Trionyx sinensis) |
| A0A218UGK2 | KPNA6 | Reptilia | Lonchura striata domestica (Bengalese finch) |
| A0A6J2H9B6 | KPNA6 | Reptilia | Pipra filicauda (Wire-tailed manakin) |
| A0A6J1UEW1 | KPNA6 | Reptilia | Notechis scutatus (mainland tiger snake) |
| U3IY73 | KPNA6 | Reptilia | Anas platyrhynchos platyrhynchos (Northern mallard) |
| K7FQT8 | KPNA6 | Reptilia | Pelodiscus sinensis (Chinese softshell turtle) (Trionyx sinensis) |
| A0A670Y1W5 | KPNA6 | Reptilia | Pseudonaja textilis (Eastern brown snake) |
| A0A670Y8U2 | KPNA6 | Reptilia | Pseudonaja textilis (Eastern brown snake) |
| V8NCM0 | KPNA6 | Reptilia | Ophiophagus hannah (King cobra) (Naja hannah) |
| A0A1V4JIW7 | KPNA6 | Reptilia | Patagioenas fasciata monilis |
| A0A6J2H8S0 | KPNA6 | Reptilia | Pipra filicauda (Wire-tailed manakin) |
| A0A670JX51 | KPNA6 | Reptilia | Podarcis muralis (Wall lizard) (Lacerta muralis) |
| A0A663EJC7 | KPNA6 | Reptilia | Aquila chrysaetos chrysaetos |
| G1N050 | KPNA6 | Reptilia | Meleagris gallopavo (Wild turkey) |
| A0A6J3E8N2 | KPNA6 | Reptilia | Aythya fuligula (Tufted duck) (Anas fuligula) |
| A0A672V4I2 | KPNA6 | Reptilia | Strigops habroptila (Kakapo) |
| A0A663EJD2 | KPNA6 | Reptilia | Aquila chrysaetos chrysaetos |
| O60684 | KPNA6 | Mammalia | Homo sapiens (Human) |
| Q3KR98 | KPNA6 | Mammalia | Rattus norvegicus (Rat) |
| A0A0G2JZS1 | KPNA6 | Mammalia | Rattus norvegicus (Rat) |
| A0A250YJI9 | KPNA6 | Mammalia | Castor canadensis (American beaver) |
| A0A6I9M8L4 | KPNA6 | Mammalia | Peromyscus maniculatus bairdii (Prairie deer mouse) |
| A0A286X7C6 | KPNA6 | Mammalia | Cavia porcellus (Guinea pig) |
| A0A3Q0D5Z9 | KPNA6 | Mammalia | Mesocricetus auratus (Golden hamster) |
| I3MIF0 | KPNA6 | Mammalia | Ictidomys tridecemlineatus (Thirteen-lined ground squirrel) (Spermophilus tridecemlineatus) |
| Q8BH30 | KPNA6 | Mammalia | Mus musculus (Mouse) |
| A0A6J0CKI2 | KPNA6 | Mammalia | Peromyscus maniculatus bairdii (Prairie deer mouse) |
| A0A6P5PM01 | KPNA6 | Mammalia | Mus caroli (Ryukyu mouse) (Ricefield mouse) |
| A0A6P6DFA1 | KPNA6 | Mammalia | Octodon degus (Degu) (Sciurus degus) |
| A0A6P5PQG2 | KPNA6 | Mammalia | Mus caroli (Ryukyu mouse) (Ricefield mouse) |
| F1LT58 | KPNA6 | Mammalia | Rattus norvegicus (Rat) |
| A0A1S3FHK2 | KPNA6 | Mammalia | Dipodomys ordii (Ord's kangaroo rat) |
| F6SAJ5 | KPNA6 | Mammalia | Ornithorhynchus anatinus (Duckbill platypus) |
| A0A3Q0EAN1 | KPNA6 | Mammalia | Carlito syrichta (Philippine tarsier) (Tarsius syrichta) |
| A0A2Y9NNA2 | KPNA6 | Mammalia | Delphinapterus leucas (Beluga whale) |
| A0A2K5K3Y4 | KPNA6 | Mammalia | Colobus angolensis palliatus (Peters' Angolan colobus) |
| A0A2K5K3W0 | KPNA6 | Mammalia | Colobus angolensis palliatus (Peters' Angolan colobus) |
| A0A1S3AP65 | KPNA6 | Mammalia | Erinaceus europaeus (Western European hedgehog) |
| A0A6I9HXL5 | KPNA6 | Mammalia | Vicugna pacos (Alpaca) (Lama pacos) |
| A0A673TMV9 | KPNA6 | Mammalia | Suricata suricatta (Meerkat) |
| A0A341D6G7 | KPNA6 | Mammalia | Neophocaena asiaeorientalis asiaeorientalis (Yangtze finless porpoise) (Neophocaena phocaenoides subsp. asiaeorientalis) |
| A0A0D9S7X9 | KPNA6 | Mammalia | Chlorocebus sabaeus (Green monkey) (Cercopithecus sabaeus) |
| A0A2K6AEJ5 | KPNA6 | Mammalia | Mandrillus leucophaeus (Drill) (Papio leucophaeus) |
| A0A4X1W0L9 | KPNA6 | Mammalia | Sus scrofa (Pig) |
| A0A6J2DI83 | KPNA6 | Mammalia | Zalophus californianus (California sealion) |
| M3Z9Q7 | KPNA6 | Mammalia | Nomascus leucogenys (Northern white-cheeked gibbon) (Hylobates leucogenys) |

| AC | GN | Class | Organism |
|---|---|---|---|
| A0A673U0E9 | KPNA6 | Mammalia | Suricata suricatta (Meerkat) |
| A0A2K6AEI0 | KPNA6 | Mammalia | Mandrillus leucophaeus (Drill) (Papio leucophaeus) |
| F6ZAL1 | KPNA6 | Mammalia | Equus caballus (Horse) |
| A0A341D6Y7 | KPNA6 | Mammalia | Neophocaena asiaeorientalis asiaeorientalis (Yangtze finless porpoise) (Neophocaena phocaenoides subsp. asiaeorientalis) |
| A0A6P5DAY8 | KPNA6 | Mammalia | Bos indicus (Zebu) |
| A0A2I2YR04 | KPNA6 | Mammalia | Gorilla gorilla gorilla (Western lowland gorilla) |
| F1PM77 | KPNA6 | Mammalia | Canis lupus familiaris (Dog) (Canis familiaris) |
| A0A2K5X9F2 | KPNA6 | Mammalia | Macaca fascicularis (Crab-eating macaque) (Cynomolgus monkey) |
| A0A6P5K253 | KPNA6 | Mammalia | Phascolarctos cinereus (Koala) |
| A0A6J3FAF5 | KPNA6 | Mammalia | Sapajus apella (Brown-capped capuchin) (Cebus apella) |
| G1TE61 | KPNA6 | Mammalia | Oryctolagus cuniculus (Rabbit) |
| A0A6P6IQI4 | KPNA6 | Mammalia | Puma concolor (Mountain lion) |
| A0A2K5X8Z3 | KPNA6 | Mammalia | Macaca fascicularis (Crab-eating macaque) (Cynomolgus monkey) |
| A0A667GXC7 | KPNA6 | Mammalia | Lynx canadensis (Canada lynx) |
| A0A6P5K276 | KPNA6 | Mammalia | Phascolarctos cinereus (Koala) |
| G1PWY7 | KPNA6 | Mammalia | Myotis lucifugus (Little brown bat) |
| A0A6G1AYV8 | KPNA6 | Mammalia | Crocuta crocuta (Spotted hyena) |
| A0A2K5X913 | KPNA6 | Mammalia | Macaca fascicularis (Crab-eating macaque) (Cynomolgus monkey) |
| G1LMT1 | KPNA6 | Mammalia | Ailuropoda melanoleuca (Giant panda) |
| A0A6J2LMK8 | KPNA6 | Mammalia | Phyllostomus discolor (pale spear-nosed bat) |
| A0A2K5RW11 | KPNA6 | Mammalia | Cebus imitator (Panamanian white-faced capuchin) (Cebus capucinus imitator) |
| A0A6J0A0G5 | KPNA6 | Mammalia | Acinonyx jubatus (Cheetah) |
| A0A6P3H7K5 | KPNA6 | Mammalia | Bison bison bison |
| A0A6P3T9P0 | KPNA6 | Mammalia | Ovis aries (Sheep) |
| A0A671F6J5 | KPNA6 | Mammalia | Rhinolophus ferrumequinum (Greater horseshoe bat) |
| W5NR48 | KPNA6 | Mammalia | Ovis aries (Sheep) |
| F7FPC0 | KPNA6 | Mammalia | Callithrix jacchus (White-tufted-ear marmoset) |
| A0A1D5QJS1 | KPNA6 | Mammalia | Macaca mulatta (Rhesus macaque) |
| A0A2K5RVX9 | KPNA6 | Mammalia | Cebus imitator (Panamanian white-faced capuchin) (Cebus capucinus imitator) |
| A0A6J0XQK1 | KPNA6 | Mammalia | Odocoileus virginianus texanus |
| A0A3Q7UAJ7 | KPNA6 | Mammalia | Vulpes vulpes (Red fox) |
| A0A6J0XSR8 | KPNA6 | Mammalia | Odocoileus virginianus texanus |
| A0A384CCX3 | KPNA6 | Mammalia | Ursus maritimus (Polar bear) (Thalarctos maritimus) |
| H9EN17 | KPNA6 | Mammalia | Macaca mulatta (Rhesus macaque) |
| A0A6J2DEH4 | KPNA6 | Mammalia | Zalophus californianus (California sealion) |
| A0A6J0A060 | KPNA6 | Mammalia | Acinonyx jubatus (Cheetah) |
| A0A2I3SZD9 | KPNA6 | Mammalia | Pan troglodytes (Chimpanzee) |
| A0A2K5MF98 | KPNA6 | Mammalia | Cercocebus atys (Sooty mangabey) (Cercocebus torquatus atys) |
| A0A2K5RW26 | KPNA6 | Mammalia | Cebus imitator (Panamanian white-faced capuchin) (Cebus capucinus imitator) |
| A0A096N2D7 | KPNA6 | Mammalia | Papio anubis (Olive baboon) |
| A0A7F8QX02 | KPNA6 | Mammalia | Leptonychotes weddellii (Weddell seal) (Otaria weddellii) |
| A0A2I2V555 | KPNA6 | Mammalia | Felis catus (Cat) (Felis silvestris catus) |
| F7E1F4 | KPNA6 | Mammalia | Monodelphis domestica (Gray short-tailed opossum) |
| A0A2I3LJG3 | KPNA6 | Mammalia | Papio anubis (Olive baboon) |
| A0A2Y9GL70 | KPNA6 | Mammalia | Neomonachus schauinslandi (Hawaiian monk seal) (Monachus schauinslandi) |
| A0A2R8ZNF5 | KPNA6 | Mammalia | Pan paniscus (Pygmy chimpanzee) (Bonobo) |
| A0A2Y9FJU4 | KPNA6 | Mammalia | Physeter macrocephalus (Sperm whale) (Physeter catodon) |
| A0A2I3LV73 | KPNA6 | Mammalia | Papio anubis (Olive baboon) |
| A0A2I3MMA4 | KPNA6 | Mammalia | Papio anubis (Olive baboon) |
| A0A2K5MFB8 | KPNA6 | Mammalia | Cercocebus atys (Sooty mangabey) (Cercocebus torquatus atys) |
| H0X0L1 | KPNA6 | Mammalia | Otolemur garnettii (Small-eared galago) (Garnett's greater bushbaby) |
| A0A455BFC9 | KPNA6 | Mammalia | Physeter macrocephalus (Sperm whale) (Physeter catodon) |
| A0A5F8GFC7 | KPNA6 | Mammalia | Monodelphis domestica (Gray short-tailed opossum) |
| A0A340Y669 | KPNA6 | Mammalia | Lipotes vexillifer (Yangtze river dolphin) |
| A0A340YBD4 | KPNA6 | Mammalia | Lipotes vexillifer (Yangtze river dolphin) |
| A0A2I3RHW2 | KPNA6 | Mammalia | Pan troglodytes (Chimpanzee) |
| H2N861 | KPNA6 | Mammalia | Pongo abelii (Sumatran orangutan) (Pongo pygmaeus abelii) |
| F7FPD9 | KPNA6 | Mammalia | Callithrix jacchus (White-tufted-ear marmoset) |
| A0A4W2FQX2 | KPNA6 | Mammalia | Bos indicus x Bos taurus (Hybrid cattle) |
| H2PYJ7 | KPNA6 | Mammalia | Pan troglodytes (Chimpanzee) |
| A0A287AEI6 | KPNA6 | Mammalia | Sus scrofa (Pig) |
| A0A6I9JKJ0 | KPNA6 | Mammalia | Chrysochloris asiatica (Cape golden mole) |
| A0A2K6AEI5 | KPNA6 | Mammalia | Mandrillus leucophaeus (Drill) (Papio leucophaeus) |
| A0A2K6AEF3 | KPNA6 | Mammalia | Mandrillus leucophaeus (Drill) (Papio leucophaeus) |
| A0A6J2DCS8 | KPNA6 | Mammalia | Zalophus californianus (California sealion) |
| A0A452ETW3 | KPNA6 | Mammalia | Capra hircus (Goat) |
| A0A6J2DI93 | KPNA6 | Mammalia | Zalophus californianus (California sealion) |
| G3VZX7 | KPNA6 | Mammalia | Sarcophilus harrisii (Tasmanian devil) (Sarcophilus laniarius) |
| A0A3Q7MUA7 | KPNA6 | Mammalia | Callorhinus ursinus (Northern fur seal) |
| A0A2U4BKQ3 | KPNA6 | Mammalia | Tursiops truncatus (Atlantic bottle-nosed dolphin) (Delphinus truncatus) |
| G3TI14 | KPNA6 | Mammalia | Loxodonta africana (African elephant) |
| A0A5G2QUV4 | KPNA6 | Mammalia | Sus scrofa (Pig) |
| A0A6J2DI60 | KPNA6 | Mammalia | Zalophus californianus (California sealion) |
| A0A6P6CCM1 | KPNA6 | Mammalia | Pteropus vampyrus (Large flying fox) |
| A0A2R8ZHT3 | KPNA6 | Mammalia | Pan paniscus (Pygmy chimpanzee) (Bonobo) |
| A0A340Y4V3 | KPNA6 | Mammalia | Lipotes vexillifer (Yangtze river dolphin) |
| A0A6P3Q105 | KPNA6 | Mammalia | Pteropus vampyrus (Large flying fox) |
| A0A3Q7P798 | KPNA6 | Mammalia | Callorhinus ursinus (Northern fur seal) |
| A0A6P6CCF2 | KPNA6 | Mammalia | Pteropus vampyrus (Large flying fox) |
| A0A3Q7MWD7 | KPNA6 | Mammalia | Callorhinus ursinus (Northern fur seal) |
| A0A2U4BKP8 | KPNA6 | Mammalia | Tursiops truncatus (Atlantic bottle-nosed dolphin) (Delphinus truncatus) |
| A0A3Q7X5I0 | KPNA6 | Mammalia | Ursus arctos horribilis |
| A0A2K6TJD1 | KPNA6 | Mammalia | Saimiri boliviensis boliviensis (Bolivian squirrel monkey) |

| AC | GN | Class | Organism |
|---|---|---|---|
| A0A1U7TY64 | KPNA6 | Mammalia | Carlito syrichta (Philippine tarsier) (Tarsius syrichta) |
| A0A1S3APL8 | KPNA6 | Mammalia | Erinaceus europaeus (Western European hedgehog) |
| A0A2K5K3W8 | KPNA6 | Mammalia | Colobus angolensis palliatus (Peters' Angolan colobus) |
| A0A2Y9NH87 | KPNA6 | Mammalia | Delphinapterus leucas (Beluga whale) |
| A0A2U3ZLS2 | KPNA6 | Mammalia | Odobenus rosmarus divergens (Pacific walrus) |
| A0A3Q0EEC2 | KPNA6 | Mammalia | Carlito syrichta (Philippine tarsier) (Tarsius syrichta) |
| A0A4X2JN23 | KPNA6 | Mammalia | Vombatus ursinus (Common wombat) |
| A0A2K6LP75 | KPNA6 | Mammalia | Rhinopithecus bieti (Black snub-nosed monkey) (Pygathrix bieti) |
| A0A452RN34 | KPNA6 | Mammalia | Ursus americanus (American black bear) (Euarctos americanus) |
| A0A2K6EGT9 | KPNA6 | Mammalia | Propithecus coquereli (Coquerel's sifaka) (Propithecus verreauxi coquereli) |
| A0A452RMT7 | KPNA6 | Mammalia | Ursus americanus (American black bear) (Euarctos americanus) |
| A0A2K6TJE2 | KPNA6 | Mammalia | Saimiri boliviensis boliviensis (Bolivian squirrel monkey) |
| A0A383Z6H8 | KPNA6 | Mammalia | Balaenoptera acutorostrata scammoni (North Pacific minke whale) (Balaenoptera davidsoni) |
| A0A4X2JNT1 | KPNA6 | Mammalia | Vombatus ursinus (Common wombat) |
| A0A383Z655 | KPNA6 | Mammalia | Balaenoptera acutorostrata scammoni (North Pacific minke whale) (Balaenoptera davidsoni) |
| A0A2K6EGU4 | KPNA6 | Mammalia | Propithecus coquereli (Coquerel's sifaka) (Propithecus verreauxi coquereli) |
| A0A6P4UL29 | KPNA6 | Mammalia | Panthera pardus (Leopard) (Felis pardus) |
| A0A2K6E6T2 | KPNA6 | Mammalia | Macaca nemestrina (Pig-tailed macaque) |
| A0A452RMZ1 | KPNA6 | Mammalia | Ursus americanus (American black bear) (Euarctos americanus) |
| A0A452RMQ4 | KPNA6 | Mammalia | Ursus americanus (American black bear) (Euarctos americanus) |
| A0A6P4UDT9 | KPNA6 | Mammalia | Panthera pardus (Leopard) (Felis pardus) |
| A0A286XBY1 | KPNA6 | Mammalia | Cavia porcellus (Guinea pig) |
| Q0V7M0 | KPNA6 | Mammalia | Bos taurus (Bovine) |
| A0A4W3JZ18 | KPNA7 | Chondrichthyes | Callorhinchus milii (Ghost shark) |
| Q803D8 | KPNA7 | Actinopterygii | Danio rerio (Zebrafish) (Brachydanio rerio) |
| A0A060YJS5 | KPNA7 | Actinopterygii | Oncorhynchus mykiss (Rainbow trout) (Salmo gairdneri) |
| A0A6P8VU10 | KPNA7 | Actinopterygii | Gymnodraco acuticeps (Antarctic dragonfish) |
| Q4QR45 | KPNA7 | Amphibia | Xenopus laevis (African clawed frog) |
| A0A1L8EXD5 | KPNA7 | Amphibia | Xenopus laevis (African clawed frog) |
| A0A6P7YUC3 | KPNA7 | Amphibia | Microcaecilia unicolor |
| B4F6X3 | KPNA7 | Amphibia | Xenopus tropicalis (Western clawed frog) (Silurana tropicalis) |
| A0A6P8SI26 | KPNA7 | Amphibia | Geotrypetes seraphini (Gaboon caecilian) (Caecilia seraphini) |
| A0A674JCE7 | KPNA7 | Reptilia | Terrapene carolina triunguis (Three-toed box turtle) |
| A0A674J8N5 | KPNA7 | Reptilia | Terrapene carolina triunguis (Three-toed box turtle) |
| A0A6P9DMK1 | KPNA7 | Reptilia | Pantherophis guttatus (Corn snake) (Elaphe guttata) |
| A0A6J1VLK5 | KPNA7 | Reptilia | Notechis scutatus (mainland tiger snake) |
| A0A151N2K4 | KPNA7 | Reptilia | Alligator mississippiensis (American alligator) |
| A0A663NCZ5 | KPNA7 | Reptilia | Athene cunicularia (Burrowing owl) (Speotyto cunicularia) |
| A0A6J2IHK9 | KPNA7 | Reptilia | Pipra filicauda (Wire-tailed manakin) |
| A0A669Q915 | KPNA7 | Reptilia | Phasianus colchicus (Common pheasant) |
| H0ZB96 | KPNA7 | Reptilia | Taeniopygia guttata (Zebra finch) (Poephila guttata) |
| A0A452IQU3 | KPNA7 | Reptilia | Gopherus agassizii (Agassiz's desert tortoise) |
| U3KHB8 | KPNA7 | Reptilia | Ficedula albicollis (Collared flycatcher) (Muscicapa albicollis) |
| E1BVP7 | KPNA7 | Reptilia | Gallus gallus (Chicken) |
| A0A6J0V1C3 | KPNA7 | Reptilia | Pogona vitticeps (central bearded dragon) |
| A0A2I0M0S3 | KPNA7 | Reptilia | Columba livia (Rock dove) |
| A0A3Q3B277 | KPNA7 | Reptilia | Gallus gallus (Chicken) |
| A0A670YIB3 | KPNA7 | Reptilia | Pseudonaja textilis (Eastern brown snake) |
| A0A6I9HHF5 | KPNA7 | Reptilia | Geospiza fortis (Medium ground-finch) |
| A0A1U7S409 | KPNA7 | Reptilia | Alligator sinensis (Chinese alligator) |
| A0A663F2M4 | KPNA7 | Reptilia | Aquila chrysaetos chrysaetos |
| A0A663F2P6 | KPNA7 | Reptilia | Aquila chrysaetos chrysaetos |
| A0A1V4KSF1 | KPNA7 | Reptilia | Patagioenas fasciata monilis |
| A0A672U8V3 | KPNA7 | Reptilia | Strigops habroptila (Kakapo) |
| G1MY68 | KPNA7 | Reptilia | Meleagris gallopavo (Wild turkey) |
| A0A6J3DSQ0 | KPNA7 | Reptilia | Aythya fuligula (Tufted duck) (Anas fuligula) |
| C0LLJ0 | KPNA7 | Mammalia | Mus musculus (Mouse) |
| A0A096MK49 | KPNA7 | Mammalia | Rattus norvegicus (Rat) |
| H0VQS0 | KPNA7 | Mammalia | Cavia porcellus (Guinea pig) |
| A0A6P7RAC8 | KPNA7 | Mammalia | Mus caroli (Ryukyu mouse) (Ricefield mouse) |
| A0A3Q0D6R9 | KPNA7 | Mammalia | Mesocricetus auratus (Golden hamster) |
| D3Z2P7 | KPNA7 | Mammalia | Mus musculus (Mouse) |
| A0A6P3VA68 | KPNA7 | Mammalia | Octodon degus (Degu) (Sciurus degus) |
| A0A6I9LF97 | KPNA7 | Mammalia | Peromyscus maniculatus bairdii (Prairie deer mouse) |
| A0A287DCQ1 | KPNA7 | Mammalia | Ictidomys tridecemlineatus (Thirteen-lined ground squirrel) (Spermophilus tridecemlineatus) |
| M3YYN0 | KPNA7 | Mammalia | Mustela putorius furo (European domestic ferret) (Mustela furo) |
| A0A6J3AJ62 | KPNA7 | Mammalia | Vicugna pacos (Alpaca) (Lama pacos) |
| A0A6I9JDZ1 | KPNA7 | Mammalia | Chrysochloris asiatica (Cape golden mole) |
| G1RIW6 | KPNA7 | Mammalia | Nomascus leucogenys (Northern white-cheeked gibbon) (Hylobates leucogenys) |
| A0A673UH42 | KPNA7 | Mammalia | Suricata suricatta (Meerkat) |
| A0A2K6A4U0 | KPNA7 | Mammalia | Mandrillus leucophaeus (Drill) (Papio leucophaeus) |
| G1SG50 | KPNA7 | Mammalia | Oryctolagus cuniculus (Rabbit) |
| A0A6P6HTK7 | KPNA7 | Mammalia | Puma concolor (Mountain lion) |
| A0A2K5UXA0 | KPNA7 | Mammalia | Macaca fascicularis (Crab-eating macaque) (Cynomolgus monkey) |
| A0A667HG78 | KPNA7 | Mammalia | Lynx canadensis (Canada lynx) |
| A0A6P5E101 | KPNA7 | Mammalia | Bos indicus (Zebu) |
| A0A6P3HNK5 | KPNA7 | Mammalia | Bison bison bison |
| A0A6P3YTR5 | KPNA7 | Mammalia | Ovis aries (Sheep) |
| A0A6J0Y4Q7 | KPNA7 | Mammalia | Odocoileus virginianus texanus |
| F6R0P5 | KPNA7 | Mammalia | Macaca mulatta (Rhesus macaque) |
| H0XT12 | KPNA7 | Mammalia | Otolemur garnettii (Small-eared galago) (Garnett's greater bushbaby) |
| A0A3Q7RIG0 | KPNA7 | Mammalia | Vulpes vulpes (Red fox) |
| A0A6J1Y007 | KPNA7 | Mammalia | Acinonyx jubatus (Cheetah) |

| AC | GN | Class | Organism |
|---|---|---|---|
| A0A096P098 | KPNA7 | Mammalia | Papio anubis (Olive baboon) |
| A0A2K5KVE0 | KPNA7 | Mammalia | Cercocebus atys (Sooty mangabey) (Cercocebus torquatus atys) |
| A0A2U3YYF1 | KPNA7 | Mammalia | Leptonychotes weddellii (Weddell seal) (Otaria weddellii) |
| A0A2Y9G9Q9 | KPNA7 | Mammalia | Neomonachus schauinslandi (Hawaiian monk seal) (Monachus schauinslandi) |
| A0A5F8GUF5 | KPNA7 | Mammalia | Monodelphis domestica (Gray short-tailed opossum) |
| M3WLV5 | KPNA7 | Mammalia | Felis catus (Cat) (Felis silvestris catus) |
| H2PLH7 | KPNA7 | Mammalia | Pongo abelii (Sumatran orangutan) (Pongo pygmaeus abelii) |
| A0A455C3E3 | KPNA7 | Mammalia | Physeter macrocephalus (Sperm whale) (Physeter catodon) |
| A0A2K6QJE3 | KPNA7 | Mammalia | Rhinopithecus roxellana (Golden snub-nosed monkey) (Pygathrix roxellana) |
| A0A4W2BKA2 | KPNA7 | Mammalia | Bos indicus x Bos taurus (Hybrid cattle) |
| A0A341BFZ0 | KPNA7 | Mammalia | Neophocaena asiaeorientalis asiaeorientalis (Yangtze finless porpoise) (Neophocaena phocaenoides subsp. asiaeorientalis) |
| H9KVH0 | KPNA7 | Mammalia | Sus scrofa (Pig) |
| A0A6P3QNL6 | KPNA7 | Mammalia | Pteropus vampyrus (Large flying fox) |
| G3SVL2 | KPNA7 | Mammalia | Loxodonta africana (African elephant) |
| A0A3Q7NGF8 | KPNA7 | Mammalia | Callorhinus ursinus (Northern fur seal) |
| F7CWU3 | KPNA7 | Mammalia | Equus caballus (Horse) |
| A0A452G6T6 | KPNA7 | Mammalia | Capra hircus (Goat) |
| A0A340WH20 | KPNA7 | Mammalia | Lipotes vexillifer (Yangtze river dolphin) |
| A0A2R9C214 | KPNA7 | Mammalia | Pan paniscus (Pygmy chimpanzee) (Bonobo) |
| A0A3Q7XXW8 | KPNA7 | Mammalia | Ursus arctos horribilis |
| A0A2U3VHW3 | KPNA7 | Mammalia | Odobenus rosmarus divergens (Pacific walrus) |
| A0A1S3AKG9 | KPNA7 | Mammalia | Erinaceus europaeus (Western European hedgehog) |
| A0A2Y9PGE5 | KPNA7 | Mammalia | Delphinapterus leucas (Beluga whale) |
| F6TJ07 | KPNA7 | Mammalia | Ornithorhynchus anatinus (Duckbill platypus) |
| A0A2K6GMC3 | KPNA7 | Mammalia | Propithecus coquereli (Coquerel's sifaka) (Propithecus verreauxi coquereli) |
| A0A384AHF6 | KPNA7 | Mammalia | Balaenoptera acutorostrata scammoni (North Pacific minke whale) (Balaenoptera davidsoni) |
| G1LJH4 | KPNA7 | Mammalia | Ailuropoda melanoleuca (Giant panda) |
| G1NZS1 | KPNA7 | Mammalia | Myotis lucifugus (Little brown bat) |
| A0A452QVF9 | KPNA7 | Mammalia | Ursus americanus (American black bear) (Euarctos americanus) |
| A0A2K6DYG1 | KPNA7 | Mammalia | Macaca nemestrina (Pig-tailed macaque) |
| A0A6P4TYG7 | KPNA7 | Mammalia | Panthera pardus (Leopard) (Felis pardus) |
| A0A6J3GXN8 | KPNA7 | Mammalia | Sapajus apella (Brown-capped capuchin) (Cebus apella) |
| A0A2K6L189 | KPNA7 | Mammalia | Rhinopithecus bieti (Black snub-nosed monkey) (Pygathrix bieti) |
| A0A384CQH9 | KPNA7 | Mammalia | Ursus maritimus (Polar bear) (Thalarctos maritimus) |
| A9QM74 | KPNA7 | Mammalia | Homo sapiens (Human) |
| C1JZ66 | KPNA7 | Mammalia | Bos taurus (Bovine) |
