## Supporting file 2 for "Biochemical propensity mapping for structural and functional anatomy of importin α IBB domain"

```
                      *        20         *        40         *        60         *        80         *
A0A1A8RH86 : -EHNLIGITMS--Q--S-----KD-NAR-LKSYKNKSLNTDEMRRRREEEGLQLRKQKKDEQL----FKR---RN----V-A--TVEDE------G--
A0A1A8JDE8 : ---------MS--Q--S-----KD-NAR-LKSYKNKSLNTDEMRRRREEEGLQLRKQKKDEQL----FKR---RN----V-A--TVEDE------G--
A0A1A8EKN4 : ---------MS--Q--S-----KD-NAR-LKSYKNKSLNTDEMRRRREEEGLQLRKQKKDEQL----FKR---RN----V-A--TVEDE------G--
A0A1A8LEE8 : -ISVLSGITMS--Q--S-----KD-NAR-LKSYKNKSLNTDEMRRRREEEGLQLRKQKKDEQL----FKR---RN----V-A--TVEDE------G--
A0A1A8A5F9 : ---------MS--Q--S-----KD-NAR-LKSYKNKSLNTDEMRRRREEEGLQLRKQKKDEQL----FKR---RN----V-A--TVEDE------G--
A0A1A8C032 : ---------MS--Q--S-----KD-NAR-LKSYKNKSLNTDEMRRRREEEGLQLRKQKKDEQL----FKR---RN----V-A--TVEDE------G--
A0A671WDK5 : ---MSR-YTMS--Q--G-----KD-NAR-LKSYKNKSLNTDEMRRRREEEGLQLRKQKKDEQL----FKR---RN----V-A--TVEDD------G--
H2LZX3     : ---------MS--Q--G-----KD-NAR-LKSYKNKSLNTDEMRRRREEEGLQLRKQKKDEQL----FKR---RN----V-A--TVEDD------G--
A0A4W6DL87 : -KTVQTCYTMS--Q--G-----KD-NAR-LKSYKNKSLNTDEMRRRREEEGLQLRKQKKDEQL----FKR---RN----V-A--TVEDD------G--
A0A4Z2JG54 : ---------MS--Q--G-----KD-NAR-LKSYKNKSLNTDEMRRRREEEGLQLRKQKKDEQL----FKR---RN----V-A--TVEDD------G--
A0A4W6DJC0 : -EPNLTGYTMS--Q--G-----KD-NAR-LKSYKNKSLNTDEMRRRREEEGLQLRKQKKDEQL----FKR---RN----V-A--TVEDD------G--
A0A6P7KJE0 : ---------MS--Q--G-----KD-NAR-LKSYKNKSLNTDEMRRRREEEGLQLRKQKKDEQL----FKR---RN----V-A--TVEDD------G--
A0A1A7Y2I3 : ---------MS--Q--S-----KD-NAR-LKSYKNKSLNTDEMRRRREEEGLQLRKQKKDEQL----FKR---RN----V-A--TVEDD------G--
A0A6J2RVP8 : ---------MS--Q--S-----KD-NAR-LKSYKNKSLNTDEMRRRREEEGLQLRKQKKDEQL----FKR---RN----V-A--TVEDD------G--
E6ZIU3     : -FCISLGCTMS--Q--G-----KD-NAR-LKSYKNKSLNTDEMRRRREEEGLQLRKQKKDEQL----FKR---RN----V-A--TVEDD------G--
A0A2I4AQA5 : ---------MS--Q--G-----KD-NAR-LKSYKNKSLNTDEMRRRREEEGLQLRKQKKDEQL----FKR---RN----V-A--TVEDD------G--
A0A6P8SZR2 : -FCISPGYTMS--Q--G-----KD-NAR-LKSYKNKSLNTDEMRRRREEEGLQLRKQKKDEQL----FKR---RN----V-A--TVEDD------G--
A0A671W6K4 : --MSYL----------G-----KD-NAR-LKSYKNKSLNTDEMRRRREEEGLQLRKQKKDEQL----FKR---RN----V-A--TVEDD------G--
A0A4W6DKG1 : --MSHQ----------G-----KD-NAR-LKSYKNKSLNTDEMRRRREEEGLQLRKQKKDEQL----FKR---RN----V-A--TVEDD------G--
A0A4W6DL44 : --YTLVKDTYAF-Q--G-----KD-NAR-LKSYKNKSLNTDEMRRRREEEGLQLRKQKKDEQL----FKR---RN----V-A--TVEDD------G--
A0A4W6DLH5 : ----L-----A-----G-----KD-NAR-LKSYKNKSLNTDEMRRRREEEGLQLRKQKKDEQL----FKR---RN----V-A--TVEDD------G--
A0A4W6DJY6 : -ELTLAK---S-----G-----KD-NAR-LKSYKNKSLNTDEMRRRREEEGLQLRKQKKDEQL----FKR---RN----V-A--TVEDD------G--
A0A4W6DL04 : ---------MT-----G-----KD-NAR-LKSYKNKSLNTDEMRRRREEEGLQLRKQKKDEQL----FKR---RN----V-A--TVEDD------G--
A0A6P3VH89 : ---------MS--Q--G-----KD-NAR-LKSYKNKSLNTDEMRRRREEEGLQLRKQKKDEQL----FKR---RN----V-A--TVEED------G--
F1QB53     : -AVLTADNIMS--Q--G-----KD-NAR-LKSYKNKSLNTDEMRRRREEEGLQLRKQKKDEQL----FKR---RN----V-A--TVEED------G--
A0A4W4H6M5 : SEVFTC-CIMS--Q--G-----KD-NAR-LRSYKNKSLNTDEMRRRREEEGLQLRKQKKDEQL----FKR---RN----V-A--TVEED------C--
W5U9X8     : ---------MS--Q--G-----KD-NAR-LRSYKNKSLNTDEMRRRREEEGLQLRKQKKDEQL----FKR---RN----V-A--TVEED------C--
A0A4W4GZA4 : FKSLIR-CIMS--Q--G-----KD-NAR-LRSYKNKSLNTDEMRRRREEEGLQLRKQKKDEQL----FKR---RN----V-A--TVEED------C--
A0A6J2UXX7 : ---------MS--Q--G-----KD-NAR-LKSYKNKSLNTEEMRRRREEEGLQLRKQKKDEQL----FKR---RN----V-A--TVEED------S--
A0A4W4H6H5 : ---MKNG--IV--Q--G-----KD-NAR-LRSYKNKSLNTDEMRRRREEEGLQLRKQKKDEQL----FKR---RN----V-A--TVEED------C--
A0A4W4H6S2 : ---MAK----V-----G-----KD-NAR-LRSYKNKSLNTDEMRRRREEEGLQLRKQKKDEQL----FKR---RN----V-A--TVEED------C--
A0A671WDU1 : -----KDAL----Q--G-----KD-NAR-LKSYKNKSLNTDEMRRRREEEGLQLRKQKKDEQ------HFFCVLSY-LHVPA--MIFSS---------
A0A4W4H6S8 : ----VLGCIMS--Q--G-----KD-NAR-LRSYKNKSLNTDEMRRRREEEGLQLRKQKKDEQ------GG--VISEDM-I-Q--MIFSS------S--
A0A671WDF6 : ----NKNYRHL--R--G-----KD-NAR-LKSYKNKSLNTDEMRRRREEEGLQLRKQKKDEQ------KK---NNPHFFC-G--GVISE------D--
A0A671W6F8 : ----MRSYTMS--Q--G-----KD-NAR-LKSYKNKSLNTDEMRRRREEEGLQLRKQKKDEQ------KK---NNPHFFC-G--GVISE------D--
A0A4W6DJI5 : --------L----D--G-----KD-NAR-LKSYKNKSLNTDEMRRRREEEGLQLRKQKKDEQL----FKR---RNV-SDV----TVFH----------
A0A1L8H5K7 : ---------MTS-S--G-----KE-NSR-LKSYKNKSLNTDEMRRRREEEGLQLRKQKREEQL----FKR---RN----V-A--CAEEA------T--
A0A6I8RJ96 : --VEAEQEIMTS-S--G-----KE-NSR-LKSYKNKSLNTDEMRRRREEEGLQLRKQKREEQL----FKR---RN----V-A--CAEEA------T--
F6TUG5     : --VYLPTEIMTS-S--G-----KE-NSR-LKSYKNKSLNTDEMRRRREEEGLQLRKQKREEQL----FKR---RN----V-A--CAEEA------T--
H3A826     : ---------MTS-S--G-----KE-NLW-QKRYKNKSLNADEMRRRREEEGLQLRKQKREEQL----FKR---RN----V-A--SAEEE------P--
A0A6I8RA27 : ---------MKN-GVYGRRPFRKE-NSR-LKSYKNKSLNTDEMRRRREEEGLQLRKQKREEQL----FKR---RN----V-A--CAEEA------T--
P83953     : ---------MST-P--G-----KE-NFR-LKSYKNKSLNPDEMRRRREEEGLQLRKQKREEQL----FKR---RN----V-A--TAEEE------T--
Q60960     : ---------MST-P--G-----KE-NFR-LKSYKNKSLNPDEMRRRREEEGLQLRKQKREEQL----FKR---RN----V-A--TAEEE------T--
A0A6P5RCS7 : ---------MST-P--G-----KE-NFR-LKSYKNKSLNPDEMRRRREEEGLQLRKQKREEQL----FKR---RN----V-A--TAEEE------T--
G1SI36     : ---------MST-P--G-----KE-NFR-LKSYKNKSLNPDEMRRRREEEGLQLRKQKREEQL----FKR---RN----V-A--TAEEE------T--
F6QFP5     : ---------MST-P--G-----KE-NFR-LKSYKNKSLNPDEMRRRREEEGLQLRKQKREEQL----FKR---RN----V-A--TAEEE------T--
A0A1S3FE51 : ---------MTT-P--G-----KE-NFR-LKSYKNKSLNPDEMRRRREEEGLQLRKQKREEQL----FKR---RN----V-A--TAEEE------T--
A0A6P4VZJ1 : --KSTSKEIMTT-P--G-----KE-NFR-LKSYKNKSLNPDEMRRRREEEGLQLRKQKREEQL----FKR---RN----V-A--TAEEE------T--
A0A6P5B348 : ---------MTT-P--G-----KE-NFR-LKSYKNKSLNPDEMRRRREEEGLQLRKQKREEQL----FKR---RN----V-A--TAEEE------T--
H2QN78     : ---------MTT-P--G-----KE-NFR-LKSYKNKSLNPDEMRRRREEEGLQLRKQKREEQL----FKR---RN----V-A--TAEEE------T--
A0A6J3H055 : ---------MTT-P--G-----KE-NFR-LKSYKNKSLNPDEMRRRREEEGLQLRKQKREEQL----FKR---RN----V-A--TAEEE------T--
A0A6P6H2Z3 : --KSTSEEIMTT-P--G-----KE-NFR-LKSYKNKSLNPDEMRRRREEEGLQLRKQKREEQL----FKR---RN----V-A--TAEEE------T--
A0A6J1ZKI4 : --KSTSEEIMTT-P--G-----KE-NFR-LKSYKNKSLNPDEMRRRREEEGLQLRKQKREEQL----FKR---RN----V-A--TAEEE------T--
A0A1S3AD74 : ---------MTT-P--G-----KE-NFR-LKSYKNKSLNPDEMRRRREEEGLQLRKQKREEQL----FKR---RN----V-A--TAEEE------T--
A0A6J2L0N9 : ---------MTT-P--G-----KE-NFR-LKSYKNKSLNPDEMRRRREEEGLQLRKQKREEQL----FKR---RN----V-A--TAEEE------T--
A0A6J2C9M7 : ---------MTT-P--G-----KE-NFR-LKSYKNKSLNPDEMRRRREEEGLQLRKQKREEQL----FKR---RN----V-A--TAEEE------T--
A0A2K5Y941 : ---------MTT-P--G-----KE-NFR-LKSYKNKSLNPDEMRRRREEEGLQLRKQKREEQL----FKR---RN----V-A--TAEEE------T--
A0A2K5IUS9 : ---------MTT-P--G-----KE-NFR-LKSYKNKSLNPDEMRRRREEEGLQLRKQKREEQL----FKR---RN----V-A--TAEEE------T--
A0A3Q7WJB8 : ---------MTT-P--G-----KE-NFR-LKSYKNKSLNPDEMRRRREEEGLQLRKQKREEQL----FKR---RN----V-A--TAEEE------T--
P52294     : ---------MTT-P--G-----KE-NFR-LKSYKNKSLNPDEMRRRREEEGLQLRKQKREEQL----FKR---RN----V-A--TAEEE------T--
G1M736     : --SLIFVENMTT-P--G-----KE-NFR-LKSYKNKSLNPDEMRRRREEEGLQLRKQKREEQL----FKR---RN----V-A--TAEEE------T--
A0A667HMJ9 : ---------MTT-P--G-----KE-NFR-LKSYKNKSLNPDEMRRRREEEGLQLRKQKREEQL----FKR---RN----V-A--TAEEE------T--
A0A2K6NKC7 : ---------MTT-P--G-----KE-NFR-LKSYKNKSLNPDEMRRRREEEGLQLRKQKREEQL----FKR---RN----V-A--TAEEE------T--
```

```
                       *         20         *         40         *         60         *         80         *
A0A2K5N988 : ---------MTT-P--G-----KE-NFR-LKSYKNKSLNPDEMRRRREEEGLQLRKQKREEQL----FKR---RN----V-A--TAEEE------T--
A0A6P3R5V2 : ---------MTT-P--G-----KE-NFR-LKSYKNKSLNPDEMRRRREEEGLQLRKQKREEQL----FKR---RN----V-A--TAEEE------T--
A0A2K6LM96 : ---------MTT-P--G-----KE-NFR-LKSYKNKSLNPDEMRRRREEEGLQLRKQKREEQL----FKR---RN----V-A--TAEEE------T--
A0A5G2QTJ8 : ------VEIMTT-P--G-----KE-NFR-LKSYKNKSLNPDEMRRRREEEGLQLRKQKREEQL----FKR---RN----V-A--TAEEE------T--
G3UJ13     : --SLNLVEIMTT-P--G-----KE-NFR-LKSYKNKSLNPDEMRRRREEEGLQLRKQKREEQL----FKR---RN----V-A--TAEEE------T--
A0A6I9K6T6 : ---------MTT-P--G-----KE-NFR-LKSYKNKSLNPDEMRRRREEEGLQLRKQKREEQL----FKR---RN----V-A--TAEEE------T--
A0A1D5QJL1 : ---------MTT-P--G-----KE-NFR-LKSYKNKSLNPDEMRRRREEEGLQLRKQKREEQL----FKR---RN----V-A--TAEEE------T--
A0A337S696 : ---------MTT-P--G-----KE-NFR-LKSYKNKSLNPDEMRRRREEEGLQLRKQKREEQL----FKR---RN----V-A--TAEEE------T--
A0A0D9R4X1 : ---------MTT-P--G-----KE-NFR-LKSYKNKSLNPDEMRRRREEEGLQLRKQKREEQL----FKR---RN----V-A--TAEEE------T--
M3Y068     : ---------MTT-P--G-----KE-NFR-LKSYKNKSLNPDEMRRRREEEGLQLRKQKREEQL----FKR---RN----V-A--TAEEE------T--
A2VE08     : ---------MTT-P--G-----KE-NFR-LKSYKNKSLNPDEMRRRREEEGLQLRKQKREEQL----FKR---RN----V-A--TAEEE------T--
A0A4W2FGB4 : --LWSLSEIMTT-P--G-----KE-NFR-LKSYKNKSLNPDEMRRRREEEGLQLRKQKREEQL----FKR---RN----V-A--TAEEE------T--
A0A2I2YX55 : ---------MTT-P--G-----KE-NFR-LKSYKNKSLNPDEMRRRREEEGLQLRKQKREEQL----FKR---RN----V-A--TAEEE------T--
A0A3Q7RJW3 : ---------MTT-P--G-----KE-NFR-LKSYKNKSLNPDEMRRRREEEGLQLRKQKREEQL----FKR---RN----V-A--TAEEE------T--
A0A2Y9HG96 : ---------MTT-P--G-----KE-NFR-LKSYKNKSLNPDEMRRRREEEGLQLRKQKREEQL----FKR---RN----V-A--TAEEE------T--
A0A2R9BX67 : ---------MTT-P--G-----KE-NFR-LKSYKNKSLNPDEMRRRREEEGLQLRKQKREEQL----FKR---RN----V-A--TAEEE------T--
A0A2K6EIA0 : ---------MTT-P--G-----KE-NFR-LKSYKNKSLNPDEMRRRREEEGLQLRKQKREEQL----FKR---RN----V-A--TAEEE------T--
Q5R909     : ---------MTT-P--G-----KE-NFR-LKSYKNKSLNPDEMRRRREEEGLQLRKQKREEQL----FKR---RN----V-A--TAEEE------T--
A0A384DH77 : --LSPFPENMTT-P--G-----KE-NFR-LKSYKNKSLNPDEMRRRREEEGLQLRKQKREEQL----FKR---RN----V-A--TAEEE------T--
F1PFK6     : ---------MTT-P--G-----KE-NFR-LKSYKNKSLNPDEMRRRREEEGLQLRKQKREEQL----FKR---RN----V-A--TAEEE------T--
A0A6J0Z446 : ---------MTT-P--G-----KE-NFR-LKSYKNKSLNPDEMRRRREEEGLQLRKQKREEQL----FKR---RN----V-A--TAEEE------T--
A0A2K5DU08 : ---------MTT-P--G-----KE-NFR-LKSYKNKSLNPDEMRRRREEEGLQLRKQKREEQL----FKR---RN----V-A--TAEEE------T--
A0A2U3VBV0 : ---------MTT-P--G-----KE-NFR-LKSYKNKSLNPDEMRRRREEEGLQLRKQKREEQL----FKR---RN----V-A--TAEEE------T--
A0A2K6DHG5 : ---------MTT-P--G-----KE-NFR-LKSYKNKSLNPDEMRRRREEEGLQLRKQKREEQL----FKR---RN----V-A--TAEEE------T--
A0A0B8RTI9 : ---------MTT-P--G-----KE-NFR-LKSYKNKSLNPDEMRRRREEEGLQLRKQKREEQL----FKR---RN----V-A--TAEEE------T--
A0A384DH80 : ------MKNMTT-P--G-----KE-NFR-LKSYKNKSLNPDEMRRRREEEGLQLRKQKREEQL----FKR---RN----V-A--TAEEE------T--
A0A6P3H099 : ---------MTT-P--G-----KE-NFR-LKSYKNKSLNPDEMRRRREEEGLQLRKQKREEQL----FKR---RN----V-A--TAEEE------T--
A0A2Y9SJ29 : ---------MTT-P--G-----KE-NFR-LKSYKNKSLNPDEMRRRREEEGLQLRKQKREEQL----FKR---RN----V-A--TAEEE------T--
A0A3Q7PAX2 : ---------MTT-P--G-----KE-NFR-LKSYKNKSLNPDEMRRRREEEGLQLRKQKREEQL----FKR---RN----V-A--TAEEE------T--
A0A452RY52 : ---------MTT-P--G-----KE-NFR-LKSYKNKSLNPDEMRRRREEEGLQLRKQKREEQL----FKR---RN----V-A--TAEEE------T--
A0A3Q0CSU4 : ---------MTT-P--G-----KE-NFR-LKSYKNKSLNPDEMRRRREEEGLQLRKQKREEQL----FKR---RN----V-A--TAEEE------T--
A0A4W2FGM4 : --YSMPCEIMTT-P--G-----KE-NFR-LKSYKNKSLNPDEMRRRREEEGLQLRKQKREEQL----FKR---RN----V-A--TAEEE------T--
G1NUG8     : ---------MTT-P--G-----KE-NFR-LKSYKNKSLNPDEMRRRREEEGLQLRKQKREEQL----FKR---RN----V-A--TAEEE------T--
A0A2U3YI58 : ---------MTT-P--G-----KE-NFR-LKSYKNKSLNPDEMRRRREEEGLQLRKQKREEQL----FKR---RN----V-A--TAEEE------T--
A0A671DWC4 : ---------MTT-P--G-----KE-NFR-LKSYKNKSLNPDEMRRRREEEGLQLRKQKREEQL----FKR---RN----V-A--TAEEE------T--
A0A2K6TSQ2 : ---------MTT-P--G-----KE-NFR-LKSYKNKSLNPDEMRRRREEEGLQLRKQKREEQL----FKR---RN----V-A--TAEEE------T--
I3N7W7     : ---------MTT-P--G-----KE-NFR-LKSYKNKSLNPDEMRRRREEEGLQLRKQKREEQL----FKR---RN----V-A--TAEEE------T--
I3L9F7     : --GRLEPEIMTT-P--G-----KE-NFR-LKSYKNKSLNPDEMRRRREEEGLQLRKQKREEQL----FKR---RN----V-A--TAEEE------T--
A0A2K5WRD1 : ---------MTT-P--G-----KE-NFR-LKSYKNKSLNPDEMRRRREEEGLQLRKQKREEQL----FKR---RN----V-A--TAEEE------T--
A0A2K5RVS8 : ---------MTT-P--G-----KE-NFR-LKSYKNKSLNPDEMRRRREEEGLQLRKQKREEQL----FKR---RN----V-A--TAEEE------T--
A0A2I3MWI4 : ---------MTT-P--G-----KE-NFR-LKSYKNKSLNPDEMRRRREEEGLQLRKQKREEQL----FKR---RN----V-A--TAEEE------T--
A0A6I9IM75 : ---------MTT-P--G-----KE-NFR-LKSYKNKSLNPDEMRRRREEEGLQLRKQKREEQL----FKR---RN----V-A--TAEEE------T--
A0A341DEI5 : ---------MTT-P--G-----RE-NFR-LKSYKNKSLNPDEMRRRREEEGLQLRKQKREEQL----FKR---RN----V-A--TAEEE------T--
A0A340WM35 : ---------MTT-P--G-----RE-NFR-LKSYKNKSLNPDEMRRRREEEGLQLRKQKREEQL----FKR---RN----V-A--TAEEE------T--
A0A2U4AAL0 : ---------MTT-P--G-----RE-NFR-LKSYKNKSLNPDEMRRRREEEGLQLRKQKREEQL----FKR---RN----V-A--TAEEE------T--
A0A2Y9MUZ0 : ---------MTT-P--G-----RE-NLR-LKSYKNKSLNPDEMRRRREEEGLQLRKQKREEQL----FKR---RN----V-A--TAEEE------T--
A0A452G7U7 : ---------MTA-P--G-----KE-NFR-LKSYKNKSLNPDEMRRRREEEGLQLRKQKREEQL----FKR---RN----V-A--TAEEE------T--
W5QFX7     : ---------MTA-P--G-----KE-NFR-LKSYKNKSLNPDEMRRRREEEGLQLRKQKREEQL----FKR---RN----V-A--TAEEE------T--
F7ISW6     : ---------MTA-P--G-----KE-NFR-LKSYKNKSLNPDEMRRRREEEGLQLRKQKREEQL----FKR---RN----V-A--TAEEE------T--
A0A673T529 : ---------MTA-P--G-----KE-NFR-LKSYKNKSLNPDEMRRRREEEGLQLRKQKREEQL----FKR---RN----V-A--TAEEE------T--
A0A6P6DCY3 : ---------MTT-P--G-----KE-NFR-LKSYKNKSLNPDEMRRRREEEGLQLRKQKREEQL----FKR---RN----V-A--TAEEE------A--
H0VB68     : ---------MTT-P--G-----KE-NFR-LKSYKNKSLNPDEMRRRREEEGLQLRKQKREEQL----FKR---RN----V-A--TAEEE------A--
A0A2I3HG58 : ---------MTT-Q--G-----KE-NFR-LKSYKNKSLNPDEMRRRREEEGLQLRKQKREEQL----FKR---RN----V-A--TAEEE------T--
A0A383ZKK2 : ---------MTT-Q--G-----KE-NFR-LKSYKNKSLNPDEMRRRREEEGLQLRKQKREEQL----FKR---RN----V-A--TAEEE------T--
A0A3Q0FNG3 : ---------MTT-S--G-----KE-NFR-LKSYKNKSLNPDEMRRRREEEGLQLRKQKREEQL----FKR---RN----V-A--TAEEE------T--
F6PVH5     : ------MVIMTT-S--G-----KE-NFR-LKSYKNKSLNPDEMRRRREEEGLQLRKQKREEQL----FKR---RN----V-A--TAEEE------T--
A0A6I9HG44 : ---------MTT-S--G-----KE-NFR-LKSYKNKSLNPDEMRRRREEEGLQLRKQKREEQL----FKR---RN----V-A--TAEEE------T--
A0A669QB17 : --WAGAAGTMTT-S--G-----KE-NFR-LKSYKNKSLNPDEMRRRREEEGLQLRKQKREEQL----FKR---RN----V-A--TAEEE------A--
A0A674I969 : ---------MTT-S--G-----KE-NFR-LKSYKNKSLNPDEMRRRREEEGLQLRKQKREEQL----FKR---RN----V-A--TAEEE------A--
A0A6J0H909 : ---------MTT-S--G-----KE-NFR-LKSYKNKSLNPDEMRRRREEEGLQLRKQKREEQL----FKR---RN----V-A--TAEEE------A--
U3IB40     : ---------MTT-S--G-----KE-NFR-LKSYKNKSLNPDEMRRRREEEGLQLRKQKREEQL----FKR---RN----V-A--TAEEE------A--
A0A663LTN7 : ---------MTT-S--G-----KE-NFR-LKSYKNKSLNPDEMRRRREEEGLQLRKQKREEQL----FKR---RN----V-A--TAEEE------A--
A0A1V4JB82 : --WKNSKGIMTT-S--G-----KE-NFR-LKSYKNKSLNPDEMRRRREEEGLQLRKQKREEQL----FKR---RN----V-A--TAEEE------A--
```

```
                     *        20         *        40         *        60         *        80         *
A0A6J2HDZ7 : ---------MTT-S--G-----KE-NFR-LKSYKNKSLNPDEMRRRREEEGLQLRKQKREEQL----FKR---RN----V-A--TAEEE------A--
K7GFN3     : ---------MTT-S--G-----KE-NFR-LKSYKNKSLNPDEMRRRREEEGLQLRKQKREEQL----FKR---RN----V-A--TAEEE------A--
A0A674GRG3 : ---------MTT-S--G-----KE-NFR-LKSYKNKSLNPDEMRRRREEEGLQLRKQKREEQL----FKR---RN----V-A--TAEEE------A--
U3JQT0     : ---MCSHGIMTT-S--G-----KE-NFR-LKSYKNKSLNPDEMRRRREEEGLQLRKQKREEQL----FKR---RN----V-A--TAEEE------A--
A0A6P5K899 : ---------MTT-S--G-----KE-NFR-LKSYKNKSLNPDEMRRRREEEGLQLRKQKREEQL----FKR---RN----V-A--TAEEE------A--
A0A218VEF9 : ---------MTT-S--G-----KE-NFR-LKSYKNKSLNPDEMRRRREEEGLQLRKQKREEQL----FKR---RN----V-A--TAEEE------A--
A0A672U9J9 : ---------MTT-S--G-----KE-NFR-LKSYKNKSLNPDEMRRRREEEGLQLRKQKREEQL----FKR---RN----V-A--TAEEE------A--
A0A452I0C3 : -GRS-ILEIMTT-S--G-----KD-NFR-LKSYKNKSLNPDEMRRRREEEGLQLRKQKREEQL----FKR---RN----V-A--TAEEE------A--
A0A452J830 : ---------MTT-S--G-----KE-NFR-LKSYKNKSLNPDEMRRRREEEGLQLRKQKREEQL----FKR---RN----V-A--TAEEE------A--
A0A2I0ML81 : ---------MTT-S--G-----KE-NFR-LKSYKNKSLNPDEMRRRREEEGLQLRKQKREEQL----FKR---RN----V-A--TAEEE------A--
A0A452I0D9 : --KTGILEIMTT-S--G-----KD-NFR-LKSYKNKSLNPDEMRRRREEEGLQLRKQKREEQL----FKR---RN----V-A--TAEEE------A--
A0A4X2KDM2 : ---------MTT-S--G-----KE-NFR-LKSYKNKSLNPDEMRRRREEEGLQLRKQKREEQL----FKR---RN----V-A--TAEEE------A--
A0A091PWE7 : ---------MTT-S--G-----KE-NFR-LKSYKNKSLNPDEMRRRREEEGLQLRKQKREEQL----FKR---RN----V-A--TAEEE------A--
A0A6J3DK49 : ---------MTT-S--G-----KE-NFR-LKSYKNKSLNPDEMRRRREEEGLQLRKQKREEQL----FKR---RN----V-A--TAEEE------A--
G1NMJ6     : ---------MTT-S--G-----KE-NFR-LKSYKNKSLNPDEMRRRREEEGLQLRKQKREEQL----FKR---RN----V-A--TAEEE------A--
A0A663FFE2 : ---------MTT-S--G-----KE-NFR-LKSYKNKSLNPDEMRRRREEEGLQLRKQKREEQL----FKR---RN----V-A--TAEEE------A--
A0A670JR46 : ---------MTT-S--G-----KE-NFR-LKSYKNKSLNPDEMRRRREEEGLQLRKQKREEQL----FKR---RN----V-A--TADEE------A--
H9GBL6     : --PAANAGTMTT-S--G-----KE-NFR-LKSYKNKSLNPDEMRRRREEEGLQLRKQKREEQL----FKR---RN----V-A--TADEE------A--
A0A6J0TGA2 : ---------MTT-S--G-----KE-NFR-LKSYKNKSLNPDEMRRRREEEGLQLRKQKREEQL----FKR---RN----V-A--TADEE------G--
A0A6J1URU5 : ---------MTS-S--G-----KE-NFR-LKSYKNKSLNPDEMRRRREEEGLQLRKQKREEQL----FKR---RN----V-A--TADEE------T--
A0A6I9XTZ4 : ---------MTS-S--G-----KE-NFR-LKSYKNKSLNPDEMRRRREEEGLQLRKQKREEQL----FKR---RN----V-A--TADEE------T--
A0A670ZFW8 : ---------MTS-S--G-----KE-NFR-LKSYKNKSLNPDEMRRRREEEGLQLRKQKREEQL----FKR---RN----V-A--TADEE------T--
V8P5N0     : ---------MTS-S--G-----KE-NFR-LKSYKNKSLNPDEMRRRREEEGLQLRKQKREEQL----FKR---RN----V-A--TADEE------T--
A0A6P9BB21 : ---------MTS-S--G-----KE-NFR-LKSYKNKSLNPDEMRRRREEEGLQLRKQKREEQL----FKR---RN----V-A--TADEE------T--
A0A6P7XWN2 : ---------MTS-S--G-----KE-NFR-LKSYKNKXLNPDEMRRRREEEGLQLRKQKREEQL----FKR---RN----V-A--NAEEE------T--
A0A6P8RC89 : ---------MTS-S--G-----KE-NFR-LKSYKNKSLNPDEMRRRREEEGLQLRKQKREEQL----FKR---RN----V-A--NAEEE------T--
A0A6G1ATK9 : ------VEIMTT-P--G-----KE-NFR-LKSYKNKSLNPDEMRRRREEEGLQLRKQKREEQI----SMI---NL----I-I--WPQP----IL-S--
A0A3Q1LXF0 : ------VEIMTT-P--G-----KE-NFR-LKSYKNKSLNPDEMRRRREEEGLQLRKQKREEQV-----M----EP----F-L--WMSSYLGGVITSDM
A0A2I3TP22 : ------VEIMTT-P--G-----KE-NFR-LKSYKNKSLNPDEMRRRREEEGLQLRKQKREEQV----------NP----LPL--FISFK---EF-NG-
A0A6I8PYZ6 : ---------MTS-S--G-----KE-NSR-LKSYKNKSLNTDEMRRRREEEGLQLRKQKREEQVCVSQFSR---NS----L-V--YNQA-------S--
A0A4Z2HHP2 : ---------MPT-K--N-----VD-DQR-LSKFKNKGKDPAKLREKRISVCVELRKAHKDESF----LKR---RN----ITL-SSLPD----------
P52170     : ----------PT-T--N-----EA-DER-MRKFKNKGKDTAELRRRRVEVSVELRKAKKDEQI----LKR---RN----VCLPEELI-----------
A0A218UWZ4 : ---------MPVRK--H-----GG-HMK-L--FKNKGKDQTTLRRQRVEVSVELRKAKKDEQI----LKR---RN----ISI--NLEKE---------
A0A2S2R7W1 : ---------MPT-K--K-----ISVNSRSFDVYQDRNKYYEGLRRKRIEVTVELRKTLRNEQM----SKH---RN----IVL--NDNDE---------
A0A6F9DGR6 : ---------MSL-T--E-----SQ-NRQNL--YKNKAGTSENLRKKRQECNVQLRKGKRDEQM----LKR---RN----INM-EDLNT----------
cIBB1      : ---------MTT-P--G-----KE-NFR-LKSYKNKSLNPDEMRRRREEEGLQLRKQKREEQL----FKR---RN----V-A--TAEEE------T--
```

```
                       *        20         *        40         *        60         *        80         *
A0A5N5TKN5 : ------PSTDGTT--------SR-----DASP-G---P-SR-IRNFK-NAGKDME----EMRRRRQETSVELRKQKKEDQLLKR-RNVT---LD-EPTS
A0A5B7IN23 : -----MPAP--AA--------AR-----DGSP-G---P-NR-LKSFK-NAGKDME----EMRRRRQETSIELRKAKKDDQLLKR-RNVT---LD-DPGS
A0A668SPP6 : -------MS--A---------GT-----D-N--G---A--R-LNQFK-NKGKDAD----ELRRRRAEVNVELRRAKKDDHILKR-RNVHSL-LD-EP--
A0A669F8L0 : -------MS--A---------GT-----D-N--G---A--R-LNQFK-NKGKDAD----ELRRRRAEVNVELRRAKKDDHILKR-RNVHSL-LD-EP--
A0A3P9AX06 : ----RSGMS--A---------GT-----D-N--G---A--R-LNQFK-NKGKDAN----ELRRRRAEVNVELRRAKKDDHILKR-RNVHSL-LD-EP--
A0A3P8NG90 : -------MS--A---------GT-----D-N--G---A--R-LNQFK-NKGKDAN----ELRRRRAEVNVELRRAKKDDHILKR-RNVHSL-LD-EP--
A0A3B4GL06 : -------MS--A---------GT-----D-N--G---A--R-LNQFK-NKGKDAN----ELRRRRAEVNVELRRAKKDDHILKR-RNVHSL-LD-EP--
A0A3Q4G708 : -------MS--A---------GT-----D-N--G---A--R-LNQFK-NKGKDAN----ELRRRRAEVNVELRRAKKDDHILKR-RNVHSL-LD-EP--
A0A3Q2VX29 : -------MS--A---------GT-----D-N--G---A--R-LNQFK-NKGKDAN----ELRRRRAEVNVELRRAKKDDHILKR-RNVHSL-LD-EP--
I3JGZ1     : ----------------------L-----A-Q--P------R-LNQFK-NKGKDAD----ELRRRRAEVNVELRRAKKDDHILKR-RNVHSL-LD-EP--
A0A668SNT5 : ----------------------L-----A-Q--P------R-LNQFK-NKGKDAD----ELRRRRAEVNVELRRAKKDDHILKR-RNVHSL-LD-EP--
A0A668SNT1 : ------NVTNEA---------VL-----K-N--PT--A--R-LNQFK-NKGKDAD----ELRRRRAEVNVELRRAKKDDHILKR-RNVHSL-LD-EP--
A0A3Q1HQH3 : -------MS--A---------GN-----D-N--G---A--R-LNQFK-NKGKDAT----ELRRRRAEVNVELRKAKKDDQILKR-RNVHSL-LD-EP--
A0A286R5L0 : -------MS--A---------GN-----D-N--G---A--R-LGQFK-NKGKDAT----ELRRRRAEVSVELRKAKKDDQILKR-RNVHSL-LE-EP--
A0A3Q0RB15 : -------MSATA---------GN-----D-S--G---A--R-LNQFK-NKGKDAN----ELRRTRAEVNVALRKAKKDDQILKR-RNVH-IPLD-EP--
A0A3B4WCW0 : -------MS--A---------GS-----D-N--G---A--R-LNQFK-NKGKDAN----ELRRRRAEVNVELRKAKKDDQILKR-RNVQSL-LD-EP--
A0A3B4V3Q9 : -------MS--A---------GS-----D-N--G---A--R-LNQFK-NKGKDAN----ELRRRRAEVNVELRKAKKDDQILKR-RNVQSL-LD-EP--
A0A4W6FTT1 : -------MS--A---------GS-----D-I--G---A--R-LTQFK-NKGKDAN----ELRRRRAEVNVELRKAKKEDQILKR-RNVHNL-SD-EP--
A0A3Q3MAK0 : RPAQRPEMS--A---------GP-----D-S--G---A--R-LTQFK-NKGKDAN----ELRRRRAEVNVELRKAKKDDQILKR-RNVHSL-LD-EP--
A0A3Q3F6G4 : -------MS--A---------GS-----D-T--E---T--R-LTQFK-NKGKDAS----ELRRRRAELNVEIRKAKKEDQILKR-RNVQSL-LD-EP--
A0A665VN51 : ----------------------I-----D-S--G---AGAR-LTQFK-NKGRDTN----ELRRRRAEVNVELRKAKKDDQILKR-RNVQTL-LD-EP--
A0A665VN42 : -------TD--A---------HFPANSPD-S--G---AGAR-LTQFK-NKGRDTN----ELRRRRAEVNVELRKAKKDDQILKR-RNVQTL-LD-EP--
A0A665VNE5 : -------MS--A---------GS-----D-S--G---AGAR-LTQFK-NKGRDTN----ELRRRRAEVNVELRKAKKDDQILKR-RNVQTL-LD-EP--
A0A3Q1B5I6 : RTVQRSGMS--A---------GN-----D-S--G---A--R-LSQFK-NKGKDASLQ--ELRRRRAEVNVELRKAKKDDQILKR-RNVQSL-DD-EP--
A0A3P8RZS7 : RTVQRSGMS--A---------GN-----D-S--G---A--R-LSQFK-NKGKDASLQ--ELRRRRAEVNVELRKAKKDDQILKR-RNVQSL-DD-EP--
A0A3B5BN85 : -------MS--A---------GN-----D-S--G---A--R-LSQFK-NKGKDAS----ELRRRRAEVNVELRKAKKDDQILKR-RNVQCL-ED-EP--
A0A672HYS4 : ----PSGMS--A---------GG-----D-N--G---A--R-LSQFK-NKGKDAH----ELRRRRAEVNVELRKAKKDDQILKR-RNVQAG-GG-IL--
A0A671TQX5 : -------MAAIV---------YN-----E-N--Q---HVDR-LNQFK-NKGKDAN----ELRRRRAEVNVELRKAKKDDQILKR-RNVQDL-LD-EP--
A0A671TQV7 : -------MS-YF---------YR-----D---------VDR-LNQFK-NKGKDAN----ELRRRRAEVNVELRKAKKDDQILKR-RNVQDL-LD-EP--
A0A671TR35 : -------MS--A---------AN-----D-S-------VDR-LNQFK-NKGKDAN----ELRRRRAEVNVELRKAKKDDQILKR-RNVQDL-LD-EP--
A0A4Z2G449 : -------MS--A---------VN-----D-S-------GDR-LPQFK-NKGKDAN----ELRRRRAEVNVELRKAKKDDQILKR-RNVQTL-LD-DP--
A0A3Q3JK14 : -------MS--A---------DN-----D-G--G---D--R-LNQFK-NKGKDAN----ELRRRRAEVSVELRKAKKDDQILKR-RNVLSL-SE-EP--
A0A3Q3B1Q0 : -------MS--G---------DN-----D-S--G---D--R-LHQFK-NKGKDAS----ELRRRRAEFSVELRKAKKDDQILKK-RNFGSL-VD-EP--
G3Q3L2     : -------MS--A---------SN-----D-D--G---DY-R-LGQFK-NKAKRVN----EMRRRRAEVNVQLRKAKKDDHILKK-RNVQSL-LD-EP--
A0A3Q3VX72 : -------MS--A---------AS-----D-S--G---D--R-LYQFK-NKGKDAN----ELRHRRVAVNMELRKAKKDDQILKK-RSVQSL-QECD---
A0A6P7J9M6 : -------MS--N---------GG-----V-S--G---D--R-LNRFK-NKGKDPN----ELRRRRAEVNVELRKAIKDEQILKR-RNVQCL-LE-EP--
A0A1A7ZW05 : -------ME-TS---------TD----YD-EL-G---T--R-LHQFK-NKGRNAT----EQTRRRKEFHVKLRKAKKDDQILER-RHVQTL-QE-EP--
A0A1A8C3Z5 : -------ME-TS---------TD----YD-EL-G---T--R-LHQFK-NKGRNAT----EQTRRRKEFHVKLRKAKKDDQILER-RHVQTL-QE-EP--
A0A1A8L5M0 : -------ME-MS---------TD----YD-EL-G---T--R-LHQFK-NKGKNAI----KQTRRRKEFHVKLRKAKKDDQILGR-RNVQTL-QE-EP--
A0A1A8EWU9 : -------ME-TS---------TD----YD-EL-G---T--R-LHQFK-NKGKNAI----KQTRRRKEFHVKLRKAKKDDQILGR-RNVQTL-QE-EP--
A0A1A8SK23 : -------ME-MS---------TD----YD-EL-G---T--R-LHQFK-NKGKNAI----KQTRRRKEFHVKLRKAKKDDQILGR-RNVQSL-QE-EP--
A0A1A8J5N3 : -------ME-MS---------TD----YD-EL-G---T--R-LHQFK-NKGKNAI----KQTRRRKEFHVKLRKAKKDDQILGR-RHVQTL-QD-EP--
A0A1A7XN45 : -------M--L----------TD----WD-EQ-G---D--R-LHQFK-NKGKDVI----EQRRRTEFHVELRKAKKDDQILKR-RNVQTL-QE-EL--
A0A3P9MSI2 : -------M---S---------AD----TD-N--G---A--R-LFQFK-NKGKDVN----ELRRRRSEFNVELRKAKRDVQILKR-RNAQSS-AE-DP--
A0A3B3TJ61 : -------M---S---------AD----TD-N--G---A--R-LFQFK-NKGKDVN----ELRRRRSEFNVELRKAKRDVQILKR-RNAQSS-AE-DP--
A0A3B3Z282 : -------M---S---------AD----TD-N--G---A--R-LFQFK-NKGKDVN----ELRRRRSEFNVELRKAKRDVQILKR-RNAQSS-AE-DP--
A0A087Y8H4 : -------M---S---------AD----TD-N--G---A--R-LFQFK-NKGKDVN----ELRRRRSEFNVELRKAKRDVQILKR-RNAQSS-AE-DP--
M4AGR3     : -------M---S---------AD----TD-N--G---A--R-LFQFK-NKGKDVN----ELRRRRSEFNVELRKAKRDVQILKR-RNAQGS-AE-DP--
A0A3Q2PPK5 : -------M---P---------GD----TD-S--G---A--R-LLQFK-NKGKDVN----ELRRRRAEFSVELRKAKKEHQILKR-RLVQSS-AD-EP--
A0A2I4CGJ0 : -------M---S---------GDN---TD--------V--R-LHQYK-NNGKDTD----EMRRRRAEVNVELRKTKRENQILKK-RNFQRD-PE-AP--
H2MF99     : -------M---S---------AGG---TD-N--P---A--R-IHLFK-NKGKCAL----ESLRRRAEVKVELRKAKKDEQIFKR-RAIQ---QD-DPAS
A0A3P9MPA0 : -------M---S---------AGG---TD-N--P---A--R-IHLFK-NKGKCAF----ESLRRRAEVKVELRKAKKDEQIFKR-RAIQ---QD-DPAS
A0A3B3BT30 : CLAQQCGM---S---------AG----TD-I--A---A--R-IHLFK-NKGKCVL----ESRRRRAEVNVELRKARKDEQILKR-RNVQSS-QD-NP--
A0A3B3ZC03 : -------ES------------TS-----R-S--AN-EA--R-LCQFK-NKGKDTN----ELRRRRAEVSVELRKAKKDEQILKR-RNVESPV-D-EH--
A0A3B3ZD83 : -------MS------------ADC----D-S--G---A--R-LCQFK-NKGKDTN----ELRRRRAEVSVELRKAKKDEQILKR-RNVESPV-D-EH--
A0A668AE89 : -------MS--A---------AN-----D-G--G---A--R-LTQFK-NKGKDAS----ELRRRRVEVNVELRKAKKDEQILKR-RNVAS-VED-EP--
A0A668A4Z8 : -----KPTNEEP---------AN-----D-G--G---A--R-LTQFK-NKGKDAS----ELRRRRVEVNVELRKAKKDEQILKR-RNVAS-VED-EP--
A0A3B4EME2 : -CYNIQKMSS-G---------SN-----D-S--V---A--R-LTQFK-NKGKDAN----ELRRRRIEVNVELRKAKKDEQILKR-RNVSCF-PD-GP--
W5L7R7     : -------MSS-G---------SN-----D-S--V---A--R-LNQFK-NKGKDAN----ELRRRRIEVSVELRKAKKDDQILKR-RNVSCF-PD-DA--
A0A2D0S4E7 : ----MEKMSSSA---------SS-----D-S--V---S--R-LNQFK-NKGKDAS----ELRRRRMEVSVELRKAKKDEQILKR-RNVSCLDPE-A---
A0A674D460 : -------MS-TT---------AN-----D-A--D------R-ITQFK-NKGKDTN----ELRRRRVEVNVELRKAKKDDQLLKR-RNVNIC-PD-EP--
A0A674BIA8 : -------MS-TT---------AN-----D-A--D------R-ITQFK-NKGKDTN----ELRRRRVEVNVELRKAKKDDQLLKR-RNVSIS-PD-EP--
A0A3P8YDR8 : -----TAMS-AT---------AN-----D-G--D------R-ISQFK-NKGKDTN----ELRRRRVEVNVELRKAKKDDQLLKR-RNVSIS-PD-EP--
```

```
                        *        20         *        40         *        60         *        80         *
A0A6Q2WUF4 : ----------VV---------SN-----E-PH-D------R-ISQFK-NKGKDTN----ELRRRRVEVNVELRKAKKDDQLLKR-RNVSIS-PD-EP--
I3NFF1     : -------MS--T----------N-----E-NA-N--SPAAR-LNRFK-NKGKDST----EMRRRRIEVNVELRKAKKDDQMLKR-RNVSS-FPD-DA--
A0A452UTX9 : ---FVSTMS--T----------N-----E-NA-N--SPAAR-LNRFK-NKGKDST----EMRRRRIEVNVELRKAKKDDQMLKR-RNVSS-FPD-DA--
A0A3Q0CQD4 : -------MS--T----------N-----E-NA-N--SPAAR-LNRFK-NKGKDST----EMRRRRIEVNVELRKAKKDDQMLKR-RNVSS-FPD-DA--
A0A1S3H041 : -------MS--T----------N-----E-NA-N--SPAAR-LNRFK-NKGKDST----EMRRRRIEVNVELRKAKKDDQMLKR-RNVSS-FPD-DA--
A0A1S3A179 : -------MS--T----------N-----E-NA-N--SPAAR-LNRFK-NKGKDST----EMRRRRIEVNVELRKAKKDDQMLKR-RNVSS-FPD-DA--
G3TP33     : -------MS--T----------N-----E-NA-N--SPAAR-LNRFK-NKGKDST----EMRRRRIEVNVELRKAKKDDQMLKR-RNVSS-FPD-DA--
E2R6L9     : -------MS--T----------N-----E-NA-N--SPAAR-LNRFK-NKGKDST----EMRRRRIEVNVELRKAKKDDQMLKR-RNVSS-FPD-DA--
A0A3Q7WT66 : -------MS--T----------N-----E-NA-N--SPAAR-LNRFK-NKGKDST----EMRRRRIEVNVELRKAKKDDQMLKR-RNVSS-FPD-DA--
A0A3Q7S7L5 : -------MS--T----------N-----E-NA-N--SPAAR-LNRFK-NKGKDST----EMRRRRIEVNVELRKAKKDDQMLKR-RNVSS-FPD-DA--
A0A341CRR5 : -------MS--T----------N-----E-NA-N--SPAAR-LNRFK-NKGKDST----EMRRRRIEVNVELRKAKKDDQMLKR-RNVSS-FPD-DA--
A0A384AFI2 : -------MS--T----------N-----E-NA-N--SPAAR-LNRFK-NKGKDST----EMRRRRIEVNVELRKAKKDDQMLKR-RNVSS-FPD-DA--
A0A2U3ZL89 : -------MS--T----------N-----E-NA-N--SPAAR-LNRFK-NKGKDST----EMRRRRIEVNVELRKAKKDDQMLKR-RNVSS-FPD-DA--
G1QDN1     : -----ITMS--T----------N-----E-NA-N--SPAAR-LNRFK-NKGKDST----EMRRRRIEVNVELRKAKKDDQMLKR-RNVSS-FPD-DA--
A0A340XCA9 : ---CLITMS--T----------N-----E-NA-N--SPAAR-LNRFK-NKGKDST----EMRRRRIEVNVELRKAKKDDQMLKR-RNVSS-FPD-DA--
A0A250YJD2 : -------MS--T----------N-----E-NA-N--SPAAR-LNRFK-NKGKDST----EMRRRRIEVNVELRKAKKDDQMLKR-RNVSS-FPD-DA--
A0A6I9KZ87 : -------MS--T----------N-----E-NA-N--SPAAR-LNRFK-NKGKDST----EMRRRRIEVNVELRKAKKDDQMLKR-RNVSS-FPD-DA--
A0A0P6IWF6 : -------MS--T----------N-----E-NA-N--SPAAR-LNRFK-NKGKDST----EMRRRRIEVNVELRKAKKDDQMLKR-RNVSS-FPD-DA--
A0A287AHI1 : -------MS--T----------N-----E-NA-N--SPAAR-LNRFK-NKGKDST----EMRRRRIEVNVELRKAKKDDQMLKR-RNVSS-FPD-DA--
A0A6G1ACR9 : -------MS--T----------N-----E-NA-N--SPAAR-LNRFK-NKGKDST----EMRRRRIEVNVELRKAKKDDQMLKR-RNVSS-FPD-DA--
G1MJD5     : -------MS--T----------N-----E-NA-N--SPAAR-LNRFK-NKGKDST----EMRRRRIEVNVELRKAKKDDQMLKR-RNVSS-FPD-DA--
A0A2Y9G4Z0 : -------MS--T----------N-----E-NA-N--SPAAR-LNRFK-NKGKDST----EMRRRRIEVNVELRKAKKDDQMLKR-RNVSS-FPD-DA--
A0A6J1YFZ8 : -------MS--T----------N-----E-NA-N--SPAAR-LNRFK-NKGKDST----EMRRRRIEVNVELRKAKKDDQMLKR-RNVSS-FPD-DA--
A0A6P3QVC4 : -------MS--T----------N-----E-NA-N--SPAAR-LNRFK-NKGKDST----EMRRRRIEVNVELRKAKKDDQMLKR-RNVSS-FPD-DA--
A0A6I9I0T4 : -------MS--T----------N-----E-NA-N--SPAAR-LNRFK-NKGKDST----EMRRRRIEVNVELRKAKKDDQMLKR-RNVSS-FPD-DA--
A0A6P6I5W2 : -------MS--T----------N-----E-NA-N--SPAAR-LNRFK-NKGKDST----EMRRRRIEVNVELRKAKKDDQMLKR-RNVSS-FPD-DA--
H0VVB1     : -------MS--T----------N-----E-NA-N--SPAAR-LNRFK-NKGKDST----EMRRRRIEVNVELRKAKKDDQMLKR-RNVSS-FPD-DA--
A0A2Y9PDA8 : -------MS--T----------N-----E-NA-N--SPAAR-LNRFK-NKGKDST----EMRRRRIEVNVELRKAKKDDQMLKR-RNVSS-FPD-DA--
A0A2U4C3S8 : -------MS--T----------N-----E-NA-N--SPAAR-LNRFK-NKGKDST----EMRRRRIEVNVELRKAKKDDQMLKR-RNVSS-FPD-DA--
A0A6P4X1F9 : -------MS--T----------N-----E-NA-N--SPAAR-LNRFK-NKGKDST----EMRRRRIEVNVELRKAKKDDQMLKR-RNVSS-FPD-DA--
A0A2U3XDN0 : -------MS--T----------N-----E-NA-N--SPAAR-LNRFK-NKGKDST----EMRRRRIEVNVELRKAKKDDQMLKR-RNVSS-FPD-DA--
A0A6P9F491 : -------MS--T----------N-----E-NA-N--SPAAR-LNRFK-NKGKDST----EMRRRRIEVNVELRKAKKDDQMLKR-RNVSS-FPD-DA--
A0A667I0F1 : -------MS--T----------N-----E-NA-N--SPAAR-LNRFK-NKGKDST----EMRRRRIEVNVELRKAKKDDQMLKR-RNVSS-FPD-DA--
A0A3Q2H8G1 : -------MS--S----------N-----E-NA-N--SPAAR-LNRFK-NKGKDST----EMRRRRIEVNVELRKAKKDDQMLKR-RNVSS-FPD-DA--
A0A6P5LDL0 : -------MS--T----------N-----E-NA-N--SPAAR-LNRFK-NKGKDST----EMRRRRIEVNVELRKAKKDDQMLKR-RNVST-FPD-DA--
A0A4X2LEJ2 : ---LNTTMS--T----------N-----E-NA-N--SLAAR-LNRFK-NKGKDST----EMRRRRIEVNVELRKAKKDDQMLKR-RNVST-FPD-DA--
F7D853     : -------MS--T----------N-----E-NA-N--SPAAR-LNRFK-NKGKDST----EMRRRRIEVNVELRKAKKDDQMLKR-RNVST-FPD-DA--
F6RK29     : -------MS--T----------N-----E-NA-N--SPAAR-LNRFK-NKGKDSS----EMRRRRIEVNVELRKAKKDDQMLKR-RNVST-FPD-DA--
Q6P6T9     : -------MS--T----------N-----E-NA-N--STAAR-LNRFK-NKGKDST----EMRRRRIEVNVELRKAKKDDQMLKR-RNVSS-FPD-DA--
A0A452G8T5 : -------MS--T----------N-----E-NA-N--SPAPR-LHRFK-NKGKDST----EMRRRRIEVNVELRKAKKDDQMLKR-RNVSS-FPD-DA--
W5Q181     : -------MS--T----------N-----E-NA-N--SPAPR-LHRFK-NKGKDST----EMRRRRIEVNVELRKAKKDDQMLKR-RNVSS-FPD-DA--
A0A6P5DHY5 : -------MS--T----------N-----E-NA-N--SPAPR-LNRFK-NKGKDST----EMRRRRIEVNVELRKAKKDDQMLKR-RNVSS-FPD-DA--
A0A6J0YM57 : -------MS--T----------N-----E-NA-N--SPAPR-LNRFK-NKGKDST----EMRRRRIEVNVELRKAKKDDQMLKR-RNVSS-FPD-DA--
Q3SYV6     : -------MS--T----------N-----E-NA-N--SPAPR-LNRFK-NKGKDST----EMRRRRIEVNVELRKAKKDDQMLKR-RNVSS-FPD-DA--
A0A6P3IU58 : -------MS--T----------N-----E-NA-N--SPAPR-LNRFK-NKGKDST----EMRRRRIEVNVELRKAKKDDQMLKR-RNVSS-FPD-DA--
A0A4W2E8W0 : -------MS--T----------N-----E-NA-N--SPAPR-LNRFK-NKGKDST----EMRRRRIEVNVELRKAKKDDQMLKR-RNVSS-FPD-DA--
A6PW68     : ---VPAAMS--T----------N-----E-NA-N--LPAAR-LNRFK-NKGKDST----EMRRRRIEVNVELRKAKKDEQMLKR-RNVSS-FPD-DA--
A0A6P5QND8 : -------MS--T----------N-----E-NA-N--LPAAR-LNRFK-NKGKDST----EMRRRRIEVNVELRKAKKDEQMLKR-RNVSS-FPD-DA--
A0A6J2MAG7 : -------MS--T----------N-----E-NA-N--SPAAR-LNRFK-NKGKDST----EMRRRRIEVNVELRKAKKDEQMLKR-RNVSS-FPD-DA--
J3KS65     : -------MS--T----------N-----E-NA-N--TPAAR-LHRFK-NKGKDST----EMRRRRIEVNVELRKAKKDDQMLKR-RNVSS-FPD-DA--
A0A2I3H982 : -------MS--T----------N-----E-NA-N--TPAAR-LHRFK-NKGKDST----EMRRRRIEVNVELRKAKKDDQMLKR-RNVSS-FPD-DA--
A0A2I2ZHY4 : -------MS--T----------N-----E-NA-N--TPAAR-LHRFK-NKGKDST----EMRRRRIEVNVELRKAKKDDQMLKR-RNVSS-FPD-DA--
F6VLU3     : -------MS--T----------N-----E-NA-N--TPAAR-LHRFK-NKGKDST----EMRRRRIEVNVELRKAKKDDQMLKR-RNVSS-FPD-DA--
K7C276     : -------MS--T----------N-----E-NA-N--TPAAR-LHRFK-NKGKDST----EMRRRRIEVNVELRKAKKDDQMLKR-RNVSS-FPD-DA--
A0A2R9CTC7 : -------MS--T----------N-----E-NA-N--TPAAR-LHRFK-NKGKDST----EMRRRRIEVNVELRKAKKDDQMLKR-RNVSS-FPD-DA--
A0A0D9QW15 : -------MS--T----------N-----E-NA-N--TPAAR-LHRFK-NKGKDST----EMRRRRIEVNVELRKAKKDDQMLKR-RNVSS-FPD-DA--
H2NUI7     : -------MS--T----------N-----E-NA-N--TPAAR-LHRFK-NKGKDST----EMRRRRIEVNVELRKAKKDEQMLKR-RNVSS-FPD-DA--
A0A6J3EXS8 : -------MS--T----------N-----E-NA-H--TPAAR-LNRFK-NKGKDST----EMRRRRIEVNVELRKAKKDDQMLKR-RNVSS-FPD-DA--
U3EQN0     : -------MS--T----------N-----E-NA-H--TPAAR-LNRFK-NKGKDST----EMRRRRIEVNVELRKAKKDDQMLKR-RNVSS-FPD-DA--
U3EY78     : -------MS--T----------N-----E-NA-N--TPDAR-PNRFK-NRGKDST----EMRCHRTEVNVELRKAKKVDQMLNR-RNISS-FPD-GA--
A0A674H5U9 : ---LTSKMS--T----------N-----E-NA-N--TPSAR-LNRFK-NKGKDST----EMRRRRIEVNVELRKAKKDDQMLKR-RNVST-LPD-DA--
U3JI37     : ---LTSKMS--T----------N-----E-NA-N--TPSAR-LNRFK-NKGKDST----EMRRRRIEVNVELRKAKKDDQMLKR-RNVST-LPD-DA--
A0A6J0GYH6 : --NLASKMS--T----------N-----E-NA-N--TP-AR-LNRFK-NKGKDST----EMRRRRIEVNVELRKAKKDDQMLKR-RNVST-LPD-DA--
```

```
                      *        20         *        40         *        60         *        80         *
A0A6J2IZT9 : --NLASKMS--T----------N-----E-NA-N--TP-AR-LNRFK-NKGKDST----EMRRRRIEVNVELRKAKKDDQMLKR-RNVST-LPD-DA--
A0A669Q9B1 : -------MS--S----------N-----E-NT-N--TA-AR-LNRFK-NKGKDTT----EMRRRRIEVNVELRKAKKDDQMLKR-RNVST-FPD-DA--
G1N6L6     : -------MS--S----------N-----E-NT-N--TP-AR-LNRFK-NKGKDTT----EMRRRRIEVNVELRKAKKDDQMLKR-RNVST-FPD-DA--
A0A1L1RX89 : -------MS--T----------N-----E-NT-N--TA-AR-LNRFK-NKGKDTT----EMRRRRIEVNVELRKAKKDDQMLKR-RNVST-FPD-DA--
A0A674IKE5 : -------MS--T----------N-----E-NA-S--AAVAR-LNRFK-NKGKDTT----EMRRRRIEVNVELRKAKKDDQMLKR-RNVST-LPD-DA--
A0A452I6U0 : -------MS--T----------N-----E-NA-S--AAVAR-LNRFK-NKGKDTT----EMRRRRIEVNVELRKAKKDDQMLKR-RNVST-LPD-DA--
K7FQJ3     : -------MS--T----------N-----E-NA-C--AAVAR-LNRFK-NKGKDTT----EMRRRRIEVNVELRKAKKDDQMLKR-RNVSI-LPD-DA--
U3IRY8     : -------MS--T----------N-----E-NA-N--AAAAR-LNRFK-NKGKDTT----EMRRRRIEVNVELRKAKKDDQMLKR-RNVST-LPD-DA--
A0A6J3E0U5 : -------MS--T----------N-----E-NA-N--AAAAR-LNRFK-NKGKDTT----EMRRRRIEVNVELRKAKKDDQMLKR-RNVST-LPD-DA--
A0A091QRY2 : -------MS--T----------N-----E-NA-N--TAAAR-LNRFK-NKGKDTT----EMRRRRIEVNVELRKAKKDDQMLKR-RNVST-LPD-DA--
A0A663E939 : -------MS--T----------N-----E-NA-N--TAAAR-LNRFK-NKGKDTT----EMRRRRIEVNVELRKAKKDDQMLKR-RNVST-LPD-DA--
A0A663M287 : -------MS--T----------N-----E-NA-N--TPAAR-LNRFK-NKGKDTT----EMRRRRIEVNVELRKAKKDDQMFKR-RNVNT-LPD-DA--
H0Z2A4     : -------MS--T----------N-----E-NA-N--TPSAR-LNRFK-NKGKDSTVSTLEMRRRRIEVNVELRKAKKDDQMLKR-RNVST-LPD-DA--
A0A672VFD3 : -------MS--T----------N-----E-NA-N--TPSAR-LNRFK-NKGKDAT----EMRRRRIEVNVELRKAKKDDQMLKR-RNVST-FPD-DA--
H9G3N8     : -------MS--S----------N-----E-NA-G--GTAVR-LNRFK-NKGKDSS----EMRRRRVEVSVELRKARKDDQMLKR-RNVGT-FPD-DA--
A0A6J0SWD6 : -------MS--T----------N-----E-NA-G--GAAVR-LNRFK-NKGKDSN----EMRRRRMEVNVELRKAKKDDQLLKR-RNVGT-FPD-DA--
A0A6I9YEM7 : -------MS--T----------N-----E-NA-G--AMAAR-LNRFK-NKGKDSN----EMRRRRMEVNVELRKAKKDDQMLKR-RNVGT-FPD-DA--
A0A6J1UUA8 : -------MS--T----------N-----E-NA-G--AMAAR-LNRFK-NKGKDSN----EMRRRRMEVNVELRKAKKDDQMLKR-RNVGT-FPD-DA--
A0A670YIX9 : -------MS--T----------N-----E-NA-A--AMAAR-LNRFK-NKGKDSN----EMRRRRMEVNVELRKAKKDDQMLKR-RNVGT-FPD-DA--
A0A6P9BCB7 : -------MS--T----------N-----E-NA-G--AMAAR-LNRFK-NKGKDSN----EMRRRRVEVNVELRKAKKDDQMLKR-RNVGT-FPD-DA--
V8PBZ1     : ---ITVAMS--T----------N-----E-NA-G--AMAAR-LNRFK-NKGKDSN----EMRRRRMEVNVELRKAKKDDQMLKR-RNVAVHWSV-E---
A0A670HXI0 : -------MS--A----------N-----E-N-------AAR-LNRFK-NKGKDSN----EMRRRRVEVNVELRKAKKDDQMLKR-RNVST-FPD-DA--
Q7ZX00     : -------MS--N----------N-----E-NG-------AR-LTRFK-NKGKDST----EMRRRRIEVNVELRKAKKDDQMLKR-RNVST-FPD-EP--
Q7ZYI0     : -------MS--S----------N-----E-NG-------AR-LTRFK-NKGKDST----EMRRRRIEVNVELRKAKKDDQLLKR-RNVST-FPD-EP--
Q66IF6     : -------MS--N----------N-----E-NA-------AR-LTRFK-NKGKDTT----EMRRRRIEVNVELRKAKKDDQMLKR-RNVST-FPD-EP--
A0A6P7YMD2 : -------MS--I----------N-----E----N----AAR-LNRFK-NKGKDSS----EMRRRRIEVNVELRKAKKDDQMFKK-RNVSS-FPD-EA--
A0A6P8SSA5 : -------MS--V----------S-----E-S--N--S-AAR-LNRFK-NKGKDSS----EMRRRRIEVNVELRKAKKDDQMFKK-RNVSS-FPD-EA--
A0A673BHL1 : -------MS-------------------E-NA-------AR-LNKFK-NKGKDVG----ECRRRRVEVNVELRKAKKDDLMCKR-RNVNS-CPD-EP--
A0A673BLF7 : -------VS--S----------------T-NA-------AR-LNKFK-NKGKDVG----ECRRRRVEVNVELRKAKKDDLMCKR-RNVNS-CPD-EP--
H3B358     : ----HTKMS--S----------------E-NA-S-----VR-LNRFK-NKGKDTT----EMRRRRFEVNVELRKAKKDEQIFKR-RNVNS-FPG-EA--
A0A1A8QNZ3 : ---------------------MS-----E-NV-A------R-LNKFK-NKGKDVN----ELRRRRAEVNVELRKAKKDDQIFKR-RNVCT-EPE-EA--
A0A1A8DGG9 : ---------------------MS-----E-NV-A------R-LNKFK-NKGKDVN----ELRRRRAEVNVELRKAKKDDQIFKR-RNVCT-EPE-EA--
A0A1A8PWJ9 : ---------------------MS-----E-NV-A------R-LNKFK-NKGKDVN----ELRRRRAEVNVELRKAKKDDQIFKR-RNVCT-EPE-EA--
A0A1A8IJ41 : ---------------------MS-----E-NV-A------R-LNKFK-NKGKDVN----ELRRRRAEVNVELRKAKKDDQIFKR-RNVCT-EPE-EA--
A0A1A8U0E9 : ---------------------MS-----E-NV-A------R-LNKFK-NKGKDVN----ELRRRRAEVNVELRKAKKDDQIFKR-RNVCT-EPE-EA--
A0A1A8GM44 : ---------------------MA-----E-NV-A------R-LNKFK-NKGKDAN----ELRRRRAEVNVELRKAKKDDQIFKR-RNVCT-EPE-EA--
A0A1A7YRI1 : ---------------------MS-----E-NV-A------R-LNKFK-NKGKDAN----ELRRRRVEVNVELRKAKKDDQIFKR-RNVCT-EPE-EA--
A0A3B3YGX7 : ------LFS--ALPNHTHTHKMS-----E-NV-A------R-LNKFK-NKGRDAN----ELRRRRVEVNVELRKAKKDDQMFKR-RNVAA-APD-EA--
A0A4Z2FV23 : ---------------------MS-----E-NV-E------R-LNKFK-NKGKDAN----ELRRRRVEVNVELRKTKKDDQMFKR-RNVSA-CPD-EA--
A0A6P8W0W9 : -------MS--A-P-------SC-----E-NA-D------R-LNKFK-NKGKDAN----ELRRRRVEVNVELRKAKKDDQMFKK-RNVAV-FPD-EA--
A0A4W3KKI1 : -------MS--A---------AN-----E-NA-S--P---R-LTQFK-NKGRDAN----ELRRRRIESSVELRKAKKDDQLFKK-RNVST-FPD-EA--
A0A673MQV0 : -------M----------------------NS-P------R-LTQFK-NKGKDVT----ELRRRRIEVNVELRRAKKDEQILKR-RNVSS-LPD-EA--
A0A671PG44 : -------MA--IC-----------------NS-P------R-LTQFK-NKGKDVT----ELRRRRIEVNVELRKAKKDEQILKR-RNVSS-LPD-EA--
A0A673MIX7 : -------MS--AA---------N-----E-NS-P------R-LTQFK-NKGKDVT----ELRRRRIEVNVELRRAKKDEQILKR-RNVSS-LPD-EA--
A0A671P8Z3 : -------MS--AA---------N-----E-NS-P------R-LTQFK-NKGKDVT----ELRRRRIEVNVELRKAKKDEQILKR-RNVSS-LPD-EA--
A0A672T5X6 : -------MS--AA---------N-----E-NS-P------R-LTQFK-NKGRDAT----ELRRRRVEVNVELRKAKKDEQILKR-RNVSS-LPD-EA--
Q6DI01     : -------MS--AA---------N-----E-NT-A------R-LTQFK-NKGKDAT----ELRRRRIEVNVELRKAKKDEQILKR-RNVSS-FPD-EA--
A0A3B3RXW3 : -------MS--SAG--------N-----E-NT-V------R-ISKFK-NKGKDAS----ELRRRRIEVNVELRKAKKDEQILKR-RNVSS-FPE-EA--
A0A6I8QW41 : -------M--------------G---QWH-SL-PWVWAFGS-LWRFF--PWQ-------EMRRRRIEVNVELRKAKKDDQMLKR-RNVST-FPD-EP--
A0A3B3RWN1 : -------MS--LLA----CVLYGGR-KWK-SL-T-VVLFNV-LVYFFLSTSK-------ELRRRRIEVNVELRKAKKDEQILKR-RNVSS-FPE-EA--
A0A2B4RY91 : -------MS-------------------E-N---------R-LRNFK-NKGKDTG----ELRRRRTDVTIELRKSKRDDQILKR-RNISQ---D-QSNS
A0A6A4VAT7 : -AGNCATMS--A----------D-----D-RS-S---A-AR-MRNFK-NRGKDSD----ELRRRRQETSVELRKAKKDDQILKR-RNIC--LTE-EDT-
P52171     : -P---TTNE--A----------D-----E-R-----------MRKFK-NKGKDTA----ELRRRRVEVSVELRKAKKDEQILKR-RNVC--FPE-ELI-
A0A3P8VN82 : -M--LSRMS--AH--------SNI---AD-R-----------IANFK-NNGKDST----EMRRRRSKVVVDLRKARKDDQILKR-RNGCS-HME-ES--
A0A2J7Q2F0 : -------MP--S--------QENY---VSGRQ-N------R-LTSFK-NKGKDAD----EMRRRRKEVSVELRKARKDDQLLKR-RNLNA--ED-DCT-
A0A2P8XZQ8 : -------MP--S--------QEN-----E-RP-N------R-LTSFK-NKGKNAD----EMRRRRNEVSVELRKARKDDQLLKR-RNLNS--ED-DSS-
A0A4Y2GZE4 : ------KLD-FS---------RT-----H-RF-T-MYAQSR-LTKFK-NKGRDQE----AMRRRRTEVNVELRKQKKDDNLSKR-RNVDA-FDD-EP--
A0A0A9Y1I6 : -------MP--S----------R-----E-N--D--VPTHR-LRAYK-NQGKDSE----EMRRRRNEVNVELRKAKKDEQLFKK-RNINW-VED-QV--
T2MFS8     : -------MP--S----------Q-----E-NV--------R-SREFK-NRGKDIS----DFRRRKNDVTVEIRKARKDEQMLKK-RNVGD--LS-EIS-
A0A2H8THT7 : -------MP--V----------K-----K-SIHN---S--RSFDVYQ-DRNKYYE----DLRRKRNDVTIELRKTLRDEQ-LKKHRNIV---LDANND-
A0A2S2NCU0 : -------MP--V----------K-----K-TILN---S--RSFDVYQ-DRNKYYE----DLRRKRNDVTIELRKTLRDEQ-LKKHRNIV---LDAHND-
cIBB2      : -------MS--T----------N-----E-NA-N---P-AR-LNRFK-NKGKDST----EMRRRRIEVNVELRKAKKDDQMLKR-RNVSS-FPD-DA--
```

```
                       *        20         *        40         *        60         *        80         *       100
A0A2S2R6K7 : --------MSDD--K---------VIG--HRLQ--S-F--K-----HRGKT---E---IERRRIRSEVT---VGIR--KNKQEQ---EL---MKRRNV--
A0A2S2NHS8 : --------MSDD--K---------VTG--HRLQ--S-F--K-----HRGKT---E---IERRRIRSEAT---VGLR--KNKQEQ---EL---MKRRNV--
G7YSA1     : --------MFDS--K---------ES----RLT--S-F--K-----NAGKSAD-E---M-RRR-RQEGQ---VELR--KNKREE---TL---QKKRNIPG
A0A1A8VCM4 : --------MADN--E------K--LDN--QRLK--N-F--K-----NKGRDL--E---T-MRRQRTEVV---VELR--KNKRDE---HL---LKRRNVPH
A0A1A8PEW1 : --------MADN--E------K--LDN--QRLK--N-F--K-----NKGRDL--E---T-MRRQRTEVV---VELR--KNKRDE---HL---LKRRNVPH
A0A1A8ILH3 : --------MADN--E------K--LDN--QRLK--N-F--K-----NKGRDL--E---T-MRRQRTEVV---VELR--KNKRDE---HL---LKRRNVPH
A0A1A8F687 : --------MADN--E------K--LDN--QRLK--N-F--K-----NKGRDL--E---T-MRRQRTEVV---VELR--KNKRDE---HL---LKRRNVPH
A0A1A8Q8V8 : --------MADN--------AG--LEN--HRIK--S-F--K-----NKGRDV--E---T-MRRHRNEVT---VELR--KNKRDE---HL---LKKRNVPQ
A0A6I8QJZ7 : WCRTAA--MADN--------AG--LEN--HRIK--S-F--K-----NKGRDV--E---T-MRRHRNEVT---VELR--KNKRDE---HL---LKKRNVPQ
A0A1A7ZEA1 : --------MADN--------AG--LEN--HRIK--S-F--K-----NKGRDV--E---T-MRRHRNEVT---VELR--KNKRDE---HL---LKKRNVPQ
A0A1A8LI71 : --------MADN--------AG--LEN--HRIK--S-F--K-----NKGRDV--E---T-MRRHRNEVT---VELR--KNKRDE---HL---LKKRNVPQ
A0A1A8JPR2 : --------MADN--------AG--LEN--HRIK--S-F--K-----NKGRDV--E---T-MRRHRNEVT---VELR--KNKRDE---HL---LKKRNVPQ
A0A1A7X5R1 : --------MADN--------AG--LEN--HRIK--S-F--K-----NKGRDV--E---T-MRRHRNEVT---VELR--KNKRDE---HL---LKKRNVPQ
A0A1A8BPJ4 : --------MADN--------AG--LEN--HRIK--S-F--K-----NKGRDV--E---T-MRRHRNEVT---VELR--KNKRDE---HL---LKKRNVPQ
A0A2I4BHC9 : --------MADN--------AG--LEN--HRIK--S-F--K-----NKGRDV--E---T-MRRHRNEVT---VELR--KNKRDE---HL---LKKRNVPQ
Q70PC6     : --------MADN--------AG--LDN--HRIK--S-F--K-----NKGRDV--E---T-MRRHRNEVT---VELR--KNKRDE---HL---LKKRNVPQ
A0A6P9CHE3 : --------MADNA-A-----AG--LEN--HRFK--S-F--K-----NKGRDV--E---T-MRRHRNEVT---IELR--KNKRDE---HL---LKKRNVPQ
A0A6J1V4X7 : --------MADNA-A-----AG--LEN--HRFK--S-F--K-----NKGRDV--E---T-MRRHRNEVT---IELR--KNKRDE---HL---LKKRNVPQ
A0A6I9X8Y6 : --------MADNA-A-----AG--LEN--HRFK--S-F--K-----NKGRDV--E---T-MRRHRNEVT---IELR--KNKRDE---HL---LKKRNVPQ
G1KPC3     : --NAAA--AAAAA-A-----AA--AEN--HRLK--S-F--K-----NKGRDV--E---T-MRRHRNEVT---IELR--KNKRDE---HL---LKKRNVPQ
A0A6J3DF17 : --------MTDNA-A-----AG--LEN--HRIK--S-F--K-----NKGRDV--E---T-MRRHRNEVT---IELR--KNKRDE---HL---LKKRNVPQ
U3IJT7     : --------MTDNA-A-----AG--LEN--HRIK--S-F--K-----NKGRDV--E---T-MRRHRNEVT---VELR--KNKRDE---HL---LKKRNVPQ
A0A6J2RYW0 : --------MAEN--A------G--LEN--HRIK--S-F--K-----NKGRDV--E---T-MRRHRNEVT---VELR--KNKRDE---HL---LKKRNVPQ
A0A667Z8B4 : --PKPCTAMAEN--A------G--LEN--HRIK--S-F--K-----NKGRDV--E---T-MRRHRNEVT---VELR--KNKRDE---HL---LKKRNVPQ
H3AV75     : --------MAEN--A------G--LEN--HRIK--S-F--K-----NKGRDV--E---T-MRRHRNEVT---VELR--KNKRDE---HL---LKKRNVPQ
A0A6P3WEA8 : --------MAEN--A------G--LEN--HRIK--S-F--K-----NKGRDV--E---T-MRRHRNEVT---VELR--KNKRDE---HL---LKKRNVPQ
A0A671MPH1 : --------MAEN--A------G--LEN--HRIK--S-F--K-----NKGRDV--E---T-MRRHRNEVT---VELR--KNKRDE---HL---LKKRNVPQ
A0A672PQJ6 : --------MAEN--A------G--LEN--HRIK--S-F--K-----NKGRDV--E---T-MRRHRNEVT---VELR--KNKRDE---HL---LKKRNVPQ
A0A6P7HJQ6 : --------MAEN--A------G--LEN--HRIK--S-F--K-----NKGRDV--E---T-MRRHRNEVT---VELR--KNKRDE---HL---LKKRNVPQ
A0A672Z3A8 : --AKPHTAMAEN--A------G--LEN--HRIK--S-F--K-----NKGRDV--E---T-MRRHRNEVT---VELR--KNKRDE---HL---LKKRNVPQ
A0A665V3R5 : --------MAEN--A------G--LEN--HRIK--S-F--K-----NKGRDV--E---T-MRRHRNEVT---VELR--KNKRDE---HL---LKKRNVPQ
Q6IRB7     : --------MAEN--A------G--LEN--HRIK--S-F--K-----NKGRDV--E---T-MRRHRNEVT---VELR--KNKRDE---HL---LKKRNVPQ
A0A6P8RJ78 : --------MAEN--A------G--LEN--HRIK--S-F--K-----NKGRDV--E---T-MRRHRNEVT---VELR--KNKRDE---HL---LKKRNVPQ
A0A6J2VU04 : --------MAEN--A------G--LEN--HRIK--S-F--K-----NKGRDV--E---T-MRRHRNEVT---VELR--KNKRDE---HL---LKKRNVPQ
A0A4W4HIJ1 : --QRENTAMAEN--A------G--LEN--HRIK--S-F--K-----NKGRDV--E---T-MRRHRNEVT---VELR--KNKRDE---HL---LKKRNVPQ
W5UB95     : --------MAEN--A------G--LEN--HRIK--S-F--K-----NKGRDV--E---T-MRRHRNEVT---VELR--KNKRDE---HL---LKKRNVPQ
A0A6P7LAX0 : --------MAEN--A------G--LEN--HRIK--S-F--K-----NKGRDV--E---T-MRRHRNEVT---VELR--KNKRDE---HL---LKKRNVPQ
Q8JFT7     : --------MAEN--A------G--LEN--HRIK--S-F--K-----NKGRDV--E---T-MRRHRNEVT---VELR--KNKRDE---HL---LKKRNVPQ
A0A672HEZ1 : --------MAEN--A------G--LEN--HRIK--S-F--K-----NKGRDV--E---T-MRRHRNEVT---VELR--KNKRDE---HL---LKKRNVPQ
A0A4Z2GJN9 : --------MAEN--A------G--LEN--HRIK--S-F--K-----NKGRDV--E---T-MRRHRNEVT---VELR--KNKRDE---HL---LKKRNVPQ
H0V3K6     : --------MAEN--P------G--LEN--HRIK--S-F--K-----NK----------T-MRRHRNEVT---VELR--KNKRDE---HL---LKKRNVPQ
A0A3Q7UKJ0 : --------MAEN--P------G--LEN--HRIK--S-F--K-----NKGRDV--E---T-MRRHRNEVT---VELR--KNKRDE---HL---LKKRNVPQ
A0A6P4VPZ6 : --------MAEN--P------G--LEN--HRIK--S-F--K-----NKGRDV--E---T-MRRHRNEVT---VELR--KNKRDE---HL---LKKRNVPQ
A0A2U3VVQ1 : --------MAEN--P------G--LEN--HRIK--S-F--K-----NKGRDV--E---T-MRRHRNEVT---VELR--KNKRDE---HL---LKKRNVPQ
A0A2Y9NMF0 : --------MAEN--P------G--LEN--HRIK--S-F--K-----NKGRDV--E---T-MRRHRNEVT---VELR--KNKRDE---HL---LKKRNVPQ
A0A3Q7PA97 : --------MAEN--P------G--LEN--HRIK--S-F--K-----NKGRDV--E---T-MRRHRNEVT---VELR--KNKRDE---HL---LKKRNVPQ
A0A6J2CSB0 : --------MAEN--P------G--LEN--HRIK--S-F--K-----NKGRDV--E---T-MRRHRNEVT---VELR--KNKRDE---HL---LKKRNVPQ
A0A2U4BGI2 : --------MAEN--P------G--LEN--HRIK--S-F--K-----NKGRDV--E---T-MRRHRNEVT---VELR--KNKRDE---HL---LKKRNVPQ
A0A2Y9EMI4 : --------MAEN--P------G--LEN--HRIK--S-F--K-----NKGRDV--E---T-MRRHRNEVT---VELR--KNKRDE---HL---LKKRNVPQ
A0A6J1ZVC0 : --------MAEN--P------G--LEN--HRIK--S-F--K-----NKGRDV--E---T-MRRHRNEVT---VELR--KNKRDE---HL---LKKRNVPQ
M3WQJ0     : --------MAEN--P------G--LEN--HRIK--S-F--K-----NKGRDV--E---T-MRRHRNEVT---VELR--KNKRDE---HL---LKKRNVPQ
A0A2Y9H0K2 : --------MAEN--P------G--LEN--HRIK--S-F--K-----NKGRDV--E---T-MRRHRNEVT---VELR--KNKRDE---HL---LKKRNVPQ
A0A4W2E6Z1 : --------MAEN--P------G--LEN--HRIK--S-F--K-----NKGRDV--E---T-MRRHRNEVT---VELR--KNKRDE---HL---LKKRNVPQ
A0A6P3YND9 : --------MAEN--P------G--LEN--HRIK--S-F--K-----NKGRDV--E---T-MRRHRNEVT---VELR--KNKRDE---HL---LKKRNVPQ
A0A3Q1MUS6 : --------MAEN--P------G--LEN--HRIK--S-F--K-----NKGRDV--E---T-MRRHRNEVT---VELR--KNKRDE---HL---LKKRNVPQ
A0A667GNB9 : --------MAEN--P------G--LEN--HRIK--S-F--K-----NKGRDV--E---T-MRRHRNEVT---VELR--KNKRDE---HL---LKKRNVPQ
A0A6P5CMI9 : --------MAEN--P------G--LEN--HRIK--S-F--K-----NKGRDV--E---T-MRRHRNEVT---VELR--KNKRDE---HL---LKKRNVPQ
A0A5G2R9E6 : --------MAEN--P------G--LEN--HRIK--S-F--K-----NKGRDV--E---T-MRRHRNEVT---VELR--KNKRDE---HL---LKKRNVPQ
A0A452FEC0 : --------MAEN--P------G--LEN--HRIK--S-F--K-----NKGRDV--E---T-MRRHRNEVT---VELR--KNKRDE---HL---LKKRNVPQ
A0A1S3AHI3 : --------MAEN--P------G--LEN--HRIK--S-F--K-----NKGRDV--E---T-MRRHRNEVT---VELR--KNKRDE---HL---LKKRNVPQ
A0A0P6JDA1 : --------MAEN--P------G--LEN--HRIK--S-F--K-----NKGRDV--E---T-MRRHRNEVT---VELR--KNKRDE---HL---LKKRNVPQ
A0A6P6EKZ5 : --------MAEN--P------G--LEN--HRIK--S-F--K-----NKGRDV--E---T-MRRHRNEVT---VELR--KNKRDE---HL---LKKRNVPQ
A0A250Y2A9 : --------MAEN--P------G--LEN--HRIK--S-F--K-----NKGRDV--E---T-MRRHRNEVT---VELR--KNKRDE---HL---LKKRNVPQ
```

```
                    *        20         *        40         *        60         *        80         *       100
Q9CT07      : --------MAEN--P------G--LEN--HRIK--S-F--K-----NKGRDV--E---T-MRRHRNEVT---VELR--KNKRDE---HL---LKKRNVPQ
A0A6P5QYT5  : --------MAEN--P------G--LEN--HRIK--S-F--K-----NKGRDV--E---T-MRRHRNEVT---VELR--KNKRDE---HL---LKKRNVPQ
A0A3Q0D085  : --------MAEN--P------G--LEN--HRIK--S-F--K-----NKGRDV--E---T-MRRHRNEVT---VELR--KNKRDE---HL---LKKRNVPQ
Q56R18      : --------MAEN--P------G--LEN--HRIK--S-F--K-----NKGRDV--E---T-MRRHRNEVT---VELR--KNKRDE---HL---LKKRNVPQ
A0A2K6R1J6  : --------MAEN--P------S--LEN--HRIK--S-F--K-----NKGRDV--E---T-MRRHRNEVT---VELR--KNKRDE---HL---LKKRNVPQ
A0A2K5RYQ6  : --------MAEN--P------S--LEN--HRIK--S-F--K-----NKGRDV--E---T-MRRHRNEVT---VELR--KNKRDE---HL---LKKRNVPQ
A0A5F8A5V2  : --------MAEN--P------S--LEN--HRIK--S-F--K-----NKGRDV--E---T-MRRHRNEVT---VELR--KNKRDE---HL---LKKRNVPQ
H0WKJ7      : --------MAEN--P------S--LEN--HRIK--S-F--K-----NKGRDV--E---T-MRRHRNEVT---VELR--KNKRDE---HL---LKKRNVPQ
A0A2R9AVB4  : --------MAEN--P------S--LEN--HRIK--S-F--K-----NKGRDV--E---T-MRRHRNEVT---VELR--KNKRDE---HL---LKKRNVPQ
H2Q7K7      : --------MAEN--P------S--LEN--HRIK--S-F--K-----NKGRDV--E---T-MRRHRNEVT---VELR--KNKRDE---HL---LKKRNVPQ
I7GIR2      : --------MAEN--P------S--LEN--HRIK--S-F--K-----NKGRDV--E---T-MRRHRNEVT---VELR--KNKRDE---HL---LKKRNVPQ
U3BKZ4      : --------MAEN--P------S--LEN--HRIK--S-F--K-----NKGRDV--E---T-MRRHRNEVT---VELR--KNKRDE---HL---LKKRNVPQ
A0A0D9RYP1  : --------MAEN--P------S--LEN--HRIK--S-F--K-----NKGRDV--E---T-MRRHRNEVT---VELR--KNKRDE---HL---LKKRNVPQ
A0A2K5DTK7  : --------MAEN--P------S--LEN--HRIK--S-F--K-----NKGRDV--E---T-MRRHRNEVT---VELR--KNKRDE---HL---LKKRNVPQ
A0A096NP23  : --------MAEN--P------S--LEN--HRIK--S-F--K-----NKGRDV--E---T-MRRHRNEVT---VELR--KNKRDE---HL---LKKRNVPQ
A0A024RDV7  : --------MAEN--P------S--LEN--HRIK--S-F--K-----NKGRDV--E---T-MRRHRNEVT---VELR--KNKRDE---HL---LKKRNVPQ
A0A2K6FG80  : --------MAEN--P------S--LEN--HRIK--S-F--K-----NKGRDV--E---T-MRRHRNEVT---VELR--KNKRDE---HL---LKKRNVPQ
A0A6J3J6G8  : --------MAEN--P------S--LEN--HRIK--S-F--K-----NKGRDV--E---T-MRRHRNEVT---VELR--KNKRDE---HL---LKKRNVPQ
A0A2K6DCA2  : --------MAEN--P------S--LEN--HRIK--S-F--K-----NKGRDV--E---T-MRRHRNEVT---VELR--KNKRDE---HL---LKKRNVPQ
A0A2K6MWD3  : --------MAEN--P------S--LEN--HRIK--S-F--K-----NKGRDV--E---T-MRRHRNEVT---VELR--KNKRDE---HI---LKKRNVHQ
A0A674N306  : --------MAEN--A------G--LEN--HRIK--S-F--K-----NKGRDV--E---V-SKR-AQAP----V-LDGPNNKRDE---HL---LKKRNVPL
A0A672HF09  : --------MAEN--A------G--LEN--HRIK--S-F--K-----NKGRDV--E---V-SGR-KAAA----V-----RNKRDE---HL---LKKRNVPQ
A0A670I6K3  : --NAAAAAAAAA--A------G--LEN--HRFK--S-F--K-----NKGRDV--E---VGAGRHSNALSWLFV-L---QNKRDE---HL---LKKRNVPQ
A0A2K5MEL9  : --------MAEN--P------S--LEN--HRIK--S-F--K-----NKGRDV--EG--V-S-RHRNFS----------ENKRDE---HL---LKKRNVPQ
A0A674HEZ2  : --------MAEN--A---AATG--LES--HRIK--S-F--K-----NKGRDA--E---T-MRRHRNEVT---IELR--KNKRDE---HL---LKKRNVPQ
A0A218UQY7  : --------MAEN--A---AATG--LES--HRIK--S-F--K-----NKGRDA--EV-RT-MRRHRNEVT---IELR--KNKRDE---HL---LKKRNVPQ
A0A6J0HA95  : --------MAEN--A---AA-G--LES--HRIK--S-F--K-----NKGRDA--E---T-MRRHRNEVT---IELR--KNKRDE---HL---LKKRNVPQ
A0A6J2GPV2  : --------MAEN--A---AA-G--LES--HRIK--S-F--K-----NKGRDA--E---T-MRRHRNEVT---IELR--KNKRDE---HL---LKKRNVPQ
A0A663F212  : --------MAENAAA---AAAG--LEN--HRIK--S-F--K-----NKGRDV--E---T-MRRHRNEVT---IELR--KNKRDE---HL---LKKRNVPQ
A0A672TI99  : --------MAENAAA---AAAG--LEN--HRIK--S-F--K-----NKGRDV--E---T-MRRHRNEVT---IELR--KNKRDE---HL---LKKRNVPQ
A0A669QP80  : --------MAEN-AA---AAAG--LEN--HRIK--S-F--K-----NKGRDV--E---T-MRRHRNEVT---IELR--KNKRDE---HL---LKKRNVPQ
F1NV68      : --------MAEN------AAAG--LEN--HRIK--S-F--K-----NKGRDV--E---T-MRRHRNEVT---IELR--KNKRDE---HL---LKKRNVPQ
A0A6P5KU41  : --------MAEN--------AG--LEN--HRIK--S-F--K-----NKGRDV--E---T-MRRHRNEVT---IELR--KNKRDE---HL---LKKRNVPQ
A0A6I8P1J4  : --------MAEN-------A-G--LEN--HRIK--S-F--K-----NKGRDV--E---T-MRRHRNEVT---IELR--KNKRDE---HL---LKKRNVPQ
A0A6P7Y0L1  : --------MAEN--A------G--LEN--HRIK--S-F--K-----NKGRDV--E---T-MRRHRNEVT---IELR--KNKRDE---HL---LKKRNVPQ
A0A452IFV5  : --------MAEN--AN--AAAG--LEN--HRIK--S-F--K-----NKGRDV--E---T-MRRHRNEVT---IELR--KNKRDE---HL---LKKRNVPQ
A0A5F8GJT3  : --------MAEN--A------G--LEN--HRIK--S-F--K-----NKGRDV--E---T-MRRHRNEVT---IELR--KNKRDE---HL---LKKRNVPQ
A0A4X2M2X5  : --------MAEN--A------G--LEN--HRIK--S-F--K-----NKGRDV--E---T-MRRHRNEVT---IELR--KNKRDE---HL---LKKRNVPQ
A0A6I9NPM0  : --------MAEN--A------G--LEN--HRIK--S-F--K-----NKGRDV--E---T-MRRHRNEVT---VELR--KNKRDE---HL---LKKRNVPL
A0A674NY42  : --RSEASAMAEN--A------G--LEN--HRIK--S-F--K-----NKGRDV--E---T-MRRHRNEVT---LELR--KNKRDE---HL---LKKRNVPL
A0A672Z3N3  : --------MAEN--A------G--LEN--HRIK--S-F--K-----NKGRDV--E---T-MRRHRNEVT---VELR--KVRKDSVVVYTSCSFNRQNVTL
A0A6P8UDL6  : --------MAEN--A------G--LEN--HRIK--S-F--K-----NKGRDV--E---T-MRRHRNEVT---VELR--KNKRDE---HL---LKKRNVPL
A0A4W5L687  : --------MAEN--A------D--LDN--HRIK--S-F--K-----NKGRDV--E---T-MRRHRNDVT---VELR--KNKRDK---HL---LKKRNVPQ
A0A4W5L543  : --------MAEN--A------G--LDN--HRIK--S-F--K-----NKGRDV--E---T-MRRHRNDVT---VELR--KNKRDE---HL---LKKRNVPQ
A0A4W5PPZ0  : --------MAEN--A------D--LDN--HRIK--S-F--K-----NKGRDV--E---T-MRRHRNEVT---VELR--KNKRDE---HL---LKKRNVPQ
A0A151NB90  : -----MQ-VGYR-WP------G--FNF-L-FLRGFSAF-PA-----KPHLPP--P--QT-MRRQRNEVV---VELR--KNKRDE---HL---LKRRNVPH
A0A1V4J4R5  : -----MD-LGAL-VPASPREQGKELKT-LAFQQAHSQLGPL------PRSDP--PFYQT-MRRHRNEVT---IELR--KNKRDE---HL---LKKRNVPQ
A0A670Y0Q9  : -----M--V-YK-YS-------KHTSA-LISVLHVSDF--I-------FL------FQT-MRRHRNEVT---IELR--KNKRDE---HL---LKKRNVPQ
A0A6I8PME1  : -----ML-LEAEDQI---------LSPKYHILNLPSSF-LI-----SVHHAF--T--ET-MRRHRNEVT---IELR--KNKRDE---HL---LKKRNVPQ
A0A6J3J7F8  : -----ML-LPAV-WIS--------LNP-VR-LT-PG----I-----N-HH-------KT-MRRHRNEVT---VELR--KNKRDE---HL---LKKRNVPQ
A0A5F9CSY8  : -----MEPRQCR-SP--------RLPP-ASRAQAGGGG-PQ-----R-GVGT--S--RT-MRRHRNEVT---VELR--KNKRDE---HL---LKKRNVPQ
A0A340Y4X1  : -----SPGLLCS-TP----------NPAASSTSSGSVLEPC-----LCALTTRGP--IT-MRRHRNEVT---VELR--KNKRDE---HL---LKKRNVPQ
Q71VM5      : --------MSVQ-KP---------LER--QKL-----F--K-----NKDKDT--E---S-LRQRRNEVS---VVLR--KAKRDE---AL---QKRRLVPN
A0A4C1V8D9  : --------MATD--Q---------VKN---RMQ--V-F--K-----NAGKDV--D---E-MRRRRNEVT---VELR--KNKREE---TL---QKRRNVPI
A0A0V1HFN1  : --HRLFFGFTMS--D------H--LNNT-RILH----F--K-----NKGRNAD-E-----LRRRRTDIS---FELR--KSKRED---TL---SKKRSL-H
A0A0V1N4W6  : --HRLFFGFTMS--D------H--LNNT-RILH----F--K-----NKGRNAD-E-----LRRRRTDIS---FELR--KSKRED---TL---SKKRSL-H
A0A0V1BD23  : ----LFFGFTMS--D------H--VNNT-RILH----F--K-----NKGRNAD-E-----LRRRRTDMS---FELR--KSKRED---TL---SKKRSI-H
A0A0V0TWQ0  : ----LFFGFTMS--D------H--VNNT-RILH----F--K-----NKGRNAD-E-----LRRRRTDMS---FELR--KSKRED---TL---SKKRSI-H
A0A0V1P443  : ----LFFGFTMS--D------H--VNNT-RILH----F--K-----NKGRNAD-E-----LRRRRTDMS---FELR--KSKRED---TL---SKKRSI-H
A0A0V1J5Y5  : --HRLFFGFTMS--D------H--LNNT-RILH----F--K-----NKGRNAD-E-----LRRRRTDIS---FELR--KSKRED---TL---SKKRSL-H
A0A0V1FJA8  : ----LFL-FSQT--C------T--VLN---FLH----FG-KELVVLNYEKSL--E-----LRRRRTDIS---FELR--KSKRED---TL---SKKRSL-H
cIBB3       : --------MAEN--P------G--LEN--HRIK--S-F--K-----NKGRDV--E---T-MRRHRNEVT---VELR--KNKRDE---HL---LKKRNVPQ
```

```
A0A2S2R6K7 : -GEV-MNED
A0A2S2NHS8 : -GEV-MNDD
G7YSA1     : LGEV-T---
A0A1A8VCM4 : -EDI-C-E-
A0A1A8PEW1 : -EDI-C-E-
A0A1A8ILH3 : -EDI-C-E-
A0A1A8F687 : -EDI-C-E-
A0A1A8Q8V8 : -EES-L-E-
A0A6I8QJZ7 : -EES-L-E-
A0A1A7ZEA1 : -EES-L-E-
A0A1A8LI71 : -EES-L-E-
A0A1A8JPR2 : -EES-L-E-
A0A1A7X5R1 : -EES-L-E-
A0A1A8BPJ4 : -EES-L-E-
A0A2I4BHC9 : -EES-L-E-
Q70PC6     : -EES-L-E-
A0A6P9CHE3 : -EES-L-E-
A0A6J1V4X7 : -EES-L-E-
A0A6I9X8Y6 : -EES-L-E-
G1KPC3     : -EEN-L-E-
A0A6J3DF17 : -EES-L-E-
U3IJT7     : -EES-L-E-
A0A6J2RYW0 : -EES-L-E-
A0A667Z8B4 : -EES-L-E-
H3AV75     : -EES-L-E-
A0A6P3WEA8 : -EES-L-E-
A0A671MPH1 : -EES-L-E-
A0A672PQJ6 : -EES-L-E-
A0A6P7HJQ6 : -EES-L-E-
A0A672Z3A8 : -EES-L-E-
A0A665V3R5 : -EES-L-E-
Q6IRB7     : -EES-L-E-
A0A6P8RJ78 : -EES-L-E-
A0A6J2VU04 : -EES-L-E-
A0A4W4HIJ1 : -DES-L-E-
W5UB95     : -DES-L-E-
A0A6P7LAX0 : -EES-L-E-
Q8JFT7     : -EES-L-E-
A0A672HEZ1 : -EES-L-E-
A0A4Z2GJN9 : -EES-L-E-
H0V3K6     : -EES-L-E-
A0A3Q7UKJ0 : -EES-L-E-
A0A6P4VPZ6 : -EES-L-E-
A0A2U3VVQ1 : -EES-L-E-
A0A2Y9NMF0 : -EES-L-E-
A0A3Q7PA97 : -EES-L-E-
A0A6J2CSB0 : -EES-L-E-
A0A2U4BGI2 : -EES-L-E-
A0A2Y9EMI4 : -EES-L-E-
A0A6J1ZVC0 : -EES-L-E-
M3WQJ0     : -EES-L-E-
A0A2Y9H0K2 : -EES-L-E-
A0A4W2E6Z1 : -EES-L-E-
A0A6P3YND9 : -EES-L-E-
A0A3Q1MUS6 : -EES-L-E-
A0A667GNB9 : -EES-L-E-
A0A6P5CMI9 : -EES-L-E-
A0A5G2R9E6 : -EES-L-E-
A0A452FEC0 : -EES-L-E-
A0A1S3AHI3 : -EES-L-E-
A0A0P6JDA1 : -EES-L-E-
A0A6P6EKZ5 : -EES-L-E-
A0A250Y2A9 : -EES-L-E-
Q9CT07     : -EES-L-E-
```

| | | |
|---|---|---|
| A0A6P5QYT5 | : | -EES-L-E- |
| A0A3Q0D085 | : | -EES-L-E- |
| Q56R18 | : | -EES-L-E- |
| A0A2K6R1J6 | : | -EES-L-E- |
| A0A2K5RYQ6 | : | -EES-L-E- |
| A0A5F8A5V2 | : | -EES-L-E- |
| H0WKJ7 | : | -EES-L-E- |
| A0A2R9AVB4 | : | -EES-L-E- |
| H2Q7K7 | : | -EES-L-E- |
| I7GIR2 | : | -EES-L-E- |
| U3BKZ4 | : | -EES-L-E- |
| A0A0D9RYP1 | : | -EES-L-E- |
| A0A2K5DTK7 | : | -EES-L-E- |
| A0A096NP23 | : | -EES-L-E- |
| A0A024RDV7 | : | -EES-L-E- |
| A0A2K6FG80 | : | -EES-L-E- |
| A0A6J3J6G8 | : | -EES-L-E- |
| A0A2K6DCA2 | : | -EES-L-E- |
| A0A2K6MWD3 | : | -EES-L-E- |
| A0A674N306 | : | -EES-L-E- |
| A0A672HF09 | : | -EES-L-E- |
| A0A670I6K3 | : | -EES-L-E- |
| A0A2K5MEL9 | : | -EES-L-E- |
| A0A674HEZ2 | : | -EES-L-E- |
| A0A218UQY7 | : | -EES-L-E- |
| A0A6J0HA95 | : | -EES-L-E- |
| A0A6J2GPV2 | : | -EES-L-E- |
| A0A663F212 | : | -EES-L-E- |
| A0A672TI99 | : | -EES-L-E- |
| A0A669QP80 | : | -EES-L-E- |
| F1NV68 | : | -EES-L-E- |
| A0A6P5KU41 | : | -EES-L-E- |
| A0A6I8P1J4 | : | -EES-L-E- |
| A0A6P7Y0L1 | : | -EES-L-E- |
| A0A452IFV5 | : | -EES-L-E- |
| A0A5F8GJT3 | : | -EES-L-E- |
| A0A4X2M2X5 | : | -EES-L-E- |
| A0A6I9NPM0 | : | -EES-L-E- |
| A0A674NY42 | : | -EES-L-E- |
| A0A672Z3N3 | : | --DAIL-Q- |
| A0A6P8UDL6 | : | -EDS-L-E- |
| A0A4W5L687 | : | -EDS-L-E- |
| A0A4W5L543 | : | -EDS-L-E- |
| A0A4W5PPZ0 | : | -EES-L-E- |
| A0A151NB90 | : | -EDI-C-E- |
| A0A1V4J4R5 | : | -EES-L-E- |
| A0A670Y0Q9 | : | -EES-L-E- |
| A0A6I8PME1 | : | -EES-L-E- |
| A0A6J3J7F8 | : | -EES-L-E- |
| A0A5F9CSY8 | : | -EES-L-E- |
| A0A340Y4X1 | : | -EES-L-E- |
| Q71VM5 | : | VEED---E- |
| A0A4C1V8D9 | : | -NDS-T-D- |
| A0A0V1HFN1 | : | -VQS-V-EI |
| A0A0V1N4W6 | : | -VQS-V-EI |
| A0A0V1BD23 | : | -VQS-V-EI |
| A0A0V0TWQ0 | : | -VQS-V-EI |
| A0A0V1P443 | : | -VQS-V-EI |
| A0A0V1J5Y5 | : | -VQN-V-EI |
| A0A0V1FJA8 | : | -VQN-V-EI |
| cIBB3 | : | -EES-L-E- |

```
                  *         20         *         40         *         60         *         80         *        100
H3AN54     : --------L---------------P-FS----NPL-LRD-S-IF---K-R-T------C---Q-------EY--ETMRRQRN-EVVVELRKNKRDEHLLK
A0A1U8DF45 : --------MQ---------VGYRWPGF-----NFLFLRGFS-AF-PTK-P-H------L---P-------PP--QTMRRQRN-EVVVELRKNKRDEHLLK
A0A4W3JN51 : --------MG------------RT-------------RKAP-LI-EAE-R-HE----LL---S-R----IAY-SASMRRQRN-EVVVELRKNKRVEHLFK
A0A6P7QP70 : ---SFYSSLR------------RDHQF-----AFL-HRKVQRVI-QS--C-HPPVFGIL---SLHLLVVVAVRNPTMRRQRN-EVVVELRKNKRDEHLLK
G1SG68     : --------ME---------N----------------MNNTG-KV-PI--------LMIL---TF-QLVRIS---QTMRRQRN-EVVVELRKNKRDEHLLK
A0A663DS72 : -----AIIVE---------NE---------------IN--G-KI-QVG-I-H---LRFL---SFGACV-LSF--KTMRRQRN-EVVVELRKNKRDEHLLK
A0A674PLW0 : --MNCLIIVT---------QG-----------HK-DGDNAP-FV-LCD-V-Y-----IL---SF------P---QTMRRQRT-EVVVELRKNKRDEHLLK
U3JFA4     : -------VWA---------Q------------LKFDLMNCK-FF-F-D-K-N-----ND---AW----------QTMRRQRN-EVVVELRKNKRDEHLLK
A0A673B6V3 : --------MS---------D-------------ALDC-NVK-IL-FCS--------------------------QTMRRQRT-EVVVELRKNKRDEHLLK
A0A6J1ZKW3 : ---MLTVPSS---------GLC----------FTLDIADSR--C-GSR-C-A------L----F------Q----TMRRQRN-EVVVELRKNKRDEHLLK
A0A6J2CG34 : ---MLTVPSS---------GLC----------FTLDIADSR--C-GSR-C-A------L----F------Q----TMRRQRN-EVVVELRKNKRDEHLLK
A0A6P4VIN5 : ---MLTVPSS---------GLC----------FTLDIADSR--C-GSR-C-A------L----F------Q----TMRRQRN-EVVVELRKNKRDEHLLK
A0A7E6D5F7 : ---MLTVPSC---------GLC----------FAMDIADSR--C-GSP-C-A------L----F------Q----TMRRQRN-EVVVELRKNKRDEHLLK
A0A6P6BYU9 : ----------------------------------MDIADSR--C-GSR-C-A------L----F------Q----TMRRQRN-EVVVELRKNKRDEHLLK
A0A3Q0GSB0 : -----T-PCS---------AVSR---------FGGPL-DGT--C--SD-C-F------L---NLKS---SQAS-ETMRRQRN-EVVVELRKNKRDEHLLK
A0A5F4W1H4 : -SWSLTMLLN---------PL-----------LALEIQDGP--F-RSS-S-G------L---QLTLK--VQR---TMRRQRN-EVVVELRKNKRDEHLLK
A0A673BC87 : --------------------------------VA----DS-----QSS-A-C------L---RYTT---VSRGWYTMRRQRT-EVVVELRKNKRDEHLLK
A0A6I9KX70 : SPPYLSSFTS---------PG-----------KRLPVHSS---I-GSP-KGYT----KL---SS------S---STMRRQRN-EVVVELRKNKRDEHLLK
A0A6J2QXP4 : --------MA---------DN-----------EKLDNQRLK-NF-KNK-G-R-----DL---E------------TMRRQRT-EVVVELRKNKRDEHLLK
A0A4Z2IS62 : --------MA---------DN-----------EKLDNQRLK-NF-KNK-G-R-----DL---E------------TMRRQRT-EVVVELRKNKRDEHLLK
A0A4W4FLE0 : --------MN---------DN-----------EKLDNQRLK-NF-KNK-G-R-----DL---E------------TMRRQRT-EVVVELRKNKRDEHLLK
W5UA83     : --------MN---------DN-----------EKLDNQRLK-NF-KNK-G-R-----DL---E------------TMRRQRT-EVVVELRKNKRDEHLLK
A0A6J2VJW1 : --------MT---------DN-----------DKLDNQRLK-NF-KNK-G-R-----DL---E------------TMRRQRT-EVVVELRKNKRDEHLLK
F1QEW3     : --------MS---------DS-----------EKLDKQRLK-NF-KNK-G-R-----DL---E------------TMRRQRT-EVVVELRKNKRDEHLLK
A0A6P3WAK5 : --------MA---------DN-----------EKLDNQRLK-NF-KNK-G-R-----DL---E------------TMRRQRT-EVVVELRKSKRDEHLLK
A0A4W6BU17 : --KKGTANMA---------DS-----------EKLDNQRLK-NF-KNK-G-R-----DL---E------------TMRRQRT-EVVVELRKNKRDEHLLK
A0A665UA99 : ---------S---------ERGTRKKRG----EKLDNQRLK-NF-KNK-G-R-----DL---E------------TMRRQRT-EVVVELRKNKRDEHLLK
A0A665UA62 : --------MA---------D--S---------EKLDNQRLK-NF-KNK-G-R-----DL---E------------TMRRQRT-EVVVELRKNKRDEHLLK
H2LP51     : --------MA---------DN-----------EKLDNQRLK-NF-KNK-G-R-----DL---E------------TMRRQRT-EVVVELRKNKRDEHLLK
A0A6P7JGS9 : --------MA---------DN-----------EKLDNQRLK-NF-KNK-G-R-----DL---E------------TMRRQRT-EVVVELRKNKRDEHLLK
A0A6P7PBN0 : --------MA---------DN-----------EKLDNQRLK-NF-KNK-G-R-----DL---E------------TMRRQRT-EVVVELRKNKRDEHLLK
A0A667XPZ3 : --TKAVANMA---------DN-----------EKLDNQRLK-NF-KNK-G-R-----DL---E------------TMRRQRT-EVVVELRKNKRDEHLLK
A0A1A7YVU0 : --------MA---------DN-----------EKLDNQRLK-NF-KNK-G-R-----DL---E------------TMRRQRT-EVVVELRKNKRDEHLLK
A0A1A8U2Q0 : --------MA---------DN-----------EKLDNQRLK-NF-KNK-G-R-----DL---E------------TMRRQRT-EVVVELRKNKRDEHLLK
A0A2I4CTN3 : --------MA---------DN-----------EKLDNQRLK-NF-KNK-G-R-----DL---E------------TMRRQRT-EVVVELRKNKRDEHLLK
A0A6I9NX86 : --------MA---------DN-----------EKLDNQRLK-NF-KNK-G-R-----DL---E------------TMRRQRT-EVVVELRKNKRDEHLLK
A0A673BBF6 : --VSTTANMA---------DN-----------EKLDNQRLK-NF-KNK-G-R-----DL---E------------TMRRQRT-EVVVELRKNKRDEHLLK
A0A1A8IH68 : --------MA---------DN-----------EKLDNQRLK-NF-KNK-G-R-----DL---E------------TMRRQRT-EVVVELRKNKRDEHLLK
A0A1A8FJ46 : --------MA---------DN-----------EKLDNQRLK-NF-KNK-G-R-----DL---E------------TMRRQRT-EVVVELRKNKRDEHLLK
I3J6S5     : --------MA---------DN-----------EKLDNQRLK-NF-KNK-G-R-----DL---E------------TMRRQRT-EVVVELRKNKRDEHLLK
A0A672FRF4 : --PPTTANMA---------DN-----------EKLDNQRLK-NF-KNK-G-R-----DL---E------------TMRRQRT-EVVVELRKNKRDEHLLK
A0A1A8MER6 : --------MA---------DN-----------EKLDNQRLK-NF-KNK-G-R-----DL---E------------TMRRQRT-EVVVELRKNKRDEHLLK
A0A6P8WCL6 : --------MA---------DN-----------EKLDNQRLK-NF-KNK-G-R-----DL---E------------TMRRQRT-EVVVELRKNKRDEHLLK
A0A1S3T0D1 : --------MA---------DN-----------EKLDNQRLK-NF-KNK-G-R-----DL---E------------TMRRQRT-EVVVELRKNKRDEHLLK
A0A673ZKB1 : --HPKIANMA---------DN-----------EKLDNQRLK-NF-KNK-G-R-----DL---E------------TMRRQRT-EVVVELRKNKRDEHLLK
A0A1A8QHT5 : --------MA---------DN-----------EKLDNQRLK-NF-KNK-G-R-----DL---E------------TMRRQRT-EVVVELRKNKRDEHLLK
A0A1A8DEU3 : --------MA---------DN-----------EKLDNQRLK-NF-KNK-G-R-----DL---E------------TMRRQRT-EVVVELRKNKRDEHLLK
H0VHE6     : --RITGPAMA---------DN-----------EKLDNQRLK-NF-KNK-G-R-----DL---E------------TMRRQRN-EVVVELRKNKRDEHLLK
O00629     : ---------A---------DN-----------EKLDNQRLK-NF-KNK-G-R-----DL---E------------TMRRQRN-EVVVELRKNKRDEHLLK
A0A452HUT1 : --------MA---------DN-----------EKLDNQRLK-NF-KNK-G-R-----DL---E------------TMRRQRN-EVVVELRKNKRDEHLLK
V8PFW4     : --------MA---------DN-----------EKLDNQRLK-NF-KNK-G-R-----DL---E------------TMRRQRN-EVVVELRKNKRDEHLLK
M3XV10     : --------MA---------DN-----------EKLDNQRLK-NF-KNK-G-R-----DL---E------------TMRRQRN-EVVVELRKNKRDEHLLK
A0A6J2L227 : --------MA---------DN-----------EKLDNQRLK-NF-KNK-G-R-----DL---E------------TMRRQRN-EVVVELRKNKRDEHLLK
A0A2K6NFI6 : --------MA---------DN-----------EKLDNQRLK-NF-KNK-G-R-----DL---E------------TMRRQRN-EVVVELRKNKRDEHLLK
A0A3Q7T5I3 : --------MA---------DN-----------EKLDNQRLK-NF-KNK-G-R-----DL---E------------TMRRQRN-EVVVELRKNKRDEHLLK
M3WRK2     : --------MA---------DN-----------EKLDNQRLK-NF-KNK-G-R-----DL---E------------TMRRQRN-EVVVELRKNKRDEHLLK
A0A2R8ZEY4 : --------MA---------DN-----------EKLDNQRLK-NF-KNK-G-R-----DL---E------------TMRRQRN-EVVVELRKNKRDEHLLK
A0A1S2ZT18 : --------MA---------DN-----------EKLDNQRLK-NF-KNK-G-R-----DL---E------------TMRRQRN-EVVVELRKNKRDEHLLK
A0A2K6K7P6 : --------MA---------DN-----------EKLDNQRLK-NF-KNK-G-R-----DL---E------------TMRRQRN-EVVVELRKNKRDEHLLK
A0A6J0TFY7 : --------MA---------DN-----------EKLDNQRLK-NF-KNK-G-R-----DL---E------------TMRRQRN-EVVVELRKNKRDEHLLK
H0X6W3     : --KVTGPAMA---------DN-----------EKLDNQRLK-NF-KNK-G-R-----DL---E------------TMRRQRN-EVVVELRKNKRDEHLLK
G1PMS5     : --KVTGPAMA---------DN-----------EKLDNQRLK-NF-KNK-G-R-----DL---E------------TMRRQRN-EVVVELRKNKRDEHLLK
A0A672UI36 : --------MA---------DN-----------EKLDNQRLK-NF-KNK-G-R-----DL---E------------TMRRQRN-EVVVELRKNKRDEHLLK
```

```
                         *        20         *        40         *        60         *        80         *       100
A0A6P9DVB1 : --------MA---------DN-----------EKLDNQRLK-NF-KNK-G-R-----DL---E------------TMRRQRN-EVVVELRKNKRDEHLLK
A0A250YJK4 : --------MA---------DN-----------EKLDNQRLK-NF-KNK-G-R-----DL---E------------TMRRQRN-EVVVELRKNKRDEHLLK
J9PAW2     : --------MA---------DN-----------EKLDNQRLK-NF-KNK-G-R-----DL---E------------TMRRQRN-EVVVELRKNKRDEHLLK
A0A2K5WX73 : --------MA---------DN-----------EKLDNQRLK-NF-KNK-G-R-----DL---E------------TMRRQRN-EVVVELRKNKRDEHLLK
A0A6P3YIU1 : --------MA---------DN-----------EKLDNQRLK-NF-KNK-G-R-----DL---E------------TMRRQRN-EVVVELRKNKRDEHLLK
A0A2K5NJH9 : --------MA---------DN-----------EKLDNQRLK-NF-KNK-G-R-----DL---E------------TMRRQRN-EVVVELRKNKRDEHLLK
A0A6J1ZM48 : --------MA---------DN-----------EKLDNQRLK-NF-KNK-G-R-----DL---E------------TMRRQRN-EVVVELRKNKRDEHLLK
A0A340X538 : --------MA---------DN-----------EKLDNQRLK-NF-KNK-G-R-----DL---E------------TMRRQRN-EVVVELRKNKRDEHLLK
A0A3Q7UWJ8 : --------MA---------DN-----------EKLDNQRLK-NF-KNK-G-R-----DL---E------------TMRRQRN-EVVVELRKNKRDEHLLK
A0A670Z644 : --------MA---------DN-----------EKLDNQRLK-NF-KNK-G-R-----DL---E------------TMRRQRN-EVVVELRKNKRDEHLLK
I3M4E9     : --GVTGPAMA---------DN-----------EKLDNQRLK-NF-KNK-G-R-----DL---E------------TMRRQRN-EVVVELRKNKRDEHLLK
F6VIF8     : --GVTGPAMA---------DN-----------EKLDNQRLK-NF-KNK-G-R-----DL---E------------TMRRQRN-EVVVELRKNKRDEHLLK
A0A5F9DRC8 : --------MA---------DN-----------EKLDNQRLK-NL-ENK-G-R-----DL---E------------TMRRQRN-EVVVELRKNKRDEHLLK
A0A6J1U177 : --------MA---------DN-----------EKLDNQRLK-NF-KNK-G-R-----DL---E------------TMRRQRN-EVVVELRKNKRDEHLLK
A0A6P6D5V3 : --------MA---------DN-----------EKLDNQRLK-NF-KNK-G-R-----DL---E------------TMRRQRN-EVVVELRKNKRDEHLLK
G1QXC4     : --------MA---------DN-----------EKLDNQRLK-NF-KNK-G-R-----DL---E------------TMRRQRN-EVVVELRKNKRDEHLLK
A0A2K5PA95 : --------MA---------DN-----------EKLDNQRLK-NF-KNK-G-R-----DL---E------------TMRRQRN-EVVVELRKNKRDEHLLK
H2QNN5     : --------MA---------DN-----------EKLDNQRLK-NF-KNK-G-R-----DL---E------------TMRRQRN-EVVVELRKNKRDEHLLK
A0A2Y9SU99 : --------MA---------DN-----------EKLDNQRLK-NF-KNK-G-R-----DL---E------------TMRRQRN-EVVVELRKNKRDEHLLK
A0A4W2HRN9 : --------MA---------DN-----------EKLDNQRLK-NF-KNK-G-R-----DL---E------------TMRRQRN-EVVVELRKNKRDEHLLK
A0A6J3RA32 : --------MA---------DN-----------EKLDNQRLK-NF-KNK-G-R-----DL---E------------TMRRQRN-EVVVELRKNKRDEHLLK
A0A452SUQ2 : --------MA---------DN-----------EKLDNQRLK-NF-KNK-G-R-----DL---E------------TMRRQRN-EVVVELRKNKRDEHLLK
K7FLG7     : --------MA---------DN-----------EKLDNQRLK-NF-KNK-G-R-----DL---E------------TMRRQRN-EVVVELRKNKRDEHLLK
A0A674ILH1 : --------MA---------DN-----------EKLDNQRLK-NF-KNK-G-R-----DL---E------------TMRRQRN-EVVVELRKNKRDEHLLK
A0A3Q2I6U1 : --GVAGPAMA---------DN-----------EKLDNQRLK-NF-KNK-G-R-----DL---E------------TMRRQRN-EVVVELRKNKRDEHLLK
A0A670I2H4 : --------MA---------DN-----------EKLDNQRLK-NF-KNK-G-R-----DL---E------------TMRRQRN-EVVVELRKNKRDEHLLK
A0A2U3WCB1 : --------MA---------DN-----------EKLDNQRLK-NF-KNK-G-R-----DL---E------------TMRRQRN-EVVVELRKNKRDEHLLK
A0A667HGJ4 : --------MA---------DN-----------EKLDNQRLK-NF-KNK-G-R-----DL---E------------TMRRQRN-EVVVELRKNKRDEHLLK
A0A3S5ZPE0 : --------MA---------DN-----------EKLDNQRLK-NF-KNK-G-R-----DL---E------------TMRRQRN-EVVVELRKNKRDEHLLK
A0A6J0Z606 : --------MA---------DN-----------EKLDNQRLK-NF-KNK-G-R-----DL---E------------TMRRQRN-EVVVELRKNKRDEHLLK
A0A096MXT2 : --------MA---------DN-----------EKLDNQRLK-NF-KNK-G-R-----DL---E------------TMRRQRN-EVVVELRKNKRDEHLLK
A0A6I9JJK3 : --------MA---------DN-----------EKLDNQRLK-NF-KNK-G-R-----DL---E------------TMRRQRN-EVVVELRKNKRDEHLLK
A0A3Q7PNZ1 : --------MA---------DN-----------EKLDNQRLK-NF-KNK-G-R-----DL---E------------TMRRQRN-EVVVELRKNKRDEHLLK
A0A2K6DJI1 : --------MA---------DN-----------EKLDNQRLK-NF-KNK-G-R-----DL---E------------TMRRQRN-EVVVELRKNKRDEHLLK
A0A663DS62 : --------MA---------DN-----------EKLDNQRLK-NF-KNK-G-R-----DL---E------------TMRRQRN-EVVVELRKNKRDEHLLK
A0A0B4J1E7 : --RVAGPAMA---------DN-----------EKLDNQRLK-NF-KNK-G-R-----DL---E------------TMRRQRN-EVVVELRKNKRDEHLLK
A0A6P5PG61 : --RVAGPAMA---------DN-----------EKLDNQRLK-NF-KNK-G-R-----DL---E------------TMRRQRN-EVVVELRKNKRDEHLLK
Q56R17     : --------MA---------DN-----------EKLDNQRLK-NF-KNK-G-R-----DL---E------------TMRRQRN-EVVVELRKNKRDEHLLK
A0A1V4KM60 : --------MA---------DN-----------EKLDNQRLK-NF-KNK-G-R-----DL---E------------TMRRQRN-EVVVELRKNKRDEHLLK
A0A1U7T2Y1 : --------MA---------DN-----------EKLDNQRLK-NF-KNK-G-R-----DL---E------------TMRRQRN-EVVVELRKNKRDEHLLK
A0A6P5BS84 : --------MA---------DN-----------EKLDNQRLK-NF-KNK-G-R-----DL---E------------TMRRQRN-EVVVELRKNKRDEHLLK
H9EQP0     : --------MA---------DN-----------EKLDNQRLK-NF-KNK-G-R-----DL---E------------TMRRQRN-EVVVELRKNKRDEHLLK
A0A2Y9I3B6 : --------MA---------DN-----------EKLDNQRLK-NF-KNK-G-R-----DL---E------------TMRRQRN-EVVVELRKNKRDEHLLK
F7FL50     : --------MA---------DN-----------EKLDNQRLK-NF-KNK-G-R-----DL---E------------TMRRQRN-EVVVELRKNKRDEHLLK
A0A6J2CAN4 : --------MA---------DN-----------EKLDNQRLK-NF-KNK-G-R-----DL---E------------TMRRQRN-EVVVELRKNKRDEHLLK
A0A6I9I3N0 : --------MA---------DN-----------EKLDNQRLK-NF-KNK-G-R-----DL---E------------TMRRQRN-EVVVELRKNKRDEHLLK
A0A6P4VF85 : --------MA---------DN-----------EKLDNQRLK-NF-KNK-G-R-----DL---E------------TMRRQRN-EVVVELRKNKRDEHLLK
Q5ZLF7     : --------MA---------DN-----------EKLDNQRLK-NF-KNK-G-R-----DL---E------------TMRRQRN-EVVVELRKNKRDEHLLK
A0A6J3GID2 : --------MA---------DN-----------EKLDNQRLK-NF-KNK-G-R-----DL---E------------TMRRQRN-EVVVELRKNKRDEHLLK
A0A2Y9NIB7 : --------MA---------DN-----------EKLDNQRLK-NF-KNK-G-R-----DL---E------------TMRRQRN-EVVVELRKNKRDEHLLK
A0A6P3RQQ3 : --------MA---------DN-----------EKLDNQRLK-NF-KNK-G-R-----DL---E------------TMRRQRN-EVVVELRKNKRDEHLLK
A0A2K5DHE6 : --------MA---------DN-----------EKLDNQRLK-NF-KNK-G-R-----DL---E------------TMRRQRN-EVVVELRKNKRDEHLLK
A0A7F8RJW8 : --------MA---------DN-----------EKLDNQRLK-NF-KNK-G-R-----DL---E------------TMRRQRN-EVVVELRKNKRDEHLLK
A0A671DSE8 : --------MA---------DN-----------EKLDNQRLK-NF-KNK-G-R-----DL---E------------TMRRQRN-EVVVELRKNKRDEHLLK
A0A6P5KG24 : --------MA---------DN-----------EKLDNQRLK-NF-KNK-G-R-----DL---E------------TMRRQRN-EVVVELRKNKRDEHLLK
A0A452GBK4 : --------MA---------DN-----------EKLDNQRLK-NF-KNK-G-R-----DL---E------------TMRRQRN-EVVVELRKNKRDEHLLK
A0A6I9YGU7 : --------MA---------DN-----------EKLDNQRLK-NF-KNK-G-R-----DL---E------------TMRRQRN-EVAVELRKNKRDEHLLK
A0A674G8H0 : --PWPPPAMA---------DS-----------EKLDNQRLK-NF-KNK-G-R-----DL---E------------TMRRQRN-EVVVELRKNKRDEHLLK
A0A6J2IMM6 : --------MA---------DS-----------EKLDNQRLK-NF-KNK-G-R-----DL---E------------TMRRQRN-EVVVELRKNKRDEHLLK
A0A5G2R9L2 : --------MA---------DS-----------EKLDNQRLK-NF-KNK-G-R-----DL---E------------TMRRQRN-EVVVELRKNKRDEHLLK
A0A0B8RSP3 : --------MA---------DS-----------EKLDNQRLK-NF-KNK-G-R-----DL---E------------TMRRQRN-EVVVELRKNKRDEHLLK
A0A3Q0DIG1 : --------MA---------DH-----------EKLDNQRLK-NF-KNK-G-R-----DL---E------------TMRRQRN-EVVVELRKNKRDEHLLK
F6TB69     : -RPGA-PIMA---------DN-----------DKLDNQRLK-NF-KNK-G-R-----DL---E------------SMRRQRN-EVVVELRKNKRDEHLLK
```

```
                     *         20          *         40          *         60          *         80          *        100
A0A1L8GA53 : --------MA---------DN-----------DKLDNQRLK-NF-KNK-G-R-----DL---E------------SMRRQRN-EVVVELRKNKRDEHLLK
A0A2Z2GP46 : --------MA---------DN-----------DKLDNQRLK-NF-KNK-G-R-----DL---E------------SMRRQRT-EVVVELRKNKRDEHLLK
A0A6P7Z784 : --------MA---------DN-----------EKLDNQRLK-NF-KNK-G-R-----DL---E------------SMRRQRN-EVVVELRKNKRDEHLLK
A0A6I8PKN1 : --GGAGPTMA---------DH-----------EKLDNQRLK-NF-KNK-G-R-----DL---E------------SMRRQRN-EVVVELRKNKRDEHLLK
A0A6P8S910 : --------MA---------DN-----------EKLDNQRLK-NF-KNK-G-R-----DL---E------------SMRRQRN-EVVVELRKNKRDEHLLK
U3IJM3     : -------M-A---------EQ-----------EKLDNQRLK-NF-KNK-G-R-----DP---E------------TMRRQRN-EVVVELRKNKRDEHLLK
A0A6J3DFR9 : -------M-A---------EQ-----------EKLDNQRLK-NF-KNK-G-R-----DL---E------------TMRRQRN-EVVVELRKNKRDEHLLK
A0A669Q581 : -----GLLRA---------RAPGP--------RKLDNQRLK-NF-KNK-G-R-----DL---E------------TMRRQRN-EVVVELRKNKRDEHLLK
A0A4W3JQE6 : --VPRSANMA---------EN-----------DKLENQRLK-NF-KNK-G-R-----DL---E------------SMRRQRN-EVVVELRKNKRVEHLFK
A0A4W3K100 : --------MA---------EN-----------DKLENQRLK-NF-KNK-G-R-----DL---EV-----------SGRAAASGELFIPLQ-NKRVEHLFK
A0A6F9DGB8 : ----MNGALE---------TMD----------PKQEYHRFK-NW-KHK-G-R-----DA---E------------TMRRQRN-EMSVELRKQKRGDHLLK
A0A2J7RBM0 : --------MA-------S-DQ----Q------SK--NR-MQ-VF-KNK-G-K-----DQ---D------------EMRRRRN-EVTVELRKNKREETLQK
A0A0C9RV98 : --------MA-------A-EM----Q------NK--SR-MM-VF-KNK-G-K-----DQ---D------------EMRRRRN-EVTVELRKNKREETLQK
A0A0A9WQY7 : --------MA-------S-DP----H------TK--YR-MQ-VF-KNK-G-K-----DQ---D------------EMRRRRN-EVTVELRKNKREETLLK
A0A2P8ZE70 : --------MGDGNVMLYS-DQKPSDHLSCYSGTKTHSKTCA-VY-VSETG-L-----AQGTAEH-----------EMRRRRN-EVTVELRKNKREETLQK
A0A5N5SW62 : --------MEKAQ----G-DQ-----------NR--NR-LG-VF-KNK-G-K-----DQ---E------------EMRRRRT-EYTVELRKCKREETFQK
A0A0K8WEE2 : --------MT-------SLEP-----------NV--NR-LQ-NF-KNK-G-K-----DQ---D------------EMRRRRN-EVTVELRKNKREETILK
A0A0A1XJB6 : --------MT-------SLEP-----------NV--NR-LQ-NF-KNK-G-K-----DQ---D------------EMRRRRN-EVTVELRKNKREETILK
A0A4Y2Q2Z3 : --------MS-------T-DN----------INK--SR-FQ-HF-KNK-G-K-----DQ---D------------EMRRRRN-EVTFELRKNKRDESLLK
A0A2B4S0M6 : --------MS---------T------------KSTDSRQ-R-YY-KNK-D-K-----DQ---S------------ALRQRRN-EVTIELRKAKRDDCVQK
A0A6G1S6S5 : --------MS---------D-----------------HRAR--Y-KNT-G-L-----DA---K------------ELRRRRE-ENSIQLRKQKRDDVLSK
A0A0V0RTF5 : ----MLQKLR---------NV-----------CQEKAEKYR-CFSDSQ-CPTT----ST---TR-----------ELRRRRT-DMSFELRKSKREDTLSK
cIBB4      : --------MA---------DN-----------EKLDNQRLK-NF-KNK-G-R-----DL---E------------TMRRQRN-EVVVELRKNKRDEHLLK

                     *
H3AN54     : RR--NVP-Q-EDI-CE-
A0A1U8DF45 : RR--NVP-H-EDI-CE-
A0A4W3JN51 : RR--NVP-Q-EDV-CE-
A0A6P7QP70 : RR--NVP-Q-EDI-CE-
G1SG68     : RR--NVP-H-EDI-CE-
A0A663DS72 : RR--NVP-H-EDI-CE-
A0A674PLW0 : RR--NVP-Y-EYI-CE-
U3JFA4     : RR--NVP-H-EDI-CE-
A0A673B6V3 : RR--NVP-H-EDI-CE-
A0A6J1ZKW3 : RR--NVP-H-EDI-CE-
A0A6J2CG34 : RR--NVP-H-EDI-CE-
A0A6P4VIN5 : RR--NVP-H-EDI-CE-
A0A7E6D5F7 : RR--NVP-H-EDI-CE-
A0A6P6BYU9 : RR--NVP-H-EDI-CE-
A0A3Q0GSB0 : RR--NVP-H-EDI-CE-
A0A5F4W1H4 : RR--NVP-H-EDI-CE-
A0A673BC87 : RR--NVP-H-EDI-CE-
A0A6I9KX70 : RR--NVP-H-EDI-CE-
A0A6J2QXP4 : RR--NVP-Y-ENI-CE-
A0A4Z2IS62 : RR--NVP-N-EDI-CE-
A0A4W4FLE0 : RR--NVP-N-EDI-CD-
W5UA83     : RR--NVP-H-EDI-CD-
A0A6J2VJW1 : RR--NVP-H-EDI-CD-
F1QEW3     : RR--NVP-H-EDI-CD-
A0A6P3WAK5 : RR--NVP-Q-EDI-CD-
A0A4W6BU17 : RR--NVP-H-EDI-CE-
A0A665UA99 : RR--NVP-H-EDF-CE-
A0A665UA62 : RR--NVP-H-EDF-CE-
H2LP51     : RR--NVP-H-EDL-CE-
A0A6P7JGS9 : RR--NVP-H-EDI-CE-
A0A6P7PBN0 : RR--NVP-H-EDI-CE-
A0A667XPZ3 : RR--NVP-H-EDI-CE-
A0A1A7YVU0 : RR--NVP-H-EDI-CE-
A0A1A8U2Q0 : RR--NVP-H-EDI-CE-
A0A2I4CTN3 : RR--NVP-H-EDI-CE-
A0A6I9NX86 : RR--NVP-H-EDI-CE-
A0A673BBF6 : RR--NVP-H-EDI-CE-
A0A1A8IH68 : RR--NVP-H-EDI-CE-
```

```
                          *
A0A1A8FJ46 : RR--NVP-H-EDI-CE-
I3J6S5     : RR--NVP-H-EDI-CE-
A0A672FRF4 : RR--NVP-H-EDI-CE-
A0A1A8MER6 : RR--NVP-H-EDI-CE-
A0A6P8WCL6 : RR--NVP-H-EDI-CE-
A0A1S3T0D1 : RR--NVP-H-EDI-CE-
A0A673ZKB1 : RR--NVP-H-EDI-CE-
A0A1A8QHT5 : RR--NVP-H-EDI-CE-
A0A1A8DEU3 : RR--NVP-H-EDI-CE-
H0VHE6     : RR--NVP-H-EDI-CE-
O00629     : RR--NVP-H-EDI-CE-
A0A452HUT1 : RR--NVP-H-EDI-CE-
V8PFW4     : RR--NVP-H-EDI-CE-
M3XV10     : RR--NVP-H-EDI-CE-
A0A6J2L227 : RR--NVP-H-EDI-CE-
A0A2K6NFI6 : RR--NVP-H-EDI-CE-
A0A3Q7T5I3 : RR--NVP-H-EDI-CE-
M3WRK2     : RR--NVP-H-EDI-CE-
A0A2R8ZEY4 : RR--NVP-H-EDI-CE-
A0A1S2ZT18 : RR--NVP-H-EDI-CE-
A0A2K6K7P6 : RR--NVP-H-EDI-CE-
A0A6J0TFY7 : RR--NVP-H-EDI-CE-
H0X6W3     : RR--NVP-H-EDI-CE-
G1PMS5     : RR--NVP-H-EDI-CE-
A0A672UI36 : KR--NVP-H-EDI-CE-
A0A6P9DVB1 : RR--NVP-H-EDI-CE-
A0A250YJK4 : RR--NVP-H-EDI-CE-
J9PAW2     : RR--NVP-H-EDI-CE-
A0A2K5WX73 : RR--NVP-H-EDI-CE-
A0A6P3YIU1 : RR--NVP-H-EDI-CE-
A0A2K5NJH9 : RR--NVP-H-EDI-CE-
A0A6J1ZM48 : RR--NVP-H-EDI-CE-
A0A340X538 : RR--NVP-H-EDI-CE-
A0A3Q7UWJ8 : RR--NVP-H-EDI-CE-
A0A670Z644 : RR--NVP-H-EDI-CE-
I3M4E9     : RR--NVP-H-EDI-CE-
F6VIF8     : RR--NVP-H-EDI-CE-
A0A5F9DRC8 : RR--NVP-H-EDI-CE-
A0A6J1U177 : RR--NVP-H-EDI-CE-
A0A6P6D5V3 : RR--NVP-H-EDI-CE-
G1QXC4     : RR--NVP-H-EDI-CE-
A0A2K5PA95 : RR--NVP-H-EDI-CE-
H2QNN5     : RR--NVP-H-EDI-CE-
A0A2Y9SU99 : RR--NVP-H-EDI-CE-
A0A4W2HRN9 : RR--NVP-H-EDI-CE-
A0A6J3RA32 : RR--NVP-H-EDI-CE-
A0A452SUQ2 : RR--NVP-H-EDI-CE-
K7FLG7     : RR--NVP-H-EDI-CE-
A0A674ILH1 : RR--NVP-H-EDI-CE-
A0A3Q2I6U1 : RR--NVP-H-EDV-CE-
A0A670I2H4 : RR--NVP-H-EDI-CE-
A0A2U3WCB1 : RR--NVP-H-EDI-CE-
A0A667HGJ4 : RR--NVP-H-EDI-CE-
A0A3S5ZPE0 : RR--NVP-H-EDI-CE-
A0A6J0Z606 : RR--NVP-H-EDI-CE-
A0A096MXT2 : RR--NVP-H-EDI-CE-
A0A6I9JJK3 : RR--NVP-H-EDI-CE-
A0A3Q7PNZ1 : RR--NVP-H-EDI-CE-
A0A2K6DJI1 : RR--NVP-H-EDI-CE-
A0A663DS62 : RR--NVP-H-EDI-CE-
A0A0B4J1E7 : RR--NVP-Q-EDI-CE-
A0A6P5PG61 : RR--NVP-Q-EDI-CE-
Q56R17     : RR--NVP-Q-EDI-CE-
```

```
                         *
A0A1V4KM60 : RR--NVP-H-EDI-CE-
A0A1U7T2Y1 : RR--NVP-H-EDI-CE-
A0A6P5BS84 : RR--NVP-H-EDI-CE-
H9EQP0     : RR--NVP-H-EDI-CE-
A0A2Y9I3B6 : RR--NVP-H-EDI-CE-
F7FL50     : RR--NVP-H-EDI-CE-
A0A6J2CAN4 : RR--NVP-H-EDI-CE-
A0A6I9I3N0 : RR--NVP-H-EDI-CE-
A0A6P4VF85 : RR--NVP-H-EDI-CE-
Q5ZLF7     : RR--NVP-H-EDI-CE-
A0A6J3GID2 : RR--NVP-H-EDI-CE-
A0A2Y9NIB7 : RR--NVP-H-EDI-CE-
A0A6P3RQQ3 : RR--NVP-H-EDI-CE-
A0A2K5DHE6 : RR--NVP-H-EDI-CE-
A0A7F8RJW8 : RR--NVP-H-EDI-CE-
A0A671DSE8 : RR--NVP-H-EDI-CE-
A0A6P5KG24 : RR--NVP-H-EDI-CE-
A0A452GBK4 : RR--NVP-H-EDI-CE-
A0A6I9YGU7 : RR--NVP-H-EDI-CE-
A0A674G8H0 : RR--NVP-H-EDI-CE-
A0A6J2IMM6 : RR--NVP-H-EDI-CE-
A0A5G2R9L2 : RR--NVP-H-EDI-CE-
A0A0B8RSP3 : RR--NVP-H-EDI-CE-
A0A3Q0DIG1 : RR--NVP-H-EDI-CE-
F6TB69     : RR--NVP-H-EDI-CE-
A0A1L8GA53 : RR--NVP-H-EDI-CE-
A0A2Z2GP46 : RR--NVP-H-EDI-CE-
A0A6P7Z784 : RR--NVP-H-EDI-CE-
A0A6I8PKN1 : RR--NVP-H-EDI-CE-
A0A6P8S910 : RR--NVP-H-EDI-CE-
U3IJM3     : RR--NVP-H-EDI-CE-
A0A6J3DFR9 : RR--NVP-H-EDI-CE-
A0A669Q581 : RR--NVP-H-EDI-CE-
A0A4W3JQE6 : RR--NVP-Q-EDV-CE-
A0A4W3K100 : RR--NVP-Q-EDV-CE-
A0A6F9DGB8 : RR--NVPIA-DDS-L--
A0A2J7RBM0 : RR--NVP-V-ADS-TD-
A0A0C9RV98 : RR--NVP-I-ADS-TD-
A0A0A9WQY7 : RR--NVP-T-VDS-TD-
A0A2P8ZE70 : RR--NVP-I-ADS-TD-
A0A5N5SW62 : RR--NVP-S-ADV-ID-
A0A0K8WEE2 : RR--NVP-N-MDSNT--
A0A0A1XJB6 : RR--NVP-N-MDSNT--
A0A4Y2Q2Z3 : RR--NVP-Q-TDT-TD-
A0A2B4S0M6 : KR--NVP---QDI-VDD
A0A6G1S6S5 : RRTLGAP-T-NDM----
A0A0V0RTF5 : KR--SI--HVQSV--EI
cIBB4      : RR--NVP-H-EDI-CE-
```

```
                        *        20         *        40         *        60         *        80
A0A4W3HMB2 : ---MN----DMAS-P--VKD--NCRMRSYKNKALNTQEMRRRREEEGIQLRKQKREEQL--FKRRNV-----SVPGGEE--
A0A4W3HQ66 : ---IKCIIVDMAS-P--VKD--NCRMRSYKNKALNTQEMRRRREEEGIQLRKQKREEQL--FKRRNV-----SVPGGEE--
A0A4W3HM51 : -GRTEP---EVPDRP--VKD--NCRMRSYKNKALNTQEMRRRREEEGIQLRKQKREEQL--FKRRNV-----SVPGGEE--
H3ASY1     : --IIFFL-HTMAS-P--VKD--NYRMKSYKNKALNPQEMRRRREEEGIQLRKQKREEQL--FKRRNV-----SLPLNDE--
A0A6P3VTJ8 : --MV-----TMAS-P--GKD--SYRMKSYKNKALNPQEMRRRREEEGIQLRKQKREEQL--FKRRNV-----SLPPDDE--
A0A6J2DZ74 : ------M-DAMAS-P--GKD--NYRMKSYKNKALNPQEMRRRREEEGIQLRKQKREEQL--FKRRNV-----SLPRNDE--
G1STV1     : --SSTLM-DAMAS-P--GKD--NYRMKSYKNKALNPQEMRRRREEEGIQLRKQKREEQL--FKRRNV-----SLPRNDE--
A0A6P5CAC8 : ------M-DAMAS-P--GKD--NYRMKSYKNKALNPQEMRRRREEEGIQLRKQKREEQL--FKRRNV-----SLPRNDE--
A0A6J3HK00 : ------M-DAMAS-P--GKD--NYRMKSYKNKALNPQEMRRRREEEGIQLRKQKREEQL--FKRRNV-----SLPRNDE--
A0A5F4WLV0 : ------M-DAMAS-P--GKD--NYRMKSYKNKALNPQEMRRRREEEGIQLRKQKREEQL--FKRRNV-----SLPRNDE--
W5PGW2     : ------M-DAMAS-P--GKD--NYRMKSYKNKALNPQEMRRRREEEGIQLRKQKREEQL--FKRRNV-----SLPRNDE--
A0A2K5Q2A3 : ------M-DAMAS-P--GKD--NYRMKSYKNKALNPQEMRRRREEEGIQLRKQKREEQL--FKRRNV-----SLPRNDE--
A0A671FXJ8 : ------M-DAMAS-P--GKD--NYRMKSYKNKALNPQEMRRRREEEGIQLRKQKREEQL--FKRRNV-----SLPRNDE--
A0A6J0Y751 : ------M-DAMAS-P--GKD--NYRMKSYKNKALNPQEMRRRREEEGIQLRKQKREEQL--FKRRNV-----SLPRNDE--
A0A2K5MAE7 : ------M-DAMAS-P--GKD--NYRMKSYKNKALNPQEMRRRREEEGIQLRKQKREEQL--FKRRNV-----SLPRNDE--
A0A2Y9HI85 : ------M-DAMAS-P--GKD--NYRMKSYKNKALNPQEMRRRREEEGIQLRKQKREEQL--FKRRNV-----SLPRNDE--
A0A4W2FFP7 : ------M-DAMAS-P--GKD--NYRMKSYKNKALNPQEMRRRREEEGIQLRKQKREEQL--FKRRNV-----SLPRNDE--
A0A3Q7NQD1 : ------M-DAMAS-P--GKD--NYRMKSYKNKALNPQEMRRRREEEGIQLRKQKREEQL--FKRRNV-----SLPRNDE--
F1PQD8     : ------M-DAMAS-P--GKD--NYRMKSYKNKALNPQEMRRRREEEGIQLRKQKREEQL--FKRRNV-----SLPRNDE--
A0A6I9IAP3 : ------M-DAMAS-P--GKD--NYRMKSYKNKALNPQEMRRRREEEGIQLRKQKREEQL--FKRRNV-----SLPRNDE--
A0A2K5HJB4 : ------M-DAMAS-P--GKD--NYRMKSYKNKALNPQEMRRRREEEGIQLRKQKREEQL--FKRRNV-----SLPRNDE--
A0A2K6C6R2 : ------M-DAMAS-P--GKD--NYRMKSYKNKALNPQEMRRRREEEGIQLRKQKREEQL--FKRRNV-----SLPRNDE--
A0A6P4T529 : ------M-DAMAS-P--GKD--NYRMKSYKNKALNPQEMRRRREEEGIQLRKQKREEQL--FKRRNV-----SLPRNDE--
A0A452G264 : ------M-DAMAS-P--GKD--NYRMKSYKNKALNPQEMRRRREEEGIQLRKQKREEQL--FKRRNV-----SLPRNDE--
H2PK58     : --RATLM-DAMAS-P--GKD--NYRMKSYKNKALNPQEMRRRREEEGIQLRKQKREEQL--FKRRNV-----SLPRNDE--
A0A2I3GNS8 : ------M-DAMAS-P--GKD--NYRMKSYKNKALNPQEMRRRREEEGIQLRKQKREEQL--FKRRNV-----SLPRNDE--
A0A6J2LE28 : ------M-DAMAS-P--GKD--NYRMKSYKNKALNPQEMRRRREEEGIQLRKQKREEQL--FKRRNV-----SLPRNDE--
A0A2K5TMX7 : ------M-DAMAS-P--GKD--NYRMKSYKNKALNPQEMRRRREEEGIQLRKQKREEQL--FKRRNV-----SLPRNDE--
F1N1K5     : ------M-DAMAS-P--GKD--NYRMKSYKNKALNPQEMRRRREEEGIQLRKQKREEQL--FKRRNV-----SLPRNDE--
A0A2K6NC93 : ------M-DAMAS-P--GKD--NYRMKSYKNKALNPQEMRRRREEEGIQLRKQKREEQL--FKRRNV-----SLPRNDE--
A0A6P3J9Q1 : ------M-DAMAS-P--GKD--NYRMKSYKNKALNPQEMRRRREEEGIQLRKQKREEQL--FKRRNV-----SLPRNDE--
H2QTM3     : ------M-DAMAS-P--GKD--NYRMKSYKNKALNPQEMRRRREEEGIQLRKQKREEQL--FKRRNV-----SLPRNDE--
A0A6I9ZEY2 : ------M-DAMAS-P--GKD--NYRMKSYKNKALNPQEMRRRREEEGIQLRKQKREEQL--FKRRNV-----SLPRNDE--
A0A2U3XT32 : ------M-DAMAS-P--GKD--NYRMKSYKNKALNPQEMRRRREEEGIQLRKQKREEQL--FKRRNV-----SLPRNDE--
H9EPP2     : ------M-DAMAS-P--GKD--NYRMKSYKNKALNPQEMRRRREEEGIQLRKQKREEQL--FKRRNV-----SLPRNDE--
A0A0D9RTD2 : ------M-DAMAS-P--GKD--NYRMKSYKNKALNPQEMRRRREEEGIQLRKQKREEQL--FKRRNV-----SLPRNDE--
F7DMT7     : ------M-DAMAS-P--GKD--NYRMKSYKNKALNPQEMRRRREEEGIQLRKQKREEQL--FKRRNV-----SLPRNDE--
M3YS23     : ------M-DAMAS-P--GKD--NYRMKSYKNKALNPQEMRRRREEEGIQLRKQKREEQL--FKRRNV-----SLPRNDE--
A0A2K6SJ59 : ------M-DAMAS-P--GKD--NYRMKSYKNKALNPQEMRRRREEEGIQLRKQKREEQL--FKRRNV-----SLPRNDE--
A0A6P6HZV5 : ------M-DAMAS-P--GKD--NYRMKSYKNKALNPQEMRRRREEEGIQLRKQKREEQL--FKRRNV-----SLPRNDE--
A0A667FL52 : ------M-DAMAS-P--GKD--NYRMKSYKNKALNPQEMRRRREEEGIQLRKQKREEQL--FKRRNV-----SLPRNDE--
A0A384C3E9 : ------M-DAMAS-P--GKD--NYRMKSYKNKALNPQEMRRRREEEGIQLRKQKREEQL--FKRRNV-----SLPRNDE--
G3R8B6     : ------M-DAMAS-P--GKD--NYRMKSYKNKALNPQEMRRRREEEGIQLRKQKREEQL--FKRRNV-----SLPRNDE--
A0A1U7UBB2 : ------M-DAMAS-P--GKD--NYRMKSYKNKALNPQEMRRRREEEGIQLRKQKREEQL--FKRRNV-----SLPRNDE--
A0A2U3VS73 : ------M-DAMAS-P--GKD--NYRMKSYKNKALNPQEMRRRREEEGIQLRKQKREEQL--FKRRNV-----SLPRNDE--
A0A2K5BXN1 : ------M-DAMAS-P--GKD--NYRMKSYKNKALNPQEMRRRREEEGIQLRKQKREEQL--FKRRNV-----SLPRNDE--
A0A2K6A233 : ------M-DAMAS-P--GKD--NYRMKSYKNKALNPQEMRRRREEEGIQLRKQKREEQL--FKRRNV-----SLPRNDE--
A0A3Q7TSK3 : ------M-DAMAS-P--GKD--NYRMKSYKNKALNPQEMRRRREEEGIQLRKQKREEQL--FKRRNV-----SLPRNDE--
M3WVA3     : ------M-DAMAS-P--GKD--NYRMKSYKNKALNPQEMRRRREEEGIQLRKQKREEQL--FKRRNV-----SLPRNDE--
A0A6P3QTU4 : ------M-DAMAS-P--GKD--NYRMKSYKNKALNPQEMRRRREEEGIQLRKQKREEQL--FKRRNV-----SLPRNDE--
A0A452RB17 : ----------MAS-P--GKD--NYRMKSYKNKALNPQEMRRRREEEGIQLRKQKREEQL--FKRRNV-----SLPRNDE--
A0A2K6KIL0 : ----------MAS-P--GKD--NYRMKSYKNKALNPQEMRRRREEEGIQLRKQKREEQL--FKRRNV-----SLPRNDE--
A0A096NFQ3 : ----------MAS-P--GKD--NYRMKSYKNKALNPQEMRRRREEEGIQLRKQKREEQL--FKRRNV-----SLPRNDE--
A0A2U4AW54 : ----------MAC-P--GKD--NYRMKSYKNKALNPQEMRRRREEEGIQLRKQKREEQL--FKRRNV-----SLPRNDE--
A0A341CCE6 : ----------MAS-P--GKD--NYRMKSYKNKALNPQEMRRRREEEGIQLRKQKREEQL--FKRRNV-----SLPRNDE--
A0A2I3TFK8 : ---MKYI-YAMAS-P--GKD--NYRMKSYKNKALNPQEMRRRREEEGIQLRKQKREEQL--FKRRNV-----SLPRNDE--
A0A455BMW6 : ----------MAS-P--GKD--NYRMKSYKNKALNPQEMRRRREEEGIQLRKQKREEQL--FKRRNV-----SLPRNDE--
G3ST52     : --------DTMTS-S--GKD--NYRMKNYKNKALNPQEMRRRREEEGIQLRKQKREEQL--FKRRNV-----SLPRNDE--
A0A6I9JBC6 : ----------MAS-P--GKD--NYRIKNYKNKVLNPQEMRRRSEEEGIQLRKQKREEQL--FKRRNV-----SLPRNDE--
A0A340X6R4 : ----------MAS-P--GKD--NYRMKSYKNKALNPQEMRRRREEEGIQLRKQKREEQL--FKRRNV-----SLPRNDE--
A0A1D5QVJ1 : --RTRRR-DAMAS-P--GKD--NYRMKSYKNKALNPQEMRRRREEEGIQLRKQKREEQL--FKRRNV-----SLPRNDE--
A0A2K5MAA4 : --RTRRR-DAMAS-P--GKD--NYRMKSYKNKALNPQEMRRRREEEGIQLRKQKREEQL--FKRRNV-----SLPRNDE--
D2HNW3     : --------DAMAS-P--GKD--NYRMKSYKNKALNPQEMRRRREEEGIQLRKQKREEQL--FKRRNV-----SLPRNDE--
```

```
                       *        20         *        40         *        60         *        80
A0A2Y9LPW6 : ----------MAS-P--GKD--NYRMKSYKNKALNPQEMRRRREEEGIQLRKQKREEQL--FKRRNV-----SLPRNDE--
F7A9W1     : --DSDEA-DAMAS-P--GKD--NYRMKSYKNKALNPQEMRRRREEEGIQLRKQKREEQL--FKRRNV-----SLPRNDE--
H0X5T9     : --------DAMAS-P--GKD--NYRMKSYKNKALNPQEMRRRREEEGIQLRKQKREEQL--FKRRNV-----SLPRNDE--
A0A384AUJ5 : ----------MAS-P--GKD--NYRMKSYKNKALNPQEMRRRREEEGIQLRKQKREEQL--FKRRNV-----SLPRNDE--
A0A2R9AVQ0 : ----------MAS-P--GKD--NYRMKSYKNKALNPQEMRRRREEEGIQLRKQKREEQL--FKRRNV-----SLPRNDE--
Q5TD90     : ----------MAS-P--GKD--NYRMKSYKNKALNPQEMRRRREEEGIQLRKQKREEQL--FKRRNV-----YLPRNDE--
A0A286Y281 : ----------MAS-L--GKD--NYRMKSYKNKALNPQEMRRR-EEEGIQLRKQKREEQL--FKRRNV-----SLPRNDD--
A0A1U7Q8Z8 : ------M-DSMAS-T--GKD--NYRMKSYKNKALNPQEMRRRREEEGIQLRKQKRAEQL--FKRRNV-----SLPRNDD--
A0A6I9M1J8 : ------M-DSMAS-P--GKD--NYRMKSYKNKALNPQEMRRRREEEGIQLRKQKREEQL--FKRRNV-----SLPRNDD--
A0A0H2UHW5 : ------M-DSMAS-P--GKD--NYRMKSYKNKALNPQEMRRRREEEGIQLRKQKREEQL--FKRRNV-----SLPRNDD--
A0A0P6JBC4 : ----------MAS-P--GKD--NYRMKSYKNKALNPQEMRRRREEEGIQLRKQKREEQL--FKRRNV-----SLPRNDD--
A0A1S2ZL74 : ------M-DAMAS-P--GKD--NYRMKSYKNKALNSQEMRRRREEEGIQLRKQKREEQL--FKRRNV-----SLPRGDD--
A0A673UVH5 : ------M-DAMAS-P--GRD--NYRMKSYKNKALNPQEMRRRREEEGIQLRKQKREEQL--FKRRNV-----SLPRNDD--
A0A4X2M1C7 : ------M-DAMAS-P--GKD--NYRMKSYKNKALNPQEMRRRREEEGIQLRKQKREEQL--FKRRNV-----SLPANDD--
A0A6P5J864 : ------M-DAMAS-P--GKD--NYRMKSYKNKALNPQEMRRRREEEGIQLRKQKREEQL--FKRRNV-----SLPANDE--
A0A1U7RMT4 : ------M-DTMAS-P--GKD--NYRMKSYKNNALNPQEMRRRREEEGIQLRKQKREEQL--FKRRNV-----SLPGNEE--
A0A151M3O9 : ------M-DTMAS-P--GKD--NYRMKSYKNNALNPQEMRRRREEEGIQLRKQKREEQL--FKRRNV-----SLPGNEE--
A0A663LZJ9 : --FLSLI-DNMAS-P--GKD--NFRMKSYKNNALNPQEMRRRREEEGIQLRKQKREEQL--FKRRNV-----SLPGNEE--
A0A672TRT0 : --FLFLI-DNMAS-P--GKD--NFRMKSYKNNALNPQEMRRRREEEGIQLRKQKREEQL--FKRRNV-----SLPGNEE--
A0A2I0MVK9 : ----------MAS-P--GKD--NFRMKSYKNNALNPQEMRRRREEEGIQLRKQKREEQL--FKRRNV-----SLPGNEE--
A0A6I9HDF7 : ----------MAS-P--GKD--NFRMKSYKNNALNPQEMRRRREEEGIQLRKQKREEQL--FKRRNV-----SLPGNEE--
G1NKY4     : ---XFLI-DIMSS-P--GKD--NFRMKSYKNNALNPQEMRRRREEEGIQLRKQKREEQL--FKRRNV-----SLPGNEE--
A0A6J0IEC5 : ------M-DNMAS-P--GKD--NFRMKSYKNNAHNPQEMRRRREEEGIQLRKQKREEQL--FKRRNV-----SLPGNEE--
A0A6J2ILX5 : ------M-DNMAS-P--GKD--NFRMKSYKNNAHNPQEMRRRREEEGIQLRKQKREEQL--FKRRNV-----SLPGNEE--
U3I6P6     : --GCASM-DNMSS-P--GKD--NFRMKSYKNNALNPQEMRRRREEEGIQLRKQKREEQL--FKRRNV-----SLPGNEE--
A0A6J3CN91 : ------M-DNMSS-P--GKD--NFRMKSYKNNALNPQEMRRRREEEGIQLRKQKREEQL--FKRRNV-----SLPGNEE--
A0A218UX91 : ------M-DNMAS-P--GKD--NFRMKSYKNNALNPQEMRRRREEEGIQLRKQKREEQL--FKRRNV-----SLPGNEE--
A0A663ECT7 : ------M-DNMAS-P--GKD--NFRMKSYKNNALNPQEMRRRREEEGIQLRKQKREEQL--FKRRNV-----SLPGNEE--
A0A1V4JN39 : ------M-DNMAS-L--GKD--NFRMKSYKNNALNPQEMRRRREEEGIQLRKQKREEQL--FKRRNV-----SLPGNEE--
U3K917     : ----------MAS-P--GKD--NFRMKSYKNNALNPQEMRRRREEEGIQLRKQKREEQL--FKRRNV-----SLPGNEE--
A0A672TRP4 : --LICEM-NNMAS-P--GKD--NFRMKSYKNNALNPQEMRRRREEEGIQLRKQKREEQL--FKRRNV-----SLPGNEE--
A0A669Q858 : --VPAAM-DIMSS-P--GKD--NFRMKSYKNNALNPQEMRRRREEEGIQLRKQKREEQL--FKRRNV-----SLPGNEE--
E1C4J0     : --VPAAM-DIMSS-P--GKD--NFRMKSYKNNALNPQEMRRRREEEGIQLRKQKREEQL--FKRRNV-----SLPGNEE--
A0A674I1L9 : ------M-DTMAS-P--GKD--NYRMKSYKNNALNPQEMRRRREEDGIQLRKQKREEQL--FKRRNV-----SLLGNEE--
K7FKR6     : --AASLM-DAMAS-P--GKD--SYRMKSYKNNALNPQEMRRRREEDGIQLRKQKREEQL--FKRRNV-----SLLGNEE--
A0A6P9C420 : ------M-DNMAS-P--EKD--NYRMKSYKNNALNPQEMRRRREEEGIQLRKQKREEQL--FKRRNV-----SLLSSEE--
A0A6I9XJ79 : --RNAKN-DNMAS-P--EKD--NYRMKSYKNNALNPQEMRRRREEEGIQLRKQKREEQL--FKRRNV-----SLLSSEE--
A0A6J1TT36 : ------M-DNMAS-P--EKD--NYRIKSYKNNALNPQEMRRRREEEGIQLRKQKREEQL--FKRRNV-----SLLSSEE--
A0A670Y8L6 : ------M-DNMAS-P--EKD--NYRIKSYKNNALNPQEMRRRREEEGIQLRKQKREEQL--FKRRNV-----SLLSGEE--
A0A6I9X684 : --M-----DNMAS-P--EKD--NYRMKSYKNNALNPQEMRRRREEEGIQLRKQKREEQL--FKRRNV-----SLLSSEE--
V8NNB7     : --LFFIA-DNMAS-P--EKD--NYRMKSYKNNALNPQEMRRRREEEGIQLRKQKREEQL--FKRRNV-----SLLSSEE--
G1KPI9     : ------M-DTMAS-P--EKD--HYRMKSYKNNALNPQEMRRRREEEGIQLRKQKREEQL--FKRRNV-----ALLVNEE--
A0A6J0UAQ1 : ------M-DTMAS-P--EKD--HYRMKSYKNNALNPQEMRRRREEEGIQLRKQKREEQL--FKRRNV-----YLLGNEE--
A0A0R4I9E5 : ------M-DAMAS-P--GKD--NYRMKSYKNKALNPQEMRRRREEEGIQLRKQKREEQL--FKRRNV-----SLPPGDE--
A0A4X1TCK2 : ------M-NAMAS-P--GKD--NYRMKSYKNKALNPQEMRRRREEEGIQLRKQKREEQL--FKRRNV-----SLPKSDE--
A0A4X1TD58 : ------M-DAMAS-P--GKD--NYRMKSYKNKALNPQEMRRRREEEGIQLRKQKREEQL--FKRRNV-----SLPKSDE--
G1PF61     : ----------MAY-S--GKD--NYRMKSYKNKALNPQEMRRRREEEGIQLRKQKREEQL--FKRRNV-----SLPRSDE--
A0A2D0T8X7 : ------M-GAMAS-P--GKE--NYRMKSYKNKALNPQEMRRRREEEGIQLRKQKREEQL--FKRRNV-----SAPSCDE--
W5U8I8     : ------MEGAMAS-P--GKE--NYRMKSYKNKALNPQEMRRRREEEGIQLRKQKREEQL--FKRRNV-----SAPSCDE--
A0A4W4FB30 : ---VSWKFATMAS-P--GKD--NYRMKSYKNKALNPQEMRRRREEEGIQLRKQKREEQL--FKRRNV-----CVPSSDE--
A0A6P8W9H9 : ----------MAS-P--GKD--TFRMKSYKNKALIPQEMRRRREEEGIQLRKQKREEQL--YKRRNV-----CVPSSDE--
A0A4W4FCA9 : --FLTT-EATMAS-P--GKD--NYRMKSYKNKALNPQEMRRRREEEGIQLRKQKREEQL--FKRRNV-----CVPSSDE--
A0A4W4FCE0 : ---MS--EATMAS-P--GKD--NYRMKSYKNKALNPQEMRRRREEEGIQLRKQKREEQL--FKRRNV-----CVPSSDE--
A0A6P7JY08 : ----------MAS-P--GKD--NYRMKSYKNKALNPQEMRRRREEEGIQLRKQKREEQL--FKRRNV-----CVPSTDE--
A0A4W4FAY0 : ---SENPVRKTPV-P--GKD--NYRMKSYKNKALNPQEMRRRREEEGIQLRKQKREEQL--FKRRNV-----CVPSSDE--
A0A4W4FB35 : -------L----I-P--GKD--NYRMKSYKNKALNPQEMRRRREEEGIQLRKQKREEQL--FKRRNV-----CVPSSDE--
A0A4W4F9E2 : ---LK--VMLSFL-P--GKD--NYRMKSYKNKALNPQEMRRRREEEGIQLRKQKREEQL--FKRRNV-----CVPSSDE--
A0A4W4F8U5 : ---AQSHYSETVL-P--GKD--NYRMKSYKNKALNPQEMRRRREEEGIQLRKQKREEQL--FKRRNV-----CVPSSDE--
A0A673B1Y4 : --FSPSHY------P--GKD--SYRMKSYKNKALNPQEMRRRREEEGIQLRKQKREEQL--FKRRNV-----CVPSNDE--
A0A6J2W6F1 : ----------MAS-P--GKD--NYRMKSYKNKALNPQEMRRRREEEGIQLRKQKREEQL--FKRRNV-----CMPSTDE--
A0A665TQL2 : -KHLSS-Y-AMAS-P--GKD--SYRMRNYKNKALNPQEMRRRREEEGIQLRKQKREEQL--FKRRNV-----CVPSNDE--
A0A665T7A5 : ---VD----AMAS-P--GKD--SYRMRNYKNKALNPQEMRRRREEEGIQLRKQKREEQL--FKRRNV-----CVPSNDE--
A0A673B4D3 : ----------MAS-P--GKD--SYRMKSYKNKALNPQEMRRRREEEGIQLRKQKREEQL--FKRRNV-----CVPSNDE--
```

```
                      *        20         *        40         *        60         *        80
A0A6P7KQ24 : ----------MAS-L--GKD--NYRMRSYKNKALNPQEMRRRREEEGIQLRKQKREEQL--FKRRNV-----GVPTNDE--
A0A2K6FAB4 : --LMLNS-NAMAS-P--GKD--NYRMKSYKNKALNPQEMRRRREEEGIQLRKQKREEQVGVFKYLIL-----LYLGS----
A0A2K6FAC6 : ----VKSHNAMAS-P--GKD--NYRMKSYKNKALNPQEMRRRREEEGIQLRKQKREEQVGVFKYLIL-----LYLGS----
A0A2K6FAD0 : ----------MAS-P--GKD--NYRMKSYKNKALNPQEMRRRREEEGIQLRKQKREEQVGVFKYLIL-----VIFCA----
A0A2K6FAA6 : ----------MAS-P--GKD--NYRMKSYKNKALNPQEMRRRREEEGIQLRKQKREEQVGVFKYLIFRNENSLLCNS----
A0A665T652 : ----------MAS-P--GKD--SYRMRNYKNKALNPQEMRRRREEEGIQLRKQKREEQV---KYLTS------FKSSLNIY
A0A2I2Y6F3 : -------MDAMAS-P--GKD--NYRMKSYKNKALNPQEMRRRREEEGIQLRKQKREEQ---EEVVTT-DMVQMIFSN-NAD
A0A5F4W540 : -------MDAMAS-P--GKD--NYRMKSYKNKALNPQEMRRRREEEGIQLRKQKREEQ---EEVVTT-DMVQMIFSN-NAD
A0A3Q7NFQ8 : -------MDAMAS-P--GKD--NYRMKSYKNKALNPQEMRRRREEEGIQLRKQKREEQ---EEVITT-DMVQMIFSN-NAD
A0A452RB58 : ----------MAS-P--GKD--NYRMKSYKNKALNPQEMRRRREEEGIQLRKQKREEQVG-EEVITT-DMVQMI-------
A0A452TA68 : ----------MAS-P--GKD--NYRMKSYKNKALNPQEMRRRREEEGIQLRKQKREEQVG-EEVITT-DMVQMI-------
A0A665TF58 : ---IHRLFDAMAS-P--GKD--SYRMRNYKNKALNPQEMRRRREEEGIQLRKQKREEQL--FKRRNV-----RLTEQTQ--
A0A1A7XYL4 : ---------TMAS-P--KKD--NYRMKSYKNKALNPQEMRRRREEEGIQLRKQKREEQL--FKRRNV-----SVDCPIQ--
A0A1A8AG92 : ---------MMAS-P--KKD--NYRMKSYKNKALNSQEMRRRREEEGIQLRKQKREEQL--FKRRNV-----SVPSADE--
A0A1A8NLB0 : ---VLNSSDMMAS-P--KKD--NYRMKSYKNKALNSQEMRRRREEEGIQLRKQKREEQL--FKRRNV-----SVPSADE--
A0A1A8S747 : ---------MMAS-P--KKD--NYRMKSYKNKALNSQEMRRRREEEGIQLRKQKREEQL--FKRRNV-----SVPSADE--
A0A1A8CN21 : ---------MMAS-P--KKD--NYRMKSYKNKALNSQEMRRRREEEGIQLRKQKREEQL--FKRRNV-----SVPSADE--
A0A1A8KCW0 : ---------MMAS-P--KKD--NYRMKSYKNKALNSQEMRRRREEEGIQLRKQKREEQL--FKRRNV-----SVPSADE--
A0A1A8H282 : ---------MMAS-P--KKD--NYRMKSYKNKALNSQEMRRRREEEGIQLRKQKREEQL--FKRRNV-----SVPSADE--
A0A2P8YN66 : ----------MSG-P---TH--KYR---YKNVGLDSQELRRRREEEGVQLRKQKREQQL--FKRRNV-----FMPVLPD--
A0A4Y2LEL5 : ----------MSA----------HRFR-YKNIGLDAQELRRRREEEGVQLRKQKRESEL--YKRRNL-----SSPNENG--
A0A0B2V5K4 : EGLLPSHADAL-S-P--SAQ----REAMYKNKGMTSVEMRKRRETNTAQLRKQKRESEI--SKRRNID----QNPDSS---
A0A5B6UCM2 : ----------MPLRPSNTRKEVEVRKKGYKTLVLDGEEARRRREDNLVVIRKSKRENSL--LKKRKD-----GFFVINH--
A0A5B6VL98 : ----------MSLRP-NAR--AEVRRNRYKVAV-DAEEGRRRREDNMVEIRKNKREENL--LKKRRE-----GLLSQQQ--
cIBB5      : ------M-DAMAS-P--GKD--NYRMKSYKNKALNPQEMRRRREEEGIQLRKQKREEQL--FKRRNV-----SLPRNDE--
```

```
                   *        20         *        40         *        60         *        80
A0A6J0IB16 : --------------------M-------EN-YRMKSYKNKALNLEEMRRRREEEGIQLRKQKREQQVTVILSR-LCS--ALTQPR
A0A6J2H8S0 : --------------------M-------EN-YRMKSYKNKALNLEEMRRRREEEGIQLRKQKREQQVTVILSR-LCS--ALTQPR
A0A2U3ZLS2 : --------------------MASPGK--DN-YRMKSYKNNALNPEEMRRRREEEGIQLRKQKREQQESVI-TREMVE--MLFSDD
A0A452RMZ1 : --------------------MASPGK--DN-YRMKSYKNNALNPEEMRRRREEEGIQLRKQKREQQESVI-TREMVE--MLFSDD
Q6IP69     : -------------MDT----MASPGK--DN-SRMKSYKNKALNPEEMRRRREEEGIQLRKQKREQQ---LFKRRNVE--LAPEDT
Q7OPC4     : -------------MDT----MASPGK--DI-SRMKSYKNKALNPEEMRRRREEEGIQLRKQKREQQ---LFKRRNVE--LAPEDT
A0A6I8SII8 : ----ASNT-----LDT----MASPGK--EN-SRMKSFKNKALNPEEMRRRREEEGIQLRKQKREQQ---LFKRRNVE--LAPEDS
A0A6I8T171 : ------MK-----L-Y----IY-PGK--EN-SRMKSFKNKALNPEEMRRRREEEGIQLRKQKREQQ---LFKRRNVE--LAPEDS
A0A6I8SHV4 : ------VE-----GYT----MASPGK--EN-SRMKSFKNKALNPEEMRRRREEEGIQLRKQKREQQ---LFKRRNVE--LAPEDS
A0A4W3ITE9 : ------M-------ET----MASPAK--EN-YRMNRYKNKALNPEEMRRRREEEGIQLRKQKREQQ---LFKRRNVE--FINEES
A0A4W3ITH2 : ---AISLT------ET----MASPAK--EN-YRMNRYKNKALNPEEMRRRREEEGIQLRKQKREQQ---LFKRRNVE--FINEES
A0A6P7NY74 : ----------ME--PS-----ASPAK--DN-YRMNRYKNKALNPEEMRRRREEEGIQLRKQKREQQ---LFKRRNVD--ILNEEE
A0A673AGP4 : ------VCGQIE--PS-----ASPAK--DN-YRMNRYKNKALNPEEMRRRREEEGIQLRKQKREQQ---LFKRRNVD--ILNEEE
A0A673AFB1 : ----------ME--PS-----ASPAK--DN-YRMNRYKNKALNPEEMRRRREEEGIQLRKQKREQQ---LFKRRNVD--ILNEEE
A0A665UT43 : ------ICGPSE--PS-----ASPAK--DN-YRMNRYKNKALNPEEMRRRREEEGIQLRKQKREQQ---LFKRRNVD--ILNEEE
A0A665USF0 : ------M----E--PS-----ASPAK--DN-YRMNRYKNKALNPEEMRRRREEEGIQLRKQKREQQ---LFKRRNVD--ILNEEE
A0A6P7JV58 : ------M----E--PS-----ASPAK--DN-YRMNRYKNKALNPEEMRRRREEEGIQLRKQKREQQ---LFKRRNVD--ILNEEE
A0A665USW9 : ---------LDE--PS-----ASPAK--DN-YRMNRYKNKALNPEEMRRRREEEGIQLRKQKREQQ---LFKRRNVD--ILNEEE
A0A6P7JZ47 : ------SCRLSE--PS-----ASPAK--DN-YRMNRYKNKALNPEEMRRRREEEGIQLRKQKREQQ---LFKRRNVD--ILNEEE
A0A6P7NWW0 : ---------MAE--PS-----ASPAK--DN-YRMNRYKNKALNPEEMRRRREEEGIQLRKQKREQQ---LFKRRNVD--ILNEEE
A0A665USW8 : ------MRL-L---PS-----ASPAK--DN-YRMNRYKNKALNPEEMRRRREEEGIQLRKQKREQQ---LFKRRNVD--ILNEEE
A0A673AJ84 : ------MKS-V---PS-----ASPAK--DN-YRMNRYKNKALNPEEMRRRREEEGIQLRKQKREQQ---LFKRRNVD--ILNEEE
A0A665USJ2 : ------MN------PS-----ASPAK--DN-YRMNRYKNKALNPEEMRRRREEEGIQLRKQKREQQ---LFKRRNVD--ILNEEE
A0A669ENE7 : ------VGTDLQ--PS-----ASPAK--DN-YRMNRYKNKALNADEMRRRREEEGIQLRKQKREQQ---LFKRRNVD--ILNEEE
I3JLH4     : ----------ME--PS-----ASPAK--DN-YRMNRYKNKALNADEMRRRREEEGIQLRKQKREQQ---LFKRRNVD--ILNEEE
A0A669E488 : ----------MQE-PS-----ASPAK--DN-YRMNRYKNKALNADEMRRRREEEGIQLRKQKREQQ---LFKRRNVD--ILNEEE
A0A669F393 : ------FQ--LRCYPS-----ASPAK--DN-YRMNRYKNKALNADEMRRRREEEGIQLRKQKREQQ---LFKRRNVD--ILNEEE
A0A669BJX6 : ----------I------------PAK--DN-YRMNRYKNKALNADEMRRRREEEGIQLRKQKREQQ---LFKRRNVD--ILNEEE
A0A669C9I5 : ------VQT-L---PS-----ASPAK--DN-YRMNRYKNKALNADEMRRRREEEGIQLRKQKREQQ---LFKRRNVD--ILNEEE
A0A669CXP1 : ----LNYSSH----PS-----ASPAK--DN-YRMNRYKNKALNADEMRRRREEEGIQLRKQKREQQ---LFKRRNVD--ILNEEE
A0A669C1X0 : ------YSSHV---PS-----ASPAK--DN-YRMNRYKNKALNADEMRRRREEEGIQLRKQKREQQ---LFKRRNVD--ILNEEE
A0A665URM1 : ---NKYICS-----PS-----ASPAK--DN-YRMNRYKNKALNPEEMRRRREEEGIQLRKQKREQQ---LFKRRNVD--ILNEEE
A0A673AGC1 : ------ICS-----HS------SPAK--DN-YRMNRYKNKALNPEEMRRRREEEGIQLRKQKREQQ---LFKRRNVD--ILNEEE
A0A673AFG9 : -------CS-----GA------SPAK--DN-YRMNRYKNKALNPEEMRRRREEEGIQLRKQKREQQ---LFKRRNVD--ILNEEE
A0A665UTB0 : --------------M------SSPAK--DN-YRMNRYKNKALNPEEMRRRREEEGIQLRKQKREQQ---LFKRRNVD--ILNEEE
A0A673AGD2 : ------MFN-----MY-----SSPAK--DN-YRMNRYKNKALNPEEMRRRREEEGIQLRKQKREQQ---LFKRRNVD--ILNEEE
A0A673AJD9 : ------MTA-----Q------SSPAK--DN-YRMNRYKNKALNPEEMRRRREEEGIQLRKQKREQQ---LFKRRNVD--ILNEEE
A0A673AID3 : ------M-P-----T------SSPAK--DN-YRMNRYKNKALNPEEMRRRREEEGIQLRKQKREQQ---LFKRRNVD--ILNEEE
A0A674P4V4 : ------LKS-SD--TNY----SSPAK--DN-YRMNLYKNKALNPEEMRRRREEEGIQLRKQKREQQ---LFKRRNVD--VLNEEE
A0A674PD67 : ------L-------T-------CPAK--DN-YRMNLYKNKALNPEEMRRRREEEGIQLRKQKREQQ---LFKRRNVD--VLNEEE
A0A6J2QLQ7 : ---KTANK------PS-----ASPAK--DNNYRMNRYKNKALNPEEMRRRREEEGIQLRKQKREQQ---LFKRRNVD--VLNEEE
A0A2I4CRW0 : ------ME------PS-----ASPAK--DN-YRMNRYKNKALNPEEMRRRREEEGIQLRKQKREQQ---LFKRRNVD--VLNEEE
A0A3B3IGL5 : ------ME------PS-----ASPAK--DN-YRMNRYKNKALNPEEMRRRREEEGIQLRKQKREQQ---LFKRRNVD--VLNEEE
A0A4Z2H8L4 : ------ME------PS-----ASPAK--DN-YRMNRYKNKALNPEEMRRRREEEGVQLRKQKREQQ---LFKRRNVD--VLNEEE
A0A1A8RFX3 : ------ME------PS-----TSPAK--DN-YRMNRYKNKALNPEEMRRRREEEGIQLRKQKREQQ---LFKRRNVD--VLNEEE
A0A1A8KQY4 : ------ME------PS-----TSPAK--DN-YRMNRYKNKALNPEEMRRRREEEGIQLRKQKREQQ---LFKRRNVD--VLNEEE
A0A1A8MC56 : ------ME------PS-----TSPAK--DN-YRMNRYKNKALNPEEMRRRREEEGIQLRKQKREQQ---LFKRRNVD--VLNEEE
A0A1A8AWK2 : ------ME------PS-----TSPAK--DN-YRMNRYKNKALNPEEMRRRREEEGIQLRKQKREQQ---LFKRRNVD--VLNEEE
A0A1A8D3J2 : ------ME------PS-----TSPAK--DN-YRMNRYKNKALNPEEMRRRREEEGIQLRKQKREQQ---LFKRRNVD--VLNEEE
A0A1A8FMT4 : ------ME------PS-----TSPAK--DN-YRMNRYKNKALNPEEMRRRREEEGIQLRKQKREEQ---LFKRRNVD--VLNEEE
A0A674N9Z8 : --NEHISE------PS-----ASPAK--DN-YRMNLYKNKALNPEEMRRRREEEGIQLRKQKREQQ---LFKRRNVD--VLNEEE
A0A674NDE2 : -----M-E------PS-----ASPAK--DN-YRMNLYKNKALNPEEMRRRREEEGIQLRKQKREQQ---LFKRRNVD--VLNEEE
H2TLL4     : -----MAE------PS-----ASPAK--DN-YRMNLYKNKALNPEEMRRRREEEGIQLRKQKREQQ---LFKRRNVD--VLNEEE
A0A674MZC1 : --ISYLSE------PS-----ASPAK--DN-YRMNLYKNKALNPEEMRRRREEEGIQLRKQKREQQ---LFKRRNVD--VLNEEE
A0A674N3Q7 : -----LDN------PS-----ASPAK--DN-YRMNLYKNKALNPEEMRRRREEEGIQLRKQKREQQ---LFKRRNVD--VLNEEE
A0A674PHN0 : -----FD-------PS-----ASPAK--DN-YRMNLYKNKALNPEEMRRRREEEGIQLRKQKREQQ---LFKRRNVD--VLNEEE
A0A1A7WS65 : ------ME------PS-----ASPAK--DN-YRMNRYKNKALNPEEMRRRREEEGIQLRKQKREQQ---LFKRRNVD--VLNEVV
H2MGF0     : ------MR------TE-----ASPAK--DN-YRMNRYKNKALNPEEMRRRREEEGIQLRKQKREQQ---LFKRRNVD--VLNEEE
A0A674PL91 : ------M--------------ASPAK--DN-YRMNLYKNKALNPEEMRRRREEEGIQLRKQKREQQ---LFKRRNVD--VLNEEE
A0A667ZAA2 : ------LSC-----VGSD--LSSPAK--DN-YRMNRYKNKALNPEEMRRRREEEGIQLRKQKREQQ---LFKRRNVD--ILNEEA
A0A667YPY9 : --------------------MSSPAK--DN-YRMNRYKNKALNPEEMRRRREEEGIQLRKQKREQQ---LFKRRNVD--ILNEEA
A0A667ZA63 : ------MRV-----V-----VLIPAK--DN-YRMNRYKNKALNPEEMRRRREEEGIQLRKQKREQQ---LFKRRNVD--ILNEEA
A0A667YHP8 : ------YK----------------AY--DN-YRMNRYKNKALNPEEMRRRREEEGIQLRKQKREQQ---LFKRRNVD--ILNEEA
```

```
                           *        20         *        40         *        60         *        80
A0A667YPD2 : ------QTG-----H-----ITSPAK--DN-YRMNRYKNKALNPEEMRRRREEEGIQLRKQKREQQ---LFKRRNVD--ILNEEA
A0A667Z1Z9 : ------L-E-----P-----SASPAK--DN-YRMNRYKNKALNPEEMRRRREEEGIQLRKQKREQQ---LFKRRNVD--ILNEEA
A0A667Z1G5 : ------IEE-----P-----SASPAK--DN-YRMNRYKNKALNPEEMRRRREEEGIQLRKQKREQQ---LFKRRNVD--ILNEEA
A0A667Z113 : ------M-E-----P-----SASPAK--DN-YRMNRYKNKALNPEEMRRRREEEGIQLRKQKREQQ---LFKRRNVD--ILNEEA
E7F7U9     : ------ME------PS-----GSPAK--DN-FRMNRYKNKALNLEEMRRRREEEGIQLRKQKREQQ---LFKRRNVE--LLGEEG
A0A2D0S2W3 : ------ME------PS-----ASPAK--DN-YRMNRYKNKALNLEEMRRRREEEGIQLRKQKREQQ---LFKRRNVE--LLNEEG
A0A6J2WLL7 : ------ME------PA-----ASPAK--DN-YRMNRYKNKALNLEEMRRRREEEGIQLRKQKREQQ---LFKRRNVE--MLNEEA
A0A6P3VJS0 : ------MD------TA-----ASPAK--DN-YRMNRYKNKALNLEEMRRRREEEGIQLRKQKRELQ---LFKRRNVE--LLNEEA
H0YT19     : --------------------M-------EN-YRMKSYKNKALNPEEMRRRREEEGIQLRKQKREQQ---LFKRRNVE--LINEDD
A0A218UGK2 : --------------------M-------EN-YRMKSYKNKALNPEEMRRRREEEGIQLRKQKREQQ---LFKRRNVE--LINEDD
A0A6J0IB21 : --------------------M-------EN-YRMKSYKNKALNLEEMRRRREEEGIQLRKQKREQQ---LFKRRNVE--LINEDD
A0A6J2H9B6 : --------------------M-------EN-YRMKSYKNKALNLEEMRRRREEEGIQLRKQKREQQ---LFKRRNVE--LINEDD
A0A670Y1W5 : -----CGSS-----ET----MASPAK--ENNYRMKSYKNKALNPEEMRRRREEEGIQLRKQKREQQ---LFKRRNVE--LINEDA
V8NCM0     : --------------------MASPAK--ENNYRMKSYKNKALNPEEMRRRREEEGIQLRKQKREQQ---LFKRRNVE--LINEDA
A0A6J0U645 : -----SYLS-----ET----MASPAK--ENNYRMKSYKNKALNPEEMRRRREEEGIQLRKQKREQQ---LFKRRNVE--LINEDA
A0A6P9BDZ8 : --------M-----ET----MASPAK--ENNYRMKSYKNKALNPEEMRRRREEEGIQLRKQKREQQ---LFKRRNVE--LINEDA
H9G9G7     : --------M-----ET----MASPAK--ENNYRMKSYKNKALNPEEMRRRREEEGIQLRKQKREQQ---LFKRRNVE--LINEDA
A0A6J0U8D3 : --------M-----ET----MASPAK--ENNYRMKSYKNKALNPEEMRRRREEEGIQLRKQKREQQ---LFKRRNVE--LINEDA
A0A6I9Y5G6 : --------M-----ET----MASPAK--ENNYRMKSYKNKALNPEEMRRRREEEGIQLRKQKREQQ---LFKRRNVE--LINEDA
A0A6J1UEW1 : --------M-----ET----MASPAK--ENNYRMKSYKNKALNPEEMRRRREEEGIQLRKQKREQQ---LFKRRNVE--LINEDA
A0A670Y8U2 : --------M-----ET----MASPAK--ENNYRMKSYKNKALNPEEMRRRREEEGIQLRKQKREQQ---LFKRRNVE--LINEDA
A0A670JX51 : --------M-----ET----MASPAK--ENNYRMKSYKNKALNPEEMRRRREEEGIQLRKQKREQQ---LFKRRNVE--LINEDA
A0A2I0LNF2 : ----FFLFT-----ET----NTSPAK--DN-YRMKSYKNKALNPEEMRRRREEEGIQLRKQKREQQ---LFKRRNVE--LINEDA
A0A1V4JIW7 : --------M-----ET----NTSPAK--DN-YRMKSYKNKALNPEEMRRRREEEGIQLRKQKREQQ---LFKRRNVE--LINEDA
A0A669Q4K7 : ----RMLRA-----DS----MASPAK--DN-YRMKCYKNKALNPEEMRRRREEEGIQLRKQKREQQ---LFKRRNVE--LINEDA
A0A669PNE7 : -----M--------DS----MASPAK--DN-YRMKCYKNKALNPEEMRRRREEEGIQLRKQKREQQ---LFKRRNVE--LINEDA
F7B5S0     : --------M-----DS----MASPAK--DN-YRMKCYKNKALNPEEMRRRREEEGIQLRKQKREQQ---LFKRRNVE--LINEDA
G1N050     : ----FFLFT-----DS----MASPAK--DN-YRMKCYKNKALNPEEMRRRREEEGIQLRKQKREQQ---LFKRRNVE--LINEDA
A0A663MVT2 : --------------------MASPAK--DN-YRMKSYKNKALNPEEMRRRREEEGIQLRKQKREQQ---LFKRRNVE--LINEDA
A0A3Q0G435 : ----CFTGT-----KT----MASPAK--DN-YRMKSYKNKALNPEEMRRRREEEGIQLRKQKREQQ---LFKRRNVE--LINEDA
A0A674I4Z4 : --------M-----ET----MASPAKDKDN-YRMKSYKNKALNPEEMRRRREEEGIQLRKQKREQQ---LFKRRNVE--LINEDA
K7FQU4     : ------PFI-----ET----MASPAKDKDN-YRMKSYKNKALNPEEMRRRREEEGIQLRKQKREQQ---LFKRRNVE--LINEDA
A0A452IZ11 : --------M-----ET----MASPAKDKDN-YRMKSYKNKALNPEEMRRRREEEGIQLRKQKREQQ---LFKRRNVE--LINEDA
K7FQT8     : -------QVG----ET----MASPAKDKDN-YRMKSYKNKALNPEEMRRRREEEGIQLRKQKREQQ---LFKRRNVE--LINEDA
A0A672V4I2 : --------M-----ET----MASPAK--DN-YRMKSYKNKALNPEEMRRRREEEGIQLRKQKREQQ---LFKRRNVE--LINEDA
A0A663EJC7 : --------M-----ET----MASPAK--DN-YRMKSYKNKALNPEEMRRRREEEGIQLRKQKREQQ---LFKRRNVE--LINEDA
A0A663EJD2 : ----ACSSQ-----ET----MASPAK--DN-YRMKSYKNKALNPEEMRRRREEEGIQLRKQKREQQ---LFKRRNVE--LINEDA
A0A0G2JZS1 : ------M-------ET----MASPGK--DN-YRMKSYKNNALNPEEMRRRREEEGIQLRKQKREQQ---LFKRRNVE--LINEEA
A0A6J2DI60 : --YAAVT-------ET----MASPGK--DN-YRMKSYKNNALNPEEMRRRREEEGIQLRKQKREQQ---LFKRRNVE--LINEEA
A0A1S3FHK2 : ------M-------ET----MASPGK--DN-YRMKSYKNNALNPEEMRRRREEEGIQLRKQKREQQ---LFKRRNVE--LINEEA
A0A6P5DAY8 : ------M-------ET----MASPGK--DN-YRMKSYKNNALNPEEMRRRREEEGIQLRKQKREQQ---LFKRRNVE--LINEEA
A0A6J0XQK1 : ------M-------ET----MASPGK--DN-YRMKSYKNNALNPEEMRRRREEEGIQLRKQKREQQ---LFKRRNVE--LINEEA
A0A2Y9GL70 : ------M-------ET----MASPGK--DN-YRMKSYKNNALNPEEMRRRREEEGIQLRKQKREQQ---LFKRRNVE--LINEEA
A0A6J2DI93 : ------M-------ET----MASPGK--DN-YRMKSYKNNALNPEEMRRRREEEGIQLRKQKREQQ---LFKRRNVE--LINEEA
A0A2K6EGT9 : ------M-------ET----MASPGK--DN-YRMKSYKNNALNPEEMRRRREEEGIQLRKQKREQQ---LFKRRNVE--LINEEA
A0A6J2DEH4 : ----RSLGQ-----ET----MASPGK--DN-YRMKSYKNNALNPEEMRRRREEEGIQLRKQKREQQ---LFKRRNVE--LINEEA
A0A3Q7MUA7 : ----RNLGQ-----ET----MASPGK--DN-YRMKSYKNNALNPEEMRRRREEEGIQLRKQKREQQ---LFKRRNVE--LINEEA
O60684     : ------M-------ET----MASPGK--DN-YRMKSYKNNALNPEEMRRRREEEGIQLRKQKREQQ---LFKRRNVE--LINEEA
A0A2Y9NNA2 : ------M-------ET----MASPGK--DN-YRMKSYKNNALNPEEMRRRREEEGIQLRKQKREQQ---LFKRRNVE--LINEEA
A0A2I2YR04 : ------M-------ET----MASPGK--DN-YRMKSYKNNALNPEEMRRRREEEGIQLRKQKREQQ---LFKRRNVE--LINEEA
A0A3Q7UAJ7 : ------M-------ET----MASPGK--DN-YRMKSYKNNALNPEEMRRRREEEGIQLRKQKREQQ---LFKRRNVE--LINEEA
A0A2Y9FJU4 : ------M-------ET----MASPGK--DN-YRMKSYKNNALNPEEMRRRREEEGIQLRKQKREQQ---LFKRRNVE--LINEEA
A0A2R8ZHT3 : ------M-------ET----MASPGK--DN-YRMKSYKNNALNPEEMRRRREEEGIQLRKQKREQQ---LFKRRNVE--LINEEA
A0A2K6E6T2 : ------M-------ET----MASPGK--DN-YRMKSYKNNALNPEEMRRRREEEGIQLRKQKREQQ---LFKRRNVE--LINEEA
W5NR48     : ---SSLLP------ET----MASPGK--DN-YRMKSYKNNALNPEEMRRRREEEGIQLRKQKREQQ---LFKRRNVE--LINEEA
A0A383Z6H8 : ---SSLLP------ET----MASPGK--DN-YRMKSYKNNALNPEEMRRRREEEGIQLRKQKREQQ---LFKRRNVE--LINEEA
A0A340YBD4 : ---SPLLP------ET----MASPGK--DN-YRMKSYKNNALNPEEMRRRREEEGIQLRKQKREQQ---LFKRRNVE--LINEEA
A0A6I9M8L4 : ------M-------ET----MASPGK--DN-YRMKSYKNNALNPEEMRRRREEEGIQLRKQKREQQ---LFKRRNVE--LINEEA
A0A6I9HXL5 : ------M-------ET----MASPGK--DN-YRMKSYKNNALNPEEMRRRREEEGIQLRKQKREQQ---LFKRRNVE--LINEEA
F1PM77     : ------M-------ET----MASPGK--DN-YRMKSYKNNALNPEEMRRRREEEGIQLRKQKREQQ---LFKRRNVE--LINEEA
A0A384CCX3 : ------M-------ET----MASPGK--DN-YRMKSYKNNALNPEEMRRRREEEGIQLRKQKREQQ---LFKRRNVE--LINEEA
A0A340Y669 : ------M-------ET----MASPGK--DN-YRMKSYKNNALNPEEMRRRREEEGIQLRKQKREQQ---LFKRRNVE--LINEEA
A0A3Q7P798 : ------M-------ET----MASPGK--DN-YRMKSYKNNALNPEEMRRRREEEGIQLRKQKREQQ---LFKRRNVE--LINEEA
```

```
                       *        20         *        40         *        60         *        80
A0A286XBY1 : ------M-------ET----MASPGK--DN-YRMKSYKNNALNPEEMRRRREEEGIQLRKQKREQQ---LFKRRNVE--LINEEA
A0A455BFC9 : -----LMHN----KET----MASPGK--DN-YRMKSYKNNALNPEEMRRRREEEGIQLRKQKREQQ---LFKRRNVE--LINEEA
A0A2Y9NH87 : -----LMHN----KET----MASPGK--DN-YRMKSYKNNALNPEEMRRRREEEGIQLRKQKREQQ---LFKRRNVE--LINEEA
A0A3Q0D5Z9 : ------M-------ET----MASPGK--DN-YRMKSYKNNALNPEEMRRRREEEGIQLRKQKREQQ---LFKRRNVE--LINEEA
A0A673TMV9 : ------M-------ET----MASPGK--DN-YRMKSYKNNALNPEEMRRRREEEGIQLRKQKREQQ---LFKRRNVE--LINEEA
A0A2K5X9F2 : ------M-------ET----MASPGK--DN-YRMKSYKNNALNPEEMRRRREEEGIQLRKQKREQQ---LFKRRNVE--LINEEA
H9EN17     : ------M-------ET----MASPGK--DN-YRMKSYKNNALNPEEMRRRREEEGIQLRKQKREQQ---LFKRRNVE--LINEEA
H2N861     : ------M-------ET----MASPGK--DN-YRMKSYKNNALNPEEMRRRREEEGIQLRKQKREQQ---LFKRRNVE--LINEEA
A0A3Q7X5I0 : ------M-------ET----MASPGK--DN-YRMKSYKNNALNPEEMRRRREEEGIQLRKQKREQQ---LFKRRNVE--LINEEA
I3MIF0     : ------M-------ET----MASPGK--DN-YRMKSYKNNALNPEEMRRRREEEGIQLRKQKREQQ---LFKRRNVE--LINEEA
A0A6J2DCS8 : --PCDHK-------ET----MASPGK--DN-YRMKSYKNNALNPEEMRRRREEEGIQLRKQKREQQ---LFKRRNVE--LINEEA
A0A0D9S7X9 : ------M-------ET----MASPGK--DN-YRMKSYKNNALNPEEMRRRREEEGIQLRKQKREQQ---LFKRRNVE--LINEEA
A0A6J3FAF5 : ------M-------ET----MASPGK--DN-YRMKSYKNNALNPEEMRRRREEEGIQLRKQKREQQ---LFKRRNVE--LINEEA
A0A6J0A060 : ------M-------ET----MASPGK--DN-YRMKSYKNNALNPEEMRRRREEEGIQLRKQKREQQ---LFKRRNVE--LINEEA
F7FPD9     : ------M-------ET----MASPGK--DN-YRMKSYKNNALNPEEMRRRREEEGIQLRKQKREQQ---LFKRRNVE--LINEEA
A0A2K6TJD1 : ------M-------ET----MASPGK--DN-YRMKSYKNNALNPEEMRRRREEEGIQLRKQKREQQ---LFKRRNVE--LINEEA
A0A383Z655 : ------M-------ET----MASPGK--DN-YRMKSYKNNALNPEEMRRRREEEGIQLRKQKREQQ---LFKRRNVE--LINEEA
Q8BH30     : -----M--------ET----MASPGK--DN-YRMKSYKNNALNPEEMRRRREEEGIQLRKQKREQQ---LFKRRNVE--LINEEA
G3TI14     : --PSFFP-------ET----MASPGK--DN-YRMKSYKNNALNPEEMRRRREEEGIQLRKQKREQQ---LFKRRNVE--PINEEA
F6ZAL1     : -----M--------ET----MASPGK--DN-YRMKSYKNNALNPEEMRRRREEEGIQLRKQKREQQ---LFKRRNVE--LINEEA
A0A667GXC7 : -----M--------ET----MASPGK--DN-YRMKSYKNNALNPEEMRRRREEEGIQLRKQKREQQ---LFKRRNVE--LINEEA
A0A096N2D7 : -----M--------ET----MASPGK--DN-YRMKSYKNNALNPEEMRRRREEEGIQLRKQKREQQ---LFKRRNVE--LINEEA
H2PYJ7     : -----M--------ET----MASPGK--DN-YRMKSYKNNALNPEEMRRRREEEGIQLRKQKREQQ---LFKRRNVE--LINEEA
G1PWY7     : ------M-------ET----MASPGK--DN-YRMKSYKNNALNPEEMRRRREEEGIQLRKQKREQQ---LFKRRNVE--MINEEA
A0A671F6J5 : -SG-AAM-------ET----MASPGK--DN-YRMKSYKNNALNPEEMRRRREEEGIQLRKQKREQQ---LFKRRNVE--MINEEA
A0A6P3Q105 : ------M-------ET----MASPGK--DN-YRMKSYKNNALNPEEMRRRREEEGIQLRKQKREQQ---LFKRRNVE--MINEEA
A0A6J2LMK8 : ------M-------ET----MASPGK--DN-YRMKSYKNNALNPEEMRRRREEEGIQLRKQKREQQ---LFKRRNVE--MINEEA
A0A6P6CCF2 : NFP--NR-------ET----MASPGK--DN-YRMKSYKNNALNPEEMRRRREEEGIQLRKQKREQQ---LFKRRNVE--MINEEA
A0A6P6CCM1 : -HP-CNK-------ET----MASPGK--DN-YRMKSYKNNALNPEEMRRRREEEGIQLRKQKREQQ---LFKRRNVE--MINEEA
M3Z9Q7     : ------M-------ET----MASPGK--DN-YRMKSYKNNALNPEEMRRRREEEGIQLRKQKREQQ---LFKRRNVE--LINEEA
G1TE61     : ------M-------ET----MASPGK--DN-YRMKSYKNNALNPEEMRRRREEEGIQLRKQKREQQ---LFKRRNVE--LINEEA
A0A2K5RW26 : ------M-------ET----MASPGK--DN-YRMKSYKNNALNPEEMRRRREEEGIQLRKQKREQQ---LFKRRNVE--LINEEA
A0A4W2FQX2 : ------M-------ET----MASPGK--DN-YRMKSYKNNALNPEEMRRRREEEGIQLRKQKREQQ---LFKRRNVE--LINEEA
A0A2U4BKP8 : ------M-------ET----MASPGK--DN-YRMKSYKNNALNPEEMRRRREEEGIQLRKQKREQQ---LFKRRNVE--LINEEA
A0A6P4UL29 : ------M-------ET----MASPGK--DN-YRMKSYKNNALNPEEMRRRREEEGIQLRKQKREQQ---LFKRRNVE--LINEEA
Q3KR98     : ---KNYVS------ET----MASPGK--DN-YRMKSYKNNALNPEEMRRRREEEGIQLRKQKREQQ---LFKRRNVE--LINEEA
A0A6J0CKI2 : ---KNYVS------ET----MASPGK--DN-YRMKSYKNNALNPEEMRRRREEEGIQLRKQKREQQ---LFKRRNVE--LINEEA
A0A6P5PQG2 : ---KNYVA------ET----MASPGK--DN-YRMKSYKNNALNPEEMRRRREEEGIQLRKQKREQQ---LFKRRNVE--LINEEA
F1LT58     : ---KTMVS------ET----MASPGK--DN-YRMKSYKNNALNPEEMRRRREEEGIQLRKQKREQQ---LFKRRNVE--LINEEA
A0A673U0E9 : -----M-S------ET----MASPGK--DN-YRMKSYKNNALNPEEMRRRREEEGIQLRKQKREQQ---LFKRRNVE--LINEEA
A0A6J2DI83 : ---KNHVS------ET----MASPGK--DN-YRMKSYKNNALNPEEMRRRREEEGIQLRKQKREQQ---LFKRRNVE--LINEEA
A0A3Q0EEC2 : ---KNHVS------ET----MASPGK--DN-YRMKSYKNNALNPEEMRRRREEEGIQLRKQKREQQ---LFKRRNVE--LINEEP
A0A6J0XSR8 : ---KNHVS------ET----MASPGK--DN-YRMKSYKNNALNPEEMRRRREEEGIQLRKQKREQQ---LFKRRNVE--LINEEA
A0A5G2QUV4 : ---KNHVS------ET----MASPGK--DN-YRMKSYKNNALNPEEMRRRREEEGIQLRKQKREQQ---LFKRRNVE--LINEEA
A0A340Y4V3 : ---KNHVS------ET----MASPGK--DN-YRMKSYKNNALNPEEMRRRREEEGIQLRKQKREQQ---LFKRRNVE--LINEEA
A0A3Q7MWD7 : ---KNHVS------ET----MASPGK--DN-YRMKSYKNNALNPEEMRRRREEEGIQLRKQKREQQ---LFKRRNVE--LINEEA
A0A6P4UDT9 : ---KNHVS------ET----MASPGK--DN-YRMKSYKNNALNPEEMRRRREEEGIQLRKQKREQQ---LFKRRNVE--LINEEA
A0A6J0A0G5 : ---RNHVS------ET----MASPGK--DN-YRMKSYKNNALNPEEMRRRREEEGIQLRKQKREQQ---LFKRRNVE--LINEEA
A0A1D5QJS1 : ---KSHVS------ET----MASPGK--DN-YRMKSYKNNALNPEEMRRRREEEGIQLRKQKREQQ---LFKRRNVE--LINEEA
A0A2K5RVX9 : ---KSHVS------ET----MASPGK--DN-YRMKSYKNNALNPEEMRRRREEEGIQLRKQKREQQ---LFKRRNVE--LINEEA
A0A1S3AP65 : ---KNHMS------ET----MASPGK--DN-YRMKSYKNNALNPEEMRRRREEEGIQLRKQKREQQ---LFKRRNVE--LINDEA
A0A1S3APL8 : ------M-------ET----MASPGK--DN-YRMKSYKNNALNPEEMRRRREEEGIQLRKQKREQQ---LFKRRNVE--LINDEA
A0A341D6G7 : ------ME------ET----MASPGK--DN-YRMKSYKNNALNPEEMRRRREEEGIQLRKQKREQQ---LFKRRNVE--LINEEA
A0A6P5PM01 : ------M-------ET----MASPGK--DN-YRMKSYKNNALNPEEMRRRREEEGIQLRKQKREQQ---LFKRRNVE--LINEEA
A0A341D6Y7 : ------M-------ET----MASPGK--DN-YRMKSYKNNALNPEEMRRRREEEGIQLRKQKREQQ---LFKRRNVE--LINEEA
A0A6P3H7K5 : ------M-------ET----MASPGK--DN-YRMKSYKNNALNPEEMRRRREEEGIQLRKQKREQQ---LFKRRNVE--LINEEA
A0A7F8QX02 : ------M-------ET----MASPGK--DN-YRMKSYKNNALNPEEMRRRREEEGIQLRKQKREQQ---LFKRRNVE--LINEEA
A0A287AEI6 : ------M-------ET----MASPGK--DN-YRMKSYKNNALNPEEMRRRREEEGIQLRKQKREQQ---LFKRRNVE--LINEEA
A0A2K5K3W8 : ------M-------ET----MASPGK--DN-YRMKSYKNNALNPEEMRRRREEEGIQLRKQKREQQ---LFKRRNVE--LINEEA
Q0V7M0     : ------M-------ET----MASPGK--DN-YRMKSYKNNALNPEEMRRRREEEGIQLRKQKREQQ---LFKRRNVE--LINEEA
A0A2K6LP75 : ------M-------ET----MASPGK--DN-YRMKSYKNNALNPEEMRRRREEEGIQLRKQKREQQ---LFKRRNVE--LINEEA
A0A2K6AEI5 : ------M-------ET----MASPGK--DN-YRMKSYKNNALNPEEMRRRREEEGIQLRKQKREQQ---LFKRRNVE--LINEEA
A0A2I2V555 : ------M-------ET----MASPGK--DN-YRMKSYKNNALNPEEMRRRREEEGIQLRKQKREQQ---LFKRRNVE--LINEEA
```

```
                      *        20         *        40         *        60         *        80
A0A6P3T9P0 : ------M-------ET----MASPGK--DN-YRMKSYKNNALNPEEMRRRREEEGIQLRKQKREQQ---LFKRRNVE--LINEEA
A0A3Q0EAN1 : ---SHQV------AQT----MASPGK--DN-YRMKSYKNNALNPEEMRRRREEEGIQLRKQKREQQ---LFKRRNVE--LINEEP
A0A1U7TY64 : -------------MET----MASPGK--DN-YRMKSYKNNALNPEEMRRRREEEGIQLRKQKREQQ---LFKRRNVE--LINEEP
A0A6P6DFA1 : -------------MET----MASPGK--DN-YRMKSYKNNALNPEEMRRRREEEGIQLRKQKREQQ---LFKRRNVE--LINEET
A0A250YJI9 : ------M-------EP----MASPGK--DN-YRMKSYKNNALNPEEMRRRREEEGIQLRKQKREQQ---LFKRRNVE--LINEEA
A0A286X7C6 : ------L-------ET----MASPGK--DN-YRMKSYKNNALNPEEMRRRREEEGIQLRKQKREQQ---LFKRRNVE--LINEEA
G1LMT1     : --LSLFL-------ET----MASPGK--DN-YRMKSYKNNALNPEEMRRRREEEGIQLRKQKREQQ---LFKRRNVE--LINEEA
A0A2K6AEI0 : ------L-------ET----MASPGK--DN-YRMKSYKNNALNPEEMRRRREEEGIQLRKQKREQQ---LFKRRNVE--LINEEA
F7FPC0     : ------L-------ET----MASPGK--DN-YRMKSYKNNALNPEEMRRRREEEGIQLRKQKREQQ---LFKRRNVE--LINEEA
A0A2K5K3W0 : ---SFLL-------ET----MASPGK--DN-YRMKSYKNNALNPEEMRRRREEEGIQLRKQKREQQ---LFKRRNVE--LINEEA
A0A2K5RW11 : --VSFLL-------ET----MASPGK--DN-YRMKSYKNNALNPEEMRRRREEEGIQLRKQKREQQ---LFKRRNVE--LINEEA
H0X0L1     : --FSFLL-------ET----MASPGK--DN-YRMKSYKNNALNPEEMRRRREEEGIQLRKQKREQQ---LFKRRNVE--LINEEA
A0A4X1W0L9 : ------L-------ET----MASPGK--DN-YRMKSYKNNALNPEEMRRRREEEGIQLRKQKREQQ---LFKRRNVE--LINEEA
A0A6G1AYV8 : ------L-------ET----MASPGK--DN-YRMKSYKNNALNPEEMRRRREEEGIQLRKQKREQQ---LFKRRNVE--LINEEA
A0A2I3LV73 : ------L-------ET----MASPGK--DN-YRMKSYKNNALNPEEMRRRREEEGIQLRKQKREQQ---LFKRRNVE--LINEEA
A0A2I3RHW2 : ------L-------ET----MASPGK--DN-YRMKSYKNNALNPEEMRRRREEEGIQLRKQKREQQ---LFKRRNVE--LINEEA
A0A452RN34 : ------L-------ET----MASPGK--DN-YRMKSYKNNALNPEEMRRRREEEGIQLRKQKREQQ---LFKRRNVE--LINEEA
A0A2K6TJE2 : ------L-------ET----MASPGK--DN-YRMKSYKNNALNPEEMRRRREEEGIQLRKQKREQQ---LFKRRNVE--LINEEA
A0A2U4BKQ3 : --MT-TLG------ET----MASPGK--DN-YRMKSYKNNALNPEEMRRRREEEGIQLRKQKREQQ---LFKRRNVE--LINEEA
A0A2K5K3Y4 : ------LK-----SLT----MASPGK--DN-YRMKSYKNNALNPEEMRRRREEEGIQLRKQKREQQ---LFKRRNVE--LINEEA
A0A2K6AEJ5 : ------LK-----SLT----MASPGK--DN-YRMKSYKNNALNPEEMRRRREEEGIQLRKQKREQQ---LFKRRNVE--LINEEA
A0A2K5X913 : ------LK-----SLT----MASPGK--DN-YRMKSYKNNALNPEEMRRRREEEGIQLRKQKREQQ---LFKRRNVE--LINEEA
A0A2I3LJG3 : ------LK-----SLT----MASPGK--DN-YRMKSYKNNALNPEEMRRRREEEGIQLRKQKREQQ---LFKRRNVE--LINEEA
A0A452RMT7 : ------LR-------T----MASPGK--DN-YRMKSYKNNALNPEEMRRRREEEGIQLRKQKREQQ---LFKRRNVE--LINEEA
A0A2K6EGU4 : ------FKEK---NHT----MASPGK--DN-YRMKSYKNNALNPEEMRRRREEEGIQLRKQKREQQ---LFKRRNVE--LINEEA
A0A2K5X8Z3 : -MK---VK------GT----MASPGK--DN-YRMKSYKNNALNPEEMRRRREEEGIQLRKQKREQQ---LFKRRNVE--LINEEA
A0A2K5MF98 : -MK---VK------GT----MASPGK--DN-YRMKSYKNNALNPEEMRRRREEEGIQLRKQKREQQ---LFKRRNVE--LINEEA
A0A2I3MMA4 : -MK---VK------GT----MASPGK--DN-YRMKSYKNNALNPEEMRRRREEEGIQLRKQKREQQ---LFKRRNVE--LINEEA
A0A2K6AEF3 : --K---VKE----RKT----MASPGK--DN-YRMKSYKNNALNPEEMRRRREEEGIQLRKQKREQQ---LFKRRNVE--LINEEA
A0A452ETW3 : --------------------MASPGK--DN-YRMKSYKNNALNPEEMRRRREEEGIQLRKQKREQQ---LFKRRNVE--LINEEA
A0A2I3SZD9 : --KSNLIK-------T----MASPGK--DN-YRMKSYKNNALNPEEMRRRREEEGIQLRKQKREQQ---LFKRRNVE--LINEEA
A0A2R8ZNF5 : --KSNLIK-------T----MASPGK--DN-YRMKSYKNNALNPEEMRRRREEEGIQLRKQKREQQ---LFKRRNVE--LINEEA
A0A6I9JKJ0 : --------------------MASPGK--DN-YRMKSYKNNALNPEEMRRRREEEGIQLRKQKREQQ---LFKRRNVE--LINEEA
A0A6P6IQI4 : --------------------MASPGK--DN-YRMKSYKNNALNPEEMRRRREEEGIQLRKQKREQQ---LFKRRNVE--LINEEA
A0A2K5MFB8 : -----SLVS----TKT----MASPGK--DN-YRMKSYKNNALNPEEMRRRREEEGIQLRKQKREQQ---LFKRRNVE--LINEEA
A0A452RMQ4 : ------MKT----TST----MASPGK--DN-YRMKSYKNNALNPEEMRRRREEEGIQLRKQKREQQ---LFKRRNVE--LINEEA
F6SAJ5     : ------M-------ET----MASPGK--DN-YRMKSYKNKALNPEEMRRRREEEGIQLRKQKREQQ---LFKRRNVE--LINEDA
A0A6J3E8N2 : ------M-------ET----MASPGK--DN-YRMKSYKNKALNPEEMRRRREEEGIQLRKQKREQQ---LFKRRNVE--LINEDA
A0A6P5K253 : ------M-------ET----MASPGK--DN-YRMKSYKNKALNPEEMRRRREEEGIQLRKQKREQQ---LFKRRNVE--LISEDA
A0A5F8GFC7 : ------M-------ET----MASPGK--DN-YRMKSYKNKALNPEEMRRRREEEGIQLRKQKREQQ---LFKRRNVE--LISEDA
A0A4X2JNT1 : ------M-------ET----MASPGK--DN-YRMKSYKNKALNPEEMRRRREEEGIQLRKQKREQQ---LFKRRNVE--LISEDA
A0A6P5K276 : ---KKQLS------ET----MASPGK--DN-YRMKSYKNKALNPEEMRRRREEEGIQLRKQKREQQ---LFKRRNVE--LISEDA
F7E1F4     : ---KKQLS------ET----MASPGK--DN-YRMKSYKNKALNPEEMRRRREEEGIQLRKQKREQQ---LFKRRNVE--LISEDA
A0A4X2JN23 : ---KKQLS------ET----MASPGK--DN-YRMKSYKNKALNPEEMRRRREEEGIQLRKQKREQQ---LFKRRNVE--LISEDA
G3VZX7     : ---------------T----MASPGK--DN-YRMKSYKNKALNPEEMRRRREEEGIQLRKQKREQQ---LFKRRNVE--LISEDA
U3IY73     : --------------------MASPGK--DN-YRMKSYKNKALNPEEMRRRREEEGIQLRKQKREQQ---LFKRRNVE--LINEDA
A0A6P8S1I7 : ---QCEMG------DT----MASPGK--DN-YRMKSYKNKALNPEEMRRRREEEGIQLRKQKREQQ---LFKRRNVE--LITDET
A0A6P7ZKX7 : --------------------MASPGK--DN-YRMKSYKNKALNPEEMRRRREEEGIQLRKQKREQQ---LFKRRNVE--LITDDA
A0A6F9DGL9 : --------------------MAS--K--E---RMKIYKNKAKDQEELRRKREEEGIQLRKQKRDEQ---VFKRRNVT--LNNEEI
A0A2B4SYN2 : --------------------MAT--R--AN-ARLKSYKNKALDTAEMRRRREEEGVQLRKAKREEQ---MFKRRNVD--MTGKSS
A0A4C1U0C7 : ------MTHWTYLRSQ----MSGG-------HK-HRYKNAGLSADELRRRREEEGVQLRKQKREQQ---LFKRRNVNLPAPNA--
A0A0C9QJY4 : ------M-------S------SGPQ------HK-HRYKNVGLDAQELRRRREEEGVQLRKQKREQQ---LSKRRNV--PTIGTDD
A0A0A9XP06 : ------M-------S-----MTS--------HK-NRYKNAGLDSQELRRRREEEGIQLRKQKREQQ---LFKRRNVNIPIID-D-
A0A2S2R7M9 : --------------------MSNT-------HR-MRYKNEGVDSKELRRRREEEGIQLRKEKRDIQ---LFKRRNLN-EFIQ-ND
A0A2H8U0D0 : --------------------MSNT-------HR-MRYKNEGVDSKELRRRREEEGIQLRKQKRDIQ---LFKRRNIN-ETIQ-NE
A0A6A4WGB8 : ------MFN-IY-HSQHPVTMSS--------HR-HRYKNEGLDSAELRRRREEEGVQLRKQKRDRQ---LNKRRNVP-P-LQPAA
A0A6G1S6W7 : --------------------MSD--------HR-ARYKNTGLDAKELRRRREEDSVQLRKQKRDDV---LSKRRTLG-P-VSPRD
A0A0A1XSA1 : --------------------MSAT-------HR-QRYKNTPLDSTEMRRRREEEGVQIRKQKREQQ---LIKRRNVV-SEVH-QS
A0A0K8VAU4 : --------------------MSTT-------HR-HRYKNTPLDSTEMRRRREEEGVQIRKQKREQQ---LIKRRNVV-SEVQ-QS
T2MID0     : --------------AQ----MAT---------RINKYKNQSFDSQELRRRREEVGVQLRKTKRDEQ---LSKRRNVEVQGVE-N-
cIBB6      : ------M-------ET----MASPGK--DN-YRMKSYKNNALNPEEMRRRREEEGIQLRKQKREQQ---LFKRRNVE--LINEEA
```

```
                          *        20         *        40         *        60         *        80
A0A674JCE7 : --MADHYKMPIFW---SC----SITQNLPQSPTPNNGWHLQSLI-RQRMEVNVELRKAKKDEQI-LKRRN-IS-FSS-M-GE-
A0A674J8N5 : ---------M-IQWL--EV----KVRQ-I-Q--T-GNSVNLTAMR-RQRMEVNVELRKAKKDEQI-LKRRN-IS-FSS-M-GE-
A0A6J1VLK5 : ---------MSTTKE--QD-E--RMRK-F-K----NKGKDLAEIR-RRKIEVRVELRKARKDEQI-LKRRN-IC-FL--A-ESS
A0A670YIB3 : ---------MSTTKE--QD-E--HMRK-F-K----NKGKDIAEIR-RRKIEVRVELRKARKDEQI-LKRRN-IC-FS--A-ESS
A0A6P9DMK1 : ---------MPTTNE--QD-G--RMRK-F-K----NKGKNVAEIR-RRKIEVRVELRKARKDEQI-LKRRN-IC-SL--A-EAS
Q4QR45     : ---------MPTTNE--AD-E--RMRK-F-K----NKGKDTAELR-RRRVEVSVELRKAKKDEQI-LKRRN-VC-LP--E-ELI
A0A1L8EXD5 : ---------MPTTNE--AD-E--RMRK-F-K----NKGKDTAELR-RRRVEVSVELRKAKKDEQI-LKRRN-VC-FP--E-ELI
B4F6X3     : ---------MPTTNE--AD-E--RMRK-F-K----NKGKDTAELR-RRRVEVNVELRKAKKDEQI-LKRRN-VC-FP--E-ELV
A0A6J0V1C3 : ---------MPTTTE--QD-E--RMRK-F-K----NRGKDLAVMRVQRR-EVGVELRKARKDEQI-LKRRN-IC-FT--A-ENT
A0A452IQU3 : ---------MPTTNE--QD-E--RMKK-F-K----NKGKDTAAMR-RQRMEVNVELRKAKKDEQI-LKRRN-IS-FSSMG-E--
A0A1U7S409 : ---------MPGMNE--QD-E--RMRK-F-K----NKGKDAAAMR-RQRVEVNVELRKAKKDEQI-LKRRN-IT-IT-LE-DR-
A0A151N2K4 : ---------M-SANENAAAAA-ARLNR-F-K----NKGKDTTEMR-RRRIEVNVELRKAKKDDQM-LKRRN-VS-TL--P-DDA
A0A4W3JZ18 : ---------M-IAKMSTSEAENTRMKI-F-K----NKGRDFVELR-RRRIEINVELRKAKKDEQI-LKRRN-VG-TLR-E-DD-
A0A6P7YUC3 : ---------MPLANE--SE-E--RIKK-F-K----NKGKDTAELR-RRRVEVSVELRKARKDEQI-LKRRN-VI-SL--P-DEG
A0A6P8SI26 : ---------MPLASE--SE-E--RIKK-F-K----NKGKNTAELR-RRRMEVSVELRKARKDEQI-LKRRN-VI-SL--P-DES
A0A663F2M4 : -------AKMPVVNE--HS-E--RMRK-F-K----NKGKDAAALR-RQRVEVNIELRKAKKDEQI-LKRRS-IS-FSL-M-DE-
A0A663F2P6 : --------KMPVVNE--HS-E--RMRK-F-K----NKGKD--ALR-RQRVEVNIELRKAKKDEQI-LKRRS-IS-FSL-M-DE-
A0A663NCZ5 : -------AKMPVVGK--DS-E--RMKK-F-K----NKGKDAAALR-RQRVEVNIELRKAKKDEQI-LKRRS-IL-FRL-M-DD-
A0A6J3DSQ0 : ---------MPVVNE--PS-E--RMKK-F-K----NRGKDAAALR-RQRVEVKVELRKAQKDEQI-LKRRS-IS-ITL-M-EK-
E1BVP7     : -----PAKMPVVHE--HG-E--RMKR-F-K----NKGKDTA-LR-RQRVEVNIELRKAKKDEQI-LKRRS-IS-IVS-L-EK-
G1MY68     : ---------MPVVHE--HG-E--RMKR-F-K----NKGKDTATLR-RQRVEVNIELRKAKKDEQI-LKRRS-IS-IVS-L-EK-
A0A669Q915 : ---------MPVVHE--HG-E--RMKR-F-K----NKGKDAAALR-RQRVEVNIELRKAKKDEQI-LKRRS-IS-IVS-L-EK-
A0A3Q3B277 : ---------MPVVHE--HG-E--RMKR-F-K----NKGKDTA-LR-RQRVEVNIELRKAKKDEQI-LKRRS-IS-ISP-S-PA-
A0A2I0M0S3 : ---------MPVLAK--QS-E--RMRQ-F-K----NKGKNAADLR-RQRVEVNVELRKAKKDEQI-LKRRN-IS-FSL-M-EE-
A0A1V4KSF1 : ---------MPVLAK--QS-E--RMRQ-F-K----NKGKNAADLR-RQRVEVNVELRKAKKDEQI-LKRRN-IS-FSL-M-EE-
U3KHB8     : ---------MPV-RK--HS-Q--HMKL-F-K----NKGKDETTLR-RQRVEVSIELRKAKKDEQI-LKRRN-IS-IQL-K-EE-
A0A6I9HHF5 : ---------MPV-RK--HS-Q--HMKL-F-K----NKGKDETTLR-RQRAEVSVELRKAKKDEQL-LKRRN-LS-VHE-R-EE-
H0ZB96     : ---------MPV-RK--HD-G--HMKL-F-K----NKGKDQTTLR-RQRVEVSVELRKAKKDEQI-LKRRN-IS-INL-E-KE-
A0A6J2IHK9 : ---------MEV-GK--QE-Q--RMRK-F-K----NKGKDAAALR-MQRVEMSIELRKAKKDEQI-LKRRN-VS-LHV-K-DE-
A0A672U8V3 : -----PAKMSLLIK--HN-E--RLRK-F-K----NKGKDPTALR-RQRVEVIVELRKAKKDEQI-LKRRN-IE-QNK-M-EM-
C0LLJ0     : ---------MATSKA--PK-E--RLKN-Y-K----YRGKEMSLPR-QQRIASSLQLRKTRKDEQV-LKRRN-ID-LFS-S-DM-
A0A6P7RAC8 : ---------MATSKA--PK-E--RLKN-Y-K----YXGKEMSLPR-QQRIASSLQLRKTRKDEQV-LKRRN-ID-LFS-S-DM-
D3Z2P7     : ---------MATSKA--PK-E--RLKN-Y-K----YRGKEMSLPR-QQRIASSLQLRKTRKDEQVNFTLDDIIQ-AVN-S-S--
A0A096MK49 : ---------MATSEA--PE-E--RLKK-F-K----YRGKEMSLRR-QQRIDSSLQLRKSRKDEQA-LKRRN-IG-LFS-S-DV-
A0A2Y9G9Q9 : ---------MPTLDA--PE-E--RLRK-F-K----YRGKDASMRR-QQRMAVSLELRKAKKDEQA-LKRRN-IT-NFS-P-DP-
A0A2U3VHW3 : ---------MPTLDA--PE-E--RLRK-F-K----YRGKDASMRR-QQRMAVSLELRKAKKDEQA-LKRRN-IT-NFS-P-DP-
A0A2U3YYF1 : ---------MPTLDA--PE-E--RLRK-F-K----YRGKDASMRR-QQRMAVSLELRKAKKDEQA-LKRRN-IT-NFS-P-DP-
A0A3Q7XXW8 : ---------MPTLDA--PE-E--RMRK-F-K----YRGKDASMRR-QQRMAVSLELRKAKKDEQA-LKRRN-IT-NFS-P-DP-
G1LJH4     : ---------MPTLDA--PE-E--RMRK-F-K----YRGKDASMRR-QQRMAVSLELRKAKKDEQA-LKRRN-IT-NFS-P-DP-
A0A452QVF9 : ---------MPTLDA--PE-E--RMRK-F-K----YRGKDASMRR-QQRMAVSLELRKAKKDEQA-LKRRN-IT-NFS-P-DP-
A0A384CQH9 : ---------MPTLDA--PE-E--RMRK-F-K----YRGKDASMRR-QQRMAVSLELRKAKKDEQA-LKRRN-IT-NFS-P-DP-
A0A3Q7NGF8 : ---------MPTLDA--PE-E--RLRK-F-K----YRGKDASMRR-QQRMAVSLELRKAKKEEQA-LKRRN-IT-NFS-P-DP-
G1RIW6     : ---------MPTLDA--LE-E--RRRK-F-K----YRGKDASLRR-QQRMAVSLELRKAKKDEQT-LKRRN-IT-SFC-P-DT-
H2PLH7     : ---------MPTLDA--PE-E--RRRK-F-K----YRGKDASLRR-QQRMAVSLELRKAKKDEQT-LKRRN-IT-SFC-P-DT-
A0A2R9C214 : ---------MPTLDA--PE-E--RRRK-F-K----YRGKDASLRR-QQRMAVSLELRKAKKDEQT-LKRRN-IM-SFC-P-DT-
A9QM74     : ---------MPTLDA--PE-E--RRRK-F-K----YRGKDVSLRR-QQRMAVSLELRKAKKDEQT-LKRRN-IT-SFC-P-DT-
A0A2K6A4U0 : ---------MPTLDA--PE-D--RRRK-F-K----YCGKDASLRR-QQRMAVSLELRKAKKDEQT-LKRRN-IT-SFC-L-DT-
A0A2K5UXA0 : ---------MPTLDA--PE-D--RRRK-F-K----YCGKDASLRR-QQRMAVSLELRKAKKDEQT-LKRRN-IT-SFC-L-DT-
F6R0P5     : ---------MPTLDA--PE-D--RRRK-F-K----YCGKDASLRR-QQRMAVSLELRKAKKDEQT-LKRRN-IT-SFC-L-DT-
A0A096P098 : ---------MPTLDA--PE-D--RRRK-F-K----YCGKDASLRR-QQRMAVSLELRKAKKDEQT-LKRRN-IT-SFC-L-DT-
A0A2K5KVE0 : ---------MPTLDA--PE-D--RRRK-F-K----YCGKDASLRR-QQRMAVSLELRKAKKDEQT-LKRRN-IT-SFC-L-DT-
A0A2K6DYG1 : ---------MPTLDA--PE-D--RRRK-F-K----YCGKDASLRR-QQRMAVSLELRKAKKDEQT-LKRRN-IT-SFC-L-DT-
A0A2K6QJE3 : ---------MPTLDA--PE-D--RRRK-F-K----YCGKDASLRR-QQRMAVSLELRKAKKDEQT-LKRRN-IT-SFC-P-DT-
A0A2K6L189 : ---------MPTLDA--PE-D--RRRK-F-K----YCGKDASLRR-QQRMAVSLELRKAKKDEQT-LKRRN-IT-SFC-P-DT-
A0A6J3GXN8 : -NLLLPTNMPSLDA--PE-E--RLRK-F-K----YRGKDASLRR-QQRMAVSLKLRKAKKDEQT-LKRRN-IT-NFF-P-DT-
M3YYN0     : ---------MPTLDA--PE-G--RLRK-F-K----YRGKDASIRR-HQRMAVSLELRKAKKDEQA-LKRRN-IT-IFS-P-EP-
A0A6P6HTK7 : ---------MPTLDA--PE-E--RLRK-F-K----YRGKDASMRR-QQRMAVSLELRKAKKDEQA-LKRRN-IT-NVS-S-DP-
A0A667HG78 : ---------MPTLDA--PE-E--RLRK-F-K----YRGKDASMRR-QQRMAVSLELRKAKKDEQA-LKRRN-IT-NVS-S-DP-
A0A6J1Y007 : ---------MPTLDA--PE-E--RLRK-F-K----YRGKDASMRR-QQRMAVSLELRKAKKDEQA-LKRRN-IT-NVS-S-DP-
M3WLV5     : ---------MPTLDA--PE-E--RLRK-F-K----YRGKDASMRR-QQRMAVSLELRKAKKDEQA-LKRRN-IT-NVS-S-DP-
A0A6P4TYG7 : ---------MPTLDA--PE-E--RLRK-F-K----YRGKDASMRR-QQRMAVSLELRKAKKDEQA-LKRRN-IT-NVS-S-DP-
A0A673UH42 : ---------MPTLDA--PE-E--RLRK-F-K----YRGKDTSMRR-QQRMAVSLELRKAKKEEQA-LKRRN-IT-NLS-S-DP-
A0A3Q7RIG0 : ---------MPTLHA--PE-E--RLRK-F-K----YRGKDVSMRR-QQRIAVSLELRKAKKDEQA-LKRRN-IA-NFS-T-DP-
```

```
                       *         20         *         40         *         60         *         80
A0A287DCQ1 : --------MSTLDV--PE-E--RLRK-F-K----YRGKDASMRR-QQRIIVSVELRKARKDEQV-LKRRN-IN-HVS-A-DP-
H0XT12     : ----LVS-MSNSDA--PE-E--RLRK-F-K----YRGKDASMRR-QQRIVVSQKLRKVKKDEQT-FKRRN-IV-SFS-H-DP-
A0A2K6GMC3 : --------MPTLDV--PE-E--RLRK-F-K----YRGKDASMRR-QQRMVVSLELRKARKDEQT-LKRRN-II-DSC-L-DL-
A0A6P5E101 : --------MPTLDA--PE-E--RLRK-F-K----YRGKDASARR-QQRIAVSLELRKAKKDEQA-LKRRN-IT-DVS-L-DP-
A0A6P3HNK5 : --------MPTLDA--PE-E--RLRK-F-K----YRGKDASARR-QQRIAVSLELRKAKKDEQA-LKRRN-IT-DVS-L-DP-
A0A4W2BKA2 : --------MPTLDA--PE-E--RLRK-F-K----YRGKDASARR-QQRIAVSLELRKAKKDEQA-LKRRN-IT-DVS-L-DP-
C1JZ66     : --------MPTLDA--PE-E--RLRK-F-K----YRGKDASARR-QQRIAVSLELRKAKKDEQA-LKRRN-IT-DVS-L-DP-
A0A6P3YTR5 : --------MPTSDA--PE-E--RLRK-F-K----YRGKDASARR-QQRIAVSLELRKAKKDEQA-LKRRN-IT-DVS-L-DP-
A0A452G6T6 : --------MPTSDA--PE-E--RLRK-F-K----YRGKDASARR-QQRIAVSLELRKAKKDEQA-LKRRN-IT-DVS-L-DP-
A0A6J0Y4Q7 : -TLLLPVNMPTLDA--PE-E--RLRK-F-K----YRGKDASARR-QQRFAVSLELRKARKDEQA-LKRRN-IT-SVS-H-DP-
A0A6J3AJ62 : --------MPTLEA--PE-E--RLRK-F-K----YRGKDASVRR-QQRIAVSLELRKAKKDEQA-LKRRN-IT-SVS-P-DP-
A0A6P3QNL6 : --------MPTLEA--PE-E--RLRK-F-K----YRGKDASARR-QQRIVVSLQLRKAKKDEQA-LKRRN-IT-SLS-P-DP-
A0A455C3E3 : -ALLLLVNMPTLEA--PG-E--RLRK-F-K----YRGKDASMRR-QQRIAVSLELRKAKKDEQS-LKRRN-IT-DVS-P-DP-
A0A340WH20 : -ALLLLVNMPTLEA--PE-E--RLRK-F-K----YRGKDASMRR-QQRIAVSLELRKAKKDEQS-LKRRN-IT-DVS-P-DP-
A0A384AHF6 : -ALLLLVNMPTLEA--PE-E--RLRK-F-K----YRGKDASMRR-QQRIAVSLELRKAKKDEQS-LKRRN-IT-DVS-P-DP-
A0A341BFZ0 : --------MPTSEA--PE-E--RLRK-F-K----YRGKDASMRR-QQRIAVSLELRKAKKDEQS-LKRRN-IT-DVS-P-DA-
A0A2Y9PGE5 : -ALLLLVNMPTLEA--PE-E--RLRK-F-K----YRGKDASMRR-QQRIAVSLELRKAKKDEQS-LKRRN-IT-DVS-P-DA-
G1SG50     : --------MPTLDV--PE-E--RLRK-F-K----YRGKEASVRR-QQRVAVSLELRKARKEEQA-LKRRN-IT-DCS-P-EP-
G3SVL2     : --------MATLEA--PE-E--RLRK-F-K----YRGKDASVRR-QQRIAVSLELRKAKKEEQA-LKRRN-IT-DLS-P-DP-
H9KVH0     : --------MPILEA--PE-E--RLRK-F-K----YRGKDASVRR-QQRLAVSLELRKAKKDEQA-LKRRN-IT-TAS---DPF
A0A6P3VA68 : RQFSS-VNMAAVGY--PK-W--RLNQ-F-K----YRGKNISVRR-RQRIAFSLELRKAKKDEQA-FKRRN-IT-YLS-P-YP-
F7CWU3     : -------NMLT-GA--PE-G--RLRK-F-K----YRGKD-TVRR-QQRIAVSLELRKAKKDEQA-LKRRN-IT-PLF-P-DP-
H0VQS0     : -DFLPLVNMPAMDT--TE-E--RLKK-F-K----YRGKDASVRR-QQRVALSLELRKAKKDEQA-LKRRN-IT-CPS-P-DP-
A0A1S3AKG9 : --------MPTPEA--PE-E--RLKK-F-K----YRGKDASALR-QQRIAVILELRKAKKDEQS-LKRRN-IT-N-G-PLNS-
A0A6I9JDZ1 : --------MPISEA--PE-G--RLKQ-F-K----YRGRDASVRR-QQRVTVSLVLRKAKKHEQV-LKRRN-IT-SCS-S-SP-
G1NZS1     : ----LPVNMSC-EA--PE-D--RLRK-F-K----YRGKDASMIR-QHRIAVNVQLRKAKKDEQV-LKRRN-IT-NLC-S-DS-
A0A3Q0D6R9 : --------MPALEA--PE-D--RLKK-F-K----YRGKDVSARR-QQRIAANLQLRKAKKDEQA-LKRRN-IS-QCL-S-YL-
A0A6I9LF97 : --------MPALEA--PE-E--RLKK-F-K----YRGKDVSARR-QERIATNLQLRKAKKDEQA-LKRRN-IS-QSS-N-DL-
A0A5F8GUF5 : ----MSA--SG--I--QN-E--RLKK-F-K----YRGRNMGLIR-QQRIAVSVELRKAKKEEQI-LKKRN-I---FN-TLHSL
F6TJ07     : ICPSLAANMSSCEV--Q--E--RMKM-F-K----NKGKNTELRR-QQRLEVNVQLRKAKKEEQI-LKKRN-IS-DF--PLME-
Q803D8     : --------MPTENL--SD-K--RMSK-F-K----NKGKEPTKLR-DRRVAECVEFRKAQRVEGI-MKRRN-IT-SLG-D-EE-
A0A060YJS5 : --------MPTTSA--TD-D--RLNK-F-K----NKGKEPSKLR-ERRIAECVELRKAAKNENF-LKRRN-IT-TLP-E-DA-
A0A6P8VU10 : --------MPTKNV--DD-A--RISK-F-K----NKGKDPAKLR-EKRISACVELRKANKDENF-LKRRN-ISVDLS-S-D--
cIBB7      : --------MPTLDA--PE-E--RLRK-F-K----YRGKDASMRR-QQRIAVSLELRKAKKDEQA-LKRRN-IT-SFS-P-DP-
```
