## Supporting file 3 for "Biochemical propensity mapping for structural and functional anatomy of importin α IBB domain"

```
                       *        20         *        40         *        60         *        80         *       100
cKPNA7_22 : ---MPTLD-APEERLRKFKYRGKD-AS-MRRQQRIAVSLELRKAKKDEQALKRRNI--------TSFSPD----PASEQ--TKG-----VSLTLQEIING
cKPNA7_24 : ---MPTLD-APEERLRKFKYRGKD-AS-MRRQQRIAVSLELRKAKKDEQALKRRNI--------TSFSPD----PASEQ--TKG-----VSLTLQEIING
cKPNA7_1  : ---MPTLD-APEERLRKFKYRGKD-AS-MRRQQRIAVSLELRKAKKDEQALKRRNI--------TSFSPD----PASEQ--TKG-----VSLTLQEIING
cKPNA7_3  : ---MPTLD-APEERLRKFKYRGKD-AS-MRRQQRIAVSLELRKAKKDEQALKRRNI--------TSFSPD----PASEQ--TKG-----VSLTLQEIING
cKPNA7_9  : ---MPTLD-APEERLRKFKYRGKD-AS-MRRQQRIAVSLELRKAKKDEQALKRRNI--------TSFSPD----PASEQ--TKG-----VSLTLQEIING
cKPNA7_11 : ---MPTLD-APEERLRKFKYRGKD-AS-MRRQQRIAVSLELRKAKKDEQALKRRNI--------TSFSPD----PASEQ--TKG-----VSLTLQEIING
cKPNA7_17 : ---MPTLD-APEERLRKFKYRGKD-AS-MRRQQRIAVSLELRKAKKDEQALKRRNI--------TSFSPD----PASEQ--TKG-----VSLTLQEIING
cKPNA7_19 : ---MPTLD-APEERLRKFKYRGKD-AS-MRRQQRIAVSLELRKAKKDEQALKRRNI--------TSFSPD----PASEQ--TKG-----VSLTLQEIING
cKPNA7_5  : ---MPTLD-APEERLRKFKYRGKD-AS-MRRQQRIAVSLELRKAKKDEQALKRRNI--------TSFSPD----PASEQ--TKG-----VSLTLQEIING
cKPNA7_7  : ---MPTLD-APEERLRKFKYRGKD-AS-MRRQQRIAVSLELRKAKKDEQALKRRNI--------TSFSPD----PASEQ--TKG-----VSLTLQEIING
cKPNA7_13 : ---MPTLD-APEERLRKFKYRGKD-AS-MRRQQRIAVSLELRKAKKDEQALKRRNI--------TSFSPD----PASEQ--TKG-----VSLTLQEIING
cKPNA7_15 : ---MPTLD-APEERLRKFKYRGKD-AS-MRRQQRIAVSLELRKAKKDEQALKRRNI--------TSFSPD----PASEQ--TKG-----VSLTLQEIING
cKPNA7_21 : ---MPTLD-APEERLRKFKYRGKD-AS-MRRQQRIAVSLELRKAKKDEQALKRRNI--------TSFSPD----PASEQ--TKG-----VSLTLQEIING
cKPNA7_23 : ---MPTLD-APEERLRKFKYRGKD-AS-MRRQQRIAVSLELRKAKKDEQALKRRNI--------TSFSPD----PASEQ--TKG-----VSLTLQEIING
cKPNA7_2  : ---MPTLD-APEERLRKFKYRGKD-AS-MRRQQRIAVSLELRKAKKDEQALKRRNI--------TSFSPD----PASEQ--TKG-----VSLTLQEIING
cKPNA7_4  : ---MPTLD-APEERLRKFKYRGKD-AS-MRRQQRIAVSLELRKAKKDEQALKRRNI--------TSFSPD----PASEQ--TKG-----VSLTLQEIING
cKPNA7_10 : ---MPTLD-APEERLRKFKYRGKD-AS-MRRQQRIAVSLELRKAKKDEQALKRRNI--------TSFSPD----PASEQ--TKG-----VSLTLQEIING
cKPNA7_12 : ---MPTLD-APEERLRKFKYRGKD-AS-MRRQQRIAVSLELRKAKKDEQALKRRNI--------TSFSPD----PASEQ--TKG-----VSLTLQEIING
cKPNA7_18 : ---MPTLD-APEERLRKFKYRGKD-AS-MRRQQRIAVSLELRKAKKDEQALKRRNI--------TSFSPD----PASEQ--TKG-----VSLTLQEIING
cKPNA7_20 : ---MPTLD-APEERLRKFKYRGKD-AS-MRRQQRIAVSLELRKAKKDEQALKRRNI--------TSFSPD----PASEQ--TKG-----VSLTLQEIING
cKPNA7_6  : ---MPTLD-APEERLRKFKYRGKD-AS-MRRQQRIAVSLELRKAKKDEQALKRRNI--------TSFSPD----PASEQ--TKG-----VSLTLQEIING
cKPNA7_8  : ---MPTLD-APEERLRKFKYRGKD-AS-MRRQQRIAVSLELRKAKKDEQALKRRNI--------TSFSPD----PASEQ--TKG-----VSLTLQEIING
cKPNA7_14 : ---MPTLD-APEERLRKFKYRGKD-AS-MRRQQRIAVSLELRKAKKDEQALKRRNI--------TSFSPD----PASEQ--TKG-----VSLTLQEIING
cKPNA7_16 : ---MPTLD-APEERLRKFKYRGKD-AS-MRRQQRIAVSLELRKAKKDEQALKRRNI--------TSFSPD----PASEQ--TKG-----VSLTLQEIING
cKPNA2_1  : ---MSTNENNSAARLNRFKNKGKD-STEMRRR-RIEVNVELRKAKKDDQMLKRRNV--------SSF-PDDATSPLQENRNNQGT----VHWSVEEIVKG
cKPNA2_2  : ---MSTNENNSAARLNRFKNKGKD-STEMRRR-RIEVNVELRKAKKDDQMLKRRNV--------SSF-PDDATSPLQENRNNQGT----VHWSVEEIVKG
cKPNA3    : ---MAENAGLENHRIKSFKNKGRDVET-MRRH-RNEVTVELRKNKRDEHLLKKRNVPQ------EESLED---SDVDAD--FKA-----QNVTLEAILQN
cKPNA4    : ---MADNEKLDNQRLKNFKNKGRDLET-MRRQ-RNEVVVELRKNKRDEHLLKRRNVPH------EDICED---SDIDGD--YRV-----QNTSLEAIVQN
cKPNA5    : -DAMASPG-KDNYRMKSYKNKALNPQE-MRRR-REEEGIQLRKQKREEQLFKRRNVSLP--RNDES-MLE---SPIQDPDISSTVPIPEEEVITTDMVQM
cKPNA6    : METMASPG-KDNYRMKSYKNNALNPEE-MRRR-REEEGIQLRKQKREQQLFKRRNVEL---INEEAAMFD---SLLMDSYVSSTT---GESVITREMVEM
cKPNA1    : ---MTTPG-KENFRLKSYKNKSLNPDE-MRRR-REEEGLQLRKQKREEQLFKRRNVATAEEETEEEVMSD---GGFHEAQINNMEMAPG-GVITSDMIEM
                       *       120         *       140         *       160         *       180         *       200
cKPNA7_22 : VNSSDPELCFQATQAARKMLSREKNPPLKLIIEA-GLIPRLVEFL-KSSLHPCLQFEAAWALTNIASGTSEQTRAVVEGGAIQPLVELLSSPHMTVCEQA
cKPNA7_24 : VNSSDPELCFQATQAARKMLSREKNPPLKLIIEA-GLIPRLVEFL-KSSLHPCLQFEAAWALTNIASGTSEQTRAVVEGGAIQPLVELLSSPHMTVCEQA
cKPNA7_1  : VNSSDPELCFQATQAARKMLSREKNPPLKLIIEA-GLIPRLVEFL-KSSLHPCLQFEAAWALTNIASGTSEQTRAVVEGGAIQPLVELLSSPHMTVCEQA
cKPNA7_3  : VNSSDPELCFQATQAARKMLSREKNPPLKLIIEA-GLIPRLVEFL-KSSLHPCLQFEAAWALTNIASGTSEQTRAVVEGGAIQPLVELLSSPHMTVCEQA
cKPNA7_9  : VNSSDPELCFQATQAARKMLSREKNPPLKLIIEA-GLIPRLVEFL-KSSLHPCLQFEAAWALTNIASGTSEQTRAVVEGGAIQPLVELLSSPHMTVCEQA
cKPNA7_11 : VNSSDPELCFQATQAARKMLSREKNPPLKLIIEA-GLIPRLVEFL-KSSLHPCLQFEAAWALTNIASGTSEQTRAVVEGGAIQPLVELLSSPHMTVCEQA
cKPNA7_17 : VNSSDPELCFQATQAARKMLSREKNPPLKLIIEA-GLIPRLVEFL-KSSLHPCLQFEAAWALTNIASGTSEQTRAVVEGGAIQPLVELLSSPHMTVCEQA
cKPNA7_19 : VNSSDPELCFQATQAARKMLSREKNPPLKLIIEA-GLIPRLVEFL-KSSLHPCLQFEAAWALTNIASGTSEQTRAVVEGGAIQPLVELLSSPHMTVCEQA
cKPNA7_5  : VNSSDPELCFQATQAARKMLSREKNPPLKLIIEA-GLIPRLVEFL-KSSLHPCLQFEAAWALTNIASGTSEQTRAVVEGGAIQPLVELLSSPHMTVCEQA
cKPNA7_7  : VNSSDPELCFQATQAARKMLSREKNPPLKLIIEA-GLIPRLVEFL-KSSLHPCLQFEAAWALTNIASGTSEQTRAVVEGGAIQPLVELLSSPHMTVCEQA
cKPNA7_13 : VNSSDPELCFQATQAARKMLSREKNPPLKLIIEA-GLIPRLVEFL-KSSLHPCLQFEAAWALTNIASGTSEQTRAVVEGGAIQPLVELLSSPHMTVCEQA
cKPNA7_15 : VNSSDPELCFQATQAARKMLSREKNPPLKLIIEA-GLIPRLVEFL-KSSLHPCLQFEAAWALTNIASGTSEQTRAVVEGGAIQPLVELLSSPHMTVCEQA
cKPNA7_21 : VNSSDPELCFQATQAARKMLSREKNPPLKLIIEA-GLIPRLVEFL-KSSLHPCLQFEAAWALTNIASGTSEQTRAVVEGGAIQPLVELLSSPHMTVCEQA
cKPNA7_23 : VNSSDPELCFQATQAARKMLSREKNPPLKLIIEA-GLIPRLVEFL-KSSLHPCLQFEAAWALTNIASGTSEQTRAVVEGGAIQPLVELLSSPHMTVCEQA
cKPNA7_2  : VNSSDPELCFQATQAARKMLSREKNPPLKLIIEA-GLIPRLVEFL-KSSLHPCLQFEAAWALTNIASGTSEQTRAVVEGGAIQPLVELLSSPHMTVCEQA
cKPNA7_4  : VNSSDPELCFQATQAARKMLSREKNPPLKLIIEA-GLIPRLVEFL-KSSLHPCLQFEAAWALTNIASGTSEQTRAVVEGGAIQPLVELLSSPHMTVCEQA
cKPNA7_10 : VNSSDPELCFQATQAARKMLSREKNPPLKLIIEA-GLIPRLVEFL-KSSLHPCLQFEAAWALTNIASGTSEQTRAVVEGGAIQPLVELLSSPHMTVCEQA
cKPNA7_12 : VNSSDPELCFQATQAARKMLSREKNPPLKLIIEA-GLIPRLVEFL-KSSLHPCLQFEAAWALTNIASGTSEQTRAVVEGGAIQPLVELLSSPHMTVCEQA
cKPNA7_18 : VNSSDPELCFQATQAARKMLSREKNPPLKLIIEA-GLIPRLVEFL-KSSLHPCLQFEAAWALTNIASGTSEQTRAVVEGGAIQPLVELLSSPHMTVCEQA
cKPNA7_20 : VNSSDPELCFQATQAARKMLSREKNPPLKLIIEA-GLIPRLVEFL-KSSLHPCLQFEAAWALTNIASGTSEQTRAVVEGGAIQPLVELLSSPHMTVCEQA
cKPNA7_6  : VNSSDPELCFQATQAARKMLSREKNPPLKLIIEA-GLIPRLVEFL-KSSLHPCLQFEAAWALTNIASGTSEQTRAVVEGGAIQPLVELLSSPHMTVCEQA
cKPNA7_8  : VNSSDPELCFQATQAARKMLSREKNPPLKLIIEA-GLIPRLVEFL-KSSLHPCLQFEAAWALTNIASGTSEQTRAVVEGGAIQPLVELLSSPHMTVCEQA
cKPNA7_14 : VNSSDPELCFQATQAARKMLSREKNPPLKLIIEA-GLIPRLVEFL-KSSLHPCLQFEAAWALTNIASGTSEQTRAVVEGGAIQPLVELLSSPHMTVCEQA
cKPNA7_16 : VNSSDPELCFQATQAARKMLSREKNPPLKLIIEA-GLIPRLVEFL-KSSLHPCLQFEAAWALTNIASGTSEQTRAVVEGGAIQPLVELLSSPHMTVCEQA
cKPNA2_1  : VNSNNLESQLQATQAARKLLSREKQPPIDNIIRA-GLIPKFVSFLGRTDCPP-IQFEAAWALTNIASGTSEQTKAVVDGGAIPAFISLLASPHAHISEQA
cKPNA2_2  : VNSNNLESQLQATQAARKLLSREKQPPIDNIIRA-GLIPKFVSFLGRTDCPP-IQFEAAWALTNIASGTSEQTKAVVDGGAIPAFISLLASPHAHISEQA
cKPNA3    : ATSDNPVVQLSAVQAARKLLSSDRNPPIDDLIKS-GILPILVKCLERDD-NPSLQFEAAWALTNIASGTSAQTQAVVQSNAVPLFLRLLHSPHQNVCEQA
cKPNA4    : ASSDNQGIQLSAVQAARKLLSSDRNPPIDDLIKS-GILPILVHCLERDD-NPSLQFEAAWALTNIASGTSEQTQAVVQSNAVPLFLRLLHSPHQNVCEQA
cKPNA5    : IFSNNADQQLTATQKFRKLLSKEPNPPIDQVIQKPGVVQRFVKFLERNE-NCTLQFEAAWALTNIASGTFLHTKVVIETGAVPIFIKLLNSEHEDVQEQA
cKPNA6    : LFSDDSDLQLATTQKFRKLLSKEPSPPIDEVINTPGVVDRFVEFLKRNE-NCTLQFEAAWALTNIASGTSQQTKIVIEAGAVPIFIELLNSDFEDVQEQA
cKPNA1    : IFSNSPEQQLSATQKFRKLLSKEPNPPIDEVISTPGVVARFVEFLKRKE-NCTLQFEAAWVLTNIASGNSLQTRIVIQAGAVPIFIELLSSEFEDVQEQA
```

```
                      *        220          *        240          *        260          *        280          *        300
cKPNA7_22 : VWALGNIAGDGPEFRDAVISSNAI-PHLLALVS-PTIP-IT--FLRNITWTLSNLCRNKNPYPCDKAVKQMLPVLSHLLQHQDKEILSDTCWALSYLTDG
cKPNA7_24 : VWALGNIAGDGPEFRDAVISSNAI-PHLLALVS-PTIP-IT--FLRNITWTLSNLCRNKNPYPCDKAVKQMLPVLSHLLQHQDKEILSDTCWALSYLTDG
cKPNA7_1  : VWALGNIAGDGPEFRDAVISSNAI-PHLLALVS-PTIP-IT--FLRNITWTLSNLCRNKNPYPCDKAVKQMLPVLLHLLQHQDKEILSDTCWALSYLTDG
cKPNA7_3  : VWALGNIAGDGPEFRDAVISSNAI-PHLLALVS-PTIP-IT--FLRNITWTLSNLCRNKNPYPCDKAVKQMLPVLLHLLQHQDKEILSDTCWALSYLTDG
cKPNA7_9  : VWALGNIAGDGPEFRDAVISSNAI-PHLLALVS-PTIP-IT--FLRNITWTLSNLCRNKNPYPCDKAVKQMLPVLFHLLQHQDKEILSDTCWALSYLTDG
cKPNA7_11 : VWALGNIAGDGPEFRDAVISSNAI-PHLLALVS-PTIP-IT--FLRNITWTLSNLCRNKNPYPCDKAVKQMLPVLFHLLQHQDKEILSDTCWALSYLTDG
cKPNA7_17 : VWALGNIAGDGPEFRDAVISSNAI-PHLLALVS-PTIP-IT--FLRNITWTLSNLCRNKNPYPCDKAVKQMLPVLSHLLQHQDKEILSDTCWALSYLTDG
cKPNA7_19 : VWALGNIAGDGPEFRDAVISSNAI-PHLLALVS-PTIP-IT--FLRNITWTLSNLCRNKNPYPCDKAVKQMLPVLSHLLQHQDKEILSDTCWALSYLTDG
cKPNA7_5  : VWALGNIAGDGPEFRDAVISSNAI-PHLLALVS-PTIP-IT--FLRNITWTLSNLCRNKNPYPCDKAVKQMLPVLLHLLQHQDKEILSDTCWALSYLTDG
cKPNA7_7  : VWALGNIAGDGPEFRDAVISSNAI-PHLLALVS-PTIP-IT--FLRNITWTLSNLCRNKNPYPCDKAVKQMLPVLLHLLQHQDKEILSDTCWALSYLTDG
cKPNA7_13 : VWALGNIAGDGPEFRDAVISSNAI-PHLLALVS-PTIP-IT--FLRNITWTLSNLCRNKNPYPCDKAVKQMLPVLFHLLQHQDKEILSDTCWALSYLTDG
cKPNA7_15 : VWALGNIAGDGPEFRDAVISSNAI-PHLLALVS-PTIP-IT--FLRNITWTLSNLCRNKNPYPCDKAVKQMLPVLFHLLQHQDKEILSDTCWALSYLTDG
cKPNA7_21 : VWALGNIAGDGPEFRDAVISSNAI-PHLLALVS-PTIP-IT--FLRNITWTLSNLCRNKNPYPCDKAVKQMLPVLSHLLQHQDKEILSDTCWALSYLTDG
cKPNA7_23 : VWALGNIAGDGPEFRDAVISSNAI-PHLLALVS-PTIP-IT--FLRNITWTLSNLCRNKNPYPCDKAVKQMLPVLSHLLQHQDKEILSDTCWALSYLTDG
cKPNA7_2  : VWALGNIAGDGPEFRDAVISSNAI-PHLLALVS-PTIP-IT--FLRNITWTLSNLCRNKNPYPCDKAVKQMLPVLLHLLQHQDKEILSDTCWALSYLTDG
cKPNA7_4  : VWALGNIAGDGPEFRDAVISSNAI-PHLLALVS-PTIP-IT--FLRNITWTLSNLCRNKNPYPCDKAVKQMLPVLLHLLQHQDKEILSDTCWALSYLTDG
cKPNA7_10 : VWALGNIAGDGPEFRDAVISSNAI-PHLLALVS-PTIP-IT--FLRNITWTLSNLCRNKNPYPCDKAVKQMLPVLFHLLQHQDKEILSDTCWALSYLTDG
cKPNA7_12 : VWALGNIAGDGPEFRDAVISSNAI-PHLLALVS-PTIP-IT--FLRNITWTLSNLCRNKNPYPCDKAVKQMLPVLFHLLQHQDKEILSDTCWALSYLTDG
cKPNA7_18 : VWALGNIAGDGPEFRDAVISSNAI-PHLLALVS-PTIP-IT--FLRNITWTLSNLCRNKNPYPCDKAVKQMLPVLSHLLQHQDKEILSDTCWALSYLTDG
cKPNA7_20 : VWALGNIAGDGPEFRDAVISSNAI-PHLLALVS-PTIP-IT--FLRNITWTLSNLCRNKNPYPCDKAVKQMLPVLSHLLQHQDKEILSDTCWALSYLTDG
cKPNA7_6  : VWALGNIAGDGPEFRDAVISSNAI-PHLLALVS-PTIP-IT--FLRNITWTLSNLCRNKNPYPCDKAVKQMLPVLLHLLQHQDKEILSDTCWALSYLTDG
cKPNA7_8  : VWALGNIAGDGPEFRDAVISSNAI-PHLLALVS-PTIP-IT--FLRNITWTLSNLCRNKNPYPCDKAVKQMLPVLLHLLQHQDKEILSDTCWALSYLTDG
cKPNA7_14 : VWALGNIAGDGPEFRDAVISSNAI-PHLLALVS-PTIP-IT--FLRNITWTLSNLCRNKNPYPCDKAVKQMLPVLFHLLQHQDKEILSDTCWALSYLTDG
cKPNA7_16 : VWALGNIAGDGPEFRDAVISSNAI-PHLLALVS-PTIP-IT--FLRNITWTLSNLCRNKNPYPCDKAVKQMLPVLFHLLQHQDKEILSDTCWALSYLTDG
cKPNA2_1  : VWALGNIAGDGSVYRDLVIKHGAVDP-LLALLAVPDLSSLACGYLRNVTWTLSNLCRNKNPAPPLEAVEQILPTLVRLLHHDDPEVLADTCWAISYLTDG
cKPNA2_2  : VWALGNIAGDGSVYRDLVIKHGAVDP-LLALLAVPDLSSLACGYLRNVTWTLSNLCRNKNPAPPLEAVEQILPTLVRLLHHDDPEVLADTCWAISYLTDG
cKPNA3    : VWALGNIIGDGPQCRDYVISLGVVKP-LLSFIN-PSIP-IT--FLRNVTWVIVNLCRNKDPPPPMETVQEILPALCVLIYHTDINILVDTVWALSYLTDG
cKPNA4    : VWALGNIIGDGPQCRDYVISLGVVKP-LLSFIS-PSIP-IT--FLRNVTWVMVNLCRHKDPPPPMETIQEILPALCVLIHHTDVNILVDTVWALSYLTDA
cKPNA5    : VWALGNIAGDNAECRDFVLNCEILPP-LLELLT-NSNR-LT--TTRNAVWALSNLCRGKNPPPNFSKVSPCLNVLSRLLFSSDPDVLADVCWALSYLSDG
cKPNA6    : VWALGNIAGDSSVCRDYVLNCSILNP-LLTLLT-KSTR-LT--MTRNAVWALSNLCRGKNPPPEFAKVSPCLPVLSRLLFSSDSDLLADACWALSYLSDG
cKPNA1    : VWALGNIAGDSTMCRDYVLDCNILPP-LLQLLS-KQNR-LT--MTRNAVWALSNLCRGKNPPPEFAKVSPCLNVLSWLLFVNDTDVLADACWALSYLSDG
                      *        320          *        340          *        360          *        380          *        400
cKPNA7_22 : SNERIGQVVDTGVLPRLVELMTSSELNVLTPSLRTVGNIVTGTDEQTQVAIDAGVLNVLPQLLRHPKPSIQKEAAWALSNVAAGPCQHIQQLIACGLLPP
cKPNA7_24 : SNERIGQVVDTGVLPRLVELMTSSELNVLTPSLRTVGNIVTGTDEQTQVAIDAGVLNVLPQLLRHPKPSIQKEAAWALSNVAAGPCQHIQQLIACGLLPP
cKPNA7_1  : SNDRIGQVVDTGVLPRLVELMTSSELNVLTPSLRTVGNIVTGTDEQTQVAIDAGVLNVLPQLLRHPKPSIQKEAAWALSNVAAGPCQHIQQLIACGLLPP
cKPNA7_3  : SNDRIGQVVDTGVLPRLVELMTSSELNVLTPSLRTVGNIVTGTDEQTQVAIDAGVLNVLPQLLRHPKPSIQKEAAWALSNVAAGPCQHIQQLIACGLLPP
cKPNA7_9  : SNDRIGQVVDTGVLPRLVELMTSSELNVLTPSLRTVGNIVTGTDEQTQVAIDAGVLNVLPQLLRHPKPSIQKEAAWALSNVAAGPCQHIQQLIACGLLPP
cKPNA7_11 : SNDRIGQVVDTGVLPRLVELMTSSELNVLTPSLRTVGNIVTGTDEQTQVAIDAGVLNVLPQLLRHPKPSIQKEAAWALSNVAAGPCQHIQQLIACGLLPP
cKPNA7_17 : SNDRIGQVVDTGVLPRLVELMTSSELNVLTPSLRTVGNIVTGTDEQTQVAIDAGVLNVLPQLLRHPKPSIQKEAAWALSNVAAGPCQHIQQLIACGLLPP
cKPNA7_19 : SNDRIGQVVDTGVLPRLVELMTSSELNVLTPSLRTVGNIVTGTDEQTQVAIDAGVLNVLPQLLRHPKPSIQKEAAWALSNVAAGPCQHIQQLIACGLLPP
cKPNA7_5  : SNERIGQVVDTGVLPRLVELMTSSELNVLTPSLRTVGNIVTGTDEQTQVAIDAGVLNVLPQLLRHPKPSIQKEAAWALSNVAAGPCQHIQQLIACGLLPP
cKPNA7_7  : SNERIGQVVDTGVLPRLVELMTSSELNVLTPSLRTVGNIVTGTDEQTQVAIDAGVLNVLPQLLRHPKPSIQKEAAWALSNVAAGPCQHIQQLIACGLLPP
cKPNA7_13 : SNERIGQVVDTGVLPRLVELMTSSELNVLTPSLRTVGNIVTGTDEQTQVAIDAGVLNVLPQLLRHPKPSIQKEAAWALSNVAAGPCQHIQQLIACGLLPP
cKPNA7_15 : SNERIGQVVDTGVLPRLVELMTSSELNVLTPSLRTVGNIVTGTDEQTQVAIDAGVLNVLPQLLRHPKPSIQKEAAWALSNVAAGPCQHIQQLIACGLLPP
cKPNA7_21 : SNERIGQVVDTGVLPRLVELMTSSELNVLTPSLRTVGNIVTGTDEQTQVAIDAGVLNVLPQLLRHPKPSIQKEAAWALSNVAAGPCQHIQQLIACGLLPP
cKPNA7_23 : SNERIGQVVDTGVLPRLVELMTSSELNVLTPSLRTVGNIVTGTDEQTQVAIDAGVLNVLPQLLRHPKPSIQKEAAWALSNVAAGPCQHIQQLIACGLLPP
cKPNA7_2  : SNDRIGQVVDTGVLPRLVELMTSSELNVLTPSLRTVGNIVTGTDEQTQVAIDAGVLNVLPQLLRHPKPSIQKEAAWALSNVAAGPCQHIQQLIACGLLPP
cKPNA7_4  : SNDRIGQVVDTGVLPRLVELMTSSELNVLTPSLRTVGNIVTGTDEQTQVAIDAGVLNVLPQLLRHPKPSIQKEAAWALSNVAAGPCQHIQQLIACGLLPP
cKPNA7_10 : SNDRIGQVVDTGVLPRLVELMTSSELNVLTPSLRTVGNIVTGTDEQTQVAIDAGVLNVLPQLLRHPKPSIQKEAAWALSNVAAGPCQHIQQLIACGLLPP
cKPNA7_12 : SNDRIGQVVDTGVLPRLVELMTSSELNVLTPSLRTVGNIVTGTDEQTQVAIDAGVLNVLPQLLRHPKPSIQKEAAWALSNVAAGPCQHIQQLIACGLLPP
cKPNA7_18 : SNDRIGQVVDTGVLPRLVELMTSSELNVLTPSLRTVGNIVTGTDEQTQVAIDAGVLNVLPQLLRHPKPSIQKEAAWALSNVAAGPCQHIQQLIACGLLPP
cKPNA7_20 : SNDRIGQVVDTGVLPRLVELMTSSELNVLTPSLRTVGNIVTGTDEQTQVAIDAGVLNVLPQLLRHPKPSIQKEAAWALSNVAAGPCQHIQQLIACGLLPP
cKPNA7_6  : SNERIGQVVDTGVLPRLVELMTSSELNVLTPSLRTVGNIVTGTDEQTQVAIDAGVLNVLPQLLRHPKPSIQKEAAWALSNVAAGPCQHIQQLIACGLLPP
cKPNA7_8  : SNERIGQVVDTGVLPRLVELMTSSELNVLTPSLRTVGNIVTGTDEQTQVAIDAGVLNVLPQLLRHPKPSIQKEAAWALSNVAAGPCQHIQQLIACGLLPP
cKPNA7_14 : SNERIGQVVDTGVLPRLVELMTSSELNVLTPSLRTVGNIVTGTDEQTQVAIDAGVLNVLPQLLRHPKPSIQKEAAWALSNVAAGPCQHIQQLIACGLLPP
cKPNA7_16 : SNERIGQVVDTGVLPRLVELMTSSELNVLTPSLRTVGNIVTGTDEQTQVAIDAGVLNVLPQLLRHPKPSIQKEAAWALSNVAAGPCQHIQQLIACGLLPP
cKPNA2_1  : SNERIEVVVKTGLVPRLVKLLGCGELPIVTPALRAIGNIVTGTDEQTQVVIDAGALAVFPSLLRHPKTNIQKEAAWTMSNITAGRQDQIQEVVNHGLVPF
cKPNA2_2  : SNERIEVVVKTGLVPRLVKLLGCGELPIVTPALRAIGNIVTGTDEQTQVVIDAGALAVFPSLLRHPKTNIQKEAAWTMSNITAGRQDQIQEVVNHGLVPF
cKPNA3    : GNEQIQMVIDSGVVPFLVPLLSHQEVKVQTAALRAVGNIVTGTDEQTQVVLNCDVLSHFPNLLTHPKEKINKEAVWFLSNITAGNQQQVQAVIDAGLIPM
cKPNA4    : GNEQIQMVIDSGIVPHLVPLLSHQEVKVQTAALRAVGNIVTGTDEQTQVVLNCDALSHFPALLTHPKEKINKEAVWFLSNITAGNQQQVQAVIDANLVPM
cKPNA5    : PNDKIQAVIDSGVCRRLVELLMHNDYKVVSPALRAVGNIVTGDDIQTQVILNCSALPCLLHLLSSPKESIRKEACWTVSNITAGNRAQIQAVIDANIFPV
cKPNA6    : PNEKIQAVIDSGVCRRLVELLMHNDYKVASPALRAVGNIVTGDDIQTQVILNCSALPCLLHLLSSPKESIRKEACWTISNITAGNRAQIQAVIDANIFPV
cKPNA1    : PNDKIQAVIDAGVCRRLVELLMHNDYKVVSPALRAVGNIVTGDDIQTQVILNCSALQSLLHLLSSPKESIKKEACWTISNITAGNRAQIQTVIDANIFPA
```

```
                     *        420         *        440         *        460         *        480         *        500
cKPNA7_22 : LVALLKNGEFKVQKEAVWTVANFTTGGTVDQLIQLVHSGVLEPLLNLLTAPDSKIVLIILDVISNILQAAEKLSEKE----N-LCLLIEELGGLDKIEAL
cKPNA7_24 : LVALLKNGEFKVQKEAVWTVANFTTGGTVDQLIQLVHSGVLEPLLNLLTAPDSKIVLIILDVISNILQAAEKLSEKE----N-LCLLIEELGGLDKIEAL
cKPNA7_1  : LVALLKNGEFKVQKEAVWTVANFTTGGTVDQLIQLVHSGVLEPLLNLLTAPDSKIVLIILDVISNILQAAEKLSEKE----N-LCLLIEELGGLDKIEAL
cKPNA7_3  : LVALLKNGEFKVQKEAVWTVANFTTGGTVDQLIQLVHSGVLEPLLNLLTAPDSKIVLIILDVISNILQAAEKLSEKE----N-LCLLIEELGGLDKIEAL
cKPNA7_9  : LVALLKNGEFKVQKEAVWTVANFTTGGTVDQLIQLVHSGVLEPLLNLLTAPDSKIVLIILDVISNILQAAEKLSEKE----N-LCLLIEELGGLDKIEAL
cKPNA7_11 : LVALLKNGEFKVQKEAVWTVANFTTGGTVDQLIQLVHSGVLEPLLNLLTAPDSKIVLIILDVISNILQAAEKLSEKE----N-LCLLIEELGGLDKIEAL
cKPNA7_17 : LVALLKNGEFKVQKEAVWTVANFTTGGTVDQLIQLVHSGVLEPLLNLLTAPDSKIVLIILDVISNILQAAEKLSEKE----N-LCLLIEELGGLDKIEAL
cKPNA7_19 : LVALLKNGEFKVQKEAVWTVANFTTGGTVDQLIQLVHSGVLEPLLNLLTAPDSKIVLIILDVISNILQAAEKLSEKE----N-LCLLIEELGGLDKIEAL
cKPNA7_5  : LVALLKNGEFKVQKEAVWTVANFTTGGTVDQLIQLVHSGVLEPLLNLLTAPDSKIVLIILDVISNILQAAEKLSEKE----N-LCLLIEELGGLDKIEAL
cKPNA7_7  : LVALLKNGEFKVQKEAVWTVANFTTGGTVDQLIQLVHSGVLEPLLNLLTAPDSKIVLIILDVISNILQAAEKLSEKE----N-LCLLIEELGGLDKIEAL
cKPNA7_13 : LVALLKNGEFKVQKEAVWTVANFTTGGTVDQLIQLVHSGVLEPLLNLLTAPDSKIVLIILDVISNILQAAEKLSEKE----N-LCLLIEELGGLDKIEAL
cKPNA7_15 : LVALLKNGEFKVQKEAVWTVANFTTGGTVDQLIQLVHSGVLEPLLNLLTAPDSKIVLIILDVISNILQAAEKLSEKE----N-LCLLIEELGGLDKIEAL
cKPNA7_21 : LVALLKNGEFKVQKEAVWTVANFTTGGTVDQLIQLVHSGVLEPLLNLLTAPDSKIVLIILDVISNILQAAEKLSEKE----N-LCLLIEELGGLDKIEAL
cKPNA7_23 : LVALLKNGEFKVQKEAVWTVANFTTGGTVDQLIQLVHSGVLEPLLNLLTAPDSKIVLIILDVISNILQAAEKLSEKE----N-LCLLIEELGGLDKIEAL
cKPNA7_2  : LVALLKNGEFKVQKEAVWTVANFTTGGTVDQLIQLVHSGVLEPLLNLLTAPDSKIVLIILDVISNILQAAEKLSEKE----N-LCLLIEELGGLDKIEAL
cKPNA7_4  : LVALLKNGEFKVQKEAVWTVANFTTGGTVDQLIQLVHSGVLEPLLNLLTAPDSKIVLIILDVISNILQAAEKLSEKE----N-LCLLIEELGGLDKIEAL
cKPNA7_10 : LVALLKNGEFKVQKEAVWTVANFTTGGTVDQLIQLVHSGVLEPLLNLLTAPDSKIVLIILDVISNILQAAEKLSEKE----N-LCLLIEELGGLDKIEAL
cKPNA7_12 : LVALLKNGEFKVQKEAVWTVANFTTGGTVDQLIQLVHSGVLEPLLNLLTAPDSKIVLIILDVISNILQAAEKLSEKE----N-LCLLIEELGGLDKIEAL
cKPNA7_18 : LVALLKNGEFKVQKEAVWTVANFTTGGTVDQLIQLVHSGVLEPLLNLLTAPDSKIVLIILDVISNILQAAEKLSEKE----N-LCLLIEELGGLDKIEAL
cKPNA7_20 : LVALLKNGEFKVQKEAVWTVANFTTGGTVDQLIQLVHSGVLEPLLNLLTAPDSKIVLIILDVISNILQAAEKLSEKE----N-LCLLIEELGGLDKIEAL
cKPNA7_6  : LVALLKNGEFKVQKEAVWTVANFTTGGTVDQLIQLVHSGVLEPLLNLLTAPDSKIVLIILDVISNILQAAEKLSEKE----N-LCLLIEELGGLDKIEAL
cKPNA7_8  : LVALLKNGEFKVQKEAVWTVANFTTGGTVDQLIQLVHSGVLEPLLNLLTAPDSKIVLIILDVISNILQAAEKLSEKE----N-LCLLIEELGGLDKIEAL
cKPNA7_14 : LVALLKNGEFKVQKEAVWTVANFTTGGTVDQLIQLVHSGVLEPLLNLLTAPDSKIVLIILDVISNILQAAEKLSEKE----N-LCLLIEELGGLDKIEAL
cKPNA7_16 : LVALLKNGEFKVQKEAVWTVANFTTGGTVDQLIQLVHSGVLEPLLNLLTAPDSKIVLIILDVISNILQAAEKLSEKE----N-LCLLIEELGGLDKIEAL
cKPNA2_1  : LVEVLSKGDFKTQKEAVWAVTNYTSGGTVEQIVYLVHCGVLEPLLNLLTAKDSKIILVILDAISNIFQAAEKLGETE----K-LCLMIEECGGLDKIEAL
cKPNA2_2  : LVEVLSKGDFKTQKEAVWAVTNYTSGGTVEQIVYLVHCGVLEPLLNLLTAKDSKIILVILDAISNIFQAAEKLGETE----K-LCLMIEECGGLDKIEAL
cKPNA3    : IIHQLAKGDFGTQKEAAWAISNLTISGRKDQVEYLVQQNVIPPFCNLLSVKDSQVVQVVLDGLKNILIMAGD--EAS----T-IAEIIEECGGLEKIEVL
cKPNA4    : IIHLLDKGDFGTQKEAAWAISNLTISGRKDQVAYLIQQNVIPPFCNLLTVKDAQVVQVVLDGLSNILKMAED--EAE----T-IANLIEECGGLEKIEQL
cKPNA5    : LIEILQKAEFRTRKEAAWAITNATSGGTPEQIRYLVALGCIKPLCDLLTVMDSKIVQVALNGLENILRLGEQESKQNGIGINPYCALIEEAYGLDKIEFL
cKPNA6    : LIEILQKAEFRTRKEAAWAITNATSGGTPEQIRYLVSLGCIKPLCDLLTVMDSKIVQVALNGLENILRLGEQEGKRSGSGVNPYCGLIEEAYGLDKIEFL
cKPNA1    : LISILQTAEFRTRKEAAWAITNATSGGSAEQIKYLVELGCIKPLCDLLTVMDSKIVQVALNGLENILRLGEQEAKRNGTGINPYCALIEEAYGLDKIEFL
                     *        520         *        540         *        560
cKPNA7_22 : QLHENRQVYRAALNIIEKHFS----EEE---TLLPAL--DQD--HEFLL-----T-----
cKPNA7_24 : QLHENHQVYRAALNIIEKHFS----EEE---TLLPAL--DQD--HEFLL-----T-----
cKPNA7_1  : QLHENRQVARAALNIIEKHFS----EEE---TLLPAL--DQD--HEFLL-----T-----
cKPNA7_3  : QLHENHQVARAALNIIEKHFS----EEE---TLLPAL--DQD--HEFLL-----T-----
cKPNA7_9  : QLHENRQVARAALNIIEKHFS----EEE---TLLPAL--DQD--HEFLL-----T-----
cKPNA7_11 : QLHENHQVARAALNIIEKHFS----EEE---TLLPAL--DQD--HEFLL-----T-----
cKPNA7_17 : QLHENRQVARAALNIIEKHFS----EEE---TLLPAL--DQD--HEFLL-----T-----
cKPNA7_19 : QLHENHQVARAALNIIEKHFS----EEE---TLLPAL--DQD--HEFLL-----T-----
cKPNA7_5  : QLHENRQVARAALNIIEKHFS----EEE---TLLPAL--DQD--HEFLL-----T-----
cKPNA7_7  : QLHENHQVARAALNIIEKHFS----EEE---TLLPAL--DQD--HEFLL-----T-----
cKPNA7_13 : QLHENRQVARAALNIIEKHFS----EEE---TLLPAL--DQD--HEFLL-----T-----
cKPNA7_15 : QLHENHQVARAALNIIEKHFS----EEE---TLLPAL--DQD--HEFLL-----T-----
cKPNA7_21 : QLHENRQVARAALNIIEKHFS----EEE---TLLPAL--DQD--HEFLL-----T-----
cKPNA7_23 : QLHENHQVARAALNIIEKHFS----EEE---TLLPAL--DQD--HEFLL-----T-----
cKPNA7_2  : QLHENRQVYRAALNIIEKHFS----EEE---TLLPAL--DQD--HEFLL-----T-----
cKPNA7_4  : QLHENHQVYRAALNIIEKHFS----EEE---TLLPAL--DQD--HEFLL-----T-----
cKPNA7_10 : QLHENRQVYRAALNIIEKHFS----EEE---TLLPAL--DQD--HEFLL-----T-----
cKPNA7_12 : QLHENHQVYRAALNIIEKHFS----EEE---TLLPAL--DQD--HEFLL-----T-----
cKPNA7_18 : QLHENRQVYRAALNIIEKHFS----EEE---TLLPAL--DQD--HEFLL-----T-----
cKPNA7_20 : QLHENHQVYRAALNIIEKHFS----EEE---TLLPAL--DQD--HEFLL-----T-----
cKPNA7_6  : QLHENRQVYRAALNIIEKHFS----EEE---TLLPAL--DQD--HEFLL-----T-----
cKPNA7_8  : QLHENHQVYRAALNIIEKHFS----EEE---TLLPAL--DQD--HEFLL-----T-----
cKPNA7_14 : QLHENRQVYRAALNIIEKHFS----EEE---TLLPAL--DQD--HEFLL-----T-----
cKPNA7_16 : QLHENHQVYRAALNIIEKHFS----EEE---TLLPAL--DQD--HEFLL-----T-----
cKPNA2_1  : QSHENESVYKASLNLIEKYFSV---EEEEDQNVVPEA--TSDG-YTFQIQDGQPT--FNF
cKPNA2_2  : QSHENESVYKASLNLIEKYFSV---EEEEDQNVVPET--TSDG-YTFQIQDGQPT--FNF
cKPNA3    : QQHENEDIYKLAFEIIDQYFSGDDIDEDP--SLIPEA--TQGGTYNFDPTANLQTKEFNF
cKPNA4    : QNHENEDIYKLAYEIIDQFFSSDDIDEDP--SLVPEA--IQGGTFGFNSSANVPTEGFQF
cKPNA5    : QSHENQEIYQKAFDLIEHYFGVE--EDDP--SIVPQVDENQQQ-FVFQQQE-APMEGFQL
cKPNA6    : QSHENQEIYQKAFDLIEHYFGVE--DDDS--SLAPQVDETQQQ-FIFQQPE-APMEGFQL
cKPNA1    : QSHENQEIYQKAFDLIEHYFGTE--DEDS--SIAPQVDLSQQQ-YIFQQCE-APMEGFQL
```
