## Supporting file 4 for "Biochemical propensity mapping for structural and functional anatomy of importin α IBB domain"

| KPNA1 IBB domain | 10 | 11 | 12 | 14 | 17 | 23 | 24 | 26 | 27 | 28 | 30 | 31 | 32 | 33 | 34 | 35 | 36 | 37 | 38 | 39 | 40 | 41 | 42 | 43 | 44 | 45 | 46 | 47 | 48 | 49 |
|---|---|---|---|---|---|---|---|---|---|---|---|---|---|---|---|---|---|---|---|---|---|---|---|---|---|---|---|---|---|---|
| Ala | 0 | 2 | 4 | 0 | 0 | 0 | 1 | 0 | 36 | 0 | 0 | 0 | 0 | 0 | 0 | 0 | 0 | 1 | 0 | 0 | 1 | 2 | 0 | 0 | 0 | 0 | 0 | 0 | 0 | 0 |
| Arg | 0 | 0 | 0 | 1 | 0 | 4 | 0 | 0 | 1 | 158 | 0 | 7 | 1 | 0 | 0 | 0 | 1 | 0 | 0 | 0 | 0 | 0 | 0 | 0 | 161 | 159 | 157 | 161 | 0 | 0 |
| Asn | 0 | 0 | 1 | 0 | 2 | 0 | 0 | 158 | 0 | 0 | 0 | 0 | 0 | 0 | 0 | 160 | 0 | 1 | 0 | 156 | 0 | 0 | 1 | 0 | 0 | 0 | 0 | 0 | 0 | 0 |
| Asp | 0 | 0 | 0 | 1 | 0 | 0 | 39 | 2 | 0 | 0 | 0 | 1 | 0 | 0 | 0 | 1 | 0 | 0 | 0 | 3 | 0 | 155 | 0 | 0 | 0 | 0 | 0 | 0 | 0 | 0 |
| Cys | 0 | 0 | 0 | 0 | 0 | 0 | 0 | 0 | 0 | 0 | 0 | 0 | 0 | 0 | 0 | 0 | 0 | 0 | 0 | 0 | 0 | 0 | 0 | 0 | 0 | 0 | 0 | 0 | 0 | 0 |
| Gln | 0 | 0 | 0 | 30 | 0 | 0 | 1 | 0 | 1 | 1 | 1 | 0 | 0 | 0 | 1 | 0 | 0 | 0 | 0 | 0 | 1 | 0 | 0 | 0 | 0 | 0 | 1 | 0 | 1 | 0 |
| Glu | 0 | 0 | 0 | 0 | 1 | 1 | 118 | 0 | 1 | 0 | 0 | 0 | 0 | 0 | 0 | 0 | 0 | 0 | 0 | 0 | 0 | 3 | 157 | 0 | 0 | 1 | 0 | 0 | 156 | 160 |
| Gly | 0 | 0 | 0 | 1 | 148 | 1 | 1 | 0 | 0 | 0 | 0 | 0 | 0 | 0 | 0 | 0 | 0 | 3 | 1 | 0 | 0 | 0 | 1 | 0 | 0 | 0 | 0 | 0 | 0 | 0 |
| His | 1 | 0 | 0 | 0 | 1 | 0 | 0 | 1 | 0 | 0 | 0 | 0 | 0 | 0 | 0 | 0 | 0 | 0 | 0 | 0 | 0 | 0 | 0 | 0 | 0 | 0 | 0 | 0 | 0 | 0 |
| Ile | 1 | 0 | 0 | 0 | 0 | 1 | 0 | 0 | 0 | 0 | 0 | 0 | 0 | 0 | 0 | 0 | 0 | 0 | 0 | 0 | 0 | 0 | 0 | 0 | 0 | 0 | 0 | 0 | 2 | 0 |
| Leu | 0 | 1 | 1 | 0 | 0 | 0 | 0 | 0 | 2 | 0 | 158 | 0 | 0 | 0 | 0 | 0 | 0 | 0 | 156 | 0 | 0 | 0 | 0 | 5 | 0 | 0 | 0 | 0 | 0 | 0 |
| Lys | 0 | 1 | 0 | 3 | 1 | 152 | 0 | 0 | 0 | 1 | 0 | 150 | 2 | 0 | 160 | 0 | 160 | 0 | 4 | 0 | 0 | 0 | 1 | 0 | 0 | 1 | 3 | 0 | 0 | 0 |
| Met | 150 | 0 | 0 | 0 | 0 | 0 | 0 | 0 | 1 | 0 | 1 | 0 | 0 | 0 | 0 | 0 | 0 | 0 | 0 | 0 | 0 | 0 | 0 | 156 | 0 | 0 | 0 | 0 | 0 | 0 |
| Phe | 0 | 0 | 1 | 0 | 0 | 0 | 0 | 0 | 113 | 0 | 1 | 0 | 0 | 3 | 0 | 0 | 0 | 0 | 0 | 0 | 0 | 0 | 0 | 0 | 0 | 0 | 0 | 0 | 0 | 0 |
| Pro | 0 | 4 | 0 | 77 | 0 | 0 | 0 | 0 | 0 | 0 | 0 | 0 | 0 | 0 | 0 | 0 | 0 | 0 | 0 | 0 | 115 | 0 | 0 | 0 | 0 | 0 | 0 | 0 | 0 | 0 |
| Ser | 0 | 32 | 12 | 40 | 8 | 1 | 1 | 0 | 6 | 0 | 0 | 1 | 155 | 0 | 0 | 0 | 0 | 155 | 0 | 0 | 1 | 0 | 0 | 0 | 0 | 0 | 0 | 0 | 0 | 1 |
| Thr | 0 | 115 | 106 | 2 | 0 | 0 | 0 | 0 | 0 | 0 | 0 | 0 | 0 | 0 | 0 | 0 | 0 | 0 | 0 | 1 | 42 | 1 | 1 | 0 | 0 | 0 | 0 | 0 | 0 | 0 |
| Trp | 0 | 0 | 0 | 0 | 0 | 0 | 0 | 0 | 0 | 1 | 0 | 0 | 0 | 0 | 0 | 0 | 0 | 0 | 0 | 0 | 0 | 0 | 0 | 0 | 0 | 0 | 0 | 0 | 0 | 0 |
| Tyr | 1 | 0 | 0 | 0 | 0 | 0 | 0 | 0 | 0 | 0 | 0 | 0 | 0 | 158 | 0 | 0 | 0 | 0 | 0 | 1 | 1 | 0 | 0 | 0 | 0 | 0 | 0 | 0 | 0 | 0 |
| Val | 0 | 2 | 1 | 0 | 0 | 1 | 0 | 0 | 0 | 0 | 0 | 0 | 1 | 0 | 0 | 0 | 0 | 0 | 0 | 0 | 0 | 0 | 0 | 0 | 0 | 0 | 0 | 0 | 2 | 0 |
| - | 8 | 4 | 35 | 6 | 0 | 0 | 0 | 0 | 0 | 0 | 0 | 2 | 2 | 0 | 0 | 0 | 0 | 0 | 0 | 0 | 0 | 0 | 0 | 0 | 0 | 0 | 0 | 0 | 0 | 0 |
| X | 0 | 0 | 0 | 0 | 0 | 0 | 0 | 0 | 0 | 0 | 0 | 0 | 0 | 0 | 0 | 0 | 0 | 1 | 0 | 0 | 0 | 0 | 0 | 0 | 0 | 0 | 0 | 0 | 0 | 0 |
| Amino acid variations | 4 | 7 | 7 | 8 | 6 | 7 | 6 | 3 | 8 | 4 | 4 | 4 | 4 | 2 | 2 | 2 | 2 | 4 | 3 | 4 | 6 | 4 | 5 | 2 | 1 | 3 | 3 | 1 | 4 | 2 |
| Most frequent amino acid | M | T | T | P | G | K | E | N | F | R | L | K | S | Y | K | N | K | S | L | N | P | D | E | M | R | R | R | R | E | E |
| Conservation (%) | 93.2 | 71.4 | 65.8 | 47.8 | 91.9 | 94.4 | 73.3 | 98.1 | 70.2 | 98.1 | 98.1 | 93.2 | 96.3 | 98.1 | 99.4 | 99.4 | 99.4 | 96.9 | 96.9 | 96.9 | 71.4 | 96.3 | 97.5 | 96.9 | 100 | 98.8 | 97.5 | 100 | 96.9 | 99.4 |

| KPNA1 IBB domain | 50 | 51 | 52 | 53 | 54 | 55 | 56 | 57 | 58 | 59 | 60 | 61 | 62 | 63 | 68 | 69 | 70 | 74 | 75 | 80 | 82 | 85 | 86 | 87 | 88 | 89 | 96 |
|---|---|---|---|---|---|---|---|---|---|---|---|---|---|---|---|---|---|---|---|---|---|---|---|---|---|---|---|
| Ala | 0 | 0 | 0 | 0 | 0 | 0 | 0 | 3 | 0 | 0 | 0 | 0 | 0 | 0 | 0 | 0 | 0 | 0 | 0 | 0 | 148 | 0 | 116 | 0 | 1 | 4 | 26 |
| Arg | 0 | 0 | 0 | 0 | 0 | 161 | 0 | 0 | 0 | 122 | 0 | 0 | 0 | 0 | 0 | 0 | 153 | 153 | 0 | 0 | 0 | 0 | 0 | 0 | 0 | 0 | 0 |
| Asn | 0 | 1 | 0 | 0 | 0 | 0 | 0 | 0 | 0 | 0 | 1 | 0 | 0 | 0 | 0 | 0 | 0 | 5 | 155 | 0 | 0 | 4 | 1 | 2 | 0 | 0 | 1 |
| Asp | 0 | 0 | 0 | 0 | 0 | 0 | 0 | 0 | 0 | 0 | 40 | 0 | 0 | 0 | 0 | 0 | 0 | 0 | 0 | 0 | 0 | 1 | 1 | 8 | 25 | 25 | 2 |
| Cys | 1 | 1 | 0 | 0 | 0 | 0 | 0 | 0 | 0 | 0 | 0 | 0 | 0 | 0 | 0 | 0 | 0 | 0 | 0 | 2 | 0 | 4 | 0 | 0 | 0 | 0 | 5 |
| Gln | 0 | 0 | 0 | 157 | 0 | 0 | 0 | 156 | 0 | 0 | 0 | 0 | 160 | 0 | 0 | 0 | 0 | 0 | 0 | 0 | 1 | 0 | 0 | 2 | 0 | 0 | 0 |
| Glu | 156 | 0 | 0 | 4 | 0 | 0 | 0 | 0 | 0 | 0 | 120 | 161 | 0 | 0 | 0 | 0 | 0 | 1 | 0 | 0 | 0 | 1 | 0 | 140 | 124 | 122 | 0 |
| Gly | 0 | 156 | 0 | 0 | 0 | 0 | 0 | 1 | 0 | 0 | 0 | 0 | 0 | 0 | 0 | 1 | 1 | 0 | 0 | 0 | 2 | 2 | 0 | 0 | 0 | 0 | 26 |
| His | 0 | 0 | 0 | 0 | 0 | 0 | 0 | 0 | 1 | 0 | 0 | 0 | 0 | 0 | 0 | 1 | 1 | 0 | 0 | 0 | 0 | 0 | 0 | 0 | 1 | 0 | 0 |
| Ile | 0 | 0 | 0 | 0 | 0 | 0 | 0 | 0 | 0 | 0 | 0 | 0 | 0 | 3 | 0 | 0 | 1 | 1 | 0 | 6 | 2 | 0 | 3 | 3 | 0 | 0 | 0 |
| Leu | 0 | 0 | 156 | 0 | 161 | 0 | 0 | 0 | 1 | 0 | 0 | 0 | 0 | 148 | 4 | 0 | 0 | 1 | 1 | 2 | 5 | 0 | 4 | 0 | 0 | 0 | 0 |
| Lys | 0 | 0 | 0 | 0 | 0 | 0 | 161 | 0 | 159 | 39 | 0 | 0 | 0 | 0 | 0 | 155 | 2 | 0 | 0 | 0 | 0 | 0 | 0 | 0 | 1 | 1 | 0 |
| Met | 0 | 0 | 0 | 0 | 0 | 0 | 0 | 0 | 0 | 0 | 0 | 0 | 0 | 2 | 0 | 2 | 0 | 0 | 0 | 0 | 1 | 2 | 1 | 0 | 0 | 0 | 0 |
| Phe | 0 | 0 | 0 | 0 | 0 | 0 | 0 | 0 | 0 | 0 | 0 | 0 | 0 | 1 | 149 | 0 | 1 | 0 | 0 | 1 | 0 | 1 | 0 | 3 | 1 | 0 | 0 |
| Pro | 0 | 0 | 0 | 0 | 0 | 0 | 0 | 0 | 0 | 0 | 0 | 0 | 0 | 0 | 0 | 0 | 0 | 0 | 2 | 0 | 0 | 0 | 1 | 1 | 1 | 0 | 1 |
| Ser | 0 | 2 | 0 | 0 | 0 | 0 | 0 | 0 | 0 | 0 | 0 | 0 | 1 | 0 | 2 | 1 | 0 | 0 | 3 | 0 | 0 | 2 | 0 | 2 | 5 | 2 | 5 |
| Thr | 0 | 1 | 0 | 0 | 0 | 0 | 0 | 1 | 0 | 0 | 0 | 0 | 0 | 0 | 0 | 0 | 0 | 0 | 0 | 0 | 0 | 141 | 0 | 0 | 1 | 0 | 88 |
| Trp | 0 | 0 | 0 | 0 | 0 | 0 | 0 | 0 | 0 | 0 | 0 | 0 | 0 | 0 | 0 | 0 | 0 | 0 | 0 | 0 | 0 | 2 | 0 | 0 | 0 | 0 | 0 |
| Tyr | 0 | 0 | 0 | 0 | 0 | 0 | 0 | 0 | 0 | 0 | 0 | 0 | 0 | 0 | 0 | 0 | 0 | 0 | 0 | 0 | 0 | 1 | 0 | 0 | 0 | 1 | 0 |
| Val | 4 | 0 | 5 | 0 | 0 | 0 | 0 | 0 | 0 | 0 | 0 | 0 | 0 | 3 | 0 | 0 | 0 | 0 | 0 | 150 | 1 | 0 | 34 | 0 | 0 | 0 | 0 |
| - | 0 | 0 | 0 | 0 | 0 | 0 | 0 | 0 | 0 | 0 | 0 | 0 | 0 | 4 | 6 | 1 | 2 | 0 | 0 | 0 | 1 | 0 | 0 | 0 | 1 | 6 | 7 |
| X | 0 | 0 | 0 | 0 | 0 | 0 | 0 | 0 | 0 | 0 | 0 | 0 | 0 | 0 | 0 | 0 | 0 | 0 | 0 | 0 | 0 | 0 | 0 | 0 | 0 | 0 | 0 |
| Amino acid variations | 3 | 5 | 2 | 2 | 1 | 1 | 1 | 4 | 3 | 2 | 3 | 1 | 2 | 5 | 3 | 5 | 6 | 5 | 4 | 5 | 7 | 11 | 8 | 8 | 9 | 6 | 8 |
| Most frequent amino acid | E | G | L | Q | L | R | K | Q | K | R | E | E | Q | L | F | K | R | R | N | V | A | T | A | E | E | E | T |
| Conservation (%) | 96.9 | 96.9 | 96.9 | 97.5 | 100 | 100 | 100 | 96.9 | 98.8 | 75.8 | 74.5 | 100 | 99.4 | 91.9 | 92.5 | 96.3 | 95.0 | 95.0 | 96.3 | 93.2 | 91.9 | 87.6 | 72.0 | 87.0 | 77.0 | 75.8 | 54.7 |

| KPNA2 IBB domain | 8 | 9 | 12 | 23 | 29 | 31 | 32 | 34 | 38 | 40 | 41 | 43 | 44 | 45 | 46 | 47 | 49 | 50 | 51 | 52 | 53 | 54 | 55 | 60 | 61 | 62 | 63 | 64 | 65 | 66 |
|---|---|---|---|---|---|---|---|---|---|---|---|---|---|---|---|---|---|---|---|---|---|---|---|---|---|---|---|---|---|---|
| Ala | 1 | 2 | 41 | 1 | 2 | 2 | 86 | 12 | 50 | 86 | 0 | 0 | 1 | 1 | 0 | 0 | 0 | 2 | 1 | 0 | 0 | 54 | 1 | 1 | 0 | 0 | 0 | 0 | 0 | 46 |
| Arg | 0 | 0 | 0 | 4 | 1 | 6 | 0 | 0 | 0 | 0 | 183 | 0 | 6 | 92 | 0 | 0 | 0 | 5 | 0 | 9 | 1 | 0 | 0 | 0 | 0 | 179 | 185 | 183 | 186 | 0 |
| Asn | 1 | 1 | 3 | 117 | 0 | 132 | 0 | 77 | 0 | 2 | 0 | 0 | 110 | 4 | 0 | 0 | 183 | 2 | 2 | 0 | 7 | 0 | 53 | 0 | 0 | 0 | 0 | 0 | 0 | 5 |
| Asp | 0 | 2 | 0 | 17 | 59 | 1 | 0 | 6 | 6 | 4 | 0 | 0 | 2 | 0 | 0 | 0 | 2 | 0 | 0 | 0 | 172 | 0 | 9 | 3 | 0 | 0 | 0 | 0 | 0 | 0 |
| Cys | 0 | 0 | 0 | 1 | 0 | 0 | 0 | 1 | 0 | 0 | 0 | 0 | 2 | 0 | 0 | 0 | 0 | 0 | 0 | 0 | 3 | 0 | 0 | 0 | 2 | 0 | 1 | 0 | 0 | 0 |
| Gln | 0 | 0 | 0 | 1 | 0 | 2 | 2 | 1 | 0 | 0 | 0 | 0 | 0 | 65 | 0 | 2 | 0 | 1 | 0 | 1 | 0 | 1 | 0 | 0 | 7 | 0 | 0 | 0 | 0 | 3 |
| Glu | 1 | 7 | 0 | 0 | 114 | 7 | 0 | 2 | 0 | 0 | 0 | 0 | 0 | 1 | 0 | 0 | 0 | 0 | 0 | 0 | 0 | 0 | 6 | 179 | 0 | 0 | 0 | 0 | 0 | 0 |
| Gly | 0 | 0 | 3 | 7 | 0 | 4 | 2 | 50 | 0 | 1 | 0 | 0 | 2 | 0 | 0 | 0 | 0 | 0 | 182 | 0 | 0 | 0 | 3 | 0 | 0 | 0 | 0 | 0 | 0 | 0 |
| His | 0 | 0 | 0 | 0 | 2 | 0 | 1 | 2 | 1 | 1 | 0 | 0 | 22 | 0 | 0 | 0 | 0 | 0 | 0 | 0 | 0 | 0 | 1 | 0 | 0 | 0 | 1 | 1 | 0 | 0 |
| Ile | 0 | 0 | 2 | 1 | 0 | 2 | 2 | 0 | 0 | 0 | 0 | 10 | 0 | 0 | 0 | 0 | 0 | 0 | 0 | 0 | 0 | 1 | 5 | 0 | 0 | 0 | 0 | 0 | 0 | 91 |
| Leu | 1 | 0 | 1 | 3 | 0 | 0 | 8 | 0 | 2 | 0 | 0 | 171 | 1 | 3 | 0 | 0 | 0 | 0 | 0 | 0 | 0 | 0 | 2 | 0 | 74 | 2 | 0 | 0 | 0 | 0 |
| Lys | 0 | 0 | 0 | 2 | 4 | 0 | 0 | 0 | 0 | 0 | 0 | 0 | 1 | 15 | 0 | 183 | 0 | 175 | 0 | 177 | 0 | 0 | 0 | 4 | 0 | 0 | 0 | 2 | 1 | 7 |
| Met | 165 | 0 | 0 | 0 | 0 | 0 | 0 | 0 | 5 | 0 | 0 | 2 | 0 | 0 | 0 | 0 | 0 | 0 | 0 | 0 | 0 | 2 | 0 | 0 | 100 | 0 | 0 | 0 | 0 | 6 |
| Phe | 1 | 0 | 1 | 1 | 0 | 0 | 1 | 0 | 0 | 0 | 0 | 2 | 5 | 0 | 183 | 2 | 0 | 0 | 0 | 0 | 0 | 0 | 1 | 0 | 1 | 0 | 0 | 0 | 0 | 1 |
| Pro | 0 | 7 | 2 | 1 | 0 | 1 | 3 | 11 | 68 | 7 | 0 | 1 | 1 | 0 | 0 | 0 | 0 | 1 | 0 | 0 | 0 | 1 | 0 | 0 | 0 | 0 | 0 | 0 | 0 | 0 |
| Ser | 1 | 141 | 28 | 17 | 1 | 25 | 6 | 5 | 3 | 2 | 1 | 1 | 7 | 3 | 0 | 0 | 1 | 0 | 1 | 0 | 0 | 79 | 14 | 0 | 3 | 0 | 0 | 0 | 0 | 6 |
| Thr | 2 | 2 | 83 | 8 | 1 | 2 | 5 | 2 | 9 | 0 | 0 | 0 | 24 | 0 | 0 | 0 | 0 | 1 | 0 | 0 | 0 | 25 | 90 | 0 | 0 | 6 | 0 | 1 | 0 | 4 |
| Trp | 0 | 0 | 0 | 0 | 0 | 0 | 0 | 0 | 0 | 0 | 0 | 0 | 1 | 0 | 0 | 0 | 0 | 0 | 1 | 0 | 0 | 0 | 0 | 0 | 0 | 0 | 0 | 0 | 0 | 0 |
| Tyr | 0 | 0 | 0 | 0 | 0 | 0 | 0 | 0 | 0 | 0 | 0 | 0 | 1 | 1 | 4 | 0 | 0 | 0 | 0 | 0 | 2 | 2 | 0 | 0 | 0 | 0 | 0 | 0 | 0 | 0 |
| Val | 2 | 0 | 5 | 0 | 1 | 0 | 10 | 4 | 1 | 3 | 1 | 0 | 1 | 2 | 0 | 0 | 0 | 0 | 0 | 0 | 0 | 20 | 0 | 0 | 0 | 0 | 0 | 0 | 0 | 18 |
| - | 12 | 25 | 18 | 6 | 2 | 3 | 61 | 14 | 42 | 81 | 2 | 0 | 0 | 0 | 0 | 0 | 1 | 0 | 0 | 0 | 2 | 2 | 2 | 0 | 0 | 0 | 0 | 0 | 0 | 0 |
| X | 0 | 0 | 0 | 0 | 0 | 0 | 0 | 0 | 0 | 0 | 0 | 0 | 0 | 0 | 0 | 0 | 0 | 0 | 0 | 0 | 0 | 0 | 0 | 0 | 0 | 0 | 0 | 0 | 0 | 0 |
| Amino acid variations | 9 | 7 | 10 | 14 | 9 | 11 | 11 | 12 | 9 | 8 | 3 | 6 | 16 | 10 | 2 | 3 | 3 | 7 | 5 | 3 | 5 | 9 | 11 | 4 | 6 | 3 | 3 | 4 | 2 | 10 |
| Most frequent amino acid | M | S | T | N | E | N | A | N | P | A | R | L | N | R | F | K | N | K | G | K | D | S | T | E | M | R | R | R | R | I |
| Conservation (%) | 88.2 | 75.4 | 44.4 | 62.6 | 61.0 | 70.6 | 46.0 | 41.2 | 36.4 | 46.0 | 97.9 | 91.4 | 58.8 | 49.2 | 97.9 | 97.9 | 97.9 | 93.6 | 97.3 | 94.7 | 92.0 | 42.2 | 48.1 | 95.7 | 53.5 | 95.7 | 98.9 | 97.9 | 99.5 | 48.7 |

| KPNA2 IBB domain | 67 | 68 | 69 | 70 | 71 | 72 | 73 | 74 | 75 | 76 | 77 | 78 | 79 | 80 | 81 | 82 | 83 | 84 | 86 | 87 | 88 | 89 | 90 | 92 | 93 | 94 | 96 | 97 |
|---|---|---|---|---|---|---|---|---|---|---|---|---|---|---|---|---|---|---|---|---|---|---|---|---|---|---|---|---|
| Ala | 1 | 0 | 0 | 0 | 1 | 0 | 0 | 0 | 180 | 0 | 0 | 0 | 0 | 0 | 0 | 0 | 0 | 0 | 0 | 2 | 5 | 5 | 5 | 1 | 6 | 0 | 2 | 106 |
| Arg | 0 | 0 | 0 | 0 | 0 | 0 | 187 | 12 | 0 | 6 | 9 | 0 | 0 | 0 | 0 | 0 | 0 | 175 | 187 | 0 | 0 | 0 | 1 | 0 | 0 | 0 | 0 | 0 |
| Asn | 0 | 0 | 157 | 0 | 0 | 0 | 0 | 0 | 0 | 0 | 0 | 0 | 1 | 1 | 0 | 0 | 1 | 0 | 0 | 180 | 0 | 8 | 1 | 0 | 0 | 0 | 2 | 2 |
| Asp | 4 | 0 | 0 | 0 | 1 | 0 | 0 | 0 | 0 | 0 | 0 | 181 | 152 | 0 | 0 | 0 | 0 | 0 | 0 | 0 | 0 | 1 | 4 | 1 | 3 | 154 | 95 | 1 |
| Cys | 0 | 0 | 0 | 0 | 0 | 0 | 0 | 0 | 0 | 0 | 0 | 0 | 0 | 0 | 0 | 2 | 0 | 0 | 0 | 0 | 0 | 10 | 5 | 3 | 0 | 0 | 0 | 1 |
| Gln | 0 | 0 | 0 | 0 | 1 | 0 | 0 | 0 | 2 | 0 | 0 | 0 | 0 | 173 | 0 | 0 | 0 | 0 | 0 | 0 | 0 | 34 | 1 | 0 | 11 | 0 | 2 | 0 |
| Glu | 181 | 0 | 0 | 0 | 178 | 0 | 0 | 0 | 0 | 0 | 0 | 5 | 28 | 0 | 0 | 0 | 2 | 0 | 0 | 0 | 0 | 2 | 0 | 7 | 6 | 29 | 82 | 0 |
| Gly | 0 | 0 | 0 | 0 | 0 | 0 | 0 | 0 | 0 | 0 | 0 | 0 | 0 | 0 | 0 | 0 | 4 | 0 | 0 | 0 | 1 | 8 | 1 | 0 | 1 | 2 | 2 | 0 |
| His | 0 | 0 | 7 | 0 | 0 | 0 | 0 | 0 | 0 | 0 | 0 | 0 | 1 | 11 | 0 | 0 | 0 | 0 | 0 | 3 | 0 | 15 | 0 | 1 | 0 | 0 | 1 | 2 |
| Ile | 0 | 0 | 0 | 4 | 0 | 2 | 0 | 0 | 0 | 1 | 0 | 0 | 0 | 0 | 78 | 0 | 0 | 0 | 0 | 0 | 8 | 0 | 5 | 0 | 0 | 0 | 1 | 1 |
| Leu | 0 | 1 | 0 | 0 | 0 | 185 | 0 | 0 | 0 | 2 | 0 | 0 | 0 | 2 | 13 | 166 | 0 | 0 | 0 | 1 | 2 | 1 | 0 | 19 | 31 | 0 | 0 | 3 |
| Lys | 1 | 0 | 2 | 0 | 6 | 0 | 0 | 175 | 0 | 178 | 178 | 0 | 0 | 0 | 0 | 0 | 180 | 12 | 0 | 0 | 0 | 0 | 0 | 0 | 0 | 0 | 0 | 0 |
| Met | 0 | 0 | 0 | 1 | 0 | 0 | 0 | 0 | 0 | 0 | 0 | 0 | 0 | 0 | 94 | 0 | 0 | 0 | 0 | 0 | 0 | 0 | 0 | 0 | 1 | 0 | 0 | 0 |
| Phe | 0 | 14 | 0 | 0 | 0 | 0 | 0 | 0 | 0 | 0 | 0 | 0 | 0 | 0 | 0 | 18 | 0 | 0 | 0 | 0 | 2 | 0 | 0 | 84 | 0 | 0 | 0 | 0 |
| Pro | 0 | 0 | 0 | 0 | 0 | 0 | 0 | 0 | 0 | 0 | 0 | 0 | 0 | 0 | 0 | 0 | 0 | 0 | 0 | 0 | 0 | 0 | 0 | 1 | 120 | 0 | 0 | 64 |
| Ser | 0 | 1 | 16 | 0 | 0 | 0 | 0 | 0 | 1 | 0 | 0 | 0 | 0 | 0 | 0 | 1 | 0 | 0 | 0 | 1 | 0 | 99 | 103 | 0 | 3 | 1 | 0 | 3 |
| Thr | 0 | 3 | 4 | 0 | 0 | 0 | 0 | 0 | 4 | 0 | 0 | 0 | 0 | 0 | 0 | 0 | 0 | 0 | 0 | 0 | 0 | 2 | 49 | 0 | 1 | 0 | 0 | 0 |
| Trp | 0 | 0 | 0 | 0 | 0 | 0 | 0 | 0 | 0 | 0 | 0 | 0 | 0 | 0 | 0 | 0 | 0 | 0 | 0 | 0 | 0 | 0 | 1 | 1 | 0 | 0 | 0 | 0 |
| Tyr | 0 | 0 | 0 | 0 | 0 | 0 | 0 | 0 | 0 | 0 | 0 | 0 | 0 | 0 | 0 | 0 | 0 | 0 | 0 | 0 | 0 | 0 | 0 | 0 | 0 | 0 | 0 | 0 |
| Val | 0 | 168 | 1 | 182 | 0 | 0 | 0 | 0 | 0 | 0 | 0 | 1 | 5 | 0 | 0 | 0 | 0 | 0 | 0 | 0 | 169 | 2 | 2 | 5 | 1 | 1 | 0 | 1 |
| - | 0 | 0 | 0 | 0 | 0 | 0 | 0 | 0 | 0 | 0 | 0 | 0 | 0 | 0 | 2 | 0 | 0 | 0 | 0 | 0 | 0 | 0 | 9 | 64 | 3 | 0 | 0 | 3 |
| X | 0 | 0 | 0 | 0 | 0 | 0 | 0 | 0 | 0 | 0 | 0 | 0 | 0 | 0 | 0 | 0 | 0 | 0 | 0 | 0 | 0 | 0 | 0 | 0 | 0 | 0 | 0 | 0 |
| Amino acid variations | 4 | 5 | 6 | 3 | 5 | 2 | 1 | 2 | 4 | 4 | 2 | 3 | 5 | 4 | 3 | 4 | 4 | 2 | 1 | 5 | 6 | 12 | 12 | 10 | 11 | 5 | 8 | 10 |
| Most frequent amino acid | E | V | N | V | E | L | R | K | A | K | K | D | D | Q | M | L | K | R | R | N | V | S | S | F | P | D | D | A |
| Conservation (%) | 96.8 | 89.8 | 84.0 | 97.3 | 95.2 | 98.9 | 100 | 93.6 | 96.3 | 95.2 | 95.2 | 96.8 | 81.3 | 92.5 | 50.3 | 88.8 | 96.3 | 93.6 | 100 | 96.3 | 90.4 | 52.9 | 55.1 | 44.9 | 64.2 | 82.4 | 50.8 | 56.7 |

| KPNA3 IBB domain | 9 | 10 | 11 | 12 | 15 | 22 | 25 | 26 | 27 | 30 | 31 | 32 | 33 | 36 | 38 | 41 | 47 | 48 | 49 | 50 | 51 | 52 | 55 | 59 | 61 | 62 | 63 | 64 | 65 | 66 |
|---|---|---|---|---|---|---|---|---|---|---|---|---|---|---|---|---|---|---|---|---|---|---|---|---|---|---|---|---|---|---|
| Ala | 2 | 104 | 5 | 2 | 45 | 1 | 1 | 0 | 1 | 1 | 0 | 1 | 0 | 0 | 0 | 1 | 0 | 2 | 1 | 0 | 1 | 11 | 0 | 0 | 1 | 0 | 0 | 0 | 1 | 1 |
| Arg | 1 | 0 | 0 | 2 | 0 | 0 | 0 | 0 | 1 | 7 | 110 | 0 | 1 | 0 | 0 | 0 | 1 | 2 | 1 | 110 | 0 | 0 | 0 | 0 | 3 | 120 | 123 | 9 | 121 | 0 |
| Asn | 0 | 0 | 0 | 103 | 0 | 0 | 0 | 9 | 109 | 0 | 0 | 0 | 1 | 4 | 0 | 0 | 116 | 0 | 0 | 0 | 6 | 0 | 0 | 0 | 0 | 0 | 0 | 0 | 0 | 108 |
| Asp | 0 | 0 | 21 | 3 | 6 | 2 | 0 | 8 | 0 | 0 | 0 | 0 | 0 | 0 | 0 | 0 | 0 | 0 | 1 | 0 | 107 | 0 | 1 | 0 | 0 | 0 | 0 | 0 | 0 | 0 |
| Cys | 0 | 0 | 2 | 0 | 1 | 0 | 0 | 0 | 0 | 0 | 0 | 0 | 0 | 0 | 0 | 1 | 0 | 1 | 0 | 0 | 0 | 0 | 0 | 0 | 0 | 0 | 0 | 0 | 0 | 0 |
| Gln | 0 | 1 | 1 | 1 | 1 | 0 | 0 | 0 | 0 | 5 | 0 | 1 | 5 | 0 | 0 | 1 | 0 | 0 | 0 | 0 | 0 | 0 | 0 | 0 | 0 | 0 | 1 | 5 | 0 | 2 |
| Glu | 0 | 1 | 85 | 1 | 4 | 0 | 1 | 98 | 0 | 0 | 0 | 0 | 0 | 0 | 0 | 0 | 0 | 0 | 1 | 0 | 0 | 0 | 115 | 1 | 0 | 0 | 0 | 0 | 0 | 0 |
| Gly | 0 | 2 | 0 | 0 | 0 | 83 | 0 | 0 | 2 | 0 | 0 | 0 | 0 | 2 | 1 | 0 | 0 | 0 | 115 | 0 | 1 | 0 | 0 | 0 | 0 | 2 | 0 | 0 | 0 | 0 |
| His | 0 | 0 | 0 | 0 | 0 | 6 | 0 | 0 | 0 | 104 | 0 | 0 | 7 | 0 | 0 | 0 | 2 | 0 | 3 | 2 | 0 | 0 | 0 | 0 | 0 | 0 | 0 | 105 | 0 | 0 |
| Ile | 0 | 0 | 0 | 0 | 2 | 0 | 0 | 1 | 0 | 1 | 7 | 96 | 0 | 0 | 0 | 3 | 0 | 0 | 0 | 0 | 0 | 0 | 0 | 2 | 0 | 0 | 0 | 2 | 0 | 0 |
| Leu | 4 | 1 | 0 | 1 | 0 | 0 | 112 | 1 | 0 | 0 | 0 | 19 | 1 | 0 | 2 | 1 | 1 | 0 | 0 | 3 | 0 | 5 | 0 | 0 | 8 | 0 | 0 | 0 | 0 | 0 |
| Lys | 0 | 0 | 0 | 1 | 3 | 4 | 0 | 2 | 0 | 0 | 1 | 0 | 105 | 0 | 0 | 117 | 1 | 112 | 0 | 6 | 0 | 0 | 0 | 0 | 0 | 1 | 0 | 0 | 1 | 0 |
| Met | 108 | 0 | 6 | 0 | 0 | 0 | 0 | 0 | 0 | 0 | 0 | 1 | 0 | 0 | 0 | 0 | 0 | 0 | 0 | 0 | 0 | 0 | 0 | 1 | 109 | 0 | 0 | 0 | 0 | 0 |
| Phe | 7 | 1 | 0 | 0 | 0 | 0 | 1 | 0 | 1 | 0 | 3 | 4 | 0 | 0 | 120 | 0 | 0 | 0 | 1 | 0 | 0 | 1 | 0 | 0 | 0 | 0 | 0 | 0 | 0 | 0 |
| Pro | 0 | 1 | 0 | 0 | 49 | 0 | 0 | 1 | 4 | 0 | 0 | 0 | 0 | 0 | 0 | 0 | 0 | 2 | 0 | 0 | 1 | 2 | 3 | 0 | 0 | 0 | 0 | 0 | 0 | 0 |
| Ser | 0 | 4 | 0 | 8 | 1 | 17 | 0 | 3 | 4 | 2 | 2 | 0 | 1 | 109 | 0 | 0 | 1 | 0 | 0 | 1 | 2 | 0 | 1 | 1 | 3 | 0 | 0 | 0 | 1 | 2 |
| Thr | 0 | 8 | 1 | 1 | 0 | 1 | 1 | 1 | 1 | 0 | 0 | 1 | 2 | 0 | 0 | 0 | 0 | 0 | 0 | 0 | 3 | 3 | 1 | 108 | 0 | 0 | 0 | 0 | 0 | 11 |
| Trp | 0 | 0 | 0 | 0 | 0 | 0 | 0 | 0 | 0 | 0 | 0 | 0 | 0 | 0 | 0 | 0 | 0 | 0 | 0 | 0 | 0 | 0 | 0 | 0 | 0 | 0 | 0 | 0 | 0 | 0 |
| Tyr | 0 | 0 | 2 | 0 | 0 | 0 | 0 | 0 | 0 | 0 | 0 | 0 | 0 | 0 | 0 | 0 | 0 | 1 | 0 | 0 | 0 | 0 | 0 | 0 | 0 | 0 | 0 | 0 | 0 | 0 |
| Val | 2 | 0 | 1 | 1 | 0 | 0 | 7 | 0 | 0 | 0 | 0 | 1 | 0 | 1 | 0 | 0 | 0 | 1 | 0 | 1 | 0 | 97 | 0 | 4 | 0 | 0 | 0 | 0 | 0 | 0 |
| - | 0 | 1 | 0 | 0 | 12 | 10 | 1 | 0 | 1 | 4 | 1 | 0 | 1 | 8 | 1 | 0 | 2 | 3 | 1 | 1 | 3 | 5 | 3 | 7 | 0 | 1 | 0 | 3 | 0 | 0 |
| X | 0 | 0 | 0 | 0 | 0 | 0 | 0 | 0 | 0 | 0 | 0 | 0 | 0 | 0 | 0 | 0 | 0 | 0 | 0 | 0 | 0 | 0 | 0 | 0 | 0 | 0 | 0 | 0 | 0 | 0 |
| Amino acid variations | 6 | 9 | 9 | 11 | 9 | 7 | 6 | 9 | 8 | 6 | 5 | 8 | 8 | 4 | 3 | 6 | 6 | 7 | 7 | 6 | 7 | 6 | 5 | 6 | 5 | 3 | 2 | 4 | 4 | 5 |
| Most frequent amino acid | M | A | E | N | P | G | L | E | N | H | R | I | K | S | F | K | N | K | G | R | D | V | E | T | M | R | R | H | R | N |
| Conservation (%) | 87.1 | 83.9 | 68.5 | 83.1 | 39.5 | 66.9 | 90.3 | 79.0 | 87.9 | 83.9 | 88.7 | 77.4 | 84.7 | 87.9 | 96.8 | 94.4 | 93.5 | 90.3 | 92.7 | 88.7 | 86.3 | 78.2 | 92.7 | 87.1 | 87.9 | 96.8 | 99.2 | 84.7 | 97.6 | 87.1 |

| KPNA3 IBB domain | 67 | 68 | 69 | 73 | 74 | 75 | 76 | 79 | 80 | 81 | 82 | 83 | 84 | 88 | 89 | 93 | 94 | 95 | 96 | 97 | 98 | 99 | 100 | 102 | 103 | 104 | 106 | 108 |
|---|---|---|---|---|---|---|---|---|---|---|---|---|---|---|---|---|---|---|---|---|---|---|---|---|---|---|---|---|
| Ala | 3 | 2 | 0 | 0 | 0 | 0 | 0 | 0 | 1 | 0 | 0 | 0 | 0 | 1 | 0 | 0 | 0 | 0 | 0 | 0 | 0 | 0 | 0 | 0 | 0 | 1 | 0 | 0 |
| Arg | 0 | 0 | 0 | 0 | 0 | 0 | 120 | 1 | 0 | 1 | 121 | 0 | 0 | 0 | 0 | 0 | 0 | 10 | 123 | 0 | 0 | 0 | 0 | 0 | 0 | 0 | 0 | 0 |
| Asn | 0 | 0 | 0 | 0 | 0 | 0 | 0 | 1 | 115 | 0 | 0 | 0 | 0 | 0 | 0 | 0 | 1 | 0 | 0 | 116 | 0 | 0 | 1 | 1 | 0 | 3 | 0 | 0 |
| Asp | 9 | 0 | 0 | 0 | 0 | 0 | 0 | 1 | 0 | 0 | 0 | 113 | 7 | 0 | 0 | 0 | 0 | 0 | 0 | 0 | 0 | 0 | 0 | 2 | 10 | 1 | 0 | 2 |
| Cys | 0 | 0 | 0 | 0 | 0 | 0 | 0 | 0 | 0 | 0 | 0 | 0 | 0 | 0 | 0 | 0 | 0 | 0 | 0 | 0 | 0 | 0 | 0 | 0 | 0 | 0 | 5 | 0 |
| Gln | 0 | 0 | 1 | 0 | 0 | 0 | 0 | 1 | 0 | 0 | 2 | 0 | 2 | 0 | 0 | 3 | 0 | 0 | 1 | 0 | 0 | 0 | 102 | 0 | 7 | 0 | 0 | 1 |
| Glu | 111 | 0 | 0 | 0 | 117 | 0 | 0 | 1 | 0 | 0 | 0 | 11 | 113 | 2 | 0 | 0 | 0 | 0 | 0 | 0 | 0 | 0 | 0 | 110 | 107 | 0 | 0 | 120 |
| Gly | 0 | 1 | 0 | 0 | 2 | 0 | 0 | 0 | 0 | 0 | 0 | 0 | 0 | 0 | 0 | 0 | 0 | 0 | 0 | 0 | 0 | 0 | 1 | 3 | 0 | 0 | 0 | 0 |
| His | 0 | 0 | 0 | 0 | 0 | 0 | 0 | 0 | 0 | 0 | 0 | 0 | 0 | 111 | 0 | 0 | 0 | 0 | 0 | 0 | 0 | 1 | 12 | 0 | 0 | 0 | 0 | 0 |
| Ile | 0 | 4 | 0 | 22 | 0 | 1 | 0 | 0 | 0 | 0 | 0 | 0 | 0 | 0 | 1 | 0 | 0 | 0 | 0 | 0 | 4 | 0 | 1 | 0 | 0 | 5 | 0 | 0 |
| Leu | 0 | 1 | 0 | 1 | 0 | 121 | 0 | 0 | 0 | 0 | 0 | 0 | 0 | 0 | 122 | 111 | 0 | 0 | 0 | 1 | 4 | 0 | 5 | 0 | 0 | 0 | 107 | 0 |
| Lys | 0 | 0 | 0 | 0 | 0 | 0 | 0 | 120 | 0 | 123 | 1 | 0 | 1 | 0 | 0 | 0 | 123 | 114 | 0 | 0 | 0 | 0 | 0 | 0 | 0 | 0 | 0 | 0 |
| Met | 0 | 3 | 0 | 0 | 0 | 0 | 0 | 0 | 0 | 0 | 0 | 0 | 0 | 0 | 0 | 2 | 0 | 0 | 0 | 0 | 0 | 0 | 0 | 0 | 0 | 0 | 2 | 0 |
| Phe | 1 | 0 | 0 | 7 | 0 | 0 | 0 | 0 | 0 | 0 | 0 | 0 | 0 | 0 | 0 | 1 | 0 | 0 | 0 | 0 | 0 | 0 | 0 | 0 | 0 | 0 | 0 | 0 |
| Pro | 0 | 1 | 0 | 0 | 0 | 0 | 0 | 0 | 0 | 0 | 0 | 0 | 0 | 0 | 0 | 0 | 0 | 0 | 0 | 0 | 0 | 113 | 0 | 0 | 0 | 0 | 0 | 0 |
| Ser | 0 | 1 | 9 | 0 | 0 | 0 | 0 | 0 | 7 | 0 | 0 | 0 | 1 | 0 | 0 | 7 | 0 | 0 | 0 | 7 | 0 | 0 | 0 | 0 | 0 | 111 | 0 | 0 |
| Thr | 0 | 0 | 106 | 0 | 0 | 0 | 0 | 0 | 0 | 0 | 0 | 0 | 0 | 9 | 1 | 0 | 0 | 0 | 0 | 0 | 0 | 1 | 0 | 0 | 0 | 0 | 2 | 0 |
| Trp | 0 | 0 | 0 | 0 | 0 | 0 | 0 | 0 | 0 | 0 | 0 | 0 | 0 | 0 | 0 | 0 | 0 | 0 | 0 | 0 | 0 | 0 | 0 | 0 | 0 | 0 | 0 | 0 |
| Tyr | 0 | 0 | 0 | 0 | 0 | 0 | 0 | 0 | 0 | 0 | 0 | 0 | 0 | 1 | 0 | 0 | 0 | 0 | 0 | 0 | 0 | 0 | 0 | 0 | 0 | 0 | 0 | 0 |
| Val | 0 | 111 | 5 | 93 | 1 | 0 | 0 | 0 | 1 | 0 | 0 | 0 | 0 | 0 | 0 | 0 | 0 | 0 | 0 | 0 | 116 | 0 | 0 | 7 | 0 | 3 | 7 | 0 |
| - | 0 | 0 | 3 | 1 | 4 | 2 | 3 | 0 | 0 | 0 | 0 | 0 | 0 | 0 | 0 | 0 | 0 | 0 | 0 | 0 | 0 | 9 | 2 | 1 | 0 | 0 | 1 | 1 |
| X | 0 | 0 | 0 | 0 | 0 | 0 | 0 | 0 | 0 | 0 | 0 | 0 | 0 | 0 | 0 | 0 | 0 | 0 | 0 | 0 | 0 | 0 | 0 | 0 | 0 | 0 | 0 | 0 |
| Amino acid variations | 4 | 8 | 4 | 4 | 3 | 2 | 2 | 5 | 4 | 2 | 3 | 2 | 5 | 5 | 3 | 5 | 2 | 2 | 2 | 3 | 3 | 3 | 6 | 5 | 3 | 6 | 5 | 3 |
| Most frequent amino acid | E | V | T | V | E | L | R | K | N | K | R | D | E | H | L | L | K | K | R | N | V | P | Q | E | E | S | L | E |
| Conservation (%) | 89.5 | 89.5 | 85.5 | 75.0 | 94.4 | 97.6 | 96.8 | 96.8 | 92.7 | 99.2 | 97.6 | 91.1 | 91.1 | 89.5 | 98.4 | 89.5 | 99.2 | 91.9 | 99.2 | 93.5 | 93.5 | 91.1 | 82.3 | 88.7 | 86.3 | 89.5 | 86.3 | 96.8 |

| KPNA4 IBB domain | 9 | 10 | 20 | 21 | 33 | 34 | 35 | 36 | 37 | 38 | 39 | 40 | 41 | 43 | 44 | 46 | 47 | 48 | 50 | 52 | 58 | 59 | 63 | 76 | 77 | 78 | 79 | 80 | 81 | 82 |
|---|---|---|---|---|---|---|---|---|---|---|---|---|---|---|---|---|---|---|---|---|---|---|---|---|---|---|---|---|---|---|
| Ala | 0 | 117 | 1 | 1 | 1 | 4 | 0 | 0 | 1 | 5 | 0 | 3 | 1 | 1 | 0 | 0 | 1 | 0 | 1 | 5 | 1 | 2 | 1 | 1 | 0 | 0 | 1 | 1 | 1 | 0 |
| Arg | 1 | 2 | 1 | 1 | 1 | 2 | 0 | 0 | 0 | 12 | 120 | 0 | 8 | 0 | 0 | 1 | 0 | 4 | 2 | 119 | 0 | 0 | 1 | 0 | 0 | 148 | 146 | 11 | 147 | 0 |
| Asn | 0 | 3 | 3 | 106 | 7 | 0 | 0 | 0 | 121 | 2 | 4 | 0 | 0 | 121 | 0 | 0 | 127 | 0 | 0 | 1 | 1 | 0 | 1 | 0 | 0 | 0 | 0 | 0 | 0 | 111 |
| Asp | 0 | 0 | 119 | 0 | 6 | 0 | 0 | 125 | 0 | 1 | 9 | 0 | 0 | 0 | 0 | 1 | 0 | 3 | 1 | 0 | 128 | 1 | 6 | 0 | 0 | 0 | 0 | 0 | 0 | 0 |
| Cys | 1 | 1 | 0 | 0 | 1 | 0 | 0 | 0 | 1 | 0 | 0 | 2 | 0 | 1 | 6 | 0 | 2 | 0 | 8 | 1 | 0 | 1 | 0 | 0 | 0 | 0 | 0 | 0 | 0 | 0 |
| Gln | 0 | 1 | 2 | 5 | 0 | 1 | 1 | 0 | 0 | 119 | 1 | 0 | 6 | 0 | 0 | 3 | 0 | 1 | 0 | 0 | 0 | 9 | 2 | 0 | 0 | 0 | 1 | 136 | 0 | 0 |
| Glu | 0 | 4 | 8 | 1 | 111 | 0 | 1 | 4 | 0 | 1 | 0 | 0 | 0 | 0 | 0 | 2 | 0 | 2 | 0 | 0 | 0 | 0 | 121 | 10 | 0 | 0 | 0 | 0 | 0 | 1 |
| Gly | 0 | 2 | 4 | 3 | 0 | 1 | 1 | 0 | 1 | 0 | 1 | 2 | 3 | 0 | 0 | 6 | 0 | 1 | 128 | 1 | 0 | 0 | 0 | 0 | 1 | 0 | 0 | 0 | 0 | 0 |
| His | 0 | 0 | 0 | 2 | 1 | 0 | 0 | 1 | 1 | 3 | 0 | 0 | 0 | 1 | 0 | 0 | 1 | 0 | 0 | 4 | 0 | 0 | 0 | 0 | 0 | 0 | 0 | 0 | 0 | 0 |
| Ile | 0 | 0 | 0 | 0 | 0 | 0 | 0 | 0 | 7 | 0 | 0 | 0 | 2 | 4 | 0 | 1 | 0 | 1 | 0 | 3 | 0 | 0 | 0 | 0 | 0 | 0 | 0 | 0 | 0 | 0 |
| Leu | 5 | 0 | 0 | 5 | 2 | 0 | 127 | 0 | 4 | 0 | 0 | 121 | 0 | 1 | 2 | 1 | 0 | 0 | 0 | 2 | 1 | 132 | 0 | 0 | 3 | 0 | 0 | 0 | 0 | 0 |
| Lys | 0 | 0 | 0 | 0 | 2 | 126 | 0 | 1 | 1 | 1 | 3 | 0 | 121 | 2 | 0 | 127 | 0 | 129 | 2 | 8 | 1 | 0 | 1 | 0 | 0 | 0 | 0 | 0 | 0 | 0 |
| Met | 127 | 0 | 0 | 2 | 0 | 0 | 2 | 0 | 1 | 1 | 0 | 3 | 1 | 0 | 0 | 0 | 0 | 0 | 0 | 0 | 0 | 0 | 0 | 0 | 144 | 0 | 0 | 0 | 0 | 0 |
| Phe | 0 | 0 | 0 | 0 | 5 | 2 | 1 | 1 | 0 | 0 | 0 | 3 | 0 | 2 | 129 | 2 | 0 | 0 | 0 | 1 | 1 | 0 | 0 | 0 | 0 | 0 | 0 | 0 | 0 | 0 |
| Pro | 0 | 0 | 2 | 3 | 1 | 1 | 0 | 2 | 0 | 0 | 0 | 0 | 3 | 0 | 0 | 2 | 0 | 2 | 1 | 0 | 0 | 1 | 1 | 0 | 0 | 0 | 0 | 0 | 0 | 0 |
| Ser | 4 | 11 | 0 | 6 | 1 | 1 | 0 | 0 | 4 | 0 | 1 | 7 | 2 | 0 | 0 | 0 | 12 | 3 | 1 | 0 | 1 | 0 | 6 | 9 | 0 | 0 | 0 | 0 | 0 | 1 |
| Thr | 1 | 4 | 2 | 0 | 2 | 3 | 2 | 0 | 0 | 0 | 1 | 1 | 1 | 0 | 0 | 0 | 1 | 1 | 0 | 2 | 0 | 1 | 2 | 128 | 0 | 0 | 0 | 0 | 0 | 35 |
| Trp | 1 | 0 | 0 | 0 | 0 | 0 | 0 | 0 | 0 | 0 | 0 | 0 | 0 | 0 | 1 | 0 | 0 | 0 | 0 | 0 | 0 | 0 | 0 | 0 | 0 | 0 | 0 | 0 | 0 | 0 |
| Tyr | 0 | 0 | 0 | 0 | 0 | 0 | 0 | 0 | 2 | 0 | 0 | 1 | 0 | 1 | 3 | 0 | 0 | 0 | 0 | 2 | 0 | 0 | 0 | 0 | 0 | 0 | 0 | 0 | 0 | 0 |
| Val | 2 | 0 | 1 | 2 | 1 | 2 | 0 | 0 | 1 | 0 | 0 | 2 | 0 | 6 | 2 | 1 | 1 | 0 | 1 | 0 | 0 | 0 | 0 | 0 | 0 | 0 | 0 | 0 | 0 | 0 |
| - | 6 | 3 | 5 | 11 | 6 | 5 | 13 | 14 | 3 | 3 | 8 | 3 | 2 | 10 | 1 | 2 | 2 | 2 | 2 | 2 | 11 | 1 | 6 | 0 | 0 | 0 | 0 | 0 | 0 | 0 |
| X | 0 | 0 | 0 | 0 | 0 | 0 | 0 | 0 | 0 | 0 | 0 | 0 | 0 | 0 | 0 | 0 | 0 | 0 | 0 | 0 | 0 | 0 | 0 | 0 | 0 | 0 | 0 | 0 | 0 | 0 |
| Amino acid variations | 8 | 9 | 10 | 12 | 14 | 10 | 7 | 6 | 12 | 9 | 8 | 10 | 9 | 10 | 7 | 10 | 8 | 9 | 10 | 11 | 8 | 7 | 10 | 4 | 3 | 1 | 3 | 3 | 2 | 4 |
| Most frequent amino acid | M | A | D | N | E | K | L | D | N | Q | R | L | K | N | F | K | N | K | G | R | D | L | E | T | M | R | R | Q | R | N |
| Conservation (%) | 85.8 | 79.1 | 80.4 | 71.6 | 75.0 | 85.1 | 85.8 | 84.5 | 81.8 | 80.4 | 81.1 | 81.8 | 81.8 | 81.8 | 87.2 | 85.8 | 85.8 | 87.2 | 86.5 | 80.4 | 86.5 | 89.2 | 81.8 | 86.5 | 97.3 | 100 | 98.6 | 91.9 | 99.3 | 75.0 |

| KPNA4 IBB domain | 84 | 85 | 86 | 87 | 88 | 89 | 90 | 91 | 92 | 93 | 94 | 95 | 96 | 97 | 98 | 99 | 100 | 101 | 102 | 105 | 106 | 107 | 109 | 111 | 112 | 113 | 115 | 116 |
|---|---|---|---|---|---|---|---|---|---|---|---|---|---|---|---|---|---|---|---|---|---|---|---|---|---|---|---|---|
| Ala | 0 | 0 | 1 | 0 | 0 | 0 | 0 | 0 | 1 | 0 | 0 | 0 | 0 | 0 | 0 | 0 | 0 | 0 | 0 | 0 | 1 | 0 | 1 | 4 | 0 | 0 | 0 | 0 |
| Arg | 0 | 0 | 0 | 0 | 0 | 0 | 147 | 0 | 0 | 0 | 148 | 0 | 0 | 0 | 0 | 0 | 0 | 145 | 148 | 0 | 0 | 0 | 0 | 0 | 0 | 0 | 0 | 0 |
| Asn | 0 | 1 | 0 | 0 | 0 | 0 | 0 | 0 | 142 | 0 | 0 | 0 | 0 | 0 | 0 | 0 | 0 | 0 | 0 | 146 | 0 | 0 | 4 | 1 | 1 | 0 | 0 | 0 |
| Asp | 1 | 0 | 0 | 0 | 0 | 0 | 0 | 0 | 0 | 0 | 0 | 136 | 4 | 0 | 0 | 0 | 0 | 0 | 0 | 0 | 0 | 0 | 0 | 1 | 145 | 0 | 0 | 12 |
| Cys | 0 | 0 | 0 | 0 | 0 | 0 | 0 | 0 | 1 | 0 | 0 | 0 | 0 | 1 | 0 | 0 | 0 | 0 | 0 | 0 | 0 | 0 | 0 | 0 | 0 | 0 | 136 | 0 |
| Gln | 0 | 0 | 0 | 0 | 1 | 0 | 1 | 0 | 2 | 0 | 0 | 0 | 0 | 0 | 0 | 5 | 0 | 0 | 0 | 0 | 0 | 0 | 10 | 2 | 0 | 0 | 0 | 0 |
| Glu | 147 | 0 | 0 | 0 | 146 | 0 | 0 | 0 | 0 | 0 | 0 | 8 | 144 | 0 | 0 | 0 | 0 | 0 | 0 | 0 | 0 | 0 | 0 | 136 | 0 | 0 | 0 | 132 |
| Gly | 0 | 0 | 0 | 0 | 0 | 0 | 0 | 0 | 0 | 0 | 0 | 1 | 0 | 0 | 0 | 0 | 0 | 0 | 0 | 1 | 0 | 0 | 0 | 0 | 0 | 0 | 0 | 0 |
| His | 0 | 0 | 0 | 0 | 0 | 0 | 0 | 0 | 0 | 0 | 0 | 0 | 0 | 137 | 0 | 0 | 0 | 0 | 0 | 0 | 0 | 0 | 124 | 0 | 0 | 0 | 0 | 0 |
| Ile | 0 | 0 | 0 | 3 | 0 | 0 | 0 | 0 | 0 | 0 | 0 | 0 | 0 | 0 | 2 | 0 | 0 | 0 | 0 | 0 | 1 | 0 | 2 | 0 | 0 | 130 | 1 | 0 |
| Leu | 0 | 1 | 0 | 0 | 0 | 148 | 0 | 0 | 0 | 0 | 0 | 0 | 0 | 0 | 144 | 138 | 0 | 0 | 0 | 0 | 0 | 0 | 0 | 0 | 0 | 1 | 1 | 0 |
| Lys | 0 | 0 | 0 | 0 | 0 | 0 | 0 | 147 | 0 | 148 | 0 | 0 | 0 | 0 | 0 | 0 | 148 | 3 | 0 | 0 | 0 | 0 | 0 | 0 | 0 | 0 | 0 | 0 |
| Met | 0 | 2 | 0 | 0 | 0 | 0 | 0 | 0 | 0 | 0 | 0 | 0 | 0 | 0 | 0 | 0 | 0 | 0 | 0 | 0 | 0 | 0 | 0 | 2 | 0 | 1 | 0 | 0 |
| Phe | 0 | 0 | 1 | 2 | 0 | 0 | 0 | 0 | 0 | 0 | 0 | 0 | 0 | 0 | 1 | 3 | 0 | 0 | 0 | 0 | 0 | 0 | 0 | 0 | 0 | 2 | 0 | 0 |
| Pro | 0 | 0 | 0 | 0 | 1 | 0 | 0 | 0 | 0 | 0 | 0 | 0 | 0 | 0 | 0 | 0 | 0 | 0 | 0 | 0 | 0 | 147 | 0 | 0 | 0 | 0 | 0 | 0 |
| Ser | 0 | 0 | 3 | 0 | 0 | 0 | 0 | 0 | 2 | 0 | 0 | 0 | 0 | 1 | 0 | 2 | 0 | 0 | 0 | 1 | 0 | 0 | 1 | 0 | 1 | 7 | 0 | 0 |
| Thr | 0 | 0 | 9 | 0 | 0 | 0 | 0 | 0 | 0 | 0 | 0 | 0 | 0 | 8 | 0 | 0 | 0 | 0 | 0 | 0 | 0 | 0 | 2 | 1 | 0 | 1 | 7 | 0 |
| Trp | 0 | 0 | 0 | 0 | 0 | 0 | 0 | 0 | 0 | 0 | 0 | 0 | 0 | 0 | 0 | 0 | 0 | 0 | 0 | 0 | 0 | 0 | 0 | 0 | 0 | 0 | 0 | 0 |
| Tyr | 0 | 1 | 0 | 0 | 0 | 0 | 0 | 0 | 0 | 0 | 0 | 0 | 0 | 0 | 0 | 0 | 0 | 0 | 0 | 0 | 0 | 0 | 2 | 0 | 1 | 0 | 0 | 0 |
| Val | 0 | 143 | 134 | 143 | 0 | 0 | 0 | 0 | 0 | 0 | 0 | 3 | 0 | 1 | 1 | 0 | 0 | 0 | 0 | 0 | 146 | 0 | 1 | 1 | 0 | 6 | 1 | 0 |
| - | 0 | 0 | 0 | 0 | 0 | 0 | 0 | 1 | 0 | 0 | 0 | 0 | 0 | 0 | 0 | 0 | 0 | 0 | 0 | 0 | 0 | 1 | 1 | 0 | 0 | 0 | 2 | 4 |
| X | 0 | 0 | 0 | 0 | 0 | 0 | 0 | 0 | 0 | 0 | 0 | 0 | 0 | 0 | 0 | 0 | 0 | 0 | 0 | 0 | 0 | 0 | 0 | 0 | 0 | 0 | 0 | 0 |
| Amino acid variations | 2 | 5 | 5 | 3 | 3 | 1 | 2 | 2 | 5 | 1 | 1 | 4 | 2 | 5 | 4 | 4 | 1 | 2 | 1 | 3 | 3 | 1 | 9 | 8 | 4 | 7 | 5 | 2 |
| Most frequent amino acid | E | V | V | V | E | L | R | K | N | K | R | D | E | H | L | L | K | R | R | N | V | P | H | E | D | I | C | E |
| Conservation (%) | 99.3 | 96.6 | 90.5 | 96.6 | 98.6 | 100 | 99.3 | 99.3 | 95.9 | 100 | 100 | 91.9 | 97.3 | 92.6 | 97.3 | 93.2 | 100 | 98.0 | 100 | 98.6 | 98.6 | 99.3 | 83.8 | 91.9 | 98.0 | 87.8 | 91.9 | 89.2 |

| KPNA5 IBB domain | 7 | 9 | 10 | 11 | 12 | 13 | 15 | 18 | 19 | 20 | 23 | 24 | 25 | 26 | 27 | 28 | 29 | 30 | 31 | 32 | 33 | 34 | 35 | 36 | 37 | 38 | 39 | 40 | 41 | 42 |
|---|---|---|---|---|---|---|---|---|---|---|---|---|---|---|---|---|---|---|---|---|---|---|---|---|---|---|---|---|---|---|
| Ala | 2 | 3 | 70 | 0 | 133 | 1 | 0 | 1 | 1 | 0 | 0 | 0 | 0 | 0 | 1 | 0 | 0 | 0 | 0 | 1 | 144 | 0 | 0 | 2 | 0 | 0 | 1 | 0 | 0 | 0 |
| Arg | 2 | 1 | 0 | 0 | 0 | 0 | 0 | 0 | 3 | 0 | 0 | 0 | 150 | 1 | 9 | 1 | 0 | 0 | 0 | 0 | 0 | 0 | 0 | 0 | 0 | 0 | 0 | 150 | 149 | 150 |
| Asn | 1 | 4 | 16 | 0 | 0 | 0 | 0 | 0 | 0 | 0 | 134 | 0 | 0 | 0 | 1 | 6 | 0 | 0 | 148 | 28 | 0 | 0 | 144 | 0 | 0 | 0 | 0 | 0 | 0 | 0 |
| Asp | 0 | 90 | 2 | 0 | 0 | 1 | 0 | 0 | 0 | 143 | 0 | 0 | 0 | 0 | 0 | 0 | 0 | 0 | 0 | 0 | 0 | 0 | 4 | 0 | 0 | 0 | 0 | 0 | 0 | 0 |
| Cys | 0 | 0 | 0 | 0 | 0 | 1 | 0 | 0 | 0 | 0 | 0 | 3 | 0 | 0 | 0 | 0 | 0 | 0 | 0 | 0 | 0 | 0 | 0 | 0 | 0 | 0 | 0 | 0 | 0 | 0 |
| Gln | 0 | 0 | 0 | 0 | 0 | 0 | 0 | 0 | 0 | 1 | 0 | 0 | 0 | 0 | 0 | 0 | 0 | 0 | 0 | 0 | 0 | 0 | 0 | 0 | 147 | 0 | 0 | 0 | 0 | 0 |
| Glu | 0 | 0 | 2 | 0 | 0 | 0 | 0 | 8 | 0 | 2 | 2 | 0 | 0 | 1 | 0 | 0 | 0 | 0 | 0 | 0 | 0 | 0 | 0 | 0 | 2 | 150 | 0 | 0 | 0 | 0 |
| Gly | 0 | 2 | 0 | 0 | 0 | 1 | 0 | 126 | 0 | 0 | 0 | 0 | 0 | 0 | 0 | 1 | 0 | 0 | 0 | 0 | 3 | 0 | 0 | 1 | 0 | 0 | 1 | 0 | 0 | 0 |
| His | 3 | 1 | 0 | 0 | 0 | 0 | 0 | 0 | 0 | 1 | 2 | 1 | 0 | 0 | 0 | 0 | 0 | 0 | 0 | 0 | 0 | 2 | 0 | 0 | 0 | 0 | 0 | 0 | 0 | 0 |
| Ile | 5 | 0 | 3 | 0 | 0 | 1 | 0 | 0 | 0 | 0 | 0 | 0 | 0 | 3 | 0 | 0 | 0 | 0 | 0 | 1 | 0 | 0 | 1 | 0 | 0 | 0 | 0 | 0 | 0 | 0 |
| Leu | 2 | 0 | 1 | 1 | 0 | 4 | 3 | 0 | 0 | 0 | 0 | 0 | 0 | 0 | 0 | 0 | 0 | 0 | 0 | 1 | 0 | 146 | 0 | 0 | 0 | 0 | 2 | 0 | 0 | 0 |
| Lys | 1 | 0 | 1 | 0 | 0 | 0 | 0 | 7 | 144 | 1 | 1 | 0 | 0 | 1 | 138 | 0 | 0 | 150 | 0 | 118 | 0 | 0 | 0 | 0 | 0 | 0 | 0 | 0 | 1 | 0 |
| Met | 76 | 1 | 6 | 143 | 0 | 0 | 0 | 0 | 0 | 0 | 0 | 0 | 0 | 142 | 0 | 1 | 0 | 0 | 0 | 0 | 0 | 1 | 0 | 0 | 0 | 0 | 146 | 0 | 0 | 0 |
| Phe | 0 | 0 | 0 | 0 | 1 | 0 | 0 | 0 | 0 | 0 | 0 | 17 | 0 | 1 | 0 | 0 | 0 | 0 | 0 | 0 | 0 | 0 | 0 | 0 | 0 | 0 | 0 | 0 | 0 | 0 |
| Pro | 1 | 0 | 0 | 0 | 3 | 0 | 143 | 0 | 0 | 0 | 0 | 0 | 0 | 0 | 0 | 0 | 0 | 0 | 0 | 0 | 0 | 0 | 0 | 135 | 0 | 0 | 0 | 0 | 0 | 0 |
| Ser | 3 | 1 | 3 | 1 | 8 | 138 | 2 | 1 | 0 | 0 | 8 | 0 | 0 | 0 | 0 | 139 | 0 | 0 | 0 | 0 | 0 | 0 | 0 | 9 | 0 | 0 | 0 | 0 | 0 | 0 |
| Thr | 0 | 0 | 12 | 2 | 1 | 0 | 1 | 1 | 1 | 0 | 1 | 0 | 0 | 0 | 0 | 0 | 0 | 0 | 1 | 0 | 0 | 0 | 1 | 3 | 0 | 0 | 0 | 0 | 0 | 0 |
| Trp | 0 | 0 | 0 | 0 | 0 | 0 | 0 | 0 | 0 | 0 | 0 | 0 | 0 | 0 | 0 | 0 | 0 | 0 | 0 | 0 | 0 | 0 | 0 | 0 | 0 | 0 | 0 | 0 | 0 | 0 |
| Tyr | 0 | 1 | 0 | 0 | 0 | 1 | 0 | 0 | 0 | 0 | 0 | 126 | 0 | 0 | 0 | 0 | 150 | 0 | 0 | 0 | 0 | 0 | 0 | 0 | 0 | 0 | 0 | 0 | 0 | 0 |
| Val | 0 | 1 | 0 | 1 | 1 | 1 | 0 | 4 | 0 | 0 | 0 | 2 | 0 | 0 | 0 | 0 | 0 | 0 | 1 | 1 | 3 | 0 | 0 | 0 | 1 | 0 | 0 | 0 | 0 | 0 |
| - | 54 | 45 | 34 | 2 | 3 | 1 | 1 | 2 | 1 | 2 | 2 | 1 | 0 | 1 | 1 | 2 | 0 | 0 | 0 | 0 | 0 | 1 | 0 | 0 | 0 | 0 | 0 | 0 | 0 | 0 |
| X | 0 | 0 | 0 | 0 | 0 | 0 | 0 | 0 | 0 | 0 | 0 | 0 | 0 | 0 | 0 | 0 | 0 | 0 | 0 | 0 | 0 | 0 | 0 | 0 | 0 | 0 | 0 | 0 | 0 | 0 |
| Amino acid variations | 10 | 10 | 10 | 5 | 6 | 9 | 4 | 7 | 4 | 5 | 6 | 5 | 1 | 6 | 4 | 5 | 1 | 1 | 3 | 6 | 3 | 3 | 4 | 5 | 3 | 1 | 4 | 1 | 2 | 1 |
| Most frequent amino acid | M | D | A | M | A | S | P | G | K | D | N | Y | R | M | K | S | Y | K | N | K | A | L | N | P | Q | E | M | R | R | R |
| Conservation (%) | 50.7 | 60.0 | 46.7 | 95.3 | 88.7 | 92.0 | 95.3 | 84.0 | 96.0 | 95.3 | 89.3 | 84.0 | 100 | 94.7 | 92.0 | 92.7 | 100 | 100 | 98.7 | 78.7 | 96.0 | 97.3 | 96.0 | 90.0 | 98.0 | 100 | 97.3 | 100 | 99.3 | 100 |

| KPNA5 IBB domain | 43 | 44 | 45 | 46 | 47 | 48 | 49 | 50 | 51 | 52 | 53 | 54 | 55 | 56 | 57 | 58 | 59 | 62 | 63 | 64 | 65 | 66 | 67 | 73 | 74 | 75 | 76 | 77 | 78 | 79 |
|---|---|---|---|---|---|---|---|---|---|---|---|---|---|---|---|---|---|---|---|---|---|---|---|---|---|---|---|---|---|---|
| Ala | 0 | 0 | 0 | 0 | 0 | 1 | 0 | 0 | 0 | 0 | 0 | 0 | 0 | 1 | 0 | 0 | 0 | 0 | 0 | 0 | 0 | 0 | 0 | 1 | 2 | 0 | 2 | 7 | 0 | 0 |
| Arg | 148 | 0 | 0 | 0 | 0 | 0 | 0 | 0 | 150 | 0 | 0 | 0 | 150 | 0 | 0 | 0 | 0 | 0 | 0 | 138 | 140 | 1 | 0 | 1 | 0 | 0 | 72 | 0 | 0 | 0 |
| Asn | 0 | 0 | 0 | 3 | 0 | 0 | 0 | 0 | 0 | 0 | 1 | 0 | 0 | 0 | 1 | 1 | 0 | 0 | 0 | 0 | 0 | 138 | 0 | 0 | 1 | 0 | 2 | 103 | 2 | 4 |
| Asp | 0 | 0 | 2 | 2 | 0 | 0 | 0 | 0 | 0 | 0 | 0 | 0 | 0 | 0 | 0 | 0 | 0 | 0 | 0 | 0 | 0 | 0 | 1 | 0 | 0 | 1 | 1 | 1 | 102 | 9 |
| Cys | 0 | 0 | 0 | 0 | 0 | 0 | 0 | 0 | 0 | 0 | 0 | 0 | 0 | 0 | 0 | 0 | 0 | 0 | 0 | 0 | 0 | 0 | 0 | 14 | 0 | 1 | 2 | 2 | 0 | 0 |
| Gln | 0 | 0 | 0 | 0 | 0 | 0 | 148 | 0 | 0 | 0 | 148 | 0 | 0 | 0 | 1 | 146 | 0 | 0 | 0 | 0 | 0 | 0 | 0 | 1 | 0 | 0 | 0 | 2 | 1 | 3 |
| Glu | 0 | 150 | 147 | 145 | 0 | 0 | 1 | 0 | 0 | 0 | 0 | 0 | 0 | 149 | 146 | 2 | 0 | 5 | 5 | 0 | 0 | 0 | 1 | 0 | 0 | 0 | 1 | 1 | 31 | 125 |
| Gly | 0 | 0 | 0 | 0 | 147 | 0 | 0 | 0 | 0 | 0 | 0 | 0 | 0 | 0 | 0 | 0 | 0 | 0 | 0 | 0 | 0 | 0 | 0 | 3 | 0 | 0 | 26 | 6 | 0 | 1 |
| His | 0 | 0 | 0 | 0 | 0 | 0 | 0 | 0 | 0 | 0 | 0 | 0 | 0 | 0 | 0 | 0 | 0 | 0 | 0 | 0 | 0 | 0 | 0 | 0 | 0 | 0 | 0 | 0 | 0 | 1 |
| Ile | 0 | 0 | 0 | 0 | 0 | 145 | 0 | 2 | 0 | 0 | 0 | 0 | 0 | 0 | 0 | 0 | 1 | 0 | 0 | 0 | 3 | 4 | 1 | 0 | 6 | 0 | 0 | 1 | 1 | 0 |
| Leu | 0 | 0 | 0 | 0 | 1 | 0 | 0 | 148 | 0 | 0 | 0 | 0 | 0 | 0 | 0 | 0 | 139 | 2 | 0 | 0 | 5 | 0 | 4 | 3 | 110 | 13 | 1 | 1 | 1 | 0 |
| Lys | 0 | 0 | 0 | 0 | 0 | 0 | 0 | 0 | 0 | 150 | 0 | 150 | 0 | 0 | 0 | 0 | 0 | 0 | 145 | 2 | 0 | 1 | 0 | 0 | 0 | 1 | 2 | 0 | 0 | 0 |
| Met | 0 | 0 | 0 | 0 | 1 | 0 | 0 | 0 | 0 | 0 | 0 | 0 | 0 | 0 | 0 | 0 | 0 | 0 | 0 | 0 | 0 | 0 | 0 | 5 | 2 | 0 | 0 | 0 | 0 | 0 |
| Phe | 0 | 0 | 0 | 0 | 0 | 0 | 0 | 0 | 0 | 0 | 0 | 0 | 0 | 0 | 0 | 0 | 0 | 139 | 0 | 0 | 0 | 0 | 1 | 1 | 2 | 5 | 0 | 0 | 0 | 0 |
| Pro | 0 | 0 | 0 | 0 | 0 | 0 | 0 | 0 | 0 | 0 | 0 | 0 | 0 | 0 | 0 | 0 | 0 | 0 | 0 | 0 | 0 | 0 | 0 | 0 | 0 | 126 | 2 | 1 | 1 | 0 |
| Ser | 1 | 0 | 0 | 0 | 0 | 0 | 0 | 0 | 0 | 0 | 1 | 0 | 0 | 0 | 2 | 1 | 0 | 1 | 0 | 0 | 0 | 0 | 1 | 117 | 1 | 0 | 33 | 21 | 1 | 0 |
| Thr | 0 | 0 | 1 | 0 | 1 | 0 | 0 | 0 | 0 | 0 | 0 | 0 | 0 | 0 | 0 | 0 | 0 | 0 | 0 | 0 | 0 | 6 | 5 | 0 | 0 | 1 | 1 | 2 | 1 | 0 |
| Trp | 0 | 0 | 0 | 0 | 0 | 0 | 0 | 0 | 0 | 0 | 0 | 0 | 0 | 0 | 0 | 0 | 0 | 0 | 0 | 0 | 0 | 0 | 0 | 0 | 0 | 0 | 0 | 0 | 0 | 0 |
| Tyr | 0 | 0 | 0 | 0 | 0 | 0 | 0 | 0 | 0 | 0 | 0 | 0 | 0 | 0 | 0 | 0 | 0 | 2 | 0 | 5 | 0 | 0 | 0 | 2 | 2 | 0 | 0 | 0 | 0 | 0 |
| Val | 0 | 0 | 0 | 0 | 0 | 4 | 1 | 0 | 0 | 0 | 0 | 0 | 0 | 0 | 0 | 0 | 7 | 0 | 0 | 5 | 2 | 0 | 136 | 1 | 24 | 0 | 3 | 0 | 0 | 0 |
| - | 1 | 0 | 0 | 0 | 0 | 0 | 0 | 0 | 0 | 0 | 0 | 0 | 0 | 0 | 0 | 0 | 3 | 1 | 0 | 0 | 0 | 0 | 0 | 1 | 0 | 2 | 2 | 2 | 9 | 7 |
| X | 0 | 0 | 0 | 0 | 0 | 0 | 0 | 0 | 0 | 0 | 0 | 0 | 0 | 0 | 0 | 0 | 0 | 0 | 0 | 0 | 0 | 0 | 0 | 0 | 0 | 0 | 0 | 0 | 0 | 0 |
| Amino acid variations | 2 | 1 | 3 | 3 | 4 | 3 | 3 | 2 | 1 | 1 | 3 | 1 | 1 | 2 | 4 | 4 | 3 | 5 | 2 | 4 | 4 | 5 | 8 | 11 | 9 | 7 | 13 | 12 | 9 | 6 |
| Most frequent amino acid | R | E | E | E | G | I | Q | L | R | K | Q | K | R | E | E | Q | L | F | K | R | R | N | V | S | L | P | R | N | D | E |
| Conservation (%) | 98.7 | 100 | 98.0 | 96.7 | 98.0 | 96.7 | 98.7 | 98.7 | 100 | 100 | 98.7 | 100 | 100 | 99.3 | 97.3 | 97.3 | 92.7 | 92.7 | 96.7 | 92.0 | 93.3 | 92.0 | 90.7 | 78.0 | 73.3 | 84.0 | 48.0 | 68.7 | 68.0 | 83.3 |

| KPNA6 IBB domain | 7 | 15 | 16 | 21 | 22 | 23 | 24 | 25 | 26 | 29 | 30 | 32 | 33 | 34 | 35 | 36 | 37 | 38 | 39 | 40 | 41 | 42 | 43 | 44 | 45 | 46 | 47 | 48 | 49 | 50 |
|---|---|---|---|---|---|---|---|---|---|---|---|---|---|---|---|---|---|---|---|---|---|---|---|---|---|---|---|---|---|---|
| Ala | 1 | 1 | 3 | 0 | 210 | 1 | 0 | 87 | 0 | 1 | 0 | 1 | 0 | 0 | 1 | 0 | 0 | 0 | 0 | 2 | 239 | 0 | 0 | 11 | 2 | 0 | 0 | 0 | 0 | 0 |
| Arg | 1 | 0 | 0 | 0 | 0 | 0 | 0 | 0 | 1 | 0 | 0 | 0 | 246 | 0 | 0 | 62 | 0 | 0 | 0 | 0 | 0 | 0 | 0 | 0 | 0 | 0 | 0 | 249 | 249 | 248 |
| Asn | 2 | 0 | 1 | 2 | 0 | 2 | 0 | 0 | 0 | 0 | 237 | 0 | 0 | 0 | 64 | 0 | 0 | 0 | 249 | 127 | 0 | 0 | 237 | 0 | 0 | 0 | 0 | 0 | 0 | 0 |
| Asp | 2 | 8 | 0 | 0 | 0 | 1 | 0 | 0 | 0 | 216 | 0 | 0 | 0 | 0 | 0 | 0 | 0 | 0 | 0 | 0 | 0 | 0 | 11 | 0 | 9 | 0 | 0 | 0 | 0 | 0 |
| Cys | 0 | 0 | 0 | 0 | 0 | 1 | 0 | 0 | 0 | 0 | 0 | 0 | 0 | 0 | 0 | 4 | 0 | 0 | 0 | 0 | 0 | 0 | 0 | 0 | 0 | 0 | 0 | 0 | 0 | 0 |
| Gln | 1 | 2 | 3 | 0 | 0 | 0 | 0 | 1 | 0 | 0 | 0 | 0 | 0 | 0 | 1 | 0 | 0 | 0 | 0 | 1 | 0 | 0 | 0 | 1 | 3 | 0 | 0 | 0 | 0 | 0 |
| Glu | 0 | 135 | 1 | 0 | 0 | 0 | 0 | 0 | 0 | 22 | 0 | 0 | 0 | 0 | 0 | 0 | 0 | 0 | 0 | 3 | 0 | 0 | 0 | 0 | 230 | 249 | 0 | 0 | 0 | 0 |
| Gly | 1 | 4 | 1 | 0 | 1 | 2 | 1 | 144 | 0 | 0 | 0 | 0 | 0 | 0 | 0 | 0 | 0 | 0 | 0 | 0 | 7 | 0 | 0 | 0 | 0 | 0 | 0 | 0 | 0 | 0 |
| His | 0 | 3 | 0 | 0 | 0 | 0 | 0 | 0 | 0 | 0 | 0 | 9 | 0 | 0 | 4 | 0 | 0 | 0 | 0 | 0 | 0 | 0 | 0 | 0 | 0 | 0 | 0 | 0 | 0 | 0 |
| Ile | 6 | 0 | 0 | 2 | 0 | 1 | 0 | 0 | 0 | 0 | 1 | 0 | 0 | 1 | 0 | 1 | 0 | 0 | 0 | 0 | 0 | 0 | 0 | 0 | 0 | 0 | 0 | 0 | 0 | 0 |
| Leu | 36 | 4 | 0 | 1 | 1 | 0 | 0 | 0 | 0 | 0 | 0 | 0 | 0 | 1 | 0 | 9 | 0 | 0 | 0 | 0 | 0 | 245 | 0 | 8 | 0 | 0 | 9 | 0 | 0 | 0 |
| Lys | 2 | 3 | 0 | 0 | 0 | 0 | 0 | 0 | 231 | 0 | 0 | 0 | 3 | 0 | 177 | 1 | 0 | 249 | 0 | 112 | 0 | 1 | 0 | 0 | 3 | 0 | 0 | 0 | 0 | 1 |
| Met | 100 | 2 | 0 | 186 | 0 | 0 | 0 | 0 | 0 | 0 | 0 | 0 | 0 | 238 | 2 | 0 | 0 | 0 | 0 | 0 | 0 | 0 | 0 | 0 | 0 | 0 | 240 | 0 | 0 | 0 |
| Phe | 2 | 0 | 0 | 0 | 0 | 0 | 0 | 0 | 0 | 0 | 0 | 1 | 0 | 0 | 0 | 0 | 3 | 0 | 0 | 0 | 0 | 1 | 0 | 0 | 0 | 0 | 0 | 0 | 0 | 0 |
| Pro | 2 | 43 | 1 | 0 | 0 | 0 | 231 | 0 | 0 | 0 | 0 | 0 | 0 | 0 | 0 | 0 | 0 | 0 | 0 | 0 | 2 | 0 | 0 | 221 | 0 | 0 | 0 | 0 | 0 | 0 |
| Ser | 4 | 5 | 44 | 3 | 15 | 229 | 0 | 0 | 0 | 0 | 0 | 5 | 0 | 0 | 0 | 172 | 0 | 0 | 0 | 0 | 1 | 0 | 1 | 7 | 0 | 0 | 0 | 0 | 0 | 0 |
| Thr | 2 | 5 | 156 | 0 | 10 | 3 | 4 | 0 | 0 | 0 | 0 | 0 | 0 | 0 | 0 | 0 | 0 | 0 | 0 | 3 | 0 | 0 | 0 | 1 | 2 | 0 | 0 | 0 | 0 | 0 |
| Trp | 0 | 0 | 0 | 0 | 0 | 0 | 0 | 0 | 0 | 0 | 0 | 0 | 0 | 0 | 0 | 0 | 0 | 0 | 0 | 0 | 0 | 0 | 0 | 0 | 0 | 0 | 0 | 0 | 0 | 0 |
| Tyr | 4 | 1 | 2 | 0 | 1 | 0 | 0 | 0 | 1 | 0 | 0 | 231 | 0 | 0 | 0 | 0 | 246 | 0 | 0 | 0 | 0 | 0 | 0 | 0 | 0 | 0 | 0 | 0 | 0 | 0 |
| Val | 23 | 2 | 0 | 1 | 0 | 0 | 0 | 0 | 0 | 0 | 0 | 0 | 0 | 0 | 0 | 0 | 0 | 0 | 0 | 1 | 0 | 2 | 0 | 0 | 0 | 0 | 0 | 0 | 0 | 0 |
| - | 60 | 31 | 37 | 54 | 11 | 9 | 13 | 17 | 16 | 10 | 11 | 2 | 0 | 9 | 0 | 0 | 0 | 0 | 0 | 0 | 0 | 0 | 0 | 0 | 0 | 0 | 0 | 0 | 0 | 0 |
| X | 0 | 0 | 0 | 0 | 0 | 0 | 0 | 0 | 0 | 0 | 0 | 0 | 0 | 0 | 0 | 0 | 0 | 0 | 0 | 0 | 0 | 0 | 0 | 0 | 0 | 0 | 0 | 0 | 0 | 0 |
| Amino acid variations | 16 | 14 | 9 | 6 | 6 | 8 | 3 | 3 | 3 | 3 | 2 | 5 | 2 | 3 | 6 | 6 | 2 | 1 | 1 | 7 | 4 | 4 | 3 | 6 | 6 | 1 | 2 | 1 | 1 | 2 |
| Most frequent amino acid | M | E | T | M | A | S | P | G | K | D | N | Y | R | M | K | S | Y | K | N | N | A | L | N | P | E | E | M | R | R | R |
| Conservation (%) | 40.2 | 54.2 | 62.7 | 74.7 | 84.3 | 92.0 | 92.8 | 57.8 | 92.8 | 86.7 | 95.2 | 92.8 | 98.8 | 95.6 | 71.1 | 69.1 | 98.8 | 100 | 100 | 51.0 | 96.0 | 98.4 | 95.2 | 88.8 | 92.4 | 100 | 96.4 | 100 | 100 | 99.6 |

| KPNA6 IBB domain | 51 | 52 | 53 | 54 | 55 | 56 | 57 | 58 | 59 | 60 | 61 | 62 | 63 | 64 | 65 | 66 | 70 | 71 | 72 | 73 | 74 | 75 | 76 | 77 | 80 | 81 | 82 | 83 | 84 | 85 |
|---|---|---|---|---|---|---|---|---|---|---|---|---|---|---|---|---|---|---|---|---|---|---|---|---|---|---|---|---|---|---|
| Ala | 0 | 0 | 0 | 0 | 0 | 0 | 0 | 0 | 0 | 0 | 1 | 0 | 0 | 0 | 0 | 0 | 0 | 0 | 0 | 0 | 0 | 0 | 0 | 0 | 3 | 5 | 0 | 1 | 1 | 168 |
| Arg | 249 | 0 | 0 | 0 | 0 | 0 | 0 | 0 | 249 | 0 | 0 | 0 | 249 | 0 | 1 | 0 | 0 | 0 | 0 | 249 | 245 | 0 | 0 | 0 | 0 | 0 | 0 | 0 | 1 | 2 |
| Asn | 0 | 0 | 0 | 0 | 0 | 0 | 0 | 0 | 0 | 0 | 0 | 0 | 0 | 0 | 0 | 0 | 0 | 1 | 0 | 0 | 0 | 244 | 0 | 4 | 0 | 1 | 220 | 0 | 3 | 0 |
| Asp | 0 | 0 | 0 | 1 | 0 | 0 | 0 | 0 | 0 | 0 | 0 | 0 | 0 | 6 | 1 | 0 | 0 | 0 | 0 | 0 | 0 | 0 | 0 | 57 | 0 | 0 | 1 | 4 | 49 | 9 |
| Cys | 0 | 0 | 0 | 0 | 0 | 0 | 0 | 0 | 0 | 0 | 0 | 0 | 0 | 0 | 0 | 0 | 0 | 0 | 0 | 0 | 0 | 0 | 2 | 0 | 0 | 0 | 0 | 0 | 0 | 0 |
| Gln | 0 | 0 | 0 | 0 | 0 | 0 | 249 | 0 | 0 | 0 | 246 | 0 | 0 | 0 | 240 | 248 | 0 | 0 | 0 | 0 | 0 | 0 | 0 | 0 | 0 | 0 | 4 | 2 | 2 | 0 |
| Glu | 0 | 249 | 249 | 247 | 0 | 0 | 0 | 0 | 0 | 0 | 1 | 0 | 0 | 243 | 4 | 0 | 0 | 0 | 0 | 0 | 2 | 0 | 0 | 180 | 2 | 0 | 1 | 230 | 188 | 48 |
| Gly | 0 | 0 | 0 | 0 | 248 | 0 | 0 | 0 | 0 | 0 | 0 | 0 | 0 | 0 | 0 | 0 | 0 | 0 | 0 | 0 | 0 | 0 | 0 | 1 | 1 | 0 | 3 | 0 | 0 | 2 |
| His | 0 | 0 | 0 | 0 | 0 | 0 | 0 | 0 | 0 | 0 | 0 | 0 | 0 | 0 | 0 | 0 | 0 | 0 | 0 | 0 | 0 | 0 | 0 | 0 | 0 | 0 | 1 | 0 | 0 | 0 |
| Ile | 0 | 0 | 0 | 0 | 0 | 240 | 0 | 2 | 0 | 0 | 0 | 0 | 0 | 0 | 2 | 0 | 4 | 2 | 0 | 0 | 0 | 0 | 1 | 0 | 36 | 172 | 0 | 0 | 0 | 1 |
| Leu | 0 | 0 | 0 | 0 | 0 | 0 | 0 | 247 | 0 | 0 | 0 | 0 | 0 | 0 | 1 | 0 | 243 | 2 | 0 | 0 | 0 | 2 | 2 | 0 | 168 | 65 | 0 | 0 | 0 | 0 |
| Lys | 0 | 0 | 0 | 0 | 0 | 0 | 0 | 0 | 0 | 249 | 0 | 249 | 0 | 0 | 0 | 0 | 0 | 0 | 245 | 0 | 0 | 0 | 0 | 0 | 0 | 0 | 0 | 1 | 0 | 0 |
| Met | 0 | 0 | 0 | 0 | 0 | 0 | 0 | 0 | 0 | 0 | 0 | 0 | 0 | 0 | 0 | 0 | 1 | 0 | 0 | 0 | 0 | 2 | 0 | 0 | 10 | 0 | 0 | 0 | 0 | 0 |
| Phe | 0 | 0 | 0 | 0 | 0 | 0 | 0 | 0 | 0 | 0 | 0 | 0 | 0 | 0 | 0 | 0 | 0 | 239 | 0 | 0 | 0 | 0 | 0 | 0 | 3 | 0 | 2 | 0 | 0 | 0 |
| Pro | 0 | 0 | 0 | 0 | 0 | 0 | 0 | 0 | 0 | 0 | 0 | 0 | 0 | 0 | 0 | 0 | 0 | 0 | 0 | 0 | 0 | 0 | 0 | 1 | 1 | 1 | 5 | 2 | 2 | 3 |
| Ser | 0 | 0 | 0 | 0 | 1 | 0 | 0 | 0 | 0 | 0 | 0 | 0 | 0 | 0 | 0 | 0 | 0 | 3 | 2 | 0 | 0 | 0 | 0 | 2 | 0 | 0 | 8 | 2 | 1 | 8 |
| Thr | 0 | 0 | 0 | 0 | 0 | 0 | 0 | 0 | 0 | 0 | 1 | 0 | 0 | 0 | 0 | 0 | 0 | 0 | 2 | 0 | 0 | 1 | 0 | 1 | 2 | 1 | 4 | 1 | 0 | 4 |
| Trp | 0 | 0 | 0 | 0 | 0 | 0 | 0 | 0 | 0 | 0 | 0 | 0 | 0 | 0 | 0 | 0 | 0 | 0 | 0 | 0 | 0 | 0 | 0 | 0 | 0 | 0 | 0 | 0 | 0 | 0 |
| Tyr | 0 | 0 | 0 | 0 | 0 | 0 | 0 | 0 | 0 | 0 | 0 | 0 | 0 | 0 | 0 | 0 | 0 | 0 | 0 | 0 | 0 | 0 | 0 | 0 | 0 | 0 | 0 | 0 | 0 | 0 |
| Val | 0 | 0 | 0 | 1 | 0 | 9 | 0 | 0 | 0 | 0 | 0 | 0 | 0 | 0 | 0 | 1 | 1 | 0 | 0 | 0 | 0 | 0 | 244 | 2 | 21 | 4 | 0 | 0 | 1 | 1 |
| - | 0 | 0 | 0 | 0 | 0 | 0 | 0 | 0 | 0 | 0 | 0 | 0 | 0 | 0 | 0 | 0 | 0 | 2 | 0 | 0 | 2 | 0 | 0 | 1 | 2 | 0 | 0 | 6 | 1 | 3 |
| X | 0 | 0 | 0 | 0 | 0 | 0 | 0 | 0 | 0 | 0 | 0 | 0 | 0 | 0 | 0 | 0 | 0 | 0 | 0 | 0 | 0 | 0 | 0 | 0 | 0 | 0 | 0 | 0 | 0 | 0 |
| Amino acid variations | 1 | 1 | 1 | 3 | 2 | 2 | 1 | 2 | 1 | 1 | 4 | 1 | 1 | 2 | 6 | 2 | 4 | 5 | 3 | 1 | 2 | 4 | 4 | 8 | 10 | 7 | 10 | 8 | 9 | 10 |
| Most frequent amino acid | R | E | E | E | G | I | Q | L | R | K | Q | K | R | E | Q | Q | L | F | K | R | R | N | V | E | L | I | N | E | E | A |
| Conservation (%) | 100 | 100 | 100 | 99.2 | 99.6 | 96.4 | 100 | 99.2 | 100 | 100 | 98.8 | 100 | 100 | 97.6 | 96.4 | 99.6 | 97.6 | 96.0 | 98.4 | 100 | 98.4 | 98.0 | 98.0 | 72.3 | 67.5 | 69.1 | 88.4 | 92.4 | 75.5 | 67.5 |

| KPNA7 IBB domain | 9 | 10 | 11 | 12 | 13 | 14 | 17 | 18 | 20 | 23 | 24 | 25 | 26 | 28 | 30 | 35 | 36 | 37 | 38 | 39 | 40 | 41 | 42 | 43 | 44 | 46 | 47 | 48 | 49 | 50 |
|---|---|---|---|---|---|---|---|---|---|---|---|---|---|---|---|---|---|---|---|---|---|---|---|---|---|---|---|---|---|---|
| Ala | 0 | 6 | 4 | 4 | 2 | 57 | 4 | 1 | 2 | 0 | 0 | 0 | 0 | 0 | 0 | 0 | 0 | 0 | 0 | 0 | 58 | 22 | 21 | 0 | 0 | 0 | 0 | 0 | 1 | 58 |
| Arg | 0 | 0 | 0 | 0 | 3 | 0 | 0 | 0 | 0 | 90 | 12 | 66 | 5 | 0 | 0 | 0 | 55 | 0 | 3 | 0 | 0 | 0 | 0 | 57 | 95 | 29 | 13 | 93 | 0 | 0 |
| Asn | 0 | 0 | 1 | 0 | 13 | 0 | 0 | 2 | 0 | 0 | 0 | 3 | 3 | 0 | 0 | 33 | 2 | 0 | 0 | 8 | 0 | 0 | 0 | 0 | 0 | 0 | 0 | 0 | 0 | 0 |
| Asp | 0 | 0 | 0 | 0 | 40 | 0 | 2 | 13 | 11 | 0 | 0 | 0 | 0 | 0 | 0 | 0 | 0 | 0 | 0 | 80 | 0 | 0 | 2 | 0 | 0 | 1 | 0 | 0 | 0 | 1 |
| Cys | 0 | 0 | 1 | 1 | 0 | 0 | 0 | 1 | 0 | 0 | 0 | 0 | 0 | 0 | 0 | 0 | 8 | 0 | 0 | 0 | 0 | 0 | 0 | 0 | 0 | 0 | 0 | 0 | 0 | 0 |
| Gln | 0 | 0 | 0 | 1 | 0 | 0 | 11 | 0 | 3 | 0 | 0 | 0 | 6 | 0 | 2 | 0 | 0 | 0 | 0 | 0 | 1 | 1 | 0 | 0 | 0 | 62 | 80 | 0 | 0 | 0 |
| Glu | 0 | 1 | 0 | 1 | 16 | 19 | 1 | 60 | 71 | 0 | 0 | 0 | 0 | 0 | 0 | 0 | 0 | 0 | 0 | 7 | 2 | 1 | 10 | 0 | 0 | 2 | 1 | 0 | 0 | 31 |
| Gly | 0 | 0 | 2 | 0 | 4 | 0 | 0 | 5 | 5 | 0 | 0 | 0 | 0 | 0 | 0 | 1 | 0 | 95 | 0 | 0 | 0 | 1 | 0 | 0 | 0 | 0 | 0 | 0 | 0 | 0 |
| His | 0 | 0 | 0 | 0 | 5 | 0 | 10 | 0 | 0 | 4 | 0 | 0 | 0 | 0 | 0 | 0 | 0 | 0 | 0 | 1 | 0 | 0 | 0 | 0 | 0 | 1 | 1 | 0 | 0 | 0 |
| Ile | 0 | 0 | 5 | 0 | 1 | 1 | 0 | 0 | 0 | 0 | 4 | 0 | 1 | 1 | 0 | 0 | 0 | 0 | 0 | 0 | 2 | 0 | 1 | 5 | 1 | 0 | 0 | 0 | 35 | 1 |
| Leu | 0 | 1 | 3 | 50 | 0 | 2 | 1 | 0 | 0 | 0 | 49 | 0 | 3 | 1 | 0 | 0 | 0 | 0 | 0 | 0 | 4 | 0 | 19 | 26 | 0 | 0 | 0 | 0 | 2 | 0 |
| Lys | 0 | 0 | 0 | 1 | 6 | 8 | 0 | 4 | 1 | 1 | 0 | 24 | 77 | 0 | 94 | 0 | 30 | 0 | 91 | 0 | 0 | 0 | 3 | 0 | 0 | 0 | 1 | 3 | 0 | 0 |
| Met | 95 | 0 | 0 | 2 | 0 | 1 | 0 | 0 | 0 | 0 | 30 | 0 | 1 | 0 | 0 | 0 | 0 | 0 | 0 | 0 | 5 | 0 | 24 | 5 | 0 | 1 | 0 | 0 | 33 | 0 |
| Phe | 0 | 0 | 0 | 1 | 0 | 0 | 0 | 0 | 0 | 0 | 0 | 0 | 0 | 91 | 0 | 0 | 0 | 0 | 0 | 0 | 1 | 0 | 0 | 0 | 0 | 0 | 0 | 0 | 1 | 0 |
| Pro | 0 | 77 | 0 | 1 | 0 | 0 | 60 | 0 | 0 | 0 | 0 | 0 | 0 | 0 | 0 | 0 | 0 | 0 | 0 | 0 | 4 | 0 | 0 | 3 | 0 | 0 | 0 | 0 | 0 | 0 |
| Ser | 0 | 8 | 3 | 9 | 2 | 0 | 5 | 8 | 0 | 1 | 0 | 2 | 0 | 0 | 0 | 0 | 0 | 1 | 0 | 0 | 0 | 61 | 1 | 0 | 0 | 0 | 0 | 0 | 0 | 1 |
| Thr | 0 | 0 | 63 | 9 | 1 | 1 | 2 | 0 | 0 | 0 | 0 | 1 | 0 | 0 | 0 | 0 | 0 | 0 | 0 | 0 | 12 | 8 | 4 | 0 | 0 | 0 | 0 | 0 | 0 | 1 |
| Trp | 0 | 0 | 0 | 0 | 2 | 0 | 0 | 0 | 1 | 0 | 0 | 0 | 0 | 0 | 0 | 0 | 0 | 0 | 1 | 0 | 0 | 0 | 0 | 0 | 0 | 0 | 0 | 0 | 0 | 0 |
| Tyr | 0 | 0 | 0 | 0 | 0 | 1 | 0 | 0 | 0 | 0 | 0 | 0 | 0 | 3 | 0 | 62 | 0 | 0 | 0 | 0 | 0 | 0 | 0 | 0 | 0 | 0 | 0 | 0 | 0 | 0 |
| Val | 0 | 0 | 14 | 9 | 0 | 5 | 0 | 1 | 0 | 0 | 1 | 0 | 0 | 0 | 0 | 0 | 0 | 0 | 1 | 0 | 5 | 1 | 9 | 0 | 0 | 0 | 0 | 0 | 23 | 3 |
| - | 1 | 3 | 0 | 7 | 1 | 1 | 0 | 1 | 2 | 0 | 0 | 0 | 0 | 0 | 0 | 0 | 0 | 0 | 0 | 0 | 2 | 1 | 2 | 0 | 0 | 0 | 0 | 0 | 1 | 0 |
| X | 0 | 0 | 0 | 0 | 0 | 0 | 0 | 0 | 0 | 0 | 0 | 0 | 0 | 0 | 0 | 0 | 1 | 0 | 0 | 0 | 0 | 0 | 0 | 0 | 0 | 0 | 0 | 0 | 0 | 0 |
| Amino acid variations | 1 | 5 | 9 | 12 | 12 | 9 | 9 | 9 | 7 | 4 | 5 | 5 | 7 | 4 | 2 | 3 | 4 | 2 | 4 | 4 | 10 | 7 | 10 | 5 | 2 | 6 | 5 | 2 | 6 | 7 |
| Most frequent amino acid | M | P | T | L | D | A | P | E | E | R | L | R | K | F | K | Y | R | G | K | D | A | S | M | R | R | Q | Q | R | I | A |
| Conservation (%) | 99.0 | 80.2 | 65.6 | 52.1 | 41.7 | 59.4 | 62.5 | 62.5 | 74.0 | 93.8 | 51.0 | 68.8 | 80.2 | 94.8 | 97.9 | 64.6 | 57.9 | 99.0 | 94.8 | 83.3 | 60.4 | 63.5 | 25.0 | 59.4 | 99.0 | 64.6 | 83.3 | 96.9 | 36.5 | 60.4 |

| KPNA7 IBB domain | 51 | 52 | 53 | 54 | 55 | 56 | 57 | 58 | 59 | 60 | 61 | 62 | 63 | 64 | 66 | 67 | 68 | 69 | 70 | 72 | 73 | 75 | 76 | 77 | 79 | 81 | 82 |
|---|---|---|---|---|---|---|---|---|---|---|---|---|---|---|---|---|---|---|---|---|---|---|---|---|---|---|---|
| Ala | 2 | 0 | 0 | 0 | 0 | 0 | 0 | 91 | 1 | 0 | 0 | 0 | 0 | 34 | 0 | 0 | 0 | 0 | 0 | 0 | 1 | 1 | 1 | 0 | 5 | 0 | 5 |
| Arg | 0 | 3 | 0 | 0 | 0 | 96 | 0 | 0 | 14 | 1 | 0 | 0 | 0 | 0 | 0 | 0 | 93 | 95 | 0 | 0 | 0 | 0 | 1 | 1 | 1 | 0 | 1 |
| Asn | 0 | 20 | 0 | 0 | 0 | 0 | 0 | 0 | 1 | 0 | 1 | 0 | 2 | 0 | 0 | 0 | 0 | 0 | 87 | 0 | 1 | 18 | 2 | 2 | 1 | 1 | 1 |
| Asp | 0 | 0 | 0 | 0 | 0 | 0 | 0 | 0 | 0 | 0 | 87 | 1 | 0 | 0 | 0 | 0 | 0 | 1 | 1 | 0 | 2 | 16 | 0 | 0 | 1 | 65 | 3 |
| Cys | 0 | 3 | 0 | 0 | 0 | 0 | 0 | 0 | 0 | 0 | 0 | 0 | 0 | 0 | 0 | 0 | 0 | 0 | 0 | 0 | 7 | 1 | 3 | 14 | 0 | 0 | 0 |
| Gln | 0 | 0 | 1 | 9 | 0 | 0 | 0 | 0 | 2 | 0 | 0 | 0 | 93 | 0 | 0 | 0 | 0 | 0 | 0 | 0 | 1 | 3 | 1 | 0 | 0 | 0 | 0 |
| Glu | 2 | 0 | 0 | 84 | 0 | 0 | 0 | 0 | 0 | 0 | 6 | 95 | 0 | 0 | 0 | 0 | 0 | 0 | 0 | 0 | 1 | 0 | 0 | 1 | 7 | 20 | 14 |
| Gly | 0 | 1 | 0 | 0 | 0 | 0 | 0 | 0 | 0 | 0 | 0 | 0 | 1 | 0 | 0 | 0 | 0 | 0 | 0 | 0 | 2 | 0 | 0 | 2 | 1 | 2 | 0 |
| His | 0 | 0 | 0 | 0 | 0 | 0 | 0 | 0 | 0 | 0 | 1 | 0 | 0 | 0 | 0 | 0 | 0 | 0 | 0 | 0 | 0 | 1 | 2 | 0 | 2 | 1 | 0 |
| Ile | 1 | 2 | 9 | 0 | 0 | 0 | 0 | 0 | 0 | 0 | 0 | 0 | 0 | 31 | 0 | 0 | 0 | 0 | 0 | 87 | 3 | 9 | 0 | 0 | 0 | 0 | 0 |
| Leu | 1 | 0 | 58 | 0 | 95 | 0 | 0 | 0 | 0 | 0 | 0 | 0 | 0 | 1 | 92 | 0 | 1 | 0 | 0 | 1 | 1 | 5 | 15 | 9 | 16 | 0 | 6 |
| Lys | 0 | 1 | 0 | 2 | 0 | 0 | 96 | 0 | 78 | 95 | 0 | 0 | 0 | 0 | 0 | 95 | 2 | 0 | 0 | 0 | 0 | 0 | 0 | 1 | 2 | 1 | 4 |
| Met | 1 | 0 | 0 | 0 | 0 | 0 | 0 | 0 | 0 | 0 | 0 | 0 | 0 | 1 | 1 | 0 | 0 | 0 | 0 | 0 | 1 | 0 | 0 | 0 | 9 | 1 | 3 |
| Phe | 1 | 0 | 0 | 0 | 1 | 0 | 0 | 0 | 0 | 0 | 0 | 0 | 0 | 2 | 3 | 0 | 0 | 0 | 0 | 0 | 0 | 13 | 29 | 2 | 0 | 0 | 0 |
| Pro | 0 | 0 | 0 | 0 | 0 | 0 | 0 | 0 | 0 | 0 | 0 | 0 | 0 | 0 | 0 | 0 | 0 | 0 | 0 | 0 | 0 | 1 | 4 | 2 | 33 | 1 | 37 |
| Ser | 4 | 66 | 0 | 0 | 0 | 0 | 0 | 1 | 0 | 0 | 0 | 0 | 0 | 6 | 0 | 0 | 0 | 0 | 8 | 0 | 21 | 21 | 11 | 49 | 15 | 2 | 5 |
| Thr | 1 | 0 | 0 | 0 | 0 | 0 | 0 | 3 | 0 | 0 | 0 | 0 | 0 | 15 | 0 | 1 | 0 | 0 | 0 | 0 | 53 | 4 | 3 | 0 | 2 | 0 | 13 |
| Trp | 0 | 0 | 0 | 0 | 0 | 0 | 0 | 0 | 0 | 0 | 0 | 0 | 0 | 0 | 0 | 0 | 0 | 0 | 0 | 0 | 0 | 0 | 0 | 0 | 0 | 0 | 0 |
| Tyr | 0 | 0 | 0 | 0 | 0 | 0 | 0 | 0 | 0 | 0 | 0 | 0 | 0 | 0 | 0 | 0 | 0 | 0 | 0 | 0 | 0 | 1 | 0 | 0 | 0 | 2 | 0 |
| Val | 83 | 0 | 28 | 1 | 0 | 0 | 0 | 1 | 0 | 0 | 1 | 0 | 0 | 6 | 0 | 0 | 0 | 0 | 0 | 8 | 1 | 1 | 23 | 1 | 0 | 0 | 1 |
| - | 0 | 0 | 0 | 0 | 0 | 0 | 0 | 0 | 0 | 0 | 0 | 0 | 0 | 0 | 0 | 0 | 0 | 0 | 0 | 0 | 1 | 1 | 1 | 12 | 1 | 0 | 3 |
| X | 0 | 0 | 0 | 0 | 0 | 0 | 0 | 0 | 0 | 0 | 0 | 0 | 0 | 0 | 0 | 0 | 0 | 0 | 0 | 0 | 0 | 0 | 0 | 0 | 0 | 0 | 0 |
| Amino acid variations | 9 | 7 | 4 | 4 | 2 | 1 | 1 | 4 | 5 | 2 | 5 | 2 | 3 | 8 | 3 | 2 | 3 | 2 | 3 | 3 | 13 | 14 | 12 | 11 | 13 | 10 | 12 |
| Most frequent amino acid | V | S | L | E | L | R | K | A | K | K | D | E | Q | A | L | K | R | R | N | I | T | S | F | S | P | D | P |
| Conservation (%) | 86.5 | 68.8 | 60.4 | 87.5 | 99.0 | 100 | 100 | 94.8 | 81.3 | 99.0 | 90.6 | 99.0 | 96.9 | 35.4 | 95.8 | 99.0 | 96.9 | 99.0 | 90.6 | 90.6 | 55.2 | 21.9 | 30.2 | 51.0 | 34.4 | 67.7 | 38.5 |
