## Supporting file 5 for "Biochemical propensity mapping for structural and functional anatomy of importin α IBB domain"

```
                                 *        20         *        40         *
cIBB1                 : MTTPGKENFRLKSYKNKSLNPDEMRRRREEEGLQLRKQKREEQLFKRRNVATAEEET
Homo_sapiens KPNA2    : ---------RLHRFKNKGKDSTEMRRRRIEVNVELRKAKKDDQMLKRRNVS------


                                 *        20         *        40         *        60
cIBB5                 : MDAMASPGKDNYRMKSYKNKALNPQEMRRRREEEGIQLRKQKREEQLFKRRNVSLPRNDE
Homo_sapiens KPNA2    : ------------RLHRFKNKGKDSTEMRRRRIEVNVELRKAKKDDQMLKRRNVS------


                                 *        20         *        40         *        60
cIBB6                 : METMASPGKDNYRMKSYKNNALNPEEMRRRREEEGIQLRKQKREQQLFKRRNVELINEEA
Homo_sapiens KPNA2    : -----------ARLHRFKNKGKDSTEMRRRRIEVNVELRKAKKDDQMLKRRNVS------


                                 *        20         *        40         *
cIBB3                 : MAENPGLENHRIKSFKNKGRDVETMRRHRNEVTVELRKNKRDEHLLKKRNVPQEESLE
Homo_sapiens KPNA2    : ----------RLHRFKNKGKDSTEMRRRRIEVNVELRKAKKDDQMLKRRNVS------


                                *        20         *        40         *
cIBB4                 : MADNEKLDNQRLKNFKNKGRDLETMRRQRNEVVVELRKNKRDEHLLKRRNVPHEDICE
Homo_sapiens KPNA2    : ---------ARLHRFKNKGKDSTEMRRRRIEVNVELRKAKKDDQMLKRRNVS------


                                *        20         *        40         *
cIBB2                 : MSTNENANPARLNRFKNKGKDSTEMRRRRIEVNVELRKAKKDDQMLKRRNVSSFPDDA
Homo_sapiens KPNA2    : --------AARLHRFKNKGKDSTEMRRRRIEVNVELRKAKKDDQMLKRRNVS------


                                *        20         *        40         *
cIBB7                 : MPTLDAPEERLRKFKYRGKDASMRRQQRIAVSLELRKAKKDEQALKRRNITSFSPDP
Homo_sapiens KPNA2    : --------ARLHRFKNKGKDSTEMRRRRIEVNVELRKAKKDDQMLKRRNVS------
```

```
                             *        20         *        40         *        60         *        80
chimeric KPNA1    : MTTPGKENFRLKSYKNKSLNPDEMRRRREEEGLQLRKQKREEQLFKRRNVATAEEETEEEVMSDGGFHEAQISNMEMAPG
S.cerevisiae SRP1 : -----------------YRRTNELRRRRDTQQVELRKAKRDEALAKRRNFQE----------------------------

                             *       100         *       120         *       140         *       160
chimeric KPNA1    : GVITSDMIEMIFSKSPEQQLSATQKFRKLLSKEPNPPIDEVISTPGVVARFVEFLKRKENCTLQFESAWVLTNIASGNSL
S.cerevisiae SRP1 : ---LPQMTQQLNSDDMQEQLSATVKFRQILSQR--PPIDVVIQ-AGVVPRLVEFMRENQPEMLQLEAAWALTNIASGTSA

                             *       180         *       200         *       220         *       240
chimeric KPNA1    : QTRIVIQAGAVPIFIELLSSEFEDVQEQAVWALGNIAGDSTMCRDYVLDCNILPPLLQLFSKQNRLTMTRNAVWALSNLC
S.cerevisiae SRP1 : QTKVVVDADAVPLFIQLLYTGSVEVKEQAIWALGNVAGDSTDYRDYVLQCNAMEPILGLFNSN-KPSLIRTATWTLSNLC

                             *       260         *       280         *       300         *       320
chimeric KPNA1    : RGKSPPPEFAKVSPCLNVLSWLLFVSDTDVLADACWALSYLSDGPNDKIQAVIDAGVCRRLVELLMHNDYKVVSPALRAV
S.cerevisiae SRP1 : RGKKPQPDWSVVSQALPTLAKLIYSMDTETLVDACWAISYLSDGPQEAIQAVIDVRIPKRLVELLSHESTLVQTPALRAV

                             *       340         *       360         *       380         *       400
chimeric KPNA1    : GNIVTGDDIQTQVILNCSALQSLLHLLSSPKESIKKEACWTISNITAGNRAQIQTVIDANIFPALISILQTAEFRTRKEA
S.cerevisiae SRP1 : GNIVTGNDLQTQVVINAGVLPALRLLLSSPKENIKKEACWTISNITAGNTEQIQAVIDANLIPPLVKLLEVAEYKTKKEA

                             *       420         *       440         *       460         *       480
chimeric KPNA1    : AWAITNATSGG--SAEQIKYLVELGCIKPLCDLLTVMDSKIVQVALNGLENILRLGEQEAKRNGTGINPYCALIEEAYGL
S.cerevisiae SRP1 : CWAISNASSGGLQRPDIIRYLVSQGCIKPLCDLLEIADNRIIEVTLDALENILKMGEADKEARGLNINENADFIEKAGGM

                             *       500         *       520         *       540
chimeric KPNA1    : DKIEFLQSHENQEIYQKAFDLIEHYFGTEDEDSSIAPQVDLNQQQYIFQQCEAPMEGFQL
S.cerevisiae SRP1 : EKIFNCQQNENDKIYEKAYKIIETYFGEEED-----------------------------
```

```
                         *        20         *        40         *        60         *        80
chimeric KPNA5     : MDAMASPGKDNYRMKSYKNKALNPQEMRRRREEEGIQLRKQKREEQLFKRRNVSLPRNDESMLESPIQDPDISSTVPIPE
S.cerevisiae SRP1 : --------------------YRRTNELRRRRDTQQVELRKAKRDEALAKRRNFQ--------------------------

                         *       100         *       120         *       140         *       160
chimeric KPNA5     : EEVVTTDMVQMIFSNNADQQLTATQKFRKLLSKEPNPPIDQVIQKPGVVQRFVKFLERNENCTLQFEAAWALTNIASGTF
S.cerevisiae SRP1 : ---ELPQMTQQLNSDDMQEQLSATVKFRQILSQR--PPIDVVI-QAGVVPRLVEFMRENQPEMLQLEAAWALTNIASGTS

                         *       180         *       200         *       220         *       240
chimeric KPNA5     : LHTKVVIETGAVPIFIKLLNSEHEDVQEQAVWALGNIAGDNAECRDFVLNCEILPPLLELLTNSNRLTTTRNAVWALSNL
S.cerevisiae SRP1 : AQTKVVVDADAVPLFIQLLYTGSVEVKEQAIWALGNVAGDSTDYRDYVLQCNAMEPILGLFNSN-KPSLIRTATWTLSNL

                         *       260         *       280         *       300         *       320
chimeric KPNA5     : CRGKNPPPNFSKVSPCLNVLSRLLFSSDPDVLADVCWALSYLSDGPNDKIQAVIDSGVCRRLVELLMHNDYKVVSPALRA
S.cerevisiae SRP1 : CRGKKPQPDWSVVSQALPTLAKLIYSMDTETLVDACWAISYLSDGPQEAIQAVIDVRIPKRLVELLSHESTLVQTPALRA

                         *       340         *       360         *       380         *       400
chimeric KPNA5     : VGNIVTGDDIQTQVILNCSALPCLLHLLSSPKESIRKEACWTVSNITAGNRAQIQAVIDANIFPVLIEILQKAEFRTRKE
S.cerevisiae SRP1 : VGNIVTGNDLQTQVVINAGVLPALRLLLSSPKENIKKEACWTISNITAGNTEQIQAVIDANLIPPLVKLLEVAEYKTKKE

                         *       420         *       440         *       460         *       480
chimeric KPNA5     : AAWAITNATSGG--TPEQIRYLVALGCIKPLCDLLTVMDSKIVQVALNGLENILRLGEQESKQNGIGINPYCALIEEAYG
S.cerevisiae SRP1 : ACWAISNASSGGLQRPDIIRYLVSQGCIKPLCDLLEIADNRIIEVTLDALENILKMGEADKEARGLNINENADFIEKAGG

                         *       500         *       520         *       540
chimeric KPNA5     : LDKIEFLQSHENQEIYQKAFDLIEHYFGVEEDDPSIVPQVDENQQQFIFQQQEAPMDGFQL
S.cerevisiae SRP1 : MEKIFNCQQNENDKIYEKAYKIIETYFGEEED-----------------------------
```

```
                             *        20         *        40         *        60         *        80
chimeric KPNA6    : METMASPGKDNYRMKSYKNNALNPEEMRRRREEEGIQLRKQKREQQLFKRRNVELINEEAAMFDSLLMDSYVSSTTGESV
S.cerevisiae SRP1 : --------------------YRRTNELRRRRDTQQVELRKAKRDEALAKRRNFQ--------------------------

                             *       100         *       120         *       140         *       160
chimeric KPNA6    : ITREMVEMLFSDDSDLQLATTQKFRKLLSKEPSPPIDEVINTPRVVDRFVEFLKRNENCTLQFEAAWALTNIASGTSQQT
S.cerevisiae SRP1 : ELPQMTQQLNSDDMQEQLSATVKFRQILSQR--PPIDVVIQ-AGVVPRLVEFMRENQPEMLQLEAAWALTNIASGTSAQT

                             *       180         *       200         *       220         *       240
chimeric KPNA6    : KIVIEAGAVPIFIELLNSDFEDVQEQAVWALGNIAGDSSVCRDYVLNCSILNPLLTLLTKSTRLTMTRNAVWALSNLCRG
S.cerevisiae SRP1 : KVVVDADAVPLFIQLLYTGSVEVKEQAIWALGNVAGDSTDYRDYVLQCNAMEPILGLFN-SNKPSLIRTATWTLSNLCRG

                             *       260         *       280         *       300         *       320
chimeric KPNA6    : KNPPPEFAKVSPCLPVLSRLLFSSDSDLLADACWALSYLSDGPNEKIQAVIDSGVCRRLVELLMHNDYKVASPALRAVGN
S.cerevisiae SRP1 : KKPQPDWSVVSQALPTLAKLIYSMDTETLVDACWAISYLSDGPQEAIQAVIDVRIPKRLVELLSHESTLVQTPALRAVGN

                             *       340         *       360         *       380         *       400
chimeric KPNA6    : IVTGDDIQTQVILNCSALPCLLHLLSSPKESIRKEACWTISNITAGNRAQIQAVIDANIFPVLIEILQKAEFRTRKEAAW
S.cerevisiae SRP1 : IVTGNDLQTQVVINAGVLPALRLLLSSPKENIKKEACWTISNITAGNTEQIQAVIDANLIPPLVKLLEVAEYKTKKEACW

                             *       420         *       440         *       460         *       480
chimeric KPNA6    : AITNATSGG--TPEQIRYLVSLGCIKPLCDLLTVMDSKIVQVALNGLENILRLGEQEGKRSGSGVNPYCGLIEEAYGLDK
S.cerevisiae SRP1 : AISNASSGGLQRPDIIRYLVSQGCIKPLCDLLEIADNRIIEVTLDALENILKMGEADKEARGLNINENADFIEKAGGMEK

                             *       500         *       520         *
chimeric KPNA6    : IEFLQSHENQEIYQKAFDLIEHYFGVEDDDSSLAPQVDETQQQFIFQQPEAPMEGFQL
S.cerevisiae SRP1 : IFNCQQNENDKIYEKAYKIIETYFGEEED-----------------------------
```

```
                          *        20         *        40         *        60         *        80
chimeric KPNA3     : MAENPGLENHRIKSFKNKGRDVETMRRHRNEVTVELRKNKRDEHLLKKRNVPQEESLEDSDVDADFKAQNVTLEAILQNA
S.cerevisiae SRP1 : -----------------EYRRTNELRRRRDTQQVELRKAKRDEALAKRRNFQ-------------------ELPQMTQQL

                          *       100         *       120         *       140         *       160
chimeric KPNA3     : TSDNPVVQLSAVQAARKLLSSDRNPPIDDLIKSGILPILVKCLERDDNPSLQFEAAWALTNIASGTSAQTQAVVQSNAVP
S.cerevisiae SRP1 : NSDDMQEQLSATVKFRQILSQR--PPIDVVIQAGVVPRLVEFMRENQPEMLQLEAAWALTNIASGTSAQTKVVVDADAVP

                          *       180         *       200         *       220         *       240
chimeric KPNA3     : LFLRLLRSPHQNVCEQAVWALGNIIGDGPQCRDYVISLGVVKPLLSFISPSIPITFLRNVTWVIVNLCRNKDPPPPMETV
S.cerevisiae SRP1 : LFIQLLYTGSVEVKEQAIWALGNVAGDSTDYRDYVLQCNAMEPILGLFNSN-KPSLIRTATWTLSNLCRGKKPQPDWSVV

                          *       260         *       280         *       300         *       320
chimeric KPNA3     : QEILPALCVLIYHTDINILVDTVWALSYLTDGGNEQIQMVIDSGVVPFLVPLLSHQEVKVQTAALRAVGNIVTGTDEQTQ
S.cerevisiae SRP1 : SQALPTLAKLIYSMDTETLVDACWAISYLSDGPQEAIQAVIDVRIPKRLVELLSHESTLVQTPALRAVGNIVTGNDLQTQ

                          *       340         *       360         *       380         *       400
chimeric KPNA3     : VVLNCDVLSHFPNLLSHPKEKINKEAVWFLSNITAGNQQQVQAVIDAGLIPMIIHQLAKGDFGTQKEAAWAISNLTISG-
S.cerevisiae SRP1 : VVINAGVLPALRLLLSSPKENIKKEACWTISNITAGNTEQIQAVIDANLIPPLVKLLEVAEYKTKKEACWAISNASSGGL

                          *       420         *       440         *       460         *       480
chimeric KPNA3     : -RKDQVEYLVQQNVIPPFCNLLSVKDSQVVQVVLDGLKNILIMAGDE-------ASTIAEIIEECGGLEKIEVLQQHENE
S.cerevisiae SRP1 : QRPDIIRYLVSQGCIKPLCDLLEIADNRIIEVTLDALENILKMGEADKEARGLNINENADFIEKAGGMEKIFNCQQNEND

                          *       500         *       520         *
chimeric KPNA3     : DIYKLAFEIIDQYFSGDDIDEDPCLIPEATQGGTYNFDPTANLQTKEFNF
S.cerevisiae SRP1 : KIYEKAYKIIETYFGEEED-------------------------------
```

```
                               *        20         *        40         *        60         *        80
chimeric KPNA4    : ICEDSDIDGDYRVQNTSLEAIVQNASSDNQGIQLSAVQAARKLLSSDRNPPIDDLIKSGILPILVHCLERDDNPSLQFEA
S.cerevisiae SRP1 : ----------------ELPQMTQQLNSDDMQEQLSATVKFRQILSQR--PPIDVVIQAGVVPRLVEFMRENQPEMLQLEA

                               *       100         *       120         *       140         *       160
chimeric KPNA4    : AWALTNIASGTSEQTQAVVQSNAVPLFLRLLHSPHQNVCEQAVWALGNIIGDGPQCRDYVISLGVVKPLLSFISPSIPIT
S.cerevisiae SRP1 : AWALTNIASGTSAQTKVVVDADAVPLFIQLLYTGSVEVKEQAIWALGNVAGDSTDYRDYVLQCNAMEPILGLFNSN-KPS

                               *       180         *       200         *       220         *       240
chimeric KPNA4    : FLRNVTWVMVNLCRHKDPPPPMETIQEILPALCVLIHHTDVNILVDTVWALSYLTDAGNEQIQMVIDSGIVPHLVPLLSH
S.cerevisiae SRP1 : LIRTATWTLSNLCRGKKPQPDWSVVSQALPTLAKLIYSMDTETLVDACWAISYLSDGPQEAIQAVIDVRIPKRLVELLSH

                               *       260         *       280         *       300         *       320
chimeric KPNA4    : QEVKVQTAALRAVGNIVTGTDEQTQVVLNCDALSHFPALLTHPKEKINKEAVWFLSNITAGNQQQVQAVIDANLVPMIIH
S.cerevisiae SRP1 : ESTLVQTPALRAVGNIVTGNDLQTQVVINAGVLPALRLLLSSPKENIKKEACWTISNITAGNTEQIQAVIDANLIPPLVK

                               *       340         *       360         *       380         *       400
chimeric KPNA4    : LLDKGDFGTQKEAAWAISNLTISG--RKDQVAYLIQQNVIPPFCNLLTVKDAQVVQVVLDGLSNILKMAEDEA-------
S.cerevisiae SRP1 : LLEVAEYKTKKEACWAISNASSGGLQRPDIIRYLVSQGCIKPLCDLLEIADNRIIEVTLDALENILKMGEADKEARGLNI

                               *       420         *       440         *       460         *
chimeric KPNA4    : ETIGNLIEECGGLEKIEQLQNHENEDIYKLAYEIIDQFFSSDDIDEDPSLVPEAIQGGTFGFNSSANVPTEGFQF
S.cerevisiae SRP1 : NENADFIEKAGGMEKIFNCQQNENDKIYEKAYKIIETYFGEEE--------------------------------
```

```
                             *        20         *        40         *        60         *        80
chimeric KPNA2     : MSTNENANPARLNRFKNKGKDSTEMRRRRIEVNVELRKAKKDDQMLKRRNVSSFPDDATSPLQENRNNQGTVNWSVDDIV
S.cerevisiae SRP1 : ------------------YRRTNELRRRRDTQQVELRKAKRDEALAKRRNFQ----------------------ELPQMT

                             *       100         *       120         *       140         *       160
chimeric KPNA2     : KGINSSNVENQLQATQAARKLLSREKQPPIDNIIRAGLIPKFVSFLGRTDCSPIQFESAWALTNIASGTSEQTKAVVDGG
S.cerevisiae SRP1 : QQLNSDDMQEQLSATVKFRQILSQR--PPIDVVIQAGVVPRLVEFMRENQPEMLQLEAAWALTNIASGTSAQTKVVVDAD

                             *       180         *       200         *       220         *       240
chimeric KPNA2     : AIPAFISLLASPHAHISEQAVWALGNIAGDGSVFRDLVIKYGAVDPLLALLAVPDMSSLACGYLRNLTWTLSNLCRNKNP
S.cerevisiae SRP1 : AVPLFIQLLYTGSVEVKEQAIWALGNVAGDSTDYRDYVLQCNAMEPILGLFNSNK-----PSLIRTATWTLSNLCRGKKP

                             *       260         *       280         *       300         *       320
chimeric KPNA2     : APPIDAVEQILPTLVRLLHHDDPEVLADTCWAISYLTDGPNERIGMVVKTGVVPQLVKLLGASELPIVTPALRAIGNIVT
S.cerevisiae SRP1 : QPDWSVVSQALPTLAKLIYSMDTETLVDACWAISYLSDGPQEAIQAVIDVRIPKRLVELLSHESTLVQTPALRAVGNIVT

                             *       340         *       360         *       380         *       400
chimeric KPNA2     : GTDEQTQVVIDAGALAVFPSLLTNPKTNIQKEATWTMSNITAGRQDQIQQVVNHGLVPFLVSVLSKADFKTQKEAVWAVT
S.cerevisiae SRP1 : GNDLQTQVVINAGVLPALRLLLSSPKENIKKEACWTISNITAGNTEQIQAVIDANLIPPLVKLLEVAEYKTKKEACWAIS

                             *       420         *       440         *       460         *       480
chimeric KPNA2     : NYTSGG--TVEQIVYLVHCGIIEPLMNLLTAKDTKIILVILDAISNIFQAAEKLG-----ETEKLSIMIEECGGLDKIEA
S.cerevisiae SRP1 : NASSGGLQRPDIIRYLVSQGCIKPLCDLLEIADNRIIEVTLDALENILKMGEADKEARGLNINENADFIEKAGGMEKIFN

                             *       500         *       520         *
chimeric KPNA2     : LQNHENESVYKASLSLIEKYFSVEEEEDQNVVPETTSEGYTFQVQDGAPGTFNF
S.cerevisiae SRP1 : CQQNENDKIYEKAYKIIETYFGEEED----------------------------
```

```
                             *        20         *        40         *        60         *        80
chimeric KPNA7    : MPTLDAPEERLRKFKYRGKDASMRRQQRIAVSLELRKAKKDEQALKRRNITSFSPDPSEKTAKGVAVSLTLGEIIKGVNS
S.cerevisiae SRP1 : ----------------EYRRTNELRRRRDTQQVELRKAKRDEALAKRRNFQE------------------LPQMTQQLNS

                             *       100         *       120         *       140         *       160
chimeric KPNA7    : SDPVLCFQATQTARKMLSQEKNPPLKLVIEAGLIPRMVEFLKSSLYPCLQFEAAWALTNIASGTSEQTRAVVEGGAIQPL
S.cerevisiae SRP1 : DDMQEQLSATVKFRQILSQ--RPPIDVVIQAGVVPRLVEFMRENQPEMLQLEAAWALTNIASGTSAQTKVVVDADAVPLF

                             *       180         *       200         *       220         *       240
chimeric KPNA7    : IELLSSSNVAVCEQAVWALGNIAGDGPEFRDNVITSNAIPHLLALISPTLPITFLRNITWTLSNLCRNKNPYPCDTAVKQ
S.cerevisiae SRP1 : IQLLYTGSVEVKEQAIWALGNVAGDSTDYRDYVLQCNAMEPILGLFNSN-KPSLIRTATWTLSNLCRGKKPQPDWSVVSQ

                             *       260         *       280         *       300         *       320
chimeric KPNA7    : ILPALLHLLQHQDSEVLSDACWALSYLTDGSNKRIGQVVNTGVLPRLVVLMTSSELNVLTPSLRTVGNIVTGTDEQTQMA
S.cerevisiae SRP1 : ALPTLAKLIYSMDTETLVDACWAISYLSDGPQEAIQAVIDVRIPKRLVELLSHESTLVQTPALRAVGNIVTGNDLQTQVV

                             *       340         *       360         *       380         *       400
chimeric KPNA7    : IDAGMLNVLPQLLQHNKPSIQKEAAWALSNVAAGPCHHIQQLLAYDVLPPLVALLKNGEFKVQKEAVWMVANFATGA--T
S.cerevisiae SRP1 : INAGVLPALRLLLSSPKENIKKEACWTISNITAGNTEQIQAVIDANLIPPLVKLLEVAEYKTKKEACWAISNASSGGLQR

                             *       420         *       440         *       460         *       480
chimeric KPNA7    : MDQLIQLVHSGVLEPLVNLLTAPDVKIVLIILDVISCILQAAEKRSE-----KENLCLLIEELGGIDRIEALQLHENRQI
S.cerevisiae SRP1 : PDIIRYLVSQGCIKPLCDLLEIADNRIIEVTLDALENILKMGEADKEARGLNINENADFIEKAGGMEKIFNCQQNENDKI

                             *       500         *       520
chimeric KPNA7    : GQSALNIIEKHFGEEEDESQTLLSQVIDQDYEFIDYECLAKK
S.cerevisiae SRP1 : YEKAYKIIETYFGEEED-------------------------
```

```
                          *        20         *        40         *        60         *        80
Homo_sapiens CSE1L : MELSDANLQTLTEYLKKTLDPDPAIRRPAEKFLESVEGNQNYPLLLLTLLE-KSQDNVIKVCASVTFKNYIKRNWRIVED
S.cerevisiae CSE1  : -----SDLETVAKFLAESVI--ASTAKTSERNLRQLETQDGFGLTLLHVIASTNLPLSTRLAGALFFKNFIKRKWVDENG

                          *       100         *       120         *       140         *       160
Homo_sapiens CSE1L : EPNKICEADRVAIKANIVHLMLSSPEQIQKQLSDAISIIGREDFPQKWPDLLTEMVNRFQSGDFHVINGVLRTAHSLFKR
S.cerevisiae CSE1  : -NHLLPANNVELIKKEIVPLMISLPNNLQVQIGEAISSIADSDFPDRWPTLLSDLASRLSNDDMVTNKGVLTVAHSIFKR

                          *       180         *       200         *       220         *       240
Homo_sapiens CSE1L : YRHEFKSNELWTEIKLVLDAFALPLTNLFKATIELCSTHANDASALRILFSSLILISKLFYSLNFQDLPEFFEDNMETWM
S.cerevisiae CSE1  : WRPLFRSDELFLEIKLVLDVFTAPFLNLLKTVDEQITANENNKASLNILFDVLLVLIKLYYDFNCQDIPEFFEDNIQVGM

                          *       260         *       280         *       300         *       320
Homo_sapiens CSE1L : NNFHTLLTLDNKLLQT-DDEEEAGLLELLKSQICDNAALYAQKYDEEFQRYLPRFVTAIWNLLVTTGQEVKYDLLVSNAI
S.cerevisiae CSE1  : GIFHKYLSYSNPLLEDPDETEHASVLIKVKSSIQELVQLYTTRYEDVFGPMINEFIQITWNLLTSISNQPKYDILVSKSL

                          *       340         *       360         *       380         *       400
Homo_sapiens CSE1L : QFLASVCERPHYKNLFEDQNTLTSICEKVIVPNMEFRAADEEAFEDNSEEYIRRDLEGSDIDTRRRAACDLVRGLCKFFE
S.cerevisiae CSE1  : SFLTAVTRIPKYFEIFNNESAMNNITEQIILPNVTLREEDVELFEDDPIEYIRRDLEG--TDTRRRACTDFLKELKEKNE

                          *       420         *       440         *       460         *       480
Homo_sapiens CSE1L : GPVTGIFSGYVNSMLQEYAKNPSVNWKHKDAAIYLVTSLASKAQTQKHGITQANELVNLTEFFVNHILPDLKSANVNEFP
S.cerevisiae CSE1  : VLVTNIFLAHMKGFVDQYMSDPSKNWKFKDLYIYLFTALAINGNITNAGVSSTNNLLNVVDFFTKEIAPDLTSNN-IPHI

                          *       500         *       520         *       540         *       560
Homo_sapiens CSE1L : VLKADGIKYIMIFRNQVPKEHLLVSIPLLINHLQAESIVVHTYAAHALERLFTMRGPN--NATLFTAAEIAPFVEILLTN
S.cerevisiae CSE1  : ILRVDAIKYIYTFRNQLTKAQLIELMPILATFLQTDEYVVYTYAAITIEKILTIRESNTSPAFIFHKEDISNSTEILLKN

                          *       580         *       600         *       620         *       640
Homo_sapiens CSE1L : LFKALTLP-----GSSENEYIMKAIMRSFSLLQEAIIPYIPTLITQLTQKLLAVSKNPSKPHFNHYMFEAICLSIRITCK
S.cerevisiae CSE1  : LIALILKHGSSPEKLAENEFLMRSIFRVLQTSEDSIQPLFPQLLAQFIEIVTIMAKNPSNPRFTHYTFESIGAILNYTQ-

                          *       660         *       680         *       700         *       720
Homo_sapiens CSE1L : ANPAAVVNFEEALFLVFTEILQNDVQEFIPYVFQVMSLLLETHKNDIPSSYMALFPHLLQPVLWERTGNIPALVRLLQAF
S.cerevisiae CSE1  : --RQNLPLLVDSMMPTFLTVFSEDIQEFIPYVFQIIAFVVEQSA-TIPESIKPLAQPLLAPNVWELKGNIPAVTRLLKSF

                          *       740         *       760         *       780         *       800
Homo_sapiens CSE1L : LERGSNTIASAAADKIPGLLGVFQKLIASKANDHQGFYLLNSIIEHMPPESVDQYRKQIFILLFQRLQNSKTTKFIKSFL
S.cerevisiae CSE1  : IKTDSSIF-----PDLVPVLGIFQRLIASKAYEVHGFDLLEHIMLLIDMNRLRPYIKQIAVLLLQRLQNSKTERYVKKLT

                          *       820         *       840         *       860         *       880
Homo_sapiens CSE1L : VFINLYCIKYGALALQEIFDGIQPKMFGMVLEKIIIPEIQKVSGNVEKKICAVGITKLLTECPPMMDTEYTKLWTPLLQS
S.cerevisiae CSE1  : VFFGLISNKLGSDFLIHFIDEVQDGLFQQIWGNFIITTLPTIGNLLDRKIALIGVLNMVINGQFF-QSKYPTLISSTMNS

                          *       900         *       920         *       940         *       960
Homo_sapiens CSE1L : LIGLFELPEDDTIPDEEHFIDIEDTPGYQTAFSQLAFAGKKEHDPVGQMVN---NPKIHLAQSLHKLSTACPGRVPSMVS
S.cerevisiae CSE1  : IIETASSIANL--K-N-D-YV-EEISTFGSHFSKLVSISEKPFDPLPE-IDVNNGVRLYVAEALNKYNAISGNTFLNTIL

                          *       980
Homo_sapiens CSE1L : TSLNAEALQYLQGYLQAASVTLL
S.cerevisiae CSE1  : PQLTQENQVKLNQLLVG------
```

```
                                      *        20         *        40         *        60         *        80
Homo_sapiens RAN            : MAAQGEPQVQFKLVLVGDGGTGKTTFVKRHLTGEFEKKYVATLGVEVHPLVFHTNRGPIKFNVWDTAGQEKFGGLRDGYY
Canis_lupus_familiaris RAN : ------PQVQFKLVLVGDGGTGKTTFVKRHLTGEFEKKYVATLGVEVHPLVFHTNRGPIKFNVWDTAGQEKFGGLRDGYY


                                      *       100         *       120         *       140         *       160
Homo_sapiens RAN            : IQAQCAIIMFDVTSRVTYKNVPNWHRDLVRVCENIPIVLCGNKVDIKDRKVKAKSIVFHRKKNLQYYDISAKSNYNFEKP
Canis_lupus_familiaris RAN : IQAQCAIIMFDVTSRVTYKNVPNWHRDLVRVCENIPIVLCGNKVDIKDRKVKAKSIVFHRKKNLQYYDISAKSNYNFEKP


                                      *       180         *       200         *
Homo_sapiens RAN            : FLWLARKLIGDPNLEFVAMPALAPPEVVMDPALAAQYEHDLEVAQTTALPDEDDDL
Canis_lupus_familiaris RAN : FLWLARKLIGDPNLEF----------------------------------------
```

```
                             *        20         *        40         *        60         *        80
chimeric  KPNA1      : MTTPGKENFRLKSYKNKSLNPDEMRRRREEEGLQLRKQKREEQLFKRRNVATAEEETEEEVMSDGGFHEAQISNMEMAPG
Mus_musculus  KPNA2 : ----------------------------------------DEQMLKRRNVSSFPDDATSPLQEN---------RNNQGTV

                             *       100         *       120         *       140         *       160
chimeric  KPNA1      : GVITSDMIEMIFSKSPEQQLSATQKFRKLLSKEPNPPIDEVISTPGVVARFVEFLKRKENCTLQFESAWVLTNIASGNSL
Mus_musculus  KPNA2 : NWSVEDIVKGINSNNLESQLQATQAARKLLSREKQPPIDNIIRAG-LIPKFVSFLGKTDCSPIQFESAWALTNIASGTSE

                             *       180         *       200         *       220         *       240
chimeric  KPNA1      : QTRIVIQAGAVPIFIELLSSEFEDVQEQAVWALGNIAGDSTMCRDYVLDCNILPPLLQLFSKQNRLTMTR----NAVWAL
Mus_musculus  KPNA2 : QTKAVVDGGAIPAFISLLASPHAHISEQAVWALGNIAGDGSAFRDLVIKHGAIDPLLALLAVPDLSTLACGYLRNLTWTL

                             *       260         *       280         *       300         *       320
chimeric  KPNA1      : SNLCRGKSPPPEFAKVSPCLNVLSWLLFVSDTDVLADACWALSYLSDGPNDKIQAVIDAGVCRRLVELLMHNDYKVVSPA
Mus_musculus  KPNA2 : SNLCRNKNPAPPLDAVEQILPTLVRLLHHNDPEVLADSCWAISYLTDGPNERIEMVVKKGVVPQLVKLLGATELPIVTPA

                             *       340         *       360         *       380         *       400
chimeric  KPNA1      : LRAVGNIVTGDDIQTQVILNCSALQSLLHLLSSPKESIKKEACWTISNITAGNRAQIQTVIDANIFPALISILQTAEFRT
Mus_musculus  KPNA2 : LRAIGNIVTGTDEQTQKVIDAGALAVFPSLLTNPKTNIQKEATWTMSNITAGRQDQIQQVVNHGLVPFLVGVLSKADFKT

                             *       420         *       440         *       460         *       480
chimeric  KPNA1      : RKEAAWAITNATSGGSAEQIKYLVELGCIKPLCDLLTVMDSKIVQVALNGLENILRLGEQEAKRNGTGINPYCALIEEAY
Mus_musculus  KPNA2 : QKEAAWAITNYTSGGTVEQIVYLVHCGIIEPLMNLLSAKDTKIIQVILDAISNIFQAAEKLG-----ETEKLSIMIEECG

                             *       500         *       520         *       540
chimeric  KPNA1      : GLDKIEFLQSHENQEIYQKAFDLIEHYFGTEDEDSSIAPQVDLNQQQYIFQQCEAPMEGFQL
Mus_musculus  KPNA2 : GLDKIEALQRHENESVYKASLNLIEKYF----------------------------------
```

```
                                *         20         *         40         *         60         *         80
chimeric  KPNA5      : MDAMASPGKDNYRMKSYKNKALNPQEMRRRREEEGIQLRKQKREEQLFKRRNVSLPRNDESMLESPIQDPDISSTVPIPE
Mus_musculus  KPNA2 : -----------------------------------------DEQMLKRRNVSSFPDDATSPLQENRNN-------QGT

                                *        100         *        120         *        140         *        160
chimeric  KPNA5      : EEVVTTDMVQMIFSNNADQQLTATQKFRKLLSKEPNPPIDQVIQKPGVVQRFVKFLERNENCTLQFEAAWALTNIASGTF
Mus_musculus  KPNA2 : VNWSVEDIVKGINSNNLESQLQATQAARKLLSREKQPPIDNIIR-AGLIPKFVSFLGKTDCSPIQFESAWALTNIASGTS

                                *        180         *        200         *        220         *        240
chimeric  KPNA5      : LHTKVVIETGAVPIFIKLLNSEHEDVQEQAVWALGNIAGDNAECRDFVLNCEILPPLLELLTNSNRLTTTR----NAVWA
Mus_musculus  KPNA2 : EQTKAVVDGGAIPAFISLLASPHAHISEQAVWALGNIAGDGSAFRDLVIKHGAIDPLLALLAVPDLSTLACGYLRNLTWT

                                *        260         *        280         *        300         *        320
chimeric  KPNA5      : LSNLCRGKNPPPNFSKVSPCLNVLSRLLFSSDPDVLADVCWALSYLSDGPNDKIQAVIDSGVCRRLVELLMHNDYKVVSP
Mus_musculus  KPNA2 : LSNLCRNKNPAPPLDAVEQILPTLVRLLHHNDPEVLADSCWAISYLTDGPNERIEMVVKKGVVPQLVKLLGATELPIVTP

                                *        340         *        360         *        380         *        400
chimeric  KPNA5      : ALRAVGNIVTGDDIQTQVILNCSALPCLLHLLSSPKESIRKEACWTVSNITAGNRAQIQAVIDANIFPVLIEILQKAEFR
Mus_musculus  KPNA2 : ALRAIGNIVTGTDEQTQKVIDAGALAVFPSLLTNPKTNIQKEATWTMSNITAGRQDQIQQVVNHGLVPFLVGVLSKADFK

                                *        420         *        440         *        460         *        480
chimeric  KPNA5      : TRKEAAWAITNATSGGTPEQIRYLVALGCIKPLCDLLTVMDSKIVQVALNGLENILRLGEQESKQNGIGINPYCALIEEA
Mus_musculus  KPNA2 : TQKEAAWAITNYTSGGTVEQIVYLVHCGIIEPLMNLLSAKDTKIIQVILDAISNIFQAAEKLG-----ETEKLSIMIEEC

                                *        500         *        520         *        540
chimeric  KPNA5      : YGLDKIEFLQSHENQEIYQKAFDLIEHYFGVEEDDPSIVPQVDENQQQFIFQQQEAPMDGFQL
Mus_musculus  KPNA2 : GGLDKIEALQRHENESVYKASLNLIEKYF----------------------------------
```

```
                            *        20         *        40         *        60         *        80
chimeric  KPNA6      : METMASPGKDNYRMKSYKNNALNPEEMRRRREEEGIQLRKQKREQQLFKRRNVELINEEAAMFDSLLMDSYVSSTTGESV
Mus_musculus  KPNA2 : -------------------------------------------DEQMLKRRNVSSFPDDATSPLQENRNNQGTVN----W

                            *       100         *       120         *       140         *       160
chimeric  KPNA6      : ITREMVEMLFSDDSDLQLATTQKFRKLLSKEPSPPIDEVINTPRVVDRFVEFLKRNENCTLQFEAAWALTNIASGTSQQT
Mus_musculus  KPNA2 : SVEDIVKGINSNNLESQLQATQAARKLLSREKQPPIDNIIRAG-LIPKFVSFLGKTDCSPIQFESAWALTNIASGTSEQT

                            *       180         *       200         *       220         *       240
chimeric  KPNA6      : KIVIEAGAVPIFIELLNSDFEDVQEQAVWALGNIAGDSSVCRDYVLNCSILNPLLTLLTKSTRLTMTR----NAVWALSN
Mus_musculus  KPNA2 : KAVVDGGAIPAFISLLASPHAHISEQAVWALGNIAGDGSAFRDLVIKHGAIDPLLALLAVPDLSTLACGYLRNLTWTLSN

                            *       260         *       280         *       300         *       320
chimeric  KPNA6      : LCRGKNPPPEFAKVSPCLPVLSRLLFSSDSDLLADACWALSYLSDGPNEKIQAVIDSGVCRRLVELLMHNDYKVASPALR
Mus_musculus  KPNA2 : LCRNKNPAPPLDAVEQILPTLVRLLHHNDPEVLADSCWAISYLTDGPNERIEMVVKKGVVPQLVKLLGATELPIVTPALR

                            *       340         *       360         *       380         *       400
chimeric  KPNA6      : AVGNIVTGDDIQTQVILNCSALPCLLHLLSSPKESIRKEACWTISNITAGNRAQIQAVIDANIFPVLIEILQKAEFRTRK
Mus_musculus  KPNA2 : AIGNIVTGTDEQTQKVIDAGALAVFPSLLTNPKTNIQKEATWTMSNITAGRQDQIQQVVNHGLVPFLVGVLSKADFKTQK

                            *       420         *       440         *       460         *       480
chimeric  KPNA6      : EAAWAITNATSGGTPEQIRYLVSLGCIKPLCDLLTVMDSKIVQVALNGLENILRLGEQEGKRSGSGVNPYCGLIEEAYGL
Mus_musculus  KPNA2 : EAAWAITNYTSGGTVEQIVYLVHCGIIEPLMNLLSAKDTKIIQVILDAISNIFQAAEKLG-----ETEKLSIMIEECGGL

                            *       500         *       520         *       540
chimeric  KPNA6      : DKIEFLQSHENQEIYQKAFDLIEHYFGVEDDDSSLAPQVDETQQQFIFQQPEAPMEGFQL
Mus_musculus  KPNA2 : DKIEALQRHENESVYKASLNLIEKYF----------------------------------
```

```
                              *        20         *        40         *        60         *        80
chimeric  KPNA3      : MAENPGLENHRIKSFKNKGRDVETMRRHRNEVTVELRKNKRDEHLLKKRNVPQEESLEDSDVDADFKAQNVT---LEAIL
Mus_musculus  KPNA2 : -----------------------------------------DEQMLKRRNVSSFPDDATSPLQENRNNQGTVNWSVEDIV

                              *       100         *       120         *       140         *       160
chimeric  KPNA3      : QNATSDNPVVQLSAVQAARKLLSSDRNPPIDDLIKSGILPILVKCLERDDNPSLQFEAAWALTNIASGTSAQTQAVVQSN
Mus_musculus  KPNA2 : KGINSNNLESQLQATQAARKLLSREKQPPIDNIIRAGLIPKFVSFLGKTDCSPIQFESAWALTNIASGTSEQTKAVVDGG

                              *       180         *       200         *       220         *       240
chimeric  KPNA3      : AVPLFLRLLRSPHQNVCEQAVWALGNIIGDGPQCRDYVISLGVVKPLLSFISP----SIPITFLRNVTWVIVNLCRNKDP
Mus_musculus  KPNA2 : AIPAFISLLASPHAHISEQAVWALGNIAGDGSAFRDLVIKHGAIDPLLALLAVPDLSTLACGYLRNLTWTLSNLCRNKNP

                              *       260         *       280         *       300         *       320
chimeric  KPNA3      : PPPMETVQEILPALCVLIYHTDINILVDTVWALSYLTDGGNEQIQMVIDSGVVPFLVPLLSHQEVKVQTAALRAVGNIVT
Mus_musculus  KPNA2 : APPLDAVEQILPTLVRLLHHNDPEVLADSCWAISYLTDGPNERIEMVVKKGVVPQLVKLLGATELPIVTPALRAIGNIVT

                              *       340         *       360         *       380         *       400
chimeric  KPNA3      : GTDEQTQVVLNCDVLSHFPNLLSHPKEKINKEAVWFLSNITAGNQQQVQAVIDAGLIPMIIHQLAKGDFGTQKEAAWAIS
Mus_musculus  KPNA2 : GTDEQTQKVIDAGALAVFPSLLTNPKTNIQKEATWTMSNITAGRQDQIQQVVNHGLVPFLVGVLSKADFKTQKEAAWAIT

                              *       420         *       440         *       460         *       480
chimeric  KPNA3      : NLTISGRKDQVEYLVQQNVIPPFCNLLSVKDSQVVQVVLDGLKNILIMAG--DEASTIAEIIEECGGLEKIEVLQQHENE
Mus_musculus  KPNA2 : NYTSGGTVEQIVYLVHCGIIEPLMNLLSAKDTKIIQVILDAISNIFQAAEKLGETEKLSIMIEECGGLDKIEALQRHENE

                              *       500         *       520         *
chimeric  KPNA3      : DIYKLAFEIIDQYFSGDDIDEDPCLIPEATQGGTYNFDPTANLQTKEFNF
Mus_musculus  KPNA2 : SVYKASLNLIEKYF------------------------------------
```

```
                               *        20         *        40         *        60         *        80
chimeric  KPNA4      : MADNEKLDNQRLKNFKNKGRDLETMRRQRNEVVVELRKNKRDEHLLKRRNVPHEDICEDSDIDGDYRVQNTS---LEAIV
Mus_musculus  KPNA2 : -----------------------------------------DEQMLKRRNVSSFPDDATSPLQENRNNQGTVNWSVEDIV

                               *       100         *       120         *       140         *       160
chimeric  KPNA4      : QNASSDNQGIQLSAVQAARKLLSSDRNPPIDDLIKSGILPILVHCLERDDNPSLQFEAAWALTNIASGTSEQTQAVVQSN
Mus_musculus  KPNA2 : KGINSNNLESQLQATQAARKLLSREKQPPIDNIIRAGLIPKFVSFLGKTDCSPIQFESAWALTNIASGTSEQTKAVVDGG

                               *       180         *       200         *       220         *       240
chimeric  KPNA4      : AVPLFLRLLHSPHQNVCEQAVWALGNIIGDGPQCRDYVISLGVVKPLLSFISP----SIPITFLRNVTWVMVNLCRHKDP
Mus_musculus  KPNA2 : AIPAFISLLASPHAHISEQAVWALGNIAGDGSAFRDLVIKHGAIDPLLALLAVPDLSTLACGYLRNLTWTLSNLCRNKNP

                               *       260         *       280         *       300         *       320
chimeric  KPNA4      : PPPMETIQEILPALCVLIHHTDVNILVDTVWALSYLTDAGNEQIQMVIDSGIVPHLVPLLSHQEVKVQTAALRAVGNIVT
Mus_musculus  KPNA2 : APPLDAVEQILPTLVRLLHHNDPEVLADSCWAISYLTDGPNERIEMVVKKGVVPQLVKLLGATELPIVTPALRAIGNIVT

                               *       340         *       360         *       380         *       400
chimeric  KPNA4      : GTDEQTQVVLNCDALSHFPALLTHPKEKINKEAVWFLSNITAGNQQQVQAVIDANLVPMIIHLLDKGDFGTQKEAAWAIS
Mus_musculus  KPNA2 : GTDEQTQKVIDAGALAVFPSLLTNPKTNIQKEATWTMSNITAGRQDQIQQVVNHGLVPFLVGVLSKADFKTQKEAAWAIT

                               *       420         *       440         *       460         *       480
chimeric  KPNA4      : NLTISGRKDQVAYLIQQNVIPPFCNLLTVKDAQVVQVVLDGLSNILKMAEDEAETIGN--LIEECGGLEKIEQLQNHENE
Mus_musculus  KPNA2 : NYTSGGTVEQIVYLVHCGIIEPLMNLLSAKDTKIIQVILDAISNIFQAAEKLGETEKLSIMIEECGGLDKIEALQRHENE

                               *       500         *       520         *
chimeric  KPNA4      : DIYKLAYEIIDQFFSSDDIDEDPSLVPEAIQGGTFGFNSSANVPTEGFQF
Mus_musculus  KPNA2 : SVYKASLNLIEKYF------------------------------------
```

```
                              *        20         *        40         *        60         *        80
chimeric  KPNA2     : MSTNENANPARLNRFKNKGKDSTEMRRRRIEVNVELRKAKKDDQMLKRRNVSSFPDDATSPLQENRNNQGTVNWSVDDIV
Mus_musculus  KPNA2 : -----------------------------------------DEQMLKRRNVSSFPDDATSPLQENRNNQGTVNWSVEDIV

                              *       100         *       120         *       140         *       160
chimeric  KPNA2     : KGINSSNVENQLQATQAARKLLSREKQPPIDNIIRAGLIPKFVSFLGRTDCSPIQFESAWALTNIASGTSEQTKAVVDGG
Mus_musculus  KPNA2 : KGINSNNLESQLQATQAARKLLSREKQPPIDNIIRAGLIPKFVSFLGKTDCSPIQFESAWALTNIASGTSEQTKAVVDGG

                              *       180         *       200         *       220         *       240
chimeric  KPNA2     : AIPAFISLLASPHAHISEQAVWALGNIAGDGSVFRDLVIKYGAVDPLLALLAVPDMSSLACGYLRNLTWTLSNLCRNKNP
Mus_musculus  KPNA2 : AIPAFISLLASPHAHISEQAVWALGNIAGDGSAFRDLVIKHGAIDPLLALLAVPDLSTLACGYLRNLTWTLSNLCRNKNP

                              *       260         *       280         *       300         *       320
chimeric  KPNA2     : APPIDAVEQILPTLVRLLHHDDPEVLADTCWAISYLTDGPNERIGMVVKTGVVPQLVKLLGASELPIVTPALRAIGNIVT
Mus_musculus  KPNA2 : APPLDAVEQILPTLVRLLHHNDPEVLADSCWAISYLTDGPNERIEMVVKKGVVPQLVKLLGATELPIVTPALRAIGNIVT

                              *       340         *       360         *       380         *       400
chimeric  KPNA2     : GTDEQTQVVIDAGALAVFPSLLTNPKTNIQKEATWTMSNITAGRQDQIQQVVNHGLVPFLVSVLSKADFKTQKEAVWAVT
Mus_musculus  KPNA2 : GTDEQTQKVIDAGALAVFPSLLTNPKTNIQKEATWTMSNITAGRQDQIQQVVNHGLVPFLVGVLSKADFKTQKEAAWAIT

                              *       420         *       440         *       460         *       480
chimeric  KPNA2     : NYTSGGTVEQIVYLVHCGIIEPLMNLLTAKDTKIILVILDAISNIFQAAEKLGETEKLSIMIEECGGLDKIEALQNHENE
Mus_musculus  KPNA2 : NYTSGGTVEQIVYLVHCGIIEPLMNLLSAKDTKIIQVILDAISNIFQAAEKLGETEKLSIMIEECGGLDKIEALQRHENE

                              *       500         *       520
chimeric  KPNA2     : SVYKASLSLIEKYFSVEEEEDQNVVPETTSEGYTFQVQDGAPGTFNF
Mus_musculus  KPNA2 : SVYKASLNLIEKYF---------------------------------
```

```
                                  *        20         *        40         *        60         *        80
chimeric  KPNA7      : MPTLDAPEERLRKFKYRGKDASMRRQQRIAVSLELRKAKKDEQALKRRNITSFSPDPSEKTAKG----VAVSLTLGEIIK
Mus_musculus  KPNA2 : ----------------------------------------DEQMLKRRNVSSFPDDATSPLQENRNNQGTVNWSVEDIVK

                                  *       100         *       120         *       140         *       160
chimeric  KPNA7      : GVNSSDPVLCFQATQTARKMLSQEKNPPLKLVIEAGLIPRMVEFLKSSLYPCLQFEAAWALTNIASGTSEQTRAVVEGGA
Mus_musculus  KPNA2 : GINSNNLESQLQATQAARKLLSREKQPPIDNIIRAGLIPKFVSFLGKTDCSPIQFESAWALTNIASGTSEQTKAVVDGGA

                                  *       180         *       200         *       220         *       240
chimeric  KPNA7      : IQPLIELLSSSNVAVCEQAVWALGNIAGDGPEFRDNVITSNAIPHLLALISP----TLPITFLRNITWTLSNLCRNKNPY
Mus_musculus  KPNA2 : IPAFISLLASPHAHISEQAVWALGNIAGDGSAFRDLVIKHGAIDPLLALLAVPDLSTLACGYLRNLTWTLSNLCRNKNPA

                                  *       260         *       280         *       300         *       320
chimeric  KPNA7      : PCDTAVKQILPALLHLLQHQDSEVLSDACWALSYLTDGSNKRIGQVVNTGVLPRLVVLMTSSELNVLTPSLRTVGNIVTG
Mus_musculus  KPNA2 : PPLDAVEQILPTLVRLLHHNDPEVLADSCWAISYLTDGPNERIEMVVKKGVVPQLVKLLGATELPIVTPALRAIGNIVTG

                                  *       340         *       360         *       380         *       400
chimeric  KPNA7      : TDEQTQMAIDAGMLNVLPQLLQHNKPSIQKEAAWALSNVAAGPCHHIQQLLAYDVLPPLVALLKNGEFKVQKEAVWMVAN
Mus_musculus  KPNA2 : TDEQTQKVIDAGALAVFPSLLTNPKTNIQKEATWTMSNITAGRQDQIQQVVNHGLVPFLVGVLSKADFKTQKEAAWAITN

                                  *       420         *       440         *       460         *       480
chimeric  KPNA7      : FATGATMDQLIQLVHSGVLEPLVNLLTAPDVKIVLIILDVISCILQAAEKRSEKENLCLLIEELGGIDRIEALQLHENRQ
Mus_musculus  KPNA2 : YTSGGTVEQIVYLVHCGIIEPLMNLLSAKDTKIIQVILDAISNIFQAAEKLGETEKLSIMIEECGGLDKIEALQRHENES

                                  *       500         *       520
chimeric  KPNA7      : IGQSALNIIEKHFGEEEDESQTLLSQVIDQDYEFIDYECLAKK
Mus_musculus  KPNA2 : VYKASLNLIEKYF------------------------------
```
